# Speciation in Nearctic oak gall wasps is frequently correlated with changes in host plant, host organ, or both

**DOI:** 10.1101/2022.02.11.480154

**Authors:** Anna K.G. Ward, Robin K. Bagley, Scott P. Egan, Glen Ray Hood, James R. Ott, Kirsten M. Prior, Sofia I. Sheikh, Kelly L. Weinersmith, Linyi Zhang, Y. Miles Zhang, Andrew A. Forbes

## Abstract

Quantifying the frequency of shifts to new host plants within diverse clades of specialist herbivorous insects is critically important to understand whether and how host shifts contribute to the origin of species. Oak gall wasps (Hymenoptera: Cynipidae: Cynipini) comprise a tribe of ~1000 species of phytophagous insects that induce gall formation on various organs of trees in the family Fagacae, — primarily the oaks (genus *Quercus*; ~435 sp). The association of oak gall wasps with oaks is ancient (~50 my), and most oak species are galled by one or more gall wasp species. Despite the diversity of both gall wasp species and their plant associations, previous phylogenetic work has not identified a strong signal of host plant shifting among oak gall wasps. However, most emphasis has been on the Western Palearctic and not the Nearctic where both oaks and oak gall wasps are considerably more species rich and where oaks are more phylogenetically diverse. We collected 86 species of Nearctic oak gall wasps from 10 of the 14 major clades of Nearctic oaks and sequenced >1000 Ultra Conserved Elements (UCEs) and flanking sequences to infer wasp phylogenies. We assessed the relationships of Nearctic gall wasps to one another and, by leveraging previously published UCE data, to the Palearctic fauna. We then used phylogenies to infer historical patterns of shifts among host tree species and tree organs. Our results indicate that oak gall wasps have moved between the Palearctic and Nearctic at least four times, that some Palearctic clades have their proximate origin in the Nearctic, and that gall wasps have shifted within and between oak tree sections, subsections, and organs considerably more often than the analysis of previous data have suggested. Given that host shifts have been demonstrated to drive reproductive isolation between host-associated populations in other phytophagous insects, our analyses of Nearctic gall wasps suggest that host shifts are key drivers of speciation in this clade, especially in hotspots of oak diversity. Though formal assessment of this hypothesis requires further study, two putatively oligophagous gall wasp species in our dataset show signals of host-associated genetic differentiation unconfounded by geographic distance, suggestive of barriers to gene flow associated with the use of alternative host plants.

## Introduction

Estimates of the number of insect species range from 1.5 – 30 million, such that by most accounts they are the most species-rich class of animal (Erwin 1982, May 1990, Stork 1993; though see Larsen et al. 2017). The vast majority of insects are parasitic and use plants, fungi, or other animals as hosts for some part of their lifecycle (Price 1980, Strong 1988), a fundamental relationship that has inspired a long history of research into the underlying causes of insects’ evolutionary success (Mitter et al. 1988, Jaenike 1990, Nosil 2002, Forister et al. 2012). One particularly compelling question is how important are shifts to new hosts to the origin of diversity? On one hand, shifts to new hosts in some insect systems can drive the evolution of reproductive isolation between insect populations using the ancestral and derived hosts (Berlocher and Feder 2002, Drès and Mallet 2002, Craig et al. 2007, Matsubayashi et al. 2010, Forbes et al. 2017). On the other hand, if such host shifts occur only rarely, most parasitic insect diversity could be a result of co-cladogenesis between insects and their hosts. Such cocladogenesis is a pattern seen in some highly specialized plant-insect systems (Roderick 1997, Machado et al. 2001, Rønsted et al. 2005, McLeish et al. 2007), though even some of these show evidence for host-shifts (Wang et al. 2016, 2021). Inferring phylogenetic relationships for large clades of parasitic insects in the context of their host use patterns can therefore provide important clues about the relative importance of these two alternative mechanisms.

The oak gall wasps (Hymenoptera: Cynipidae: Cynipini), are a species-rich clade of ~1000 phytophagous (plant-feeding) insects that primarily use *Quercus* (oak) species as hosts (Stone et al. 2002, Melika & Abrahamson 2002, Buffington et al. 2020; Melika et al. 2021a,b). Female gall wasps oviposit into meristematic tissue, inducing abnormal, but highly structured, outgrowths of tissue (galls), inside of which the larvae develop (Stone and Schönrogge 2003, Csóka et al. 2005, Martinson et al. 2021). Gall wasps can attack many parts (organs) of oaks, including leaves, stems, buds, flowers, petioles, roots, and acorns and many gall wasp species alternate between sexual (gamic) and asexual (agamic) generations (Stone et al. 2002), with each generation inducing different gall morphologies often on different organs on the same or different tree species (Pujade-Villar et al. 2001, Egan et al. 2018). These differences in gall morphology, location, and the morphology of the agamic and gamic adult gall wasps have in some instances led to different generations being classified as different species or even different genera (Lund et al. 1998, Pujade-Villar et al. 2001, Melika & Abrahamson 2002). Conversely, similarities in gall and adult wasp morphology have led to many genera describing para- or polyphyletic groups (Cooke 2018, Melika et al. 2021a). Several gall wasp species are also known from only one generation, likely because the alternate generation galls are inconspicuous or hidden, e.g., on roots or under bark (Weld 1959, Hood et al. 2018). The complexity of the life cycle and biology of gall wasps makes them fantastic systems for asking myriad ecological and evolutionary questions but has simultaneously complicated understanding of their evolution and phylogeny.

Oaks are also a species-rich and taxonomically complex group, though their phylogeny is better resolved than that of the gall wasps. There are ~435 oak species worldwide, currently organized into two subgenera (*Cerris* and *Quercus*; Hipp et al. 2020), each containing several sections further divided into subsections (Table 1). The split between subgenus *Cerris* and subgenus *Quercus* is estimated to be 40–50 Ma (McIver and Basinger 1999, Hipp et al. 2018), with subgenus *Cerris* confined entirely to the Palearctic and subgenus *Quercus* primarily in the Nearctic (Hipp et al. 2020). Only two relatively small clades of subgenus *Quercus* have dispersed back to Eurasia (the Palearctic roburoids and part of section *Ponticae*; Kremer and Hipp 2020, Manos and Hipp 2021). At the species level, oaks are considerably more diverse in the Nearctic (Kremer and Hipp 2020), where the two most species-rich sections (sect. *Quercus* [white oaks] and sect. *Lobatae* [red oaks]) broadly overlap geographically with one another and the ranges of three other smaller sections (*Virentes, Ponticae*, and *Protobalanus*; Denk et al. 2017; Hipp et al. 2018; Manos and Hipp 2021). Oak gall wasps are also most species-rich in the Nearctic, with an estimated 700 species (Krombein et al. 1979, Penzes et al. 2018; Melika et al. 2021b).

**Table 1.** Classification of the subgenera, sections and subsections of genus *Quercus* following Manos and Hipp (2021). Some names in the “subsections” column are not currently recognized as true subsections, but nevertheless represent apparently monophyletic clades of oaks (Hipp et al. 2018, 2020, Manos and Hipp 2021). Oak clades not represented by any gall wasps in this study are indicated by an asterisk.

| Subgenus | Section | Subsection (clades) | Biogeography |
| --- | --- | --- | --- |
| Cerris | *Cerris* |  | Palearctic |
|  | *Ilex* |  | Palearctic |
|  | *Cyclobalanopsis* * |  | Palearctic |
| Quercus | *Lobatae* | *Agrifoliae* | Nearctic - Pacific coast |
|  |  | *Palustres* | Nearctic - Central USA |
|  |  | *Coccineae* | Nearctic - Central USA |
|  |  | *Phellos* | Nearctic - Central USA |
|  |  | “Texas red oaks”* | Nearctic - Texas |
|  |  | *Erythromexicana* | Nearctic - Mexico |
|  | *Protobalanus* |  | Nearctic - Southwest USA and Northwest Mexico |
|  | *Ponticae* * |  | Western Palearctic and CA |
|  | *Virentes* |  | Nearctic - Southeast USA and Mexico |
|  | *Quercus* | *Dumosae* | Nearctic - Pacific coast |
|  |  | *Prinoideae* | Nearctic |
|  |  | *Albae* | Nearctic - Central USA |
|  |  | *Roburoids* (incl. *Mesobalanus*) | Palearctic |
|  |  | *Stellatae* | Nearctic |
|  |  | *Polymorphae* * | Nearctic - Southwest USA to Guatemala |
|  |  | *Leucomexicana* | Nearctic - Southwest USA and Mexico |

If host shifting has contributed to the evolution of oak gall wasp diversity, then phylogenies of oak gall wasps should recover patterns of historical movements among oak subgenera, sections, subsections, and species. To date, the most ambitious phylogenetic study directly focused on the role of host shifting in oak gall wasp speciation sequenced two nuclear genes and a mitochondrial gene for 84 species of gall wasp in the Western Palearctic (Stone et al. 2009). Resultant trees showed that this assemblage of gall wasps sorted into clades organized by host tree associations with limited evidence of regular shifts among the oak subgenera and sections. However, a test for gall wasp host shifts limited to the Palearctic — where cynipid and oak diversity are well described but oak diversity itself is relatively depauperate (Manos and Hipp 2021) — ultimately may not have adequate power to resolve the overall role of host shifting in oak gall wasp diversity at a global scale. Further, new advances in global and continental oak phylogeny that have resolved relationships below the section level (Hipp et al. 2018, 2020, Manos and Hipp 2021) now allow for assessing the role of host shifting at progressively finer scales.

Just as shifts among host trees may be correlated with diversification of oak gall wasps, the fact that different species induce galls on very different plant organs also raises the question of whether shifts to new organs – even within the same tree species – could lead to the evolution of reproductive isolation and contribute to speciation. Indeed, in one clade of Palearctic *Andricus*, shifts among tree organs do appear to be common, even as tree host shifts appear rarer (Cook et al. 2002). With a shift to new organs on the same plant, one emergent reproductive barrier could be habitat isolation (Carroll and Boyd 1992, Berlocher and Feder 2002, Feder and Forbes 2007, Hood et al. 2015), wherein discrimination among plant organs isolates insect populations from one another. Another possibility is allochronic isolation, with populations of gall wasps becoming isolated temporally due to their induction of galls on organs produced by trees at different times during the year. Temporal (allochronic) isolation has been shown to be a strong contributor to incipient speciation in other insects (Feder and Filchak 1999, Joy and Crespi 2007, Inskeep et al. 2021, Mattsson et al. 2022). New phylogenies of Nearctic gall wasps reared from a diversity of galled organs across a diversity of oak species should allow for evaluation of whether shifts among different host organs are correlated with speciation events in gall wasps.

Another question important to understanding patterns and drivers of diversification of oak gall wasps, though one entangled with host association, is what is the phylogenetic relationship of the most species-rich gall wasp assemblage — the Nearctic gall wasps — to those in the Palearctic and in other biogeographic regions? The first serious attempts at understanding the phylogeny began with Kinsey (1920, 1936) who — prior to the advent of molecular phylogeny — used wasp morphology, gall characters, geography, host ranges, and other non-molecular characters to develop his hypothesis that the cradle of gall wasp diversity is present-day Mexico. Most modern studies have employed one to three loci to infer relationships, and in general results have conflicted with Kinsey’s view, putting the likely origin of oak galling in the Palearctic, where oaks also first originated (Stone et al. 2009, Andersen et al. 2021). However, these studies have often used no (Stone et al. 2009) or a limited number (Ronquist et al. 2015, Andersen et al. 2021) of Nearctic gall wasp samples, or have used Nearctic wasps but no Palearctic wasps (Melika et al. 2021a). One recent three-locus study that included both Nearctic and Palearctic samples maintains Palearctic wasps as basal to the Nearctic wasps, but with intercalated clades of Palearctic and Nearctic wasps (Nicholls et al. 2017), suggesting the possibility of multiple movements between continents throughout their evolutionary history. However, relationships among clades in this study were not sufficiently well resolved to assess this question directly.

Three recent studies have employed a multi-locus genome-level approach using Ultra Conserved Elements (UCEs, Faircloth et al. 2012) to address oak gall wasp phylogenetic relationships. First, Cooke (2018) used 17 Palearctic and 52 Nearctic wasp species in an attempt to resolve questions about the monophyly of several oak gall wasp genera. Next, Blaimer et al. (2020) sequenced some Cynipini as part of a larger study of Cynipoidea and found a likely point of origin for oak galling approximately ~50 MYA, roughly coincident with the split between subgenus *Cerris* and subgenus *Quercus* oaks (Hipp et al. 2018). Most recently, Brandão-Dias et al. (2022) added additional data harvested from published Cynipini genomes to expand the Blaimer et al. (2020) UCE gall wasp dataset. Though not the main focus of any of these studies, in all cases the resultant phylogenies appear to show the Palearctic fauna as being paraphyletic. Cooke’s (2018) study also showed a Nearctic oak gall wasp (*Protobalandricus spectabilis*) as being basal to all other oak gall wasps. An expanded Nearctic gall wasp phylogeny will aid in resolving these relationships.

Here, we report the largest UCE-based phylogenetic study of Nearctic oak gall wasps to date. We sequence UCEs for 86 species reared from galls collected from 24 species of oak across the continental United States. We paired these with existing UCE data from Palearctic and other Nearctic oak gall wasps (Blaimer et al. 2020, Brandão-Dias et al. 2022), such that our set spanned six of the eight oak sections and 11 of the 13 subsections of oak sections *Lobatae* and *Quercus*. Combining our new collections with existing Palearctic data allow us to re-contextualize relationships among the Nearctic and Palearctic Cynipini. With this accomplished, our careful records of the host oak species and organ from which each gall was collected allow us to ask how often gall wasps have shifted among oak tree sections, subsections, and organs. After considering oak gall wasp biogeography and host shifting at a Holarctic scale, we synthesize our findings and explore, in detail, two gall wasp clades (*Andricus quercuspetiolicola* and a clade of *Disholcaspis* wasps) that show evidence of host-associated genetic differentiation indicative of ongoing or recent divergence related to host plant shifts.

## Methods

### Collections

We collected oak galls from August 2015 – September 2019. Most collection sites were located in the eastern half of the United States, supplemented by smaller collections from the Southwest and the Pacific Coast (Supplemental Table 1). We reared insects directly from galls by placing gall collections of the same type and same collection date for each locality into deli cups with the base removed and replaced with a fine gauze to allow air flow. Most collections were stored in an incubator that rotated through four three-month artificial “seasons” of light:dark (LD), temperature, and relative humidity (RH) settings, as follows: Spring: 13:12 LD, 18°C day, 5°c night, 75% RH; Summer: 15:9 LD, 24°C day, 17°C night, 85% RH, Fall: 14:10 LD, 20°C day, 14°C night, 75% RH, Winter: 10:14 LD, 10°C day, 5°C night, 75 % RH. Some galls from warmer climates were kept at room temperature on a lab bench to avoid dramatic changes in temperature, including simulated winter conditions. All cups were checked daily for up to three years for emergent insects, which were placed into 95% ethanol in microcentrifuge tubes labeled with collection metadata and the emergence date. Gall wasps were initially identified to species based on the morphological characteristics of the galls from which they emerged, and by their tree host following a variety of resources (Weld and von Dalla Torre 1952, Weld 1957, 1959, 1960, Melika & Abrahamson 2002, Russo 2021, “Gallformers.org” 2021). Though gall morphology was our primary indicator of gall wasp species identity, the assumed wasp genus was verified morphologically from reared wasps against keys (Weld and von Dalla Torre 1952, Zimmerman 2018, Melika et al. 2021a) and all wasps selected for sequencing were photographed (see below) such that their identity could be confirmed as needed. We refer to some gall wasps reared from as-yet-undescribed galls by their description on gallformers.org as of December 2021, which includes the best-attempt genus name, followed by the host tree from which that gall was first recorded, and a description of its morphology (e.g., “Callirhytis_q_stellata_pentagonal_cluster”).

### DNA extractions

We extracted DNA from the whole adult bodies of 209 individuals representing 81 named species and five undescribed oak gall wasp species, and two *Synergus* individuals to use as outgroups (Supplemental Table 1). In cases where we had reared >1 gall wasp individuals of the same species from different tree hosts and/or different geographic regions, we included these replicates. We photographed one fore wing and the lateral side of each wasp body prior to destructive extractions (Supplemental File 1) and for many species deposited voucher specimens representing the same or similar collections into the Frost Entomological Museum at Pennsylvania State University or the National Museum of Natural History (Supplemental Table 1). For 38 individuals sampled before 2018, we extracted DNA using a Qiagen DNeasy kit (Redwood City, CA). For the remaining 173 samples, we used a CTAB/PCI approach modified from Chen et al. (2010) because we found this method increases both the quality and quantity of the DNA yield.

### Ultra-Conserved Element (UCE) sequencing

We prepared libraries for each individual using the Kapa Hyper Prep library preparation kit (Kapa Biosystems Inc., Wilmington, MA, USA) and enzymatic fragmentation of DNA. To ensure the correct distribution of fragment lengths for UCE sequencing (300–800 bp), we double size-selected fragments from each individual with AMpure Beads and verified fragment distributions using a Bioanalyzer. Libraries were then pooled and hybridized following the MyBaits protocol (ArborBiosciences, Ann Arbor, MI, USA) using the Hym v2P bait set (Branstetter et al. 2017). We confirmed UCE loci enrichment for pooled samples using relative qPCR. We then sequenced the entire pooled library on one lane of a NovaSeq6000 (Illumina, Inc. San Diego, CA) at the Iowa Institute of Human Genetics at the University of Iowa. We followed the Phyluce v1.7 pipeline (Faircloth 2016) to process our UCE loci. We first trimmed remaining adapters and primers using Trimmomatic (Bolger et al. 2014). We then assembled *de novo* contigs using SPAdes (Bankevich et al. 2012). Next, the UCE loci were aligned and internally trimmed using PHYLUCE (Faircloth 2016) and MAFFT v7 (Katoh and Standley 2013).

We generated 2,719–2,347,711 raw sequencing reads per taxon (median: 692,910) which we assembled with Spades to generate 1,645–120,122 contigs per taxon (median: 9,604). One *Disholcaspis quercusglobulus* sample was removed from the data set due to low capture of UCE loci (2,719 raw reads, 175 contigs, 91 UCE loci), probably the result of too little DNA being included in the final pooled library sample. Realized capture of the 2,590 target UCE loci ranged from a minimum of 472 to a maximum of 1607 loci. Further details on UCE capture and assembly can be found in Supplemental Table 3.

To investigate evolutionary relationships between Palearctic and Nearctic gall wasps, we added UCE loci from 16 additional taxa from Branstetter et al. (2017), Blaimer et al. (2020), and Brandão-Dias et al. (2022). We did not use UCE data from Cooke (2018) because these were not publicly available. We also bioinformatically extracted UCE loci from 8 existing genomes following the protocol in Phyluce 1.7 (Faircloth 2016) again using the Hym v2P baitset. We intentionally selected gall wasp species from other geographical regions and/or not previously represented in our dataset. We also only used samples with >300 UCE loci recovered, a minimum we set arbitrarily but with the goal of maximizing data completeness. Samples added from other sources can be found in Supplemental Table 2.

### Phylogenetic inference

We constructed phylogenetic trees for the entire combined dataset (235 individuals, representing 106 named species, 5 unknown species) using both a concatenated matrix and a multiple species coalescence approach, comparing resultant tree topologies across methods. For the concatenated matrix approach we assembled three different data matrices comprised only of the UCE loci found in at least 50%, 75%, and 90% of all individuals. These three matrices resulted in 1325, 1077, and 464 loci, respectively. Total alignment lengths of sequences from the three matrices were 442,593 bp, 381,400 bp, and 184,665 bp. We used AMAS (Borowiec 2016) to determine alignment length, percent missing data, AT and GC content, number and proportion of variable sites, and the number and proportion of parsimony informative sites (Supplemental Table 4). We partitioned the matrix with SWSC-EN (Tagliacollo and Lanfear 2018) and PartitionFinder2 (Lanfear et al. 2017) using the “rclusterf” algorithm (Lanfear et al. 2014) to test three models: GTR, GTR+G, GTR+I+G. We then generated Maximum Likelihood (ML) trees with IQ-TREE2 (Minh et al. 2020) for each of the concatenated matrices with 1000 Ultrafast bootstraps (Hoang et al. 2017) and 1000 SH-likelihood ratio tests (Guindon et al. 2010). For the multiple species coalescence approach, we generated gene trees for each UCE locus with IQ-TREE with ModelFinder (Kalyaanamoorthy et al. 2017) to determine the best model of substitution for each locus. We then used ASTRAL-III (Zhang et al. 2018) to estimate a species tree from the combined pool of gene trees.

In addition to a tree containing both Palearctic and Nearctic gall wasps, we constructed a phylogeny consisting of only our new North American samples. Because we had used fresh specimens where other studies had included primarily museum samples, this “North American only” dataset has the advantage of having many more UCE loci recovered per each individual, increasing the amount of phylogenetically informative sites. We again generated phylogenies using concatenated matrices and a multiple species coalescence approach as previously described.

### Biogeography and host associations

To investigate how tree host and biogeographic history may have influenced gall wasp evolution, we mapped various levels of oak tree taxonomic organization onto our gall wasp phylogenies. The genus *Quercus* is composed of two subgenera, *Cerris* and *Quercus*, each of which is further divided into sections (Hipp et al. 2018), and further still into several subsections (Manos and Hipp 2021; Table 1). Oaks in subgenus *Cerris* are only found in the Palearctic, while members of the three sections of the subgenus *Quercus*, – *Lobatae* (red oaks), *Virentes* (live oaks), and *Protobalanus* (golden oaks) – are almost exclusively restricted to the Nearctic. Section *Quercus* (white oaks) straddles the Palearctic and Nearctic, and section *Ponticae* (intermediate oaks) is found only on the Pacific coast of North America and on the eastern coast of the Black Sea (Manos and Hipp 2021). Within the North American oaks in subgenus *Quercus*, sections and subsections tend to be restricted to one of three biogeographic regions: Eastern North America, the Southwest (representing the northern tip of a Mexican and Central American oak flora), or the Pacific coast (California, north to British Columbia) (Hipp et al. 2018, Manos and Hipp 2021).

We estimated the ancestral host associations of gall wasps using R package Phytools (Revell 2012). We coded the gall wasp-associated traits based on host plant taxonomy (*Quercus, Lobatae, Virentes, Chrysolepis, Castanea*, and *Cerris*), and for the three Western Palearctic species that alternated between *Quercus* and *Cerris* we coded the alternate host plants as separate states. We pruned the 75% complete matrix tree down to a single specimen per taxa, and performed marginal reconstructions and selected the equal rates (ER) model based on AIC score. The resulting phylogenetic tree was built using the package ‘ape’ in R v4.1.1 (R Core Team 2021).

We also investigated evolutionary switches in the location of the gall on different host tree organs (e.g., bud, leaf, stem). Due to the cyclically parthenogenetic lifecycle of most gall wasps, the two generations often induce galls on different plant organs, such that one generation could shift to a new organ while the other generation did not. We combined our own observations from collections with accounts from the literature (relying especially on gallformers.org) to map gall locations for both sexual (gamic) and sexual (agamic) gall generations for each species in our North American phylogeny, which we then used to tally a minimum number of host organ shifts for both the sexual and asexual generations.

## Results

### Phylogeny and Biogeography of Cynipini

Both the multiple species coalescence approach and concatenated matrix approaches (Supplementary Figures 1–4) generated essentially the same topology (see below). We chose the concatenated matrix approach resulting from the 75% complete data matrix (1077 loci) as the basis for figures in this manuscript because it provided a middle ground between optimizing the amount of shared data among specimens and maximizing the total number of loci. Figure 1 shows the topology of the combined Palearctic and Nearctic tree (collapsed at the species level; see Supplementary Figure 2 for full tree). Neither the Palearctic nor Nearctic gall wasps were monophyletic in any of the inferred trees (Figure 1, Supplementary Figures 1–4). The Palearctic gall wasps formed four independent clades in all tree topologies, with the Cerris-associated *Pseudoneuroterus saliens* and *Plagiotrochus suberi* and the chestnut-galling *Dryocosmus kuriphilus* being basal to all other gall wasps (Figure 1). The remaining Palearctic gall wasps split into three clades within the larger North American fauna: a clade of *Andricus* wasps galling oaks of subgenus *Quercus* or alternating between oaks of subgenera *Cerris* and *Quercus*, a single *Andricus* species (*A. inflator*) on subgenus *Quercus*, and a clade that pairs *Neuroterus quercusbaccarum* with *Cynips divisa*, both on subgenus *Quercus* (Figure 1). This topology implies a minimum of four movements of gall wasps between the Palearctic and Nearctic: one presumed original colonization of the Nearctic from the Palearctic, and then three clades subsequently “returning” to the Palearctic, either at the same or at different times.

**Figure 1.**
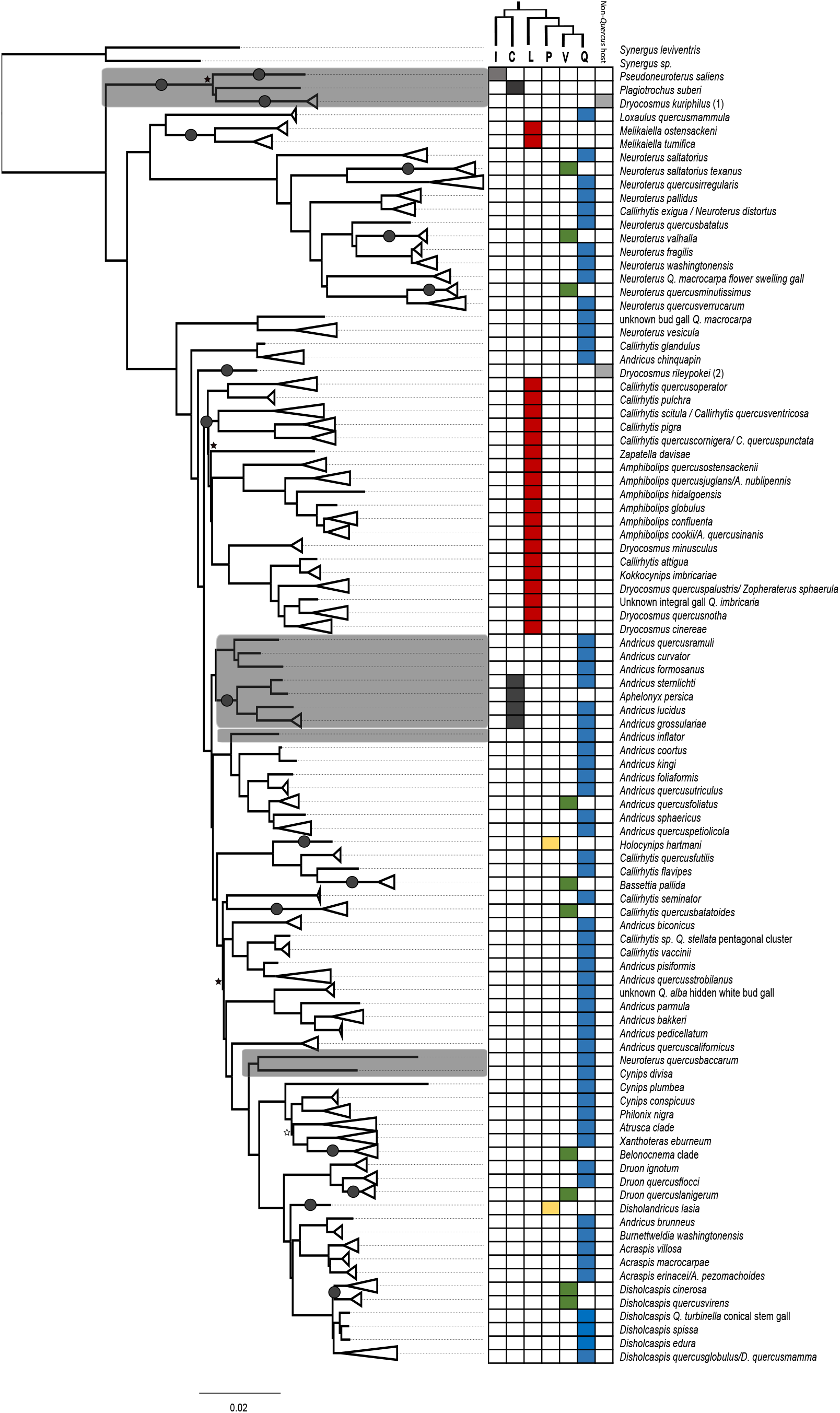
Phylogenetic tree of Nearctic and Palearctic oak gall wasps represented by the partitioned 75% complete data matrix. Clades have been collapsed to the species, or rarely genus, level (see Supplemental Figure 2 for the expanded tree). Tips with more than one species name indicate apparent polyphyly or the closing of lifecycles (see Supplemental File 2). All nodes have 100% bootstrap support, save for those indicated with black (<100%) or white (<80%) stars. Clades shaded in gray comprise Palearctic species. The grid to the right of the tree indicates the association of each gall with an oak section (or two sections, in the case of the alternating-host Palearctic *Andricus* clade). The cladogram above the right-hand grid illustrates the relationship among the oak sections: I - *Ilex*, C = *Cerris*, L = *Lobatae*, P = *Protobalanus*, V = *Virentes*, Q = *Quercus*. (1) = from galls on chestnut (*Castanea*); (2) = from galls on *Chrysolepis*. Dark circles, superficially resembling galls, on some branches of the phylogeny indicate changes in host tree section, assuming an ancestral association with oaks in subgenus *Quercus*, section *Quercus* as suggested by Ancestral State Reconstruction (Supplementary Figure 5).

As in several previous phylogenetic treatments of the Cynipini (Rokas et al. 2003, Ács et al. 2010, Ronquist et al. 2015, Cooke 2018, Andersen et al. 2021, Brandão-Dias et al. 2022), several gall wasp genera, including *Andricus, Callirhytis, Neuroterus*, and *Dryocosmus*, were found to be either para- or polyphyletic in our tree topologies (Figures 1–4). Of the genera represented in the tree by more than three species, only three genera: *Amphibolips, Acraspis*, and *Disholcaspis* maintain monophyly, the latter only because of a recent revision (Melika et al. 2021a) after Cooke (2018) had found *Disholcaspis* to be polyphyletic. Similarly, some species previously given different names were sufficiently similar such that they likely represent the alternate generations of the same species (Supplemental File 2; Supplementary trees 1–3).

**Figure 2.**
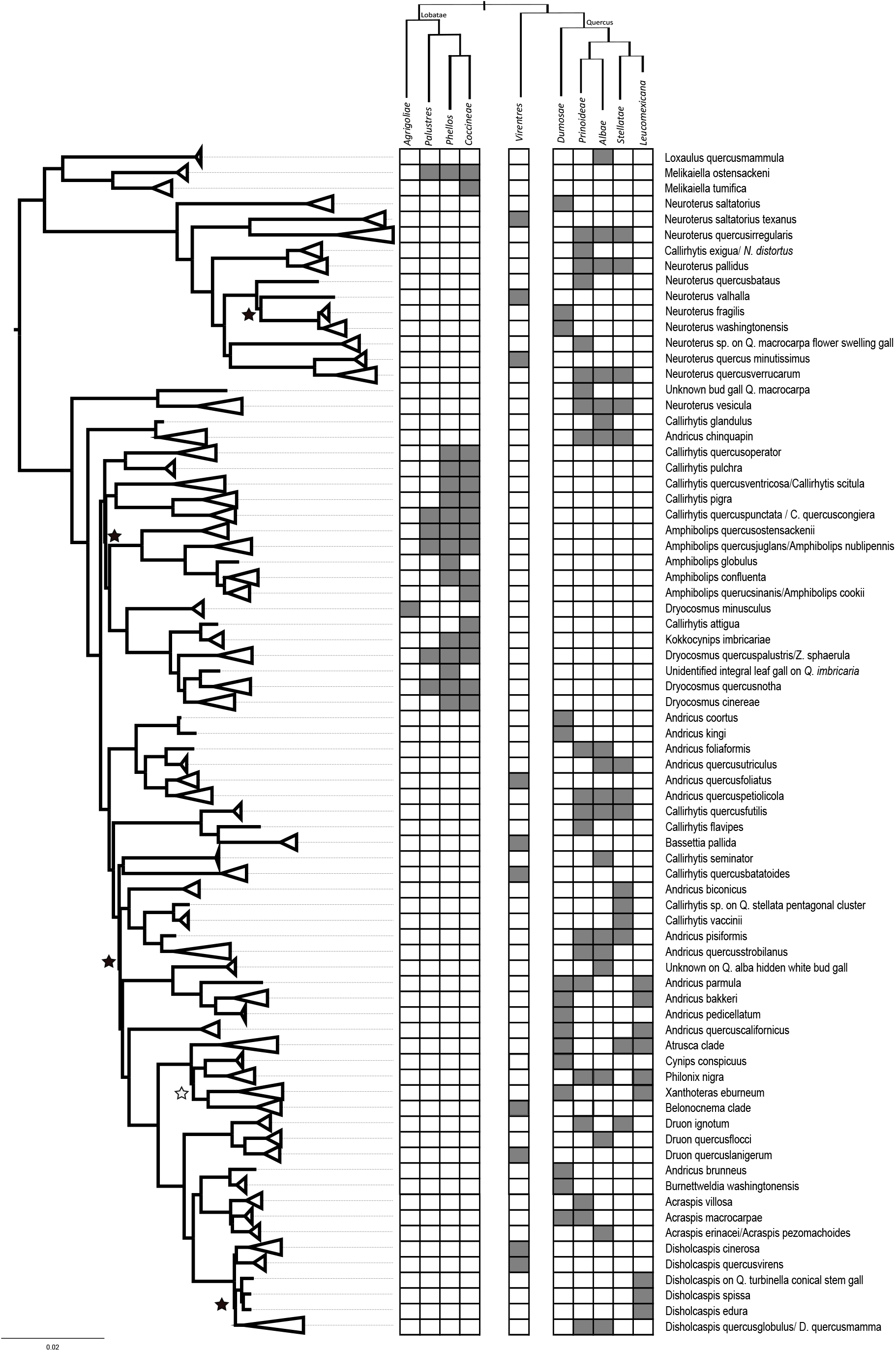
Phylogenetic representation of Nearctic gall wasps based on partitioned 75% completeness data matrix (see Supplemental Figure 7 for the expanded tree). Clades have been collapsed to the species or genus level as in Figure 1. Gray boxes in the table to the right of the tree represent known host associations – including those from which we collected insects and those from the literature – for each gall wasp species at the level of oak subsection. The cladogram above the table represents the evolutionary relationships among subsections within sections *Lobatae, Virentes*, and *Quercus* (section *Virentes* has no subsections). All nodes have bootstrap values and SH-likelihood support of 100, save for those indicated with black (<100%) or white (<80%) stars. While this tree implies that many oak gall wasp species induce galls on trees across >1 subsection, it may instead be the case that many taxa are species complexes, each species having a more restricted host range (see Discussion).

**Figure 3.**
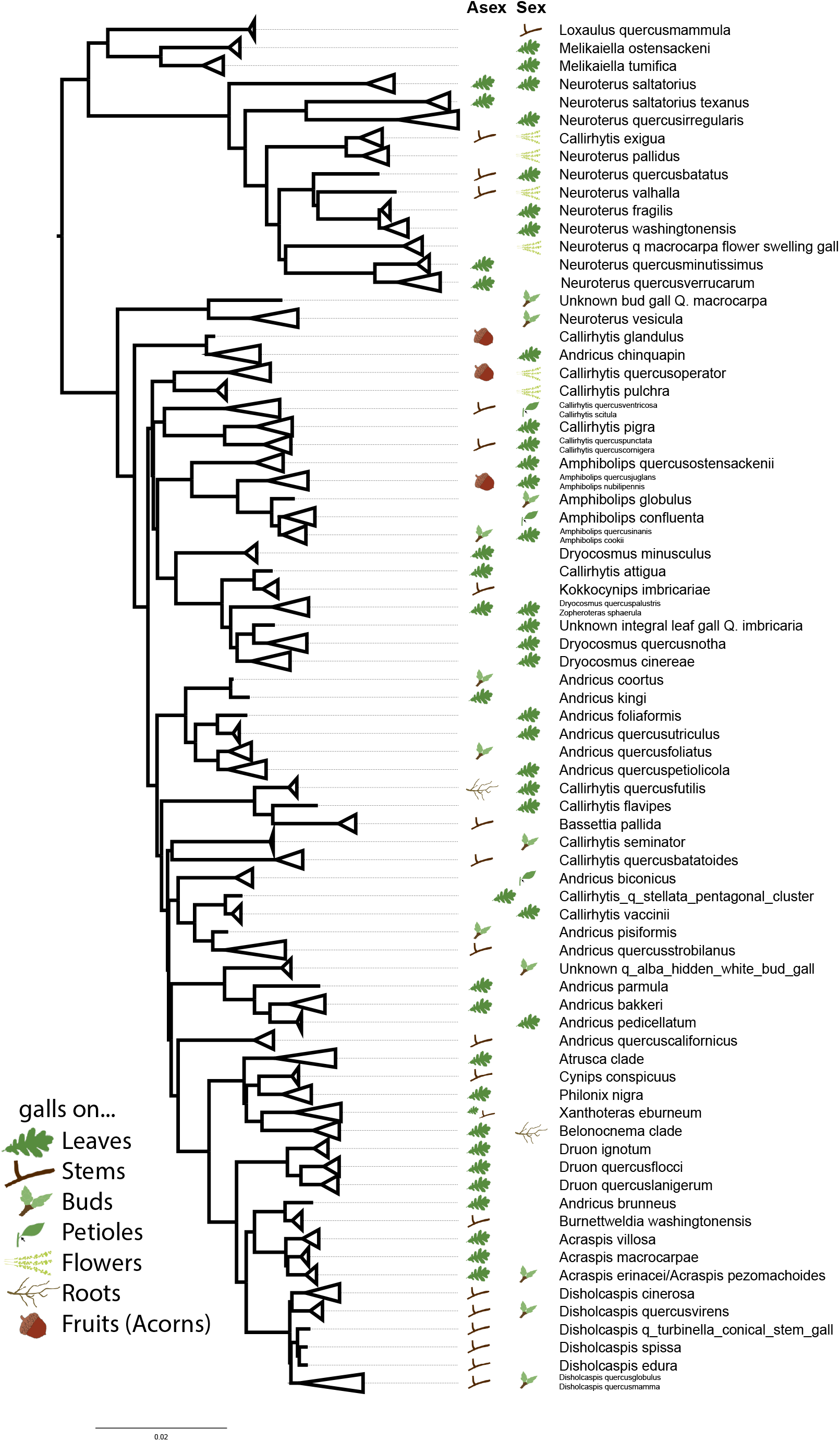
Known locations of galls on oaks for our Nearctic gall wasp set. Many cynipid gall wasps undergo cyclical parthenogenesis and have both asexual (“Asex”) and sexual (“Sex”) generations that gall different oak organs. Empty spaces indicate missing data and are in many cases due to the gall wasp having been described only from one generation. It is also possible that in some cases only one generation exists (e.g., *Andricus quercuscalifornicus*), though this is apparently rare in the Cynipini (Pujade-Villar et al. 1999; Stone et al. 2008).

**Figure 4.**
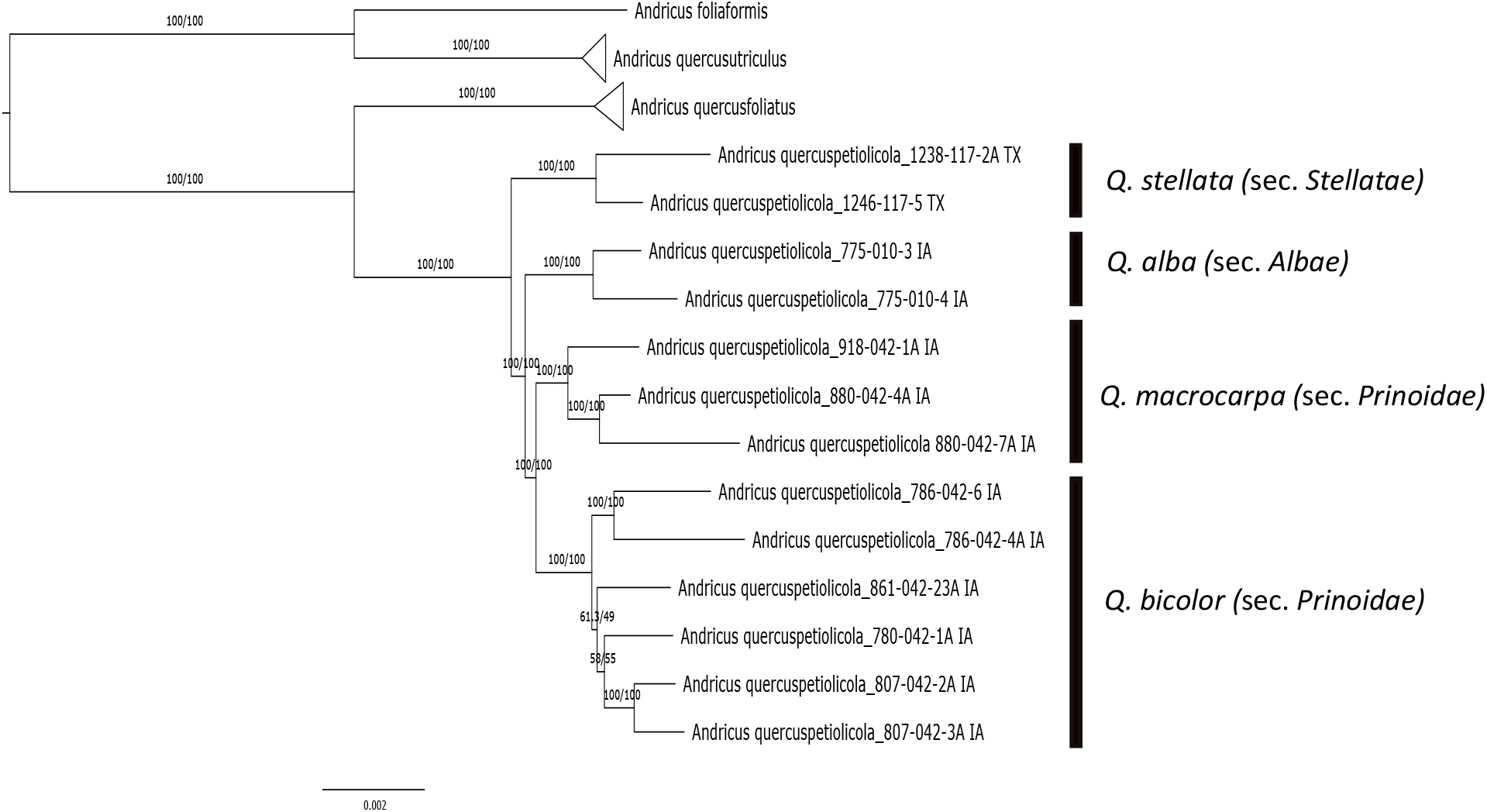
Host-associated genetic differentiation in *Andricus quercuspetiolicola*. A) phylogeny of A. *quercuspetiolicola* based on a concatenated 75% data matrix of UCE loci. Branch support are bootstrap values/ SH-likelihood values. Annotations indicate host tree species and the *Quercus* section from which we collected wasps.

Minor disagreements among trees using different approaches and data completeness matrices included: 1) the clade including *Callirhytis quercusscitula, C. quercusventricosa, C. pigra, C. quercuscornigera*, and *C. quercuspunctata* was sometimes placed as sister to the clade including *C. quercusoperator* and *C. pulchra* instead of as sister to the rest of *non-Callirhytis* gall wasps associated with section *Lobatae* oaks; 2) the clade including *Andricus biconicus*, an undescribed *Callirhytis* sp., *C. vaccinii, An. pisiformis*, and *An. quercusstrobilanus* was variably placed as sister to *C. seminator* and *C. quercusbataoides* or to a clade of *Andricus* species; 3) the *Cynips conspicuus* and *Philonix nigra* clade was sometimes sister to and sometimes outgroup to the clade consisting of the *Atrusca* gall wasps, *Xanthoteras eburneum*, and the *Belonocnema* gall wasps.

### Host association and host shifting

The ancestral state reconstruction recovered the oak subgenus *Quercus* as the ancestral host (Supplementary Figure 5. Based on this reconstruction, a minimum of 17 transitions among oaks of different sections are required to explain patterns in our tree (Figure 1; Supplementary Figure 4). Transitions between section *Lobatae* and section *Quercus* are estimated to have occurred at least twice, while those among the more closely related oak sections *Quercus* and *Virentes* have occurred at least nine times. The two gall wasp species, *Holocynips hartmani* and *Disholandricus lasia*, that were associated with oak section *Protobalanus*, were embedded in two different clades of wasps, each associated with sections *Quercus* and *Virentes* (Figure 1), implying at least two additional host shifts. Finally, a clade of Quercus-associated *Andricus* wasps has apparently shifted back to subgenus *Cerris*, section *Cerris*, but only for one of their two generations. Two other gall wasp lineages have moved from oak hosts onto more distantly related hosts (*Dryocosmus kuriphilus* onto *Castanea* and *Dryocosmus rileyipokei* onto *Chrysolepis*).

Host shifts between subsections of oaks are also apparent in the phylogeny of North American cynipids, though quantifying the incidence of shifts between subsections is complicated by the apparent sharing of hosts across several subsections by some gall wasp species (Figure 2; Supplementary Figures 6–8). While no gall wasp species in the North American assemblage apparently galls trees across multiple oak sections (Figure 1), both historical records and our own collections show that many gall wasp species induce galls on trees from >1 subsection (Figure 2). Still, the presence of gall wasps on subsections *Agrifoliae* and *Dumosae* in multiple clades cannot be explained without invoking inter-subsection host shifts.

The location at which galls are initiated and develop on host trees has also changed frequently across the Nearctic gall wasps (Figure 3). In spite of a considerable amount of missing data regarding gall location for both generations of many species, a strong signal emerges: gall wasps in this phylogeny show evidence of at least 29 changes in location of their galls on oak organs (13 in the asexual generation, 16 in the sexual generation). Within some clades of relatively recently diverged gall wasps, gall location appears to be more conserved (Figure 3). For instance, galls of asexual *Disholcaspis* (where known) are always on stems, while the galls of the corresponding sexual generation (again, where known) for all species are on buds. Similarly, species in the *Acraspis, Atrusca, Belonocnema*, and *Druon* clades show no apparent variation in organ use (Melika and Abrahamson 2002, Zhang et al. 2021) (Figure 3).

## Discussion

Our oak gall wasp phylogenies lead to several new conclusions regarding the biogeographic history and evolution of this charismatic and speciose tribe of insects. While we find additional support for a Palearctic origin of oak galling, a primary finding of our study is that some Palearctic clades appear to have their proximate origin in the Nearctic. We also uncovered evidence that host shifting is much more common than previously estimated, particularly between tree species in different subsections and among different tree organs. We discuss the implications of these results for the study of gall wasps as well as deeper implications for the role of variation in host plant use in the diversification of gall-inducing insects.

### Oak gall wasp biogeography

Our phylogenies further crystallize a biogeographic history of oak gall wasps in relation to their ultimate origin. Oak subgenus *Cerris* (Palearctic only) split from subgenus *Quercus* (primarily Nearctic, with one clade in the Palearctic) in the early to mid-Eocene, an estimated 50 Ma (Hipp et al. 2018). Likewise, the diversification of Cynipini into many of its extant lineages is estimated at ~50 Ma (Blaimer et al. 2020). Together, these lines of evidence support the hypothesis of an ancestral split between *Cerris*- and Quercus-associated gall wasps, with the origin of these gall wasps almost certainly occurring in the Palearctic (as suggested by Stone et al. 2009) before oaks spread to the Nearctic. In our dataset, two of the three most basal oak gall wasps are the Palearctic endemic species *Plagiotrochus suberi* and *Pseudoneuroterus saliens*, which are associated with cork oak (Subgenus *Cerris;* section *Ilex*) and Turkey oak (*Cerris; Cerris*), respectively (the other wasp in the clade induces galls on chestnut trees). As the Palearctic is the only biogeographic realm where subsection *Cerris* oaks are endemic, a Palearctic origin for these oak gall wasps remains the most parsimonious hypothesis. One caveat to the inference drawn from our analysis is that our dataset did not include *Protobalandricus spectabilis*, a Nearctic oak gall wasp species associated with oaks in section *Protobalanus. Protobalandricus spectabilis* was inferred to be basal to all others by the phylogeny presented in Cooke (2018). However, the support for this relationship in Cooke (2018) was lower than for other inferred relationships. Moreover, two three-gene phylogenies (Nicholls et al. 2017; Andersen et al. 2021) have instead grouped *P. spectabilis* as being sister to the other Nearctic species and not basal to all oak gall wasps. Future work that includes this taxon is required to resolve its relationship with the other oak gall wasps.

In contrast to our phylogenies’ apparently confirmatory nature with respect to the ultimate origins of oak gall wasps, our evolutionary trees rewrite the proximate origins of many extant oak gall wasp lineages. While the present study and other previous genetically based phylogenies (Stone et al. 2009; Andersen et al. 2021) refute Kinsey’s (1936) hypothesis of the Nearctic region as the point-of-origin for the oak-gall wasp association, our phylogeny also shows three Palearctic clades embedded within the Nearctic gall wasps, with high support, which strongly suggests movements of oak gall wasps from the Nearctic back to the Palearctic. Hints of this biogeographic exchange can be seen in the UCE phylogenies of Cooke (2018), Blaimer et al. (2020) and Brandão-Dias et al. (2022), with Cooke’s (2018) phylogeny suggesting as many as five separate movements to the Palearctic. Thus, although gall wasps may not have originated in the Nearctic, this region must now be reconsidered as a cradle of most or all gall wasp diversity associated with Palearctic roburoid oaks (the only Palearctic clade of oaks in section *Quercus*). Importantly, this scenario matches current biogeographic hypotheses for the origins of the Palearctic roburoids, which clearly originated in the Americas before moving into Asia and Europe (likely bringing Nearctic gall wasps with them) (Manos and Hipp 2021). One caveat is that the Eastern Palearctic remains under sampled, with just one species in our dataset (*Andricus formosanus*, from Taiwan; Blaimer et al. 2020). While in our new combined Nearctic-Palearctic phylogeny this wasp was embedded in one of the eastern Palearctic clades, more sampling should be made from these and other geographic areas.

A second caveat with respect to the combined Nearctic-Palearctic dataset is that there was considerable difference in locus recovery between our new data from recently collected material and previously published UCEs. Because most representatives from the Palearctic region had 600 or fewer UCE loci represented (Supplemental Tables 2-3), one might be mildly cautious about the placement of these wasps in our trees. We note, however, that the Nearctic representatives from Blaimer et al. (2020) (e.g., *Amphibolips hidalgoensis*) resolve within clades with their presumed close relatives and conspecifics (Figures 1–2; Supplemental Figures 1–4; 6–8). Further, the topologies of our trees are consistent with those of Blaimer et al. (2020), and the general phylogenetic relationships among named taxa are similar to those recovered for the same species using a small number of Sanger-sequenced loci (Stone et al. 2009, Ács et al. 2010, Ronquist et al. 2015, Andersen et al. 2021). For these reasons, the lower numbers of UCE loci for some samples are not likely to have caused major systematic (pun intended) errors in phylogenetic placement of the Palearctic species.

### Gall wasp host shifting across oak sections and organs

Our data provide the clearest “big picture” assessment yet of the tribe-wide relationships between host shifts and diversity in oak gall wasps. Our results demonstrate that gall wasps have moved between evolutionarily distant clades of oak tree hosts including at least 17 shifts among oaks of different sections and some non-zero number of shifts among oaks of different subsections within the same section (see below). These numbers do not include the previously published shift from section *Cerris* back to section *Ilex* by *Plagiotrochus amenti* (not present in our tree; see Stone et al. 2009). Some movements between oak sections could reasonably be interpreted as co-cladogenesis, for instance the deepest split between the three basal Palearctic gall wasps and the primarily Nearctic clade could have been coincident with subgenus *Cerris* splitting from subgenus *Quercus*. Similarly, the divergence of oak sections *Cerris* and *Ilex* in the Palearctic may be responsible for the host differences between *P. saliens* and *P. suberi* (Stone et al. 2009). However other major changes in tree association (e.g., two independent transitions from section *Quercus* to section *Lobatae*, nine transitions from section *Quercus* to section *Virentes*, and two transitions onto non-oak hosts) cannot all be explained by cospeciation of gall wasps with oaks and require at least some host shifting to have occurred.

Our conclusions differ from those of previous work that focused exclusively on Palearctic gall wasps (Stone et al. 2009, Ács et al. 2010). These studies found that gall wasps sorted largely into clades by their association with oak subgenera (*Cerris, Quercus*, or alternating between the two), and showed only one apparent complete shift among sections within *Cerris* (Stone et al. 2009). Based on this apparent minimal historical movement among hosts, these previous studies concluded that host use is strongly conserved in oak gall wasps, at least at the level of oak section. Indeed, in these previous studies, near monophyly among gall wasps associated with different tree sections has been the most reasonable emergent hypothesis based on gall wasp sampling and resultant phylogenies. In fact, if our UCE tree had been restricted to only the taxa that overlapped with those represented in the phylogenies of Andersen et al. (2021) and Stone et al. (2009), they would have shown essentially the same pattern.

Our differing conclusions from studies of primarily Palearctic species about the prevalence of host tree shifts in gall wasps are a result of our increased sampling of gall wasps from the Nearctic. Approximately 65% of oak species are found in the Nearctic (Nixon 1997; Manos et al., 1999; Hipp et al. 2017), including most of the subsections of oaks in subgenus *Quercus*, section *Quercus* (oaks from only one of these subsections occur in the western Palearctic; Manos and Hipp 2021), the entirety of section *Virentes, Protobalanus*, and all four subsections of section *Lobatae* (no *Virentes* or *Lobatae* are endemic to the Palearctic) (Table 1). Conversely, the Western Palearctic, where most phylogenetic studies of cynipids have been conducted (Cook et al. 2002, Bailey et al. 2009, Stone et al. 2009), has just 29 endemic oak species (Govaerts and Frodin 1998). Oak gall wasp richness is also correspondingly higher in the Nearctic, amounting to (70–80%) of an estimated >1000 species globally (Melika & Abrahamson 2002, Buffington et al. 2020; Melika et al. 2021b). Our samples spanned most of the Nearctic oak sections and subsections, with the exception of some oak clades in Texas, Mexico and Central America. These particular clades of oaks appear to have diversified recently and rapidly (Manos et al. 1999; Manos and Hipp 2021), such that their associated gall wasps will be an exciting future addition to these phylogenies.

Host shifts are also common between tree organs. For example, for the three species clade consisting of *Amphibolips globulus, Am. confluenta*, and *Am. quercusinanis / Am. cookii*, host tree association appears to have been somewhat conserved. *Amphibolips globulus* only induces galls on subsection *Phellos* oaks and *Am. quercusinanis* only on subsection *Coccineae* oaks, but Am. *confluenta* overlaps with both host ranges. However, these three species have shifted in their use of tree organs: *Am. globulus* induces galls on buds, *Am. confluenta* on petioles and young leaves, and *Am. quercusinanis* on leaves and sometimes flowers (Figure 3). We find at least 29 such shifts between organs in our phylogeny (Figure 3), with many more likely present but obscured because the location of development for some galls is unknown. This missing data regarding alternative galling sites, added to our incomplete sampling of the North American gall wasps, prevents a more robust analysis of whether host organ shifts occur more often with or without shifts in tree host. However, at a coarse level, many host shifts appear to occur without changes in tree section (Figures 1, 3). This pattern echoes results from the Palearctic (Cook et al. 2002), suggesting that organ shifts may be relevant to lineage divergence even in the absence of host tree shifts. Shifts across different organs on the same host plant species on the surface may not seem sufficient to lead to reproductive isolation. However, temporal differences in when oak tree organs are suitable for gall induction may separate populations allochronically and create a powerful barrier to gene flow. Temporal isolation of this kind is known from many non-galling insect systems (Malausa et al. 2005, Forbes et al. 2009, Powell et al. 2014, Boumans et al. 2017, Inskeep et al. 2021), some gall systems (Craig et al. 1993, Hood et al. 2019, Zhang et al. 2019) and even some instances where insects use the same host plants (Joy and Crespi 2007, Hippee et al. 2016, 2021).

Though the opportunities for host shifts among oaks is much higher in the Nearctic due to the higher oak diversity, our finding of frequent host shifting was by no means a foregone conclusion; differences in oak chemistry and other life history and defensive traits might still have restricted colonization of highly divergent oaks. Indeed, it is clearly rarer for gall wasps to move across subgenera than it is for them to shift sections (Figure 1). So, while the larger diversity of oaks and gall wasps in the Nearctic may simply mean that there are more opportunities for host shifts, it may also or instead be that some sections in the Nearctic are more closely related to one another than are the oak sections in the Palearctic, thereby making host shifting easier (e.g., *Quercus* and *Virentes* are sister sections and have the largest number of gall wasp shifts between them). On the other hand, similarities in oak defensive chemistry and other plant traits appear to be more related to biogeography and climate than phylogeny (Pearse and Hipp 2012, Moreira et al. 2018), so higher rates of gall wasp host shifting may really be a function of having a greater diversity of tree species in the same location.

### Host shifts across oak subsections and species

We do not directly quantify the number of host tree shifts at the levels of oak subsection or species as describing host plant ranges for gall wasps is not straightforward. For example, though no single gall wasp species induces galls of the same generation on oaks from different sections (some Palearctic *Andricus* alternate between Cerris and Quercus but show fidelity to oak section within each generation), many species induce galls on trees from different subsections within the same section (Figure 2). One interpretation of this pattern is that gall wasps may often be to some extent oligophagous, able to move freely between closely related tree species without strong barriers to gene flow evolving. This could be because trees in the same section are more often chemically similar (though see Pearse and Hipp 2012, Moreira et al. 2018) or have more similar developmental pathways, perhaps making it easier for the gall wasp species to shift to some novel hosts. If this is the case, the evolution of reproductive isolation between gall wasps may require host trees to be more different in one or more important dimensions (e.g., chemistry, phenology, development) than is commonly observed at the subsection level. A second possible explanation for the apparently broad host ranges of many gall wasp species is that the picture may be clouded by how we define gall wasp “species.” With most species names relying largely on the morphology of the adult wasp, it may be that historically the names of some gall wasps have “lumped” together closely related but reproductively isolated lineages as revealed by Zhang et al. (2021) for species of *Belonocnema*.

While we did not design this study for a detailed assessment of species boundaries needed to disentangle the two above possibilities, our data provides two case studies that hint at cryptic host-associated lineages within some gall wasp taxa. First, our samples included 13 individual *Andricus quercuspetiolicola* reared from five different oak tree species across both sympatric and allopatric sites. If we focus on this clade, we can test the hypothesis that An. *quercuspetiolicola* developing on the same hosts are more closely related to one another than sympatric wasps from different host plant species. Support for this hypothesis is shown in Figure 4: *An. quercuspetiolicola* gall wasps reared from *Quercus stellata, Q. alba*, and *Q. bicolor* sort into separate clades. Though the *Q. stellata* wasps were all from a site in Texas more than 1300 km from the others, wasps in the *Q. bicolor* clade were reared at sites within 10 miles of wasps galling *Q. alba* and *Q. macrocarpa*. The signal of closer genetic identity between allopatric populations developing on the same tree species than between sympatric populations developing on different tree species is a strong preliminary signal of host association acting as a reproductive isolating barrier.

Further support for the hypothesis that some apparently oligophagous gall wasps are actually complexes of more specialized lineages comes from the clade containing *Disholcaspis quercusmamma* and *D. quercusglobulus* (Figure 5a). Here again, wasps sort based on host tree species, independent of shared geography. *Disholcaspis quercusglobulus* reared from *Q. stellata* and *Q. alba* in Kentucky were less closely related to each other than to wasps reared from the same hosts in Missouri and Iowa, respectively. Similarly, *D. quercusmamma* collected from several sites in Iowa City, IA (3–7 km apart) on *Quercus macrocarpa* and *Q. bicolor* were less closely related to each other than to wasps from the same hosts collected in Spirit Lake, IA (352 km from Iowa City) and Reynoldsburg, OH (758 km from Iowa City). There has been some previous debate as to whether *D. quercusglobulus* and *D. quercusmamma* represent different species or simply induce galls of slightly different morphology on different oak species (McEwen et al. 2014). Our data now suggest that they are a complex of several putative species, each affiliated with different oak tree species.

**Figure 5.**
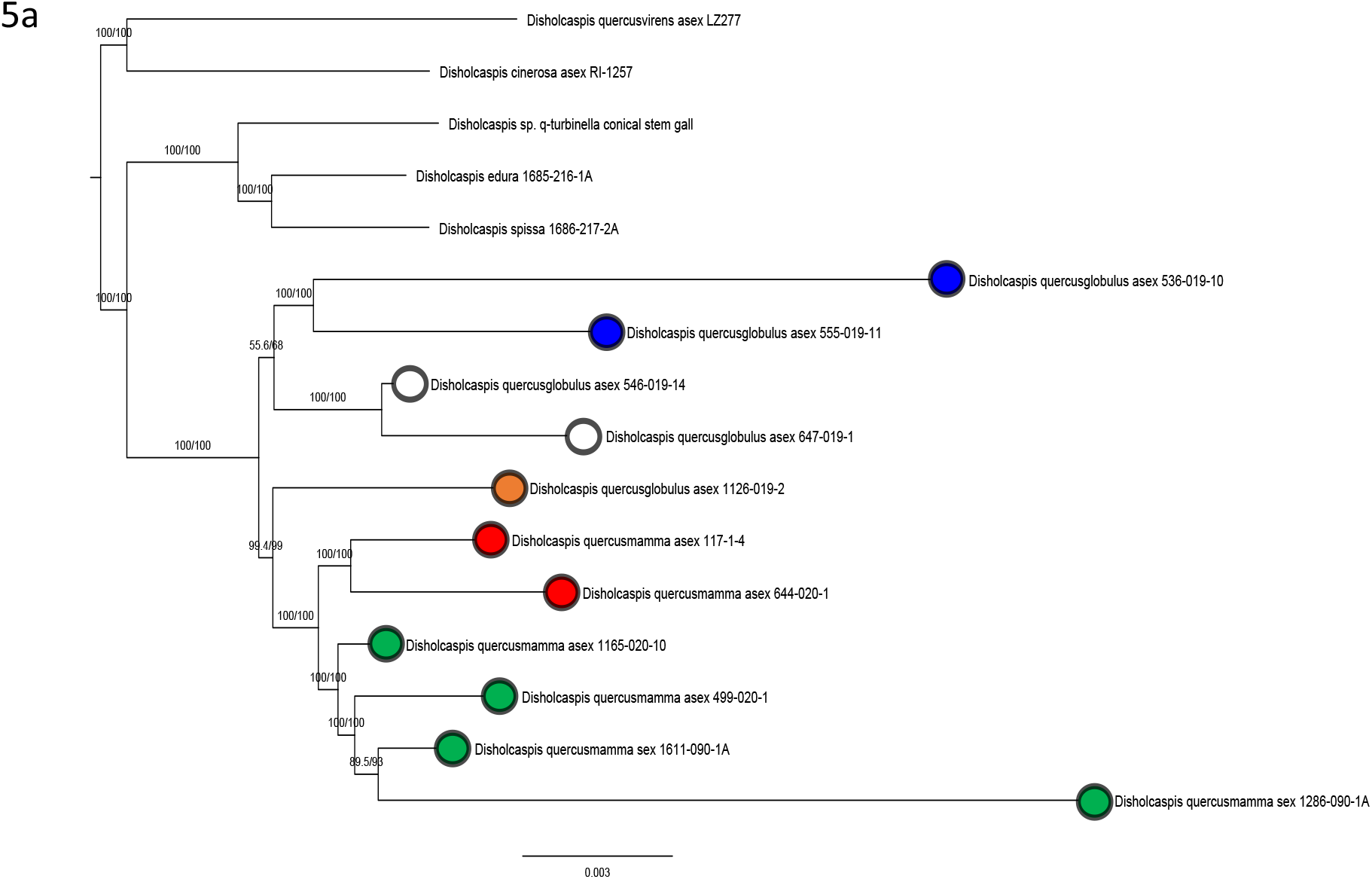
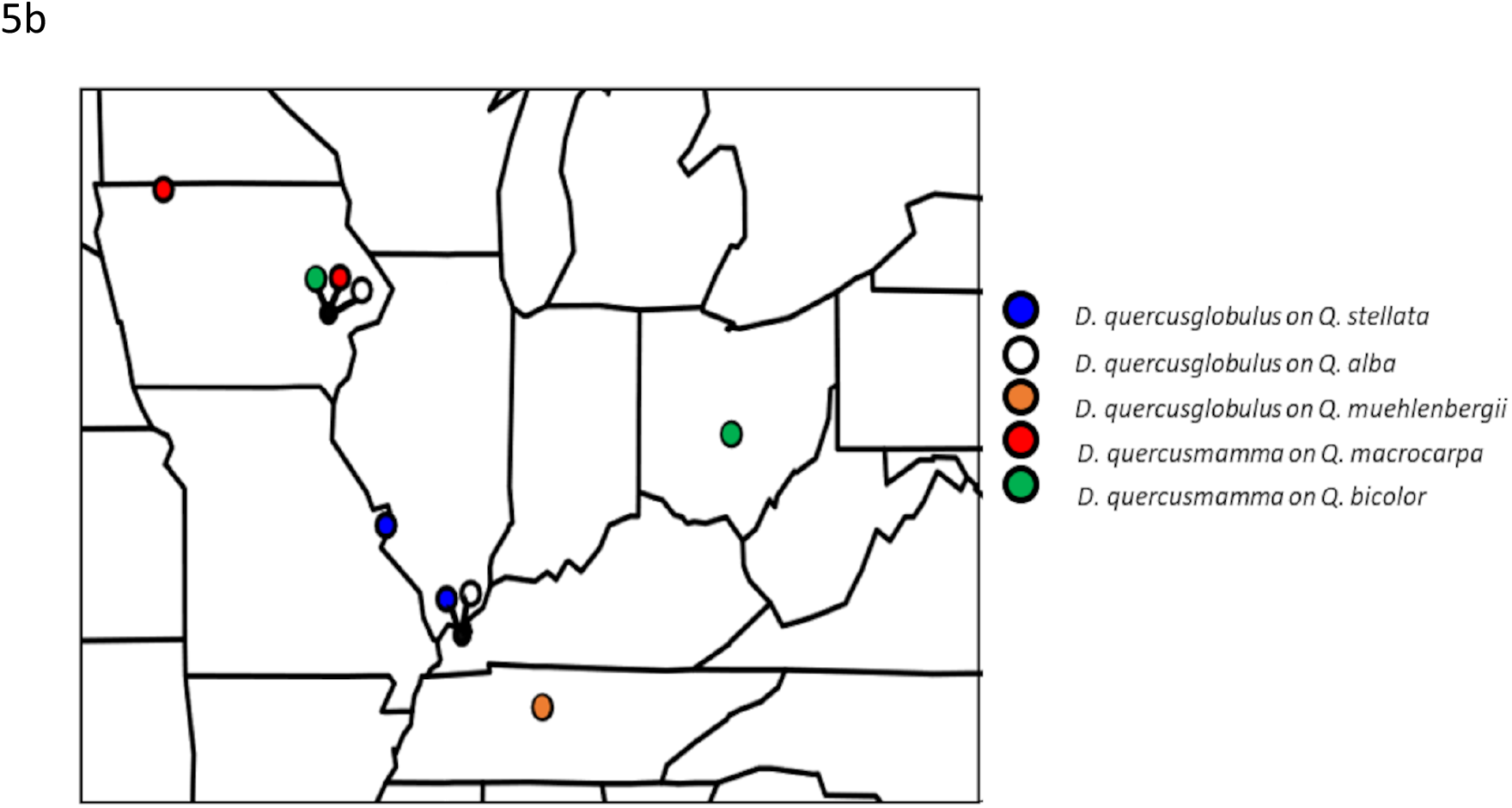
Host-associated genetic differentiation in *Disholcaspis quercusglobulus* and *Disholcaspis quercusmamma*. A) Phylogeny of *D. quercusglobulus* and *D. quercusmamma* based on a concatenated 75% data matrix of UCE loci, branch support are bootstrap/SH-likelihood values. Colored circles correspond to the gall wasp species ID and oak tree species from which each wasp was collected. B) Map of the upper Midwestern United States, showing location of collections.

If both *An. quercuspetiolicola* and the *Disholcaspis* gall wasps (Figures 4 & 5) are complexes of several different host-associated lineages, this further heightens the potential importance of host plant associations in understanding gall wasp diversity. Rather than many gall wasps being generalists with relatively broad host ranges (several tree species across >1 subsections as Figure 2 implies), there may instead be more cryptic species than have been previously recognized, each quite highly specialized and inducing galls on only one or a few closely related tree species. In this scenario, host shifts among closely related tree species may be as frequent as they are between different organs on the same tree (Cook et al. 2002; this paper), with shifts becoming less common – but still possible – as hosts become more taxonomically distant (subsections, sections, subgenera). We note as a caveat that in the ASTRAL tree (Supplemental Figure 4), *An. quercuspetiolicola and Disholcaspis* wasps did not sort as perfectly based on host tree species as in our concatenated matrix approach. This could be because these are recently diverged lineages, and resultant incomplete lineage sorting might confound resolution in multispecies coalescence approaches. Future work should use much denser sampling strategies from across larger geographic areas and multiple tree hosts to directly address host-associated differentiation in these and other Nearctic gall wasps.

Though we provide strong evidence for shifts of gall wasps among oak subgenera, sections, subsections, species, and different tree organs, our analyses do not directly address the question of whether these host shifts actually drive speciation in gall wasps. Directly confronting whether host shifts result in new insect diversity requires identifying populations that are only recently diverged and/or actively diverging (i.e., documented ongoing gene flow) or for which a host shift has been documented historically (Forbes et al. 2017). Lineages meeting these conditions can then provide the framework for the comparative and or experimental analysis of the relationships between variation in components of fitness in relation to host plant use and assessment of levels of both pre- and post-zygotic isolating mechanisms. Over a decade of work in one group of Nearctic gall wasps provides a template for addressing the role of host-plant-association in gall wasp speciation. The genus *Belonocnema* comprises three species that specialize on American live oaks (genus *Quercus;* subsection *Virentes;* Cavender-Bares et al. 2015). Two species of *Belonocnema, B. treatae* and *B.fossoria*, co-occur over a portion of their range across the coastal southeastern United States on the southern live oak, *Q virginiana*, and the sand live oak, *Q. geminata*, respectively (Zhang, M. et al. 2021), and represent two genetically diverged host-associated genetic clusters (Driscoe et al. 2019). Several other studies provide examples of how the host plant influences the evolution of reproductive isolation and promotes speciation in this genus. These two sympatric gall wasps have evolved multiple barriers to gene flow, including temporal isolation (Hood et al. 2019), sexual isolation (Egan et al. 2012a), habitat isolation (Egan et al. 2012b), immigrant inviability and reduced fecundity (Zhang et al. 2021a), and context-dependent reduced hybrid fitness (Zhang et al. 2021b). In total, these barriers combined to reduce gene flow by 95% to 99% between host-associated lineages (Hood et al. 2019; Zhang et al. 2021b). It would be fruitful to conduct similar studies in other Nearctic gall wasp genera to ascertain whether *Belonocnema* is unique or one of many similar systems.

If host shifts do prove to be a frequent driver of speciation in oak gall wasps, it will be exciting to then consider whether and how they affect the evolutionary histories of the many other insects associated with galls. Many parasitoids, inquilines, and hyperparasitoids have been reared from oak galls, with associates of some galls numbering more than 25 species (Joseph et al. 2011, Bird et al. 2013, Prior and Hellmann 2013, Forbes et al. 2016, Weinersmith et al. 2020). Recent work shows that many insect genera commonly associated with oak galls harbor species that specialize on one or a small subset of gall wasp hosts (Ward et al. 2019, 2020, Sheikh et al. 2022; Zhang et al. 2022). If gall wasps shift hosts, and are followed by concomitant shifts by natural enemies, the ecological dimensions relevant to reproductive isolation in the gall wasp might “cascade” during a host-shift and promote reproductive isolation among insect natural enemies (Blair et al. 2005, Stireman et al. 2006, Abrahamson and Blair 2008, Forbes et al. 2009, Hood et al. 2015). Indeed, Zhang et al. (2019) found that the *Synergus* inquilines associated with different gall formers on live oaks were staggered in their emergence times, with differences tracking development of galls on different tree hosts. Host shifting in gall wasps may therefore be relevant not only to their own diversity, but for biodiversity in their larger communities of predators and other gall associates. This promises to be an exciting area of future study.

## Supporting information

Supplemental figures 1-8. Phylogenetic trees

Supplemental file 2. Notes on Taxonomy

Supplemental file 1. images of gallwasps

## Acknowledgements

The authors thank Megan Blance, Amanda Driscoe, Robert Busbee, Charles Davis, Brad Foley, Elaine Hu, Rebecca Izen, Susan Lee, Kyle McElroy, Daniel McGarry, Shannon Meadley-Dunphy, Kevin Neely, Maurine Neiman, Shannon Pelini, Catherine Ruis, Monzer Shakally, Shih-An Shzu, Eric Tvedte, Joseph Verry, Heather Widmayer, and Caleb Wilson for help with gall collection and/or insect rearing. Adam Kranz helped identify some species based on their gall morphology. Steve Hendrix and Alaine Hippee provided valuable comments on an early draft of this manuscript. Funding for collections was provided to A.A.F. by the University of Iowa, to A.K.G.W. by the Center for Global and Regional Environmental Research, and to K.M.P by the National Geographic Society and Binghamton University. Funding for library preparation and sequencing was provided to A.K.G.W. via an EECG award from the American Genetics Association. Mention of trade names or commercial products in this publication is solely to provide specific information and does not imply recommendation or endorsement by the USDA. USDA is an equal opportunity provider and employer.

## Author Contributions

A.K.G.W., and A.A.F. designed the study. All authors made collections and/or reared animals. A.K.G.W and S.I.S. obtained the sequence data. A.K.G.W., S.I.S., Y.M.Z., and A.A.F. conducted the analyses, and A.K.G.W. and A.A.F. wrote the manuscript. All authors contributed to revisions and approved the final manuscript.

