## Supplemental figures 1-8. Phylogenetic trees for "Speciation in Nearctic oak gall wasps is frequently correlated with changes in host plant, host organ, or both"

Supplemental Figure 1. Maximum likelihood phylogeny based on the Nearctic and Palearctic gall wasp 50% complete data matrix. Support values show bootstrap value followed by SH-likelihood ratio. Tip labels include the abbreviated genus, species, lab code number, tree host section, tree host species, city and state of collection, where that information was known. For full sample information see Supplemental Tables 1 and 3.

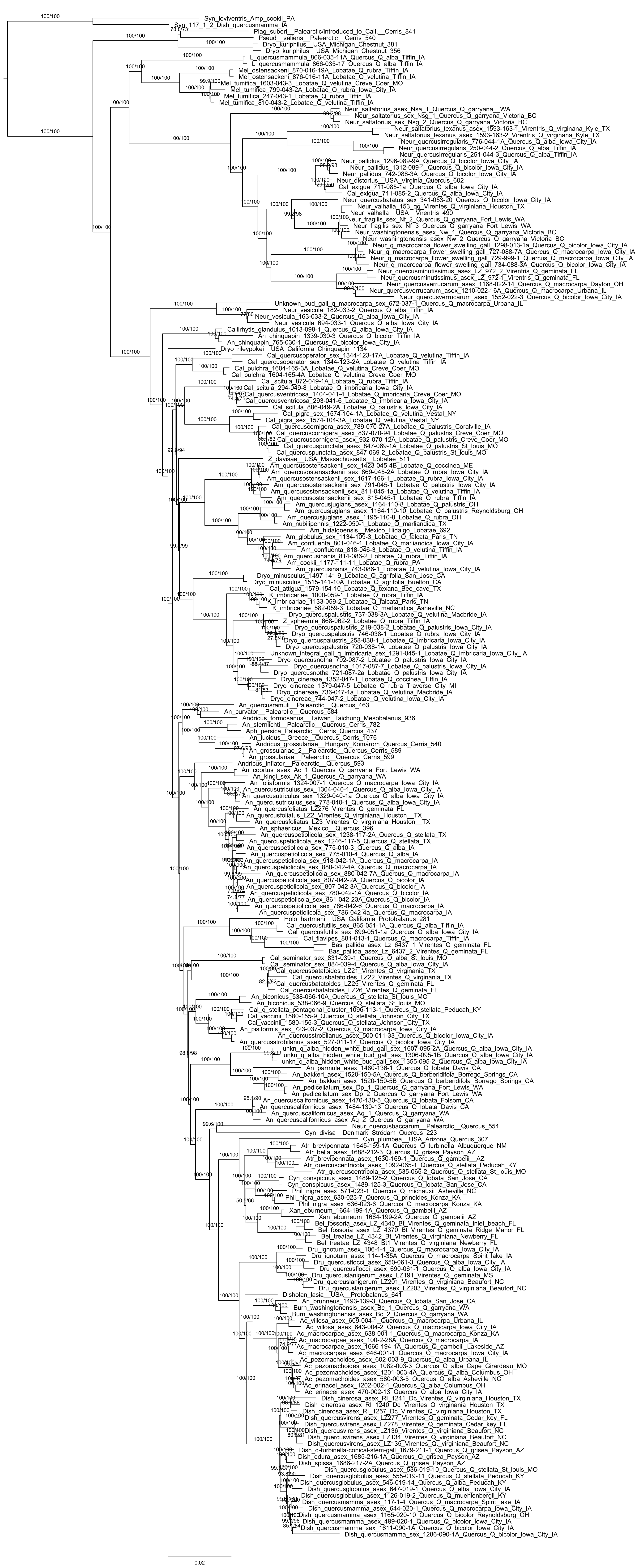

Supplemental Figure 2. Maximum likelihood phylogeny based on the Nearctic and Palearctic gall wasp 75% complete data matrix. Support values show bootstrap value followed by SH-likelihood ratio. Tip labels include the abbreviated genus, species, lab code number, tree host section, tree host species, city and state of collection, where that information was known. For full sample information see Supplemental Tables 1 and 3.

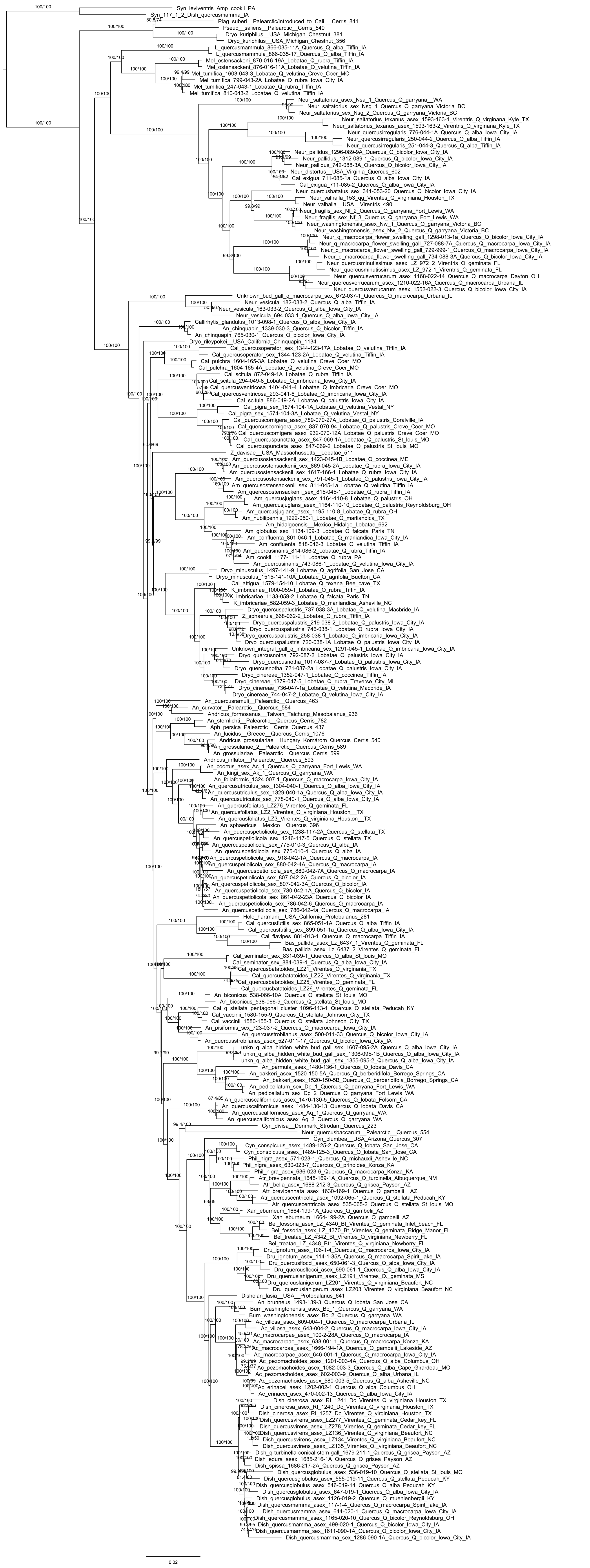

Supplemental Figure 3. Maximum likelihood phylogeny based on the Nearctic and Palearctic gall wasp 90% complete data matrix. Support values show bootstrap value followed by SH-likelihood ratio. Tip labels include the abbreviated genus, species, lab code number, tree host section, tree host species, city and state of collection, where that information was known. For full sample information see Supplemental Tables 1 and 3.

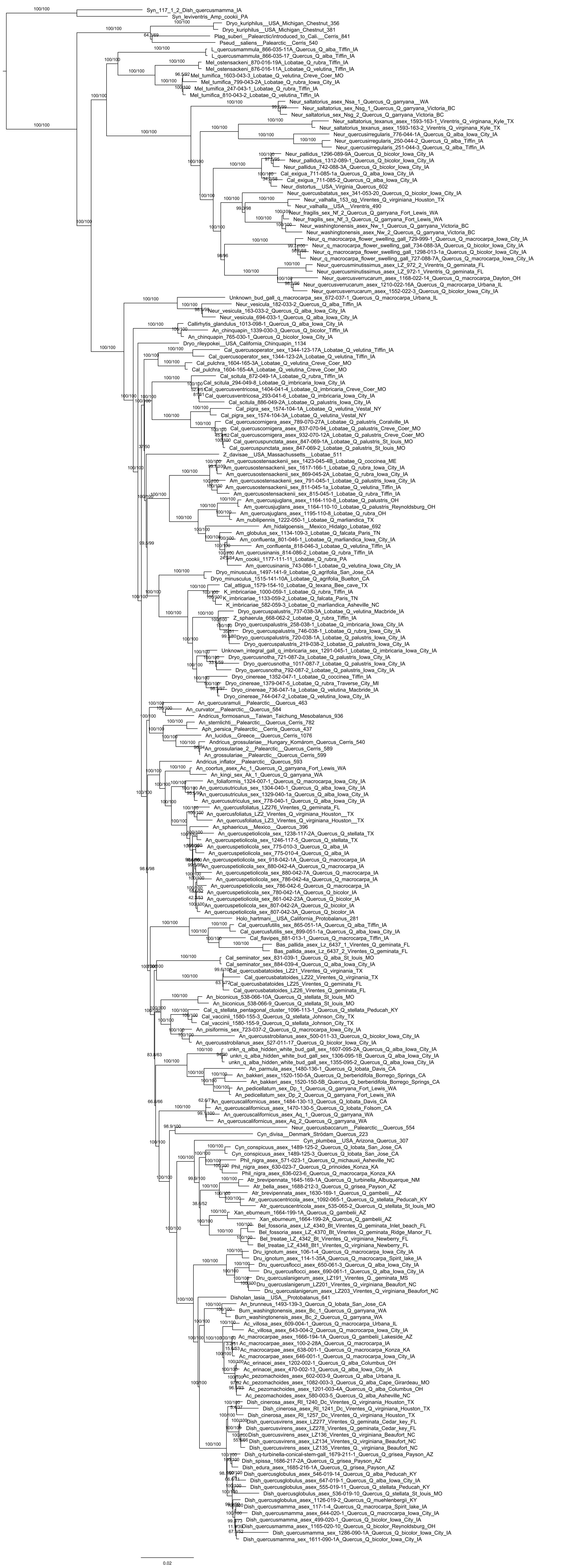

Supplemental Figure 4. ASTRAL-III phylogeny based on the combined Nearctic and Palearctic gall wasp dataset. Numbers at nodes indicate local posterior probabilities based on gene quartet frequency. Unlabeled nodes all have a probability of 1. Tip labels include the abbreviated genus, species, lab code number, tree host section, tree host species, city and state of collection, where that information was known.
For full sample information see Supplemental Tables 1 and 3.

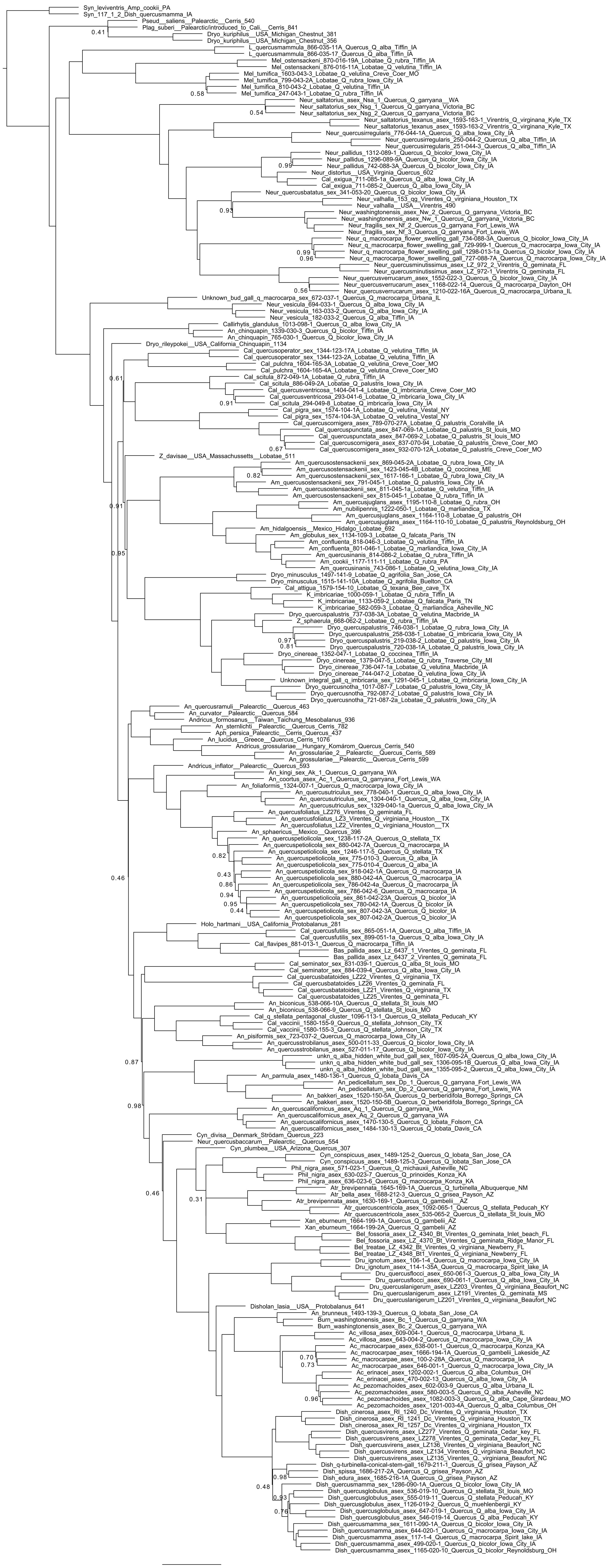

Supplemental Figure 5. Ancestral State Reconstruction (ASR) for the Palearctic + Nearctic gall wasp dataset using the 75% complete data matrix phylogeny (Supplemental Figure 2) pruned to a single specimen per taxa. Colored circles at tips indicate associations of each species with an oak section or non-*Quercus* host (see legend). Circles at tree nodes indicate reconstructions of ancestral tree hosts.

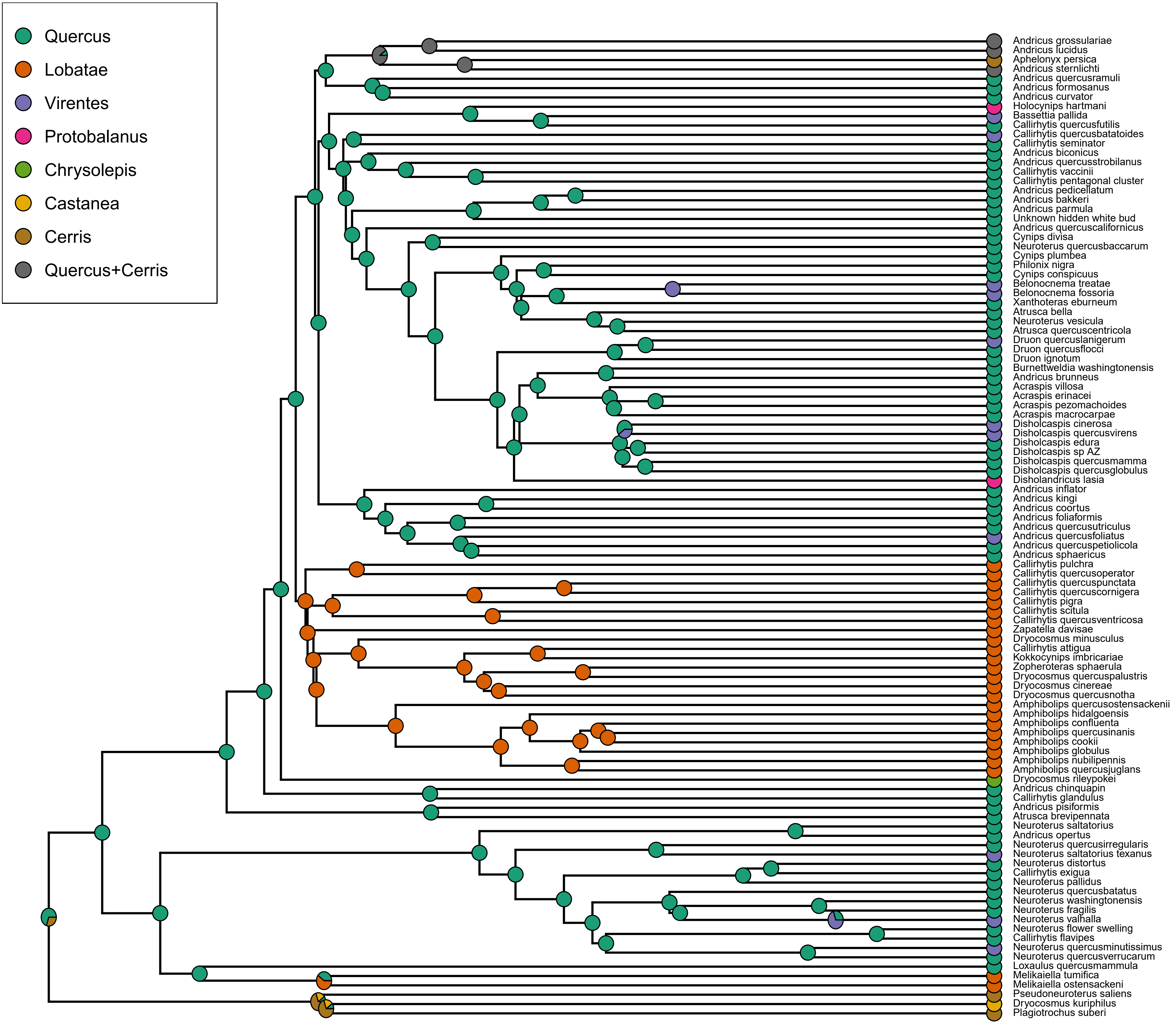

Supplemental Figure 6. Maximum likelihood phylogeny based on the Nearctic gall wasp 50% complete data matrix. Support values show bootstrap value followed by SH-likelihood ratio. Tip labels include the abbreviated genus, species, lab code number, tree host section, tree host species, city and state of collection, where that information was known. For full sample information see Supplemental Table 1.

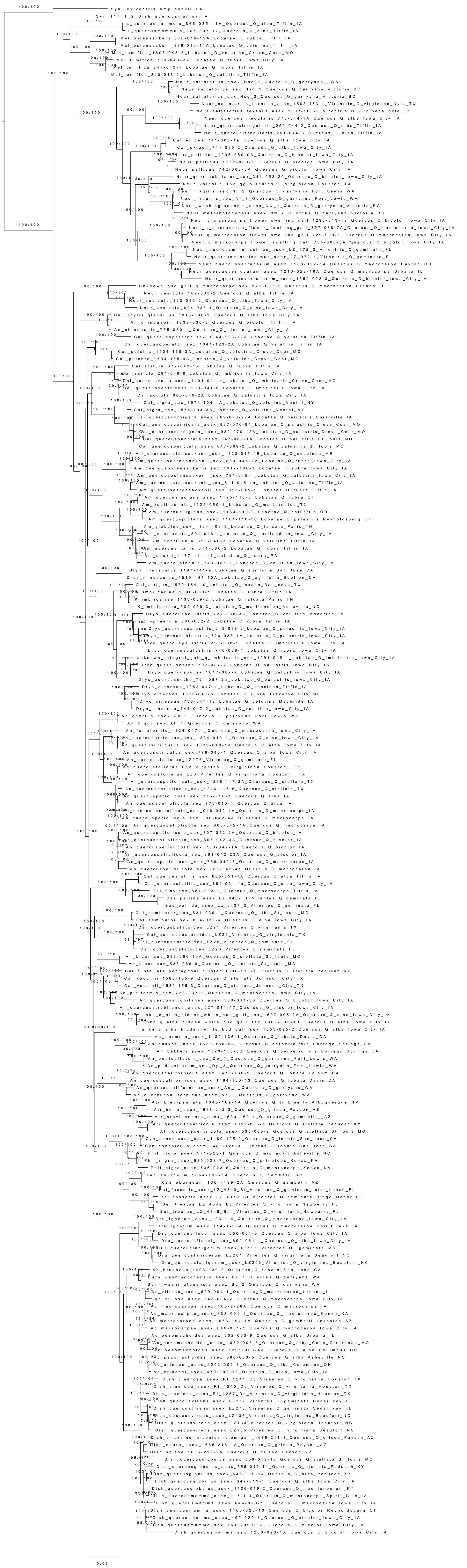

Supplemental Figure 7. Maximum likelihood phylogeny based on the Nearctic gall wasp 75% complete data matrix. Support values show bootstrap value followed by SH-likelihood ratio. Tip labels include the abbreviated genus, species, lab code number, tree host section, tree host species, city and state of collection, where that information was known. For full sample information see Supplemental Table 1.

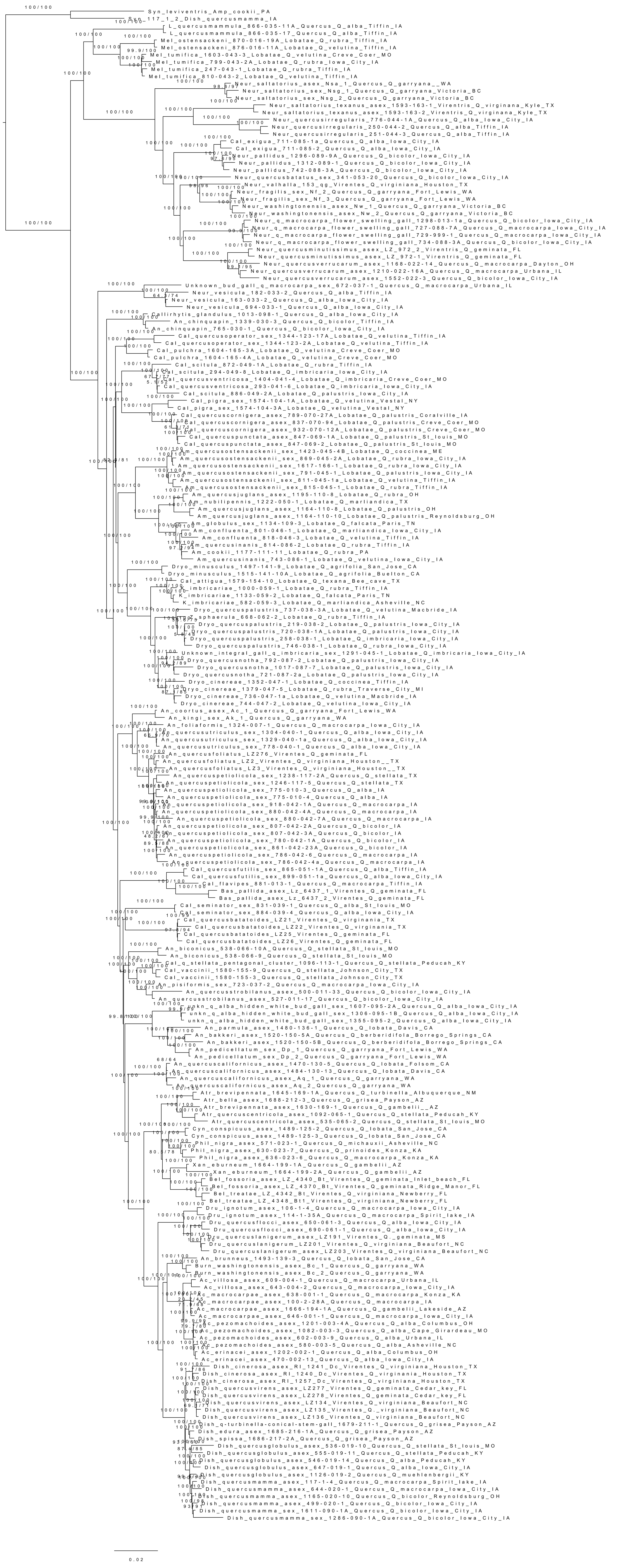

Supplemental Figure 8. Maximum likelihood phylogeny based on the Nearctic gall wasp 90% complete data matrix. Support values show bootstrap value followed by SH-likelihood ratio. Tip labels include the abbreviated genus, species, lab code number, tree host section, tree host species, city and state of collection, where that information was known. For full sample information see Supplemental Table 1.

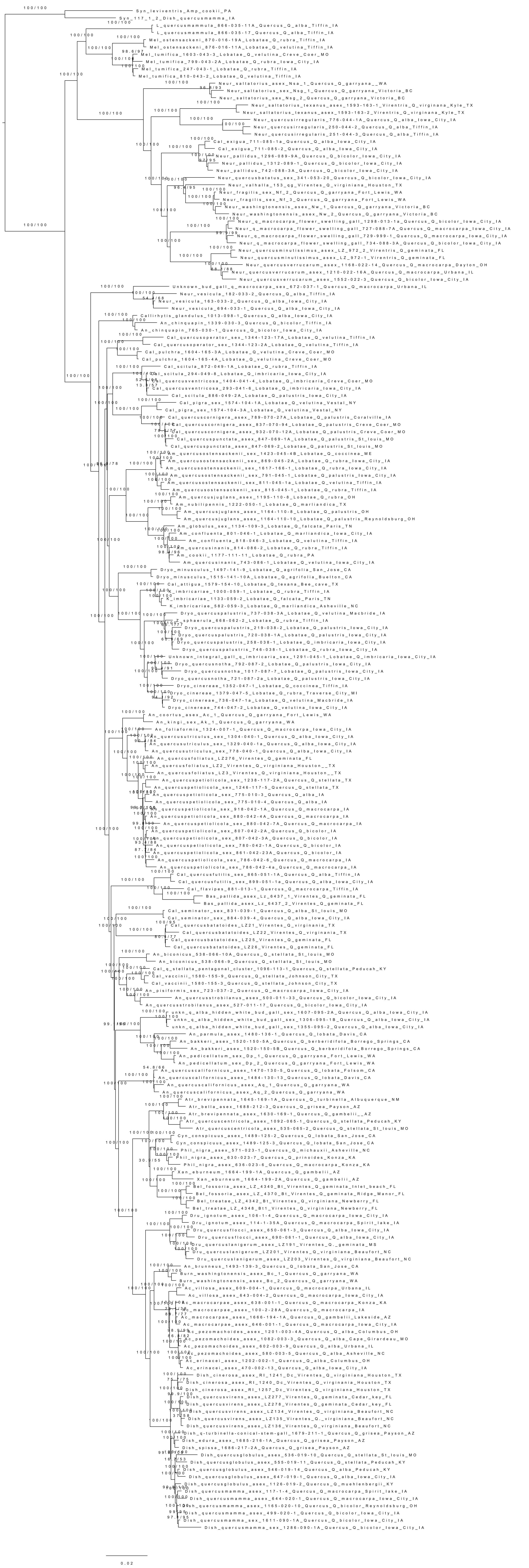
