## Supplemental file 2. Notes on Taxonomy for "Speciation in Nearctic oak gall wasps is frequently correlated with changes in host plant, host organ, or both"

**Paraphyletic genera and closed lifecycles**

A long-standing bugaboo in gall wasp research has been the accuracy of taxonomic names. As in several previous treatments of the Cynipini (Lund et al. 1998, Pujade-Villar et al. 2001, Stone et al. 2009, Cooke 2018), several gall wasp genera in our dataset, including *Andricus, Callirhytis, Neuroterus,* and *Dryocosmus*, were not paraphyletic or polyphyletic in all tree topologies. Our phylogeny also provides secondary credence to the monophyly of some Nearctic genera, including *Disholcaspis*, *Acraspis*, and *Amphibolips*, though we caution that our sampling does not encompass the full diversity of all species in any of those three genera.

Taxonomy in gall wasps has been further complicated by the common presence of both asexual and sexual generations that induce galls of different morphologies on different plant tissues. For many gall wasps, the alternate generation is currently unknown, either because it is undiscovered, because the species is entirely asexual (as has been speculated is the case for *Andricus quercuscalifornicus*; Joseph et al. 2011), or because it has been described as a different species altogether (Pujade-Villar et al. 2010). Previous studies have used molecular and ecological data to “close” life cycles, identifying the alternate generation of a named galler (e.g., Hood et al. 2018). We find four pairs of gall wasps, each previously described as different species, that may represent alternate generations of the same species (Table S2A). Though this was not a primary goal of our study, this result highlights the degree to which basic natural history remains undiscovered in this charismatic group of insects and the potential for genetic information to provide an accessible complement to morphological taxonomic studies.

**Table S2A. Four instances where UCE data suggest that two previously named gallers are actually the sexual and asexual generations of a single species (Supplementary Figures 1-6). These combinations are emergent hypotheses and should be subject to further molecular, behavioral, and ecological testing.**

| Gamic (sexual) generation | Description | Agamic (asexual) generation | Description |
| --- | --- | --- | --- |
| *Callirhytis scitula* | Bud/petiole gall; red oaks; Spring; wasps emerged in June | *Callirhytis quercusventricosa* | Stem gall; red oaks; Summer/Fall; wasps emerged in April |
| *Callirhytis exigua* | Flower gall; white oaks; Early Spring; wasps emerged in | *Neuroterus distortus* | Stem/petiole gall; white oaks; Spring; wasps emerged in |
| *Amphibolips nubilpennis* | Leaf gall; red oaks; Spring | *Amphibolips quercusjuglans* | Acorn gall; red oaks; Fall |
| *Amphibolips cookii* | Bud gall; red oaks; Summer/Fall | *Amphibolips quercusinanis* | Leaf gall; red oaks; Spring |

In one other instance, UCEs combine named species but other evidence suggests that they should be maintained as different species pending additional study. *Dryocosmus quercuspalustris* (sex) and *Zopheraterus sphaerula* (asex) might initially seem to fall into the category of a closed lifecycle, but another species that we did not sequence, *Zopheroteras guttatum,* looks similar to *Z. sphaerula* but differs in coloring (mottled vs. solid) and is found on a different subset of tree species from *Z. sphaerula*. *Dryocosmus quercuspalustris* has described host ranges that overlap with both *Z. sphaerula* and *Z. guttatum*, such that we suspect each *Zopheroteras* may actually be the asexual generation of a different cryptic species of *D. quercuspalustris*. Additional sampling and sequencing will be required to determine these relationships, so for now we consider them part of a potential complex.

**Other taxonomic notes**

The stem-girdling galls *Callirhytis quercuscornigera* and *C. quercuspunctata* do not form reciprocally monophyletic clades in any of our trees. Both of these species induce galls on trees in section *Lobatae*, and in both the sexual generation produce a small leaf gall along the vein, while the asexual generation is a swelling of the stem. It could be that these are one species with differences in asexual gall morphology on different trees resulting from host or other environmental differences.

What we have called the “*Atrusca* clade” on our trees (Figures 1-3) consists of three potentially different species, *Atrusca quercuscentricola, Atrusca bella,* and *Atrusca brevipennata*. All three species induce galls on leaves that all have internal radiating fibers. *A. quercuscentricola* forms its own clade but the other two species do not. *A. quercuscentricola* is found on *Q. stellata* in the *Stellatae* subsection of section *Quercus*, while *A. bella* and *A. brevipennata* are both found on trees in the *Leucomexicana* subsection of section *Quercus*. *Atrusca bella* and *A. brevipennata* are also both described as having similar gall morphology and have some overlap in their host associations but differ in geographic locations (Kinsey 1936). We have collected *A. bella* and *A. brevipennata* galls on three different tree hosts in Arizona and New Mexico, but they do not separate into separate clades based on UCE data (Supplementary Figures 1-4). More sequencing of galls from the southwestern U.S. and Mexico is required to resolve these relationships and potentially update species identification and host ranges.

*Acraspis erinacei* and *Acraspis pezomachoides* are collapsed in our trees (Figure 1) because they do not form reciprocally monophyletic groups. One *A. pezomachoides* sometimes groups with *A. erinacei* collected from Asheville, NC (Supplemental Figures 1-2), while in other trees the species are reciprocally monophyletic (Supplementary Figures 3-4). These two species induce galls with different morphologies though are found on the same tree host and tissue. One pre-phylogenetics era hypothesis (Kinsey 1936) posited that *A. erinacei* - based on its wide geographical distribution and morphological variability - might be prone to hybridization with other closely related *Acraspis*. This hypothesis, or alternatives involving a recent radiation or multiple cryptic species within each named species, have credence and should be investigated. Additional collections coupled with phylogenetic or population genetic analyses would be welcome in this possible species complex.

*Disholcaspis quercusglobulus* and *Disholcaspis quercusmamma* also do not form monophyletic groups in all trees (Supplemental Figures 1-3; Figure 5), Here, our collections allowed for a deeper dive into correlation between phylogeny and host association and these appear to be a complex of several host-associated species (see main paper, Figure 5 and Discussion).

Finally, *Aphelyonx persica* from the Blaimer et al. (2020) dataset did not comport with the placement of *Aphelonyx* species in previous phylogenies (e.g., Stone et al. 2009; Nicholls et al. 2017). We fished a 28S sequence from this UCE dataset and it matched 100% to an *Andricus* sp. sequence on genbank, so we suspect that this species should be moved to *Andricus*. Though *A. persica* is known only from the asexual generation (on trees in subgenus *Cerris*, section *Cerris*), its placement in our tree within a clade of host alternating species suggests that the sexual generation may be found on oaks in subgenus *Quercus*.

**Blaimer, B. B., D. Gotzek, S. G. Brady, and M. L. Buffington**. **2020**. Comprehensive phylogenomic analyses re-write the evolution of parasitism within cynipoid wasps. BMC Evol. Biol. 20: 155.

**Cooke, C. L.** 2018. Forest Micro-Hymenoptera, including those attacking trees (Cynipidae oak gall wasps) and those potentially defending them (parasitic Pteromalidae). PhD Dissertation, Univ. Maryland.

**Hood, G. R., L. Zhang, L. Topper, P.F.P. Brandão-Dias, G.A. Del Pino, M. S. Comerford, and S. P. Egan. 2018.** ‘Closing the Life Cycle’ of *Andricus quercuslanigera* (Hymenoptera: Cynipidae). Ann. Entomol. Soc. Am. 111: 103–113.

**Joseph, M. B., M. Gentles, and I.S. Pearse.** 2011. The parasitoid community of *Andricus quercuscalifornicus* and its association with gall size, phenology, and location. Biodivers. Conserv. 20: 203 – 216.

**Kinsey, A. C.** **1936**. The origin of the higher categories in *Cynips*. Indiana Univ. Publ Sci Ser. 4: 1–334.

**Lund J. N, J. R. Ott, and R. J. Lyon.** 1998. Heterogony in *Belonocnema treatae* Mayr (Hymenoptera: Cynipidae). Proc. Entomol. Soc. Wash. 100: 755–763.

**Nicholls, J. A., Melika, G. and Stone, G. N., 2017.** Sweet tetra-trophic interactions: multiple evolution of nectar secretion, a defensive extended phenotype in Cynipid gall wasps. The American Naturalist. 189: 67–77.

**Pujade-Villar, J., D. Bellido, G. Segu, and G. Melika**. **2001**. Current state of knowledge of heterogony in Cynipidae (Hymenoptera, Cynipoidea). Sess. Conjunta Entomol. 11: 87–107.

**Stone, G. N., A. Hernandez-Lopez, J. A. Nicholls, E. di Pierro, J. Pujade-Villar, G. Melika, and J. M. Cook**. **2009**. Extreme Host Plant Conservatism During at Least 20 Million Years of Host Plant Pursuit by Oak gall wasps. Evolution. 63: 854–869.

**Weld, L. H.** **1960**. Cynipid galls of the southwest. Ann Arbor MI Priv. Print. 35.
