## Supplemental file 1. images of gallwasps for "Speciation in Nearctic oak gall wasps is frequently correlated with changes in host plant, host organ, or both"

Supplemental File 1. Lateral and (where available) wing pictures for representatives of each species or species group sequenced in this study. Lab code numbers for each wasp can be used in concert with Supplemental Table 1 to find additional collection details. Female wasps are shown except in cases where no females were reared. Though these individuals were destructively sampled, vouchers specimens from many of the same collections have been deposited in the Smithsonian or the Frost Entomological Museum at the Pennsylvania State University (Supplementary Table 1).

*Acraspis erinacei* - Wasp: 1202-002-1

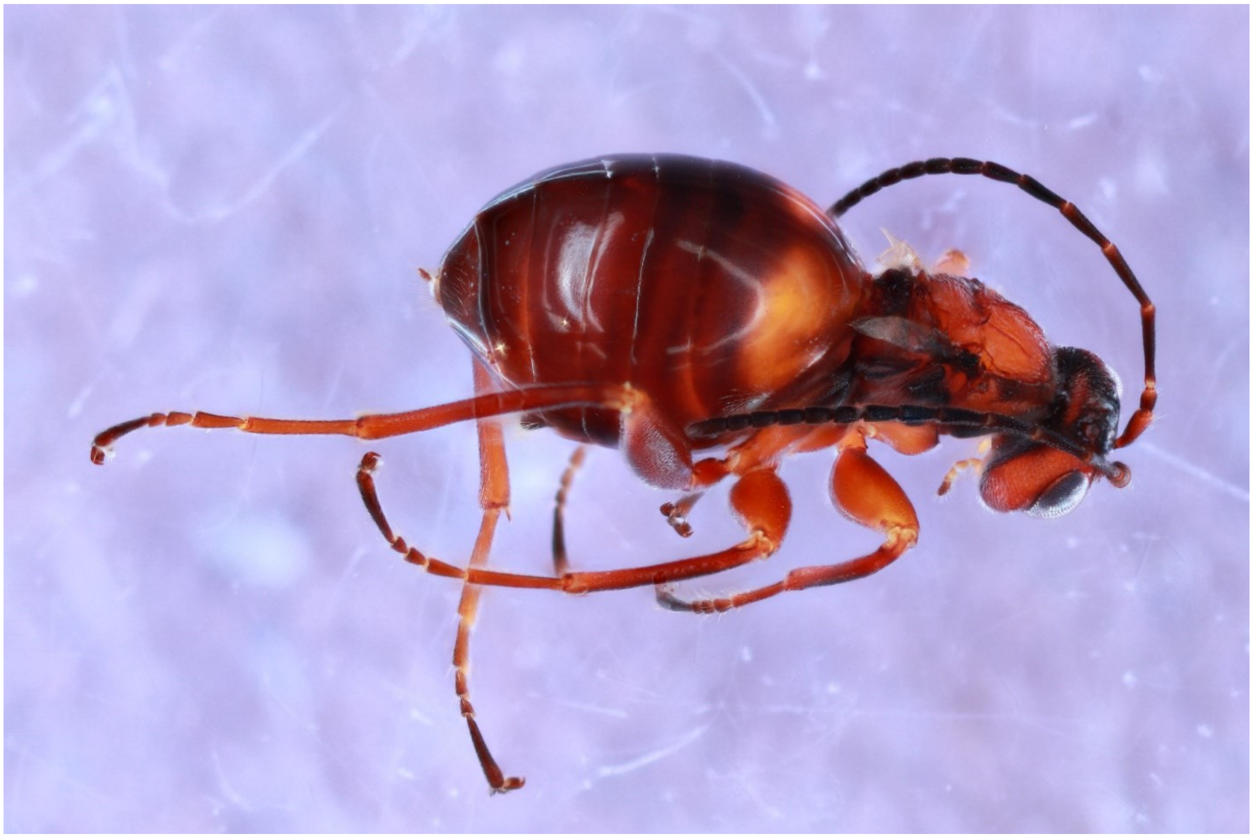

*Acraspis macrocarpae* - Wasp: 638-001-001

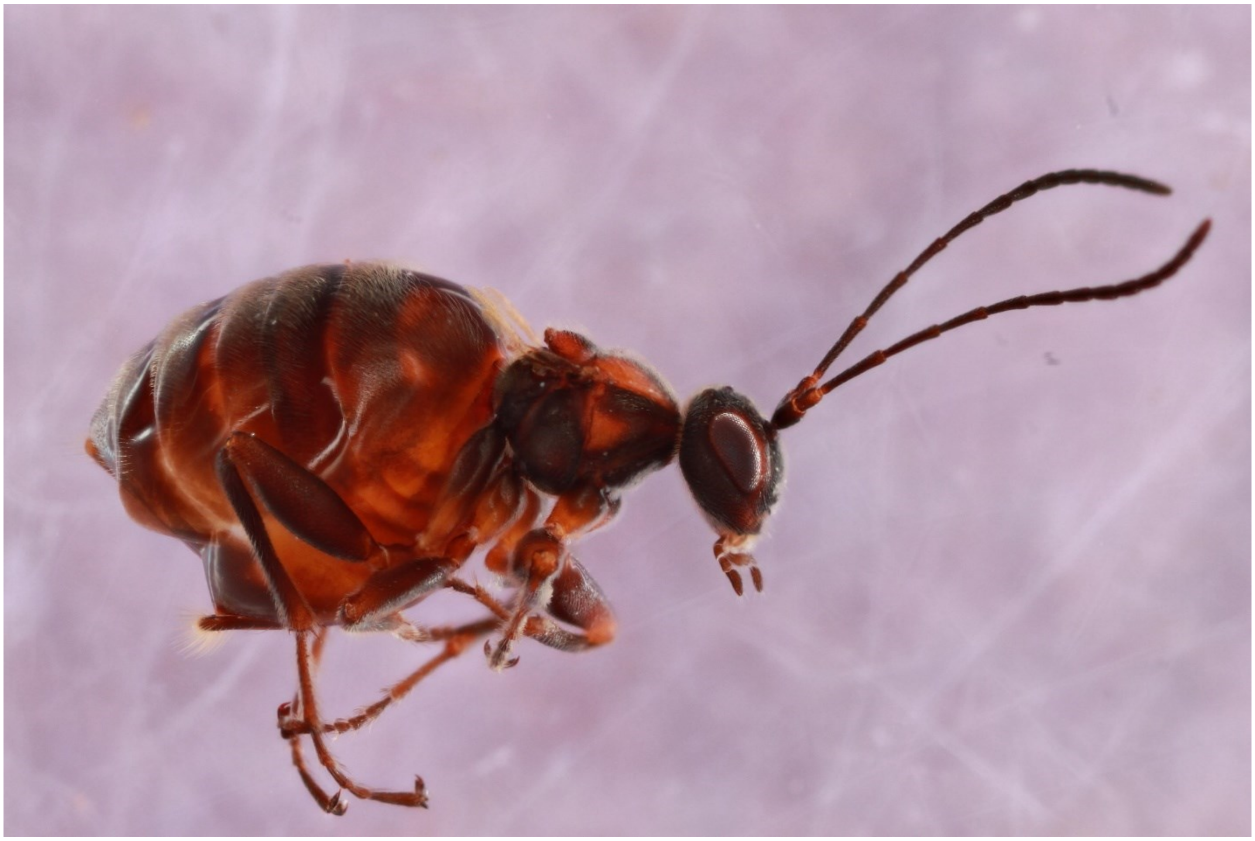

*Acraspis pezomachoides* - Wasp: 1201-003-4A

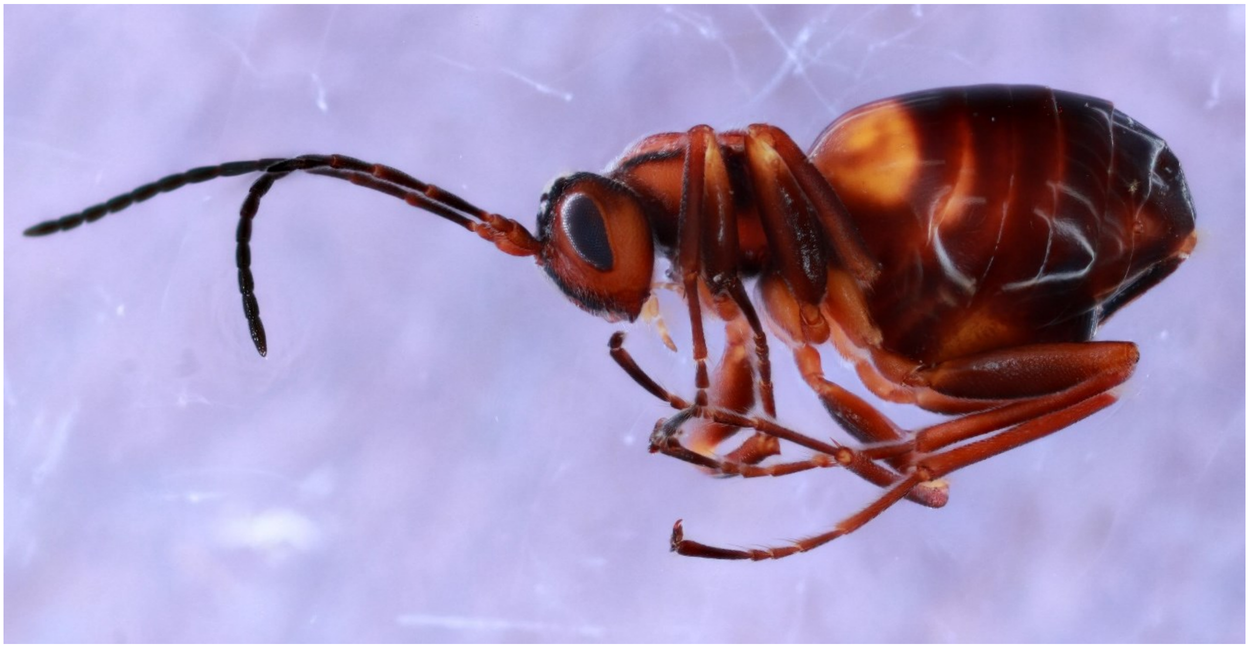

*Acraspis villosa* - Wasp: 609-004-001

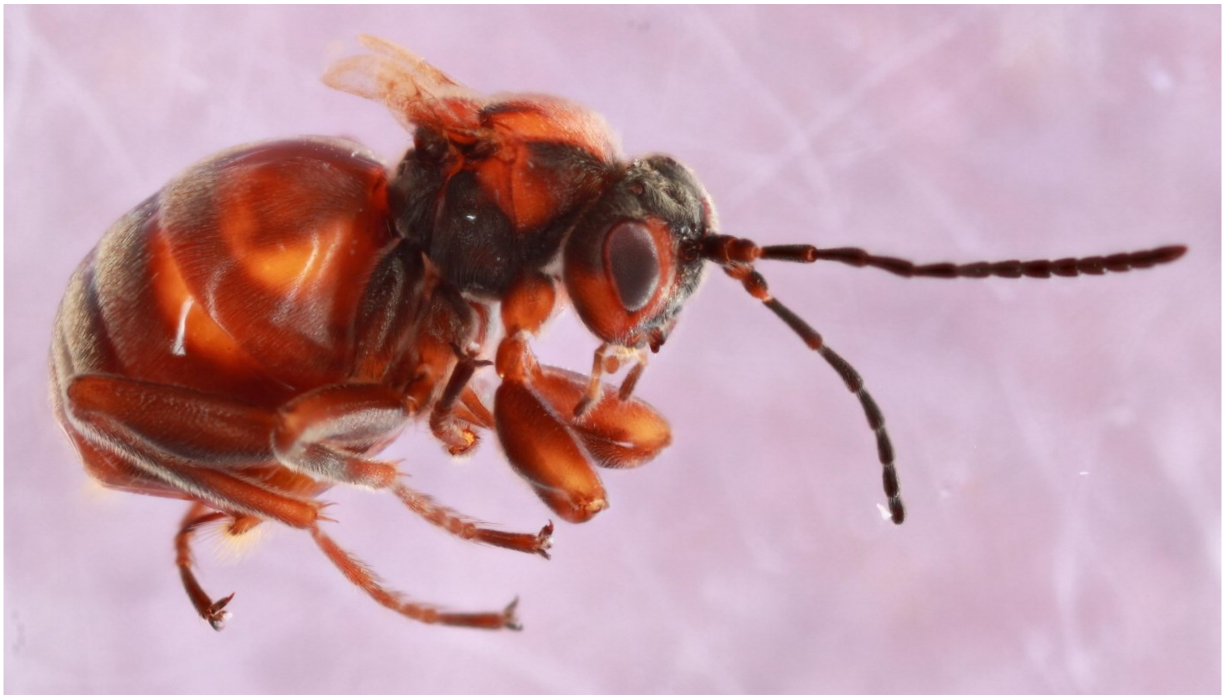

*Amphibolips cookii* - Wasp: 1177-111-11

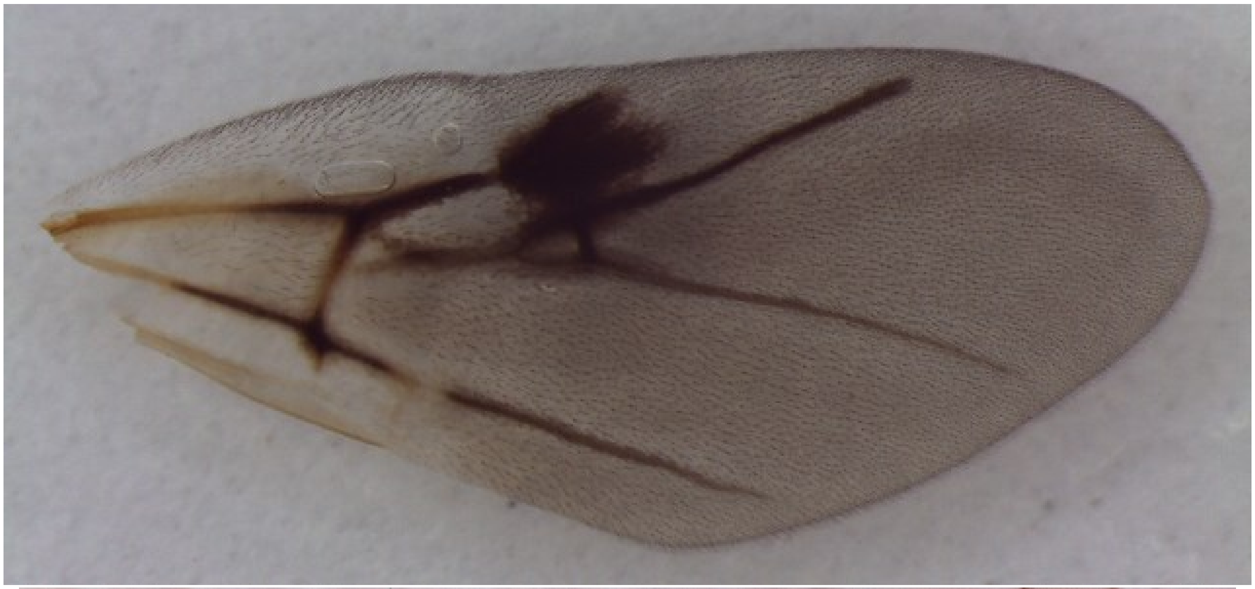

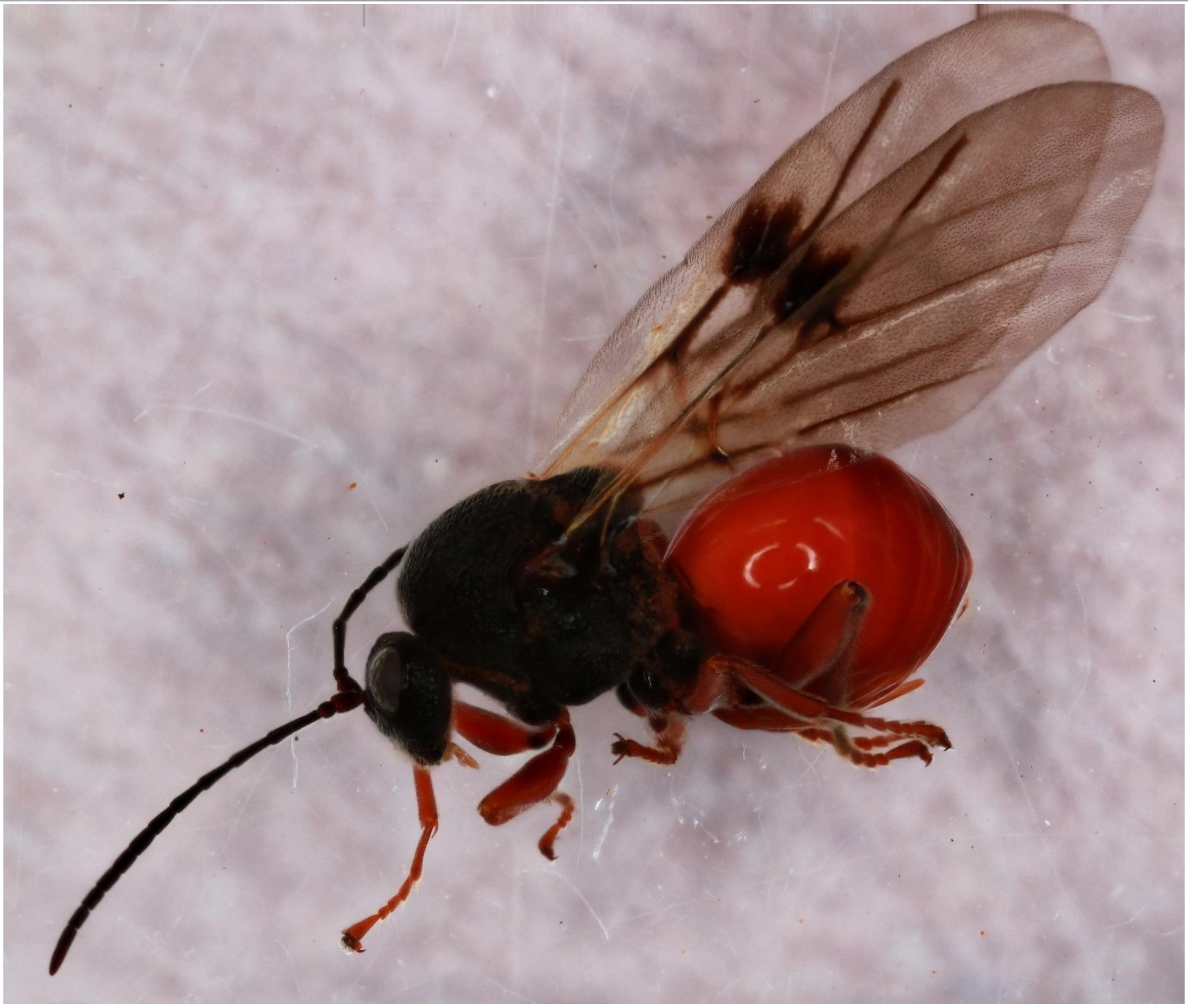

*Amphibolips confluenta* - Wasp: 818-046-5

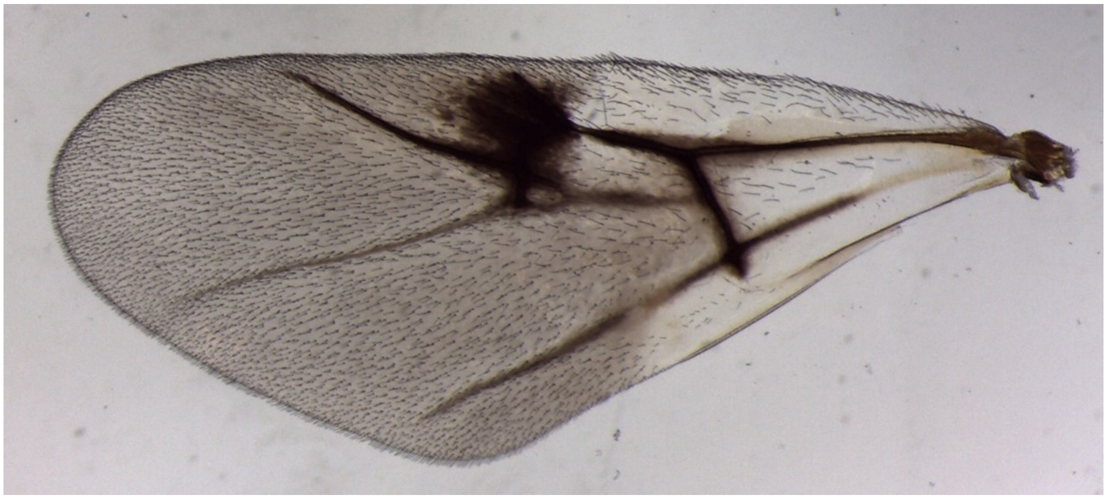

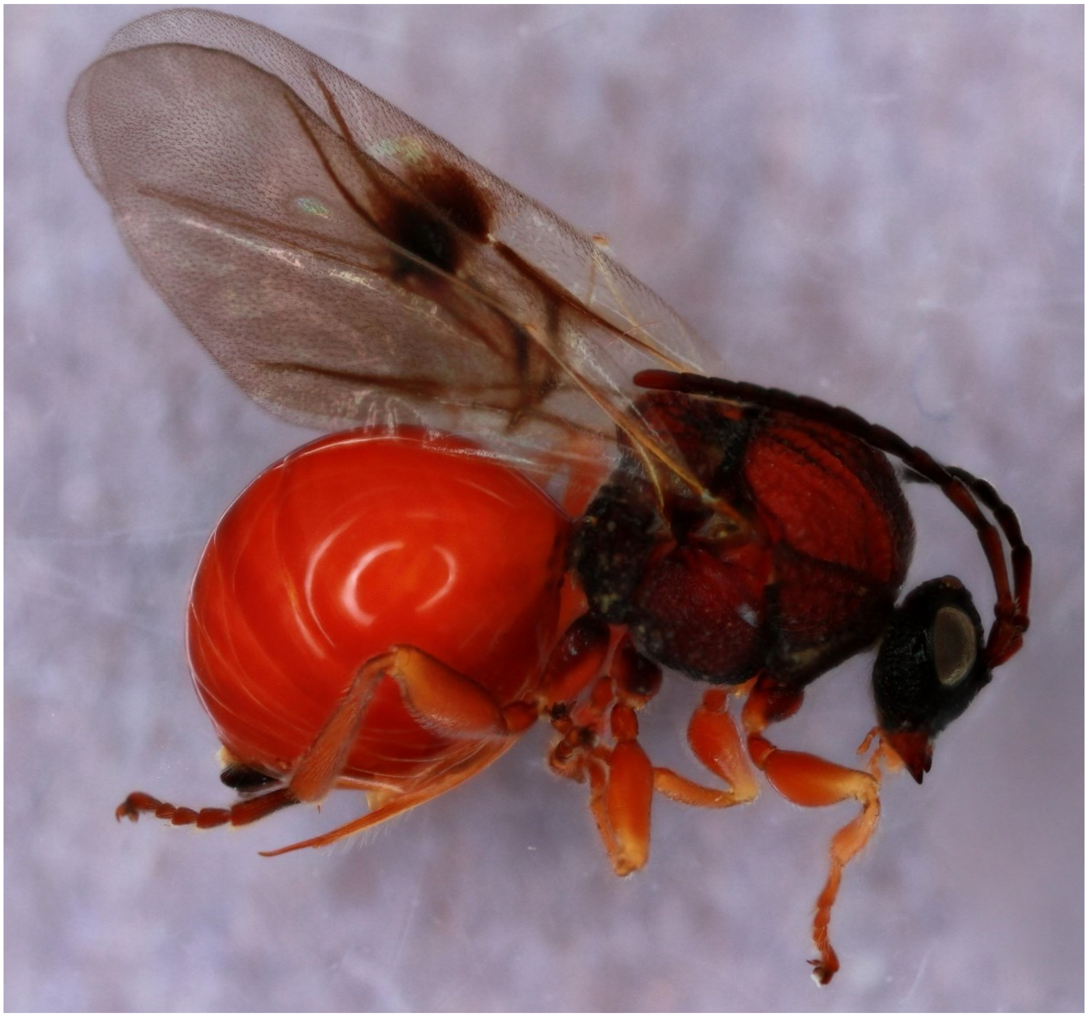

*Amphibolips nubilipennis* - Wasp: 1222-050-1

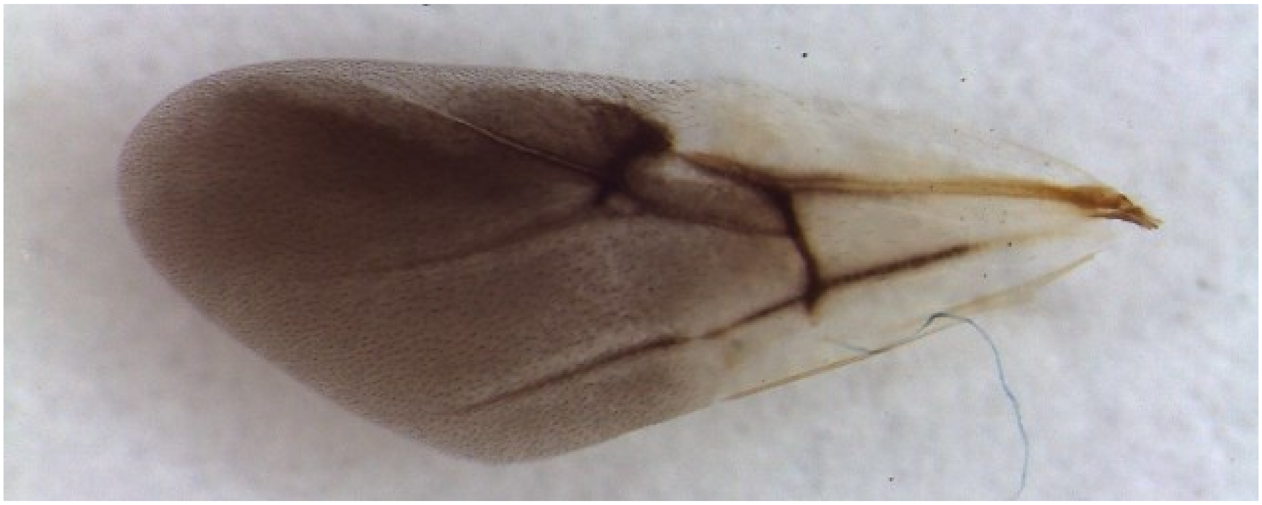

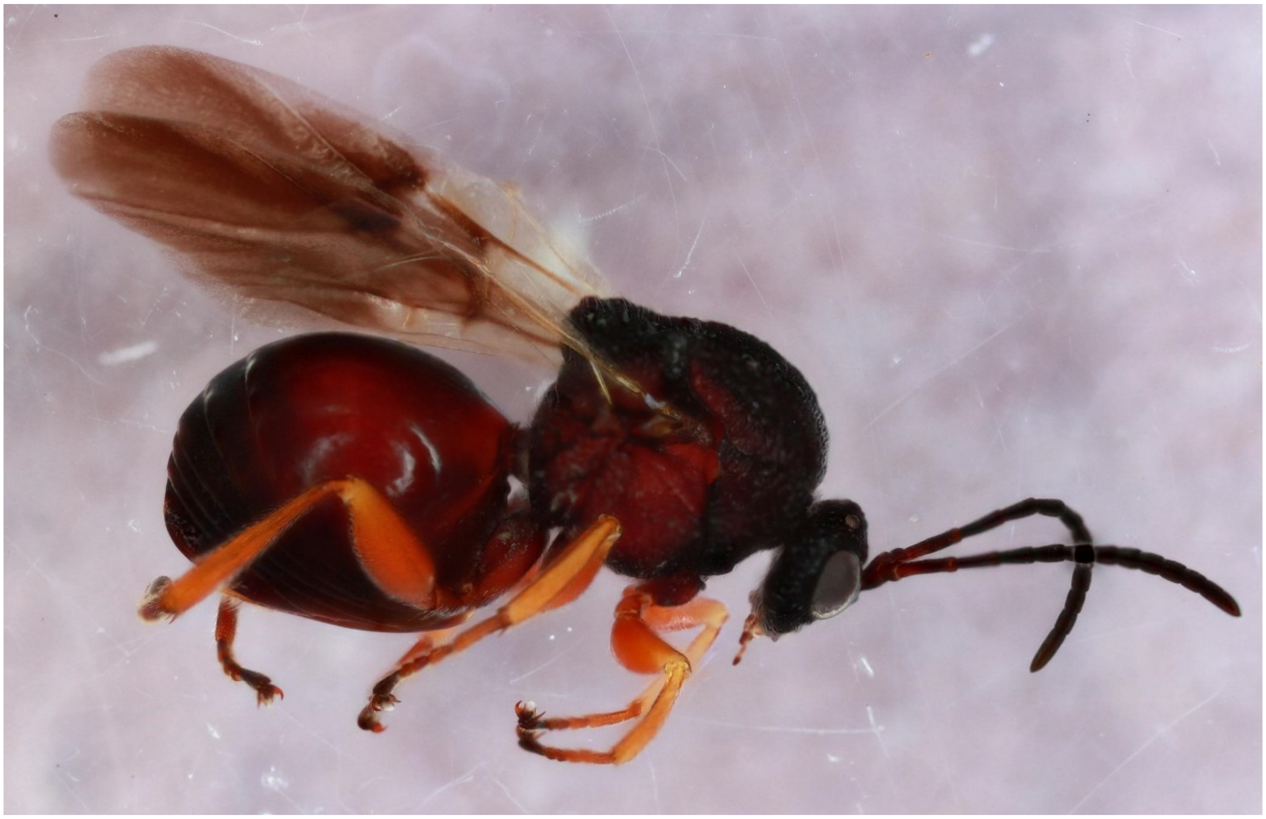

*Amphibolips quercusinanis* - Wasp: 814-086-002

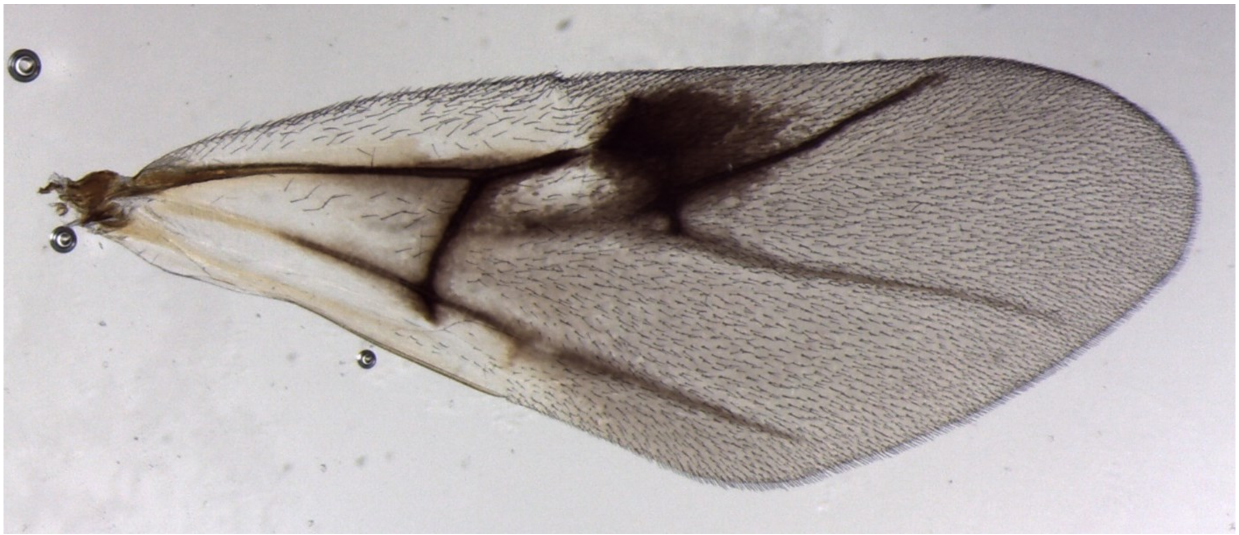

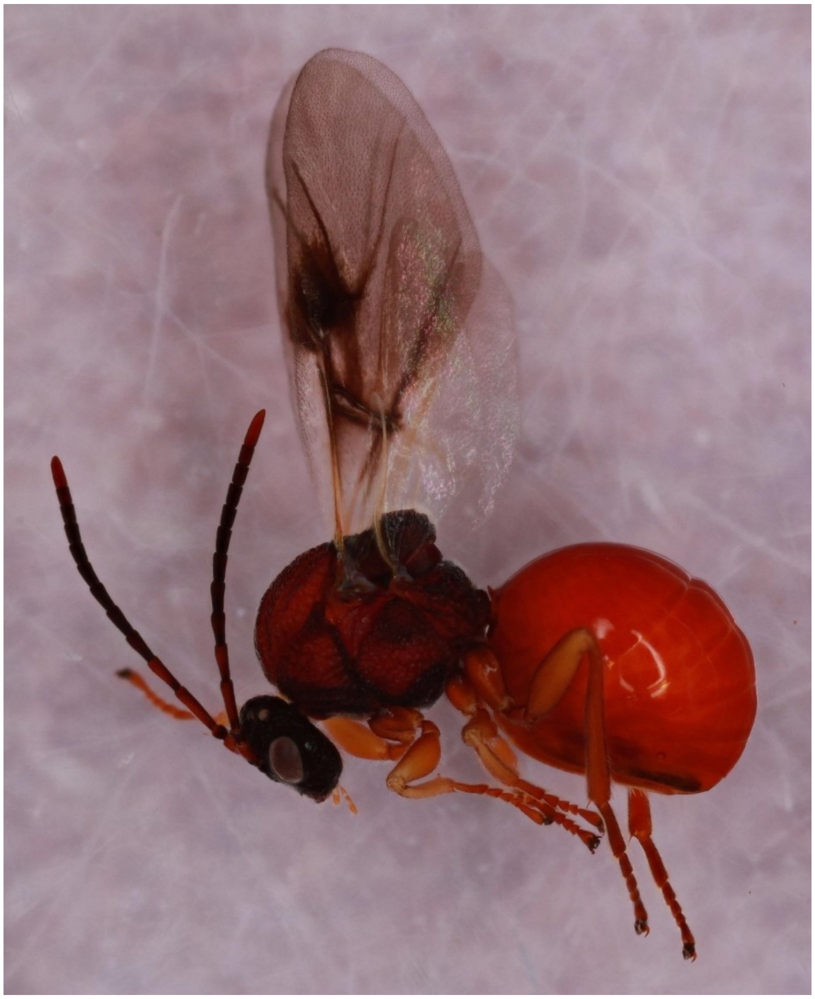

*Amphibolips quercusjugulans* - Wasp: 1195-110-8

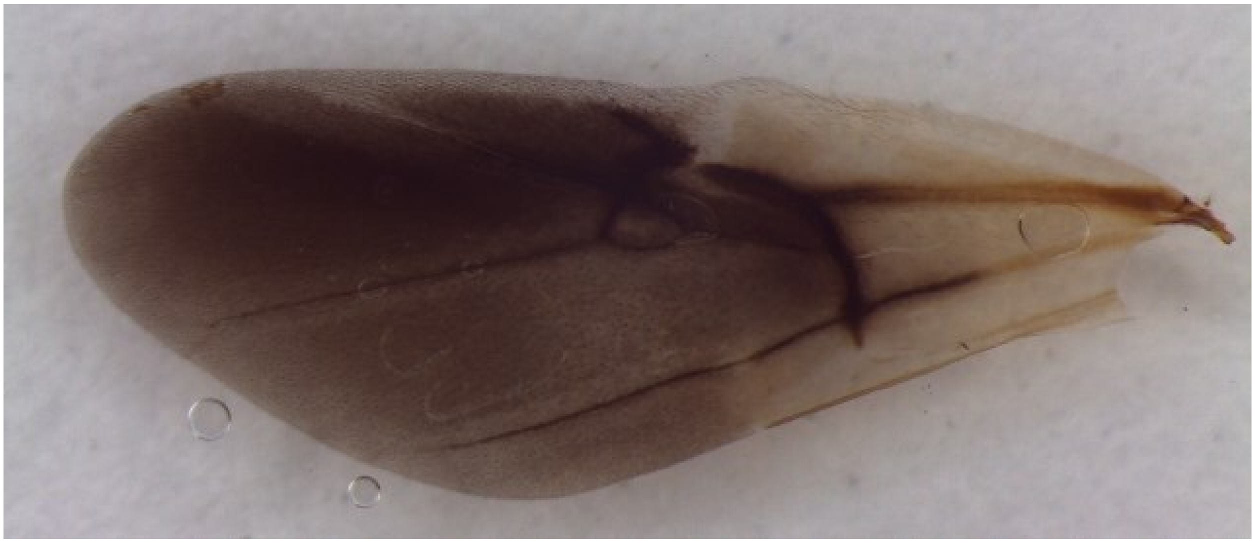

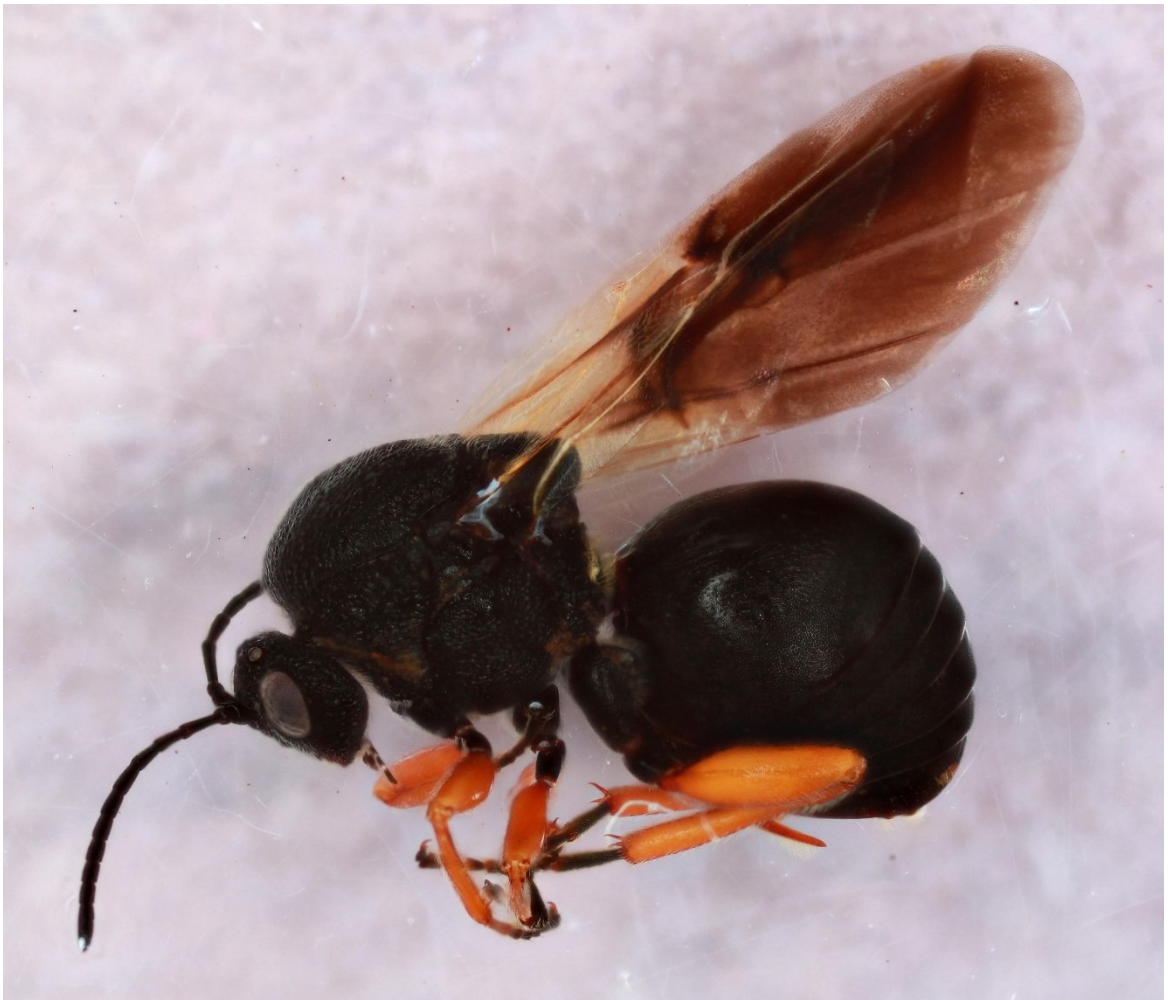

*Andricus biconicus* - Wasp: 1520-150-5B

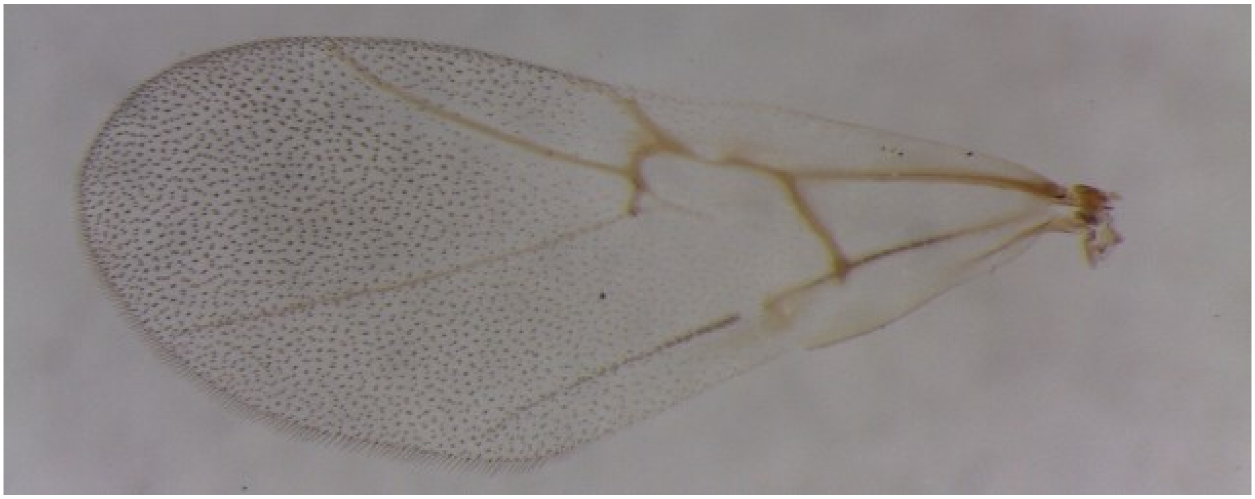

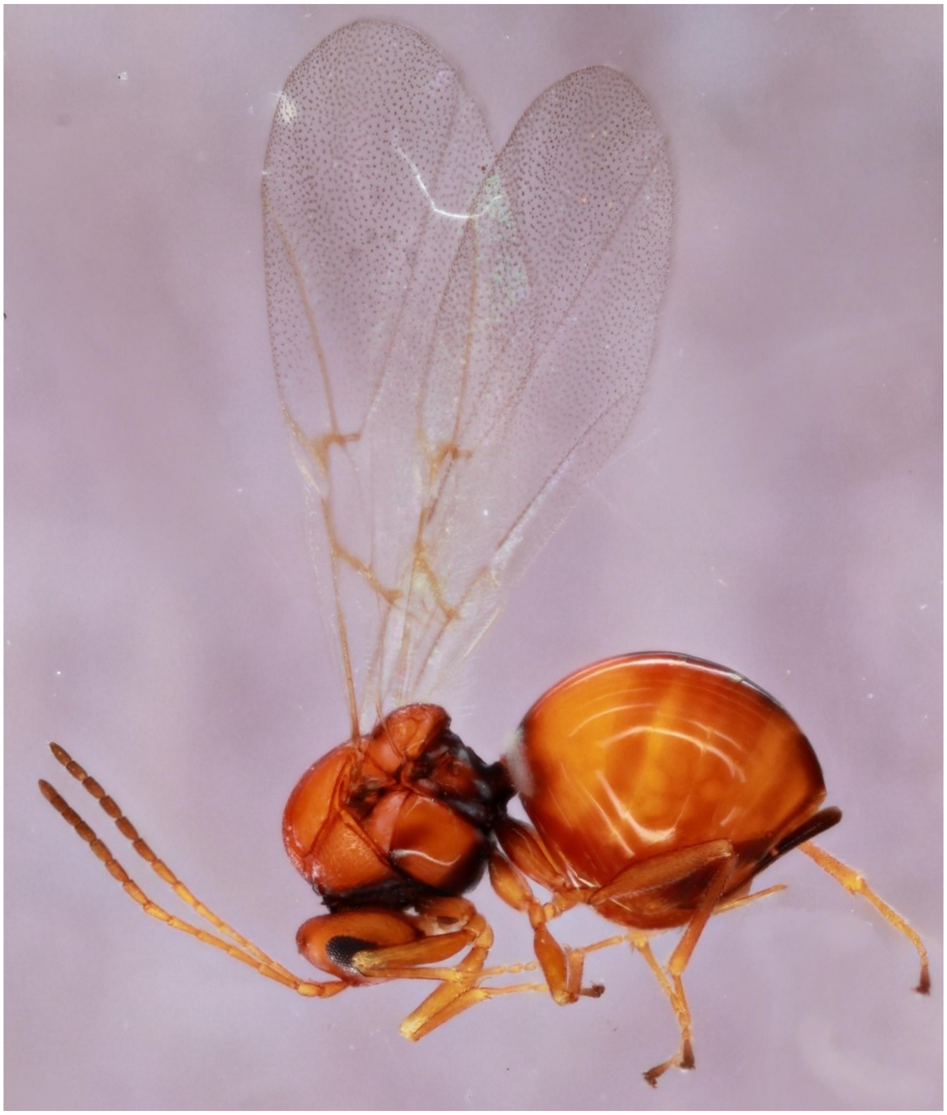

*Andricus biconicus* - Wasp: 538-066-9

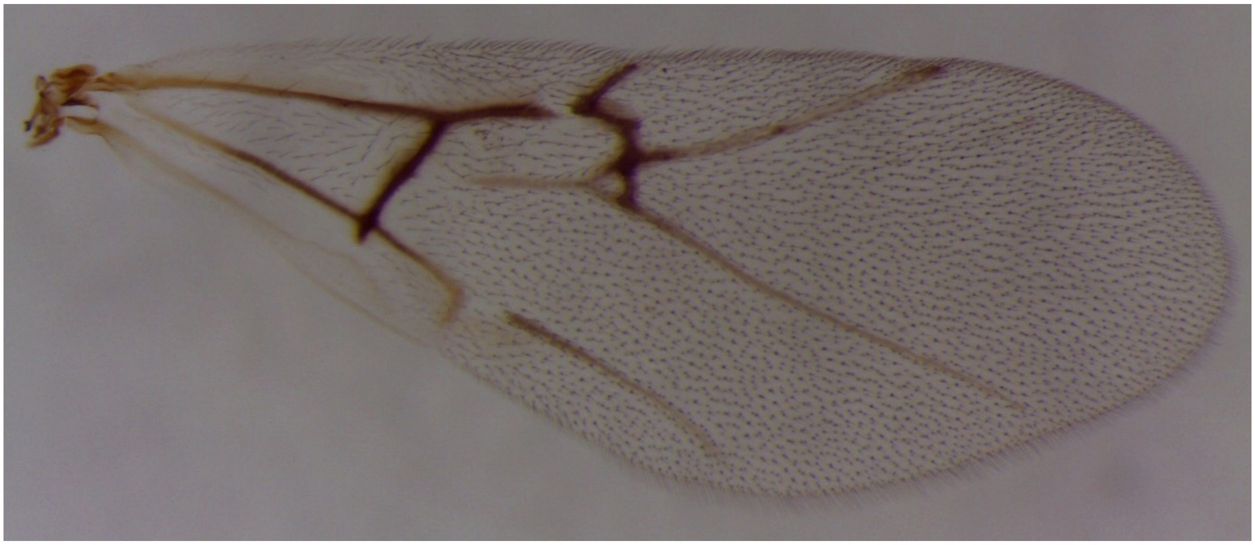

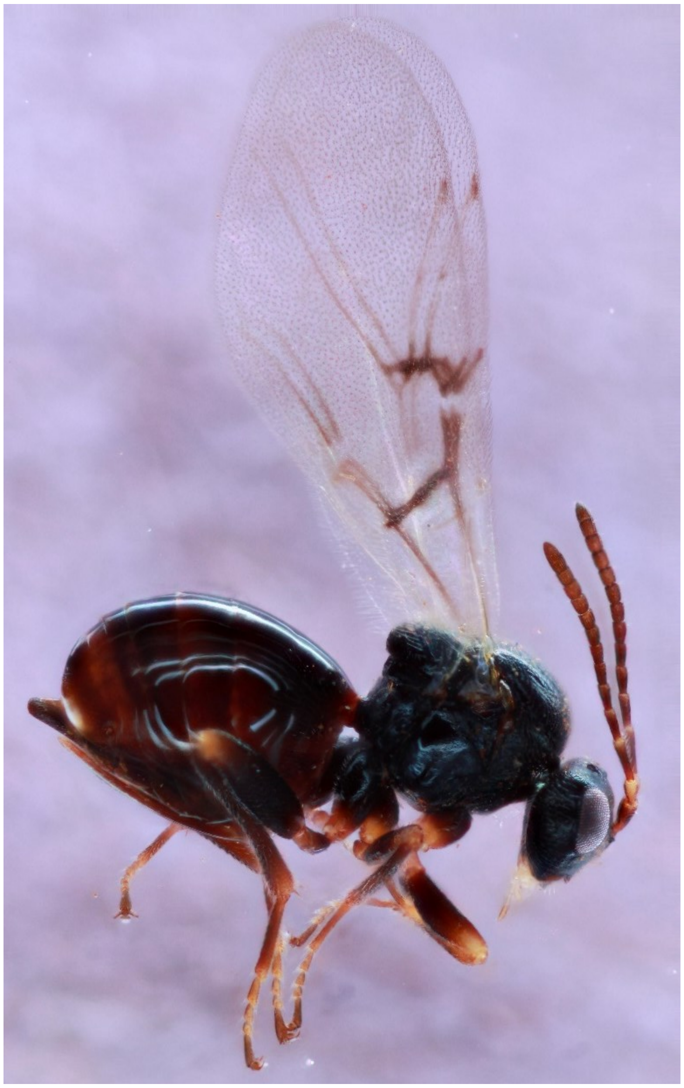

*Andricus brunneus*- Wasp: 1494-139-3

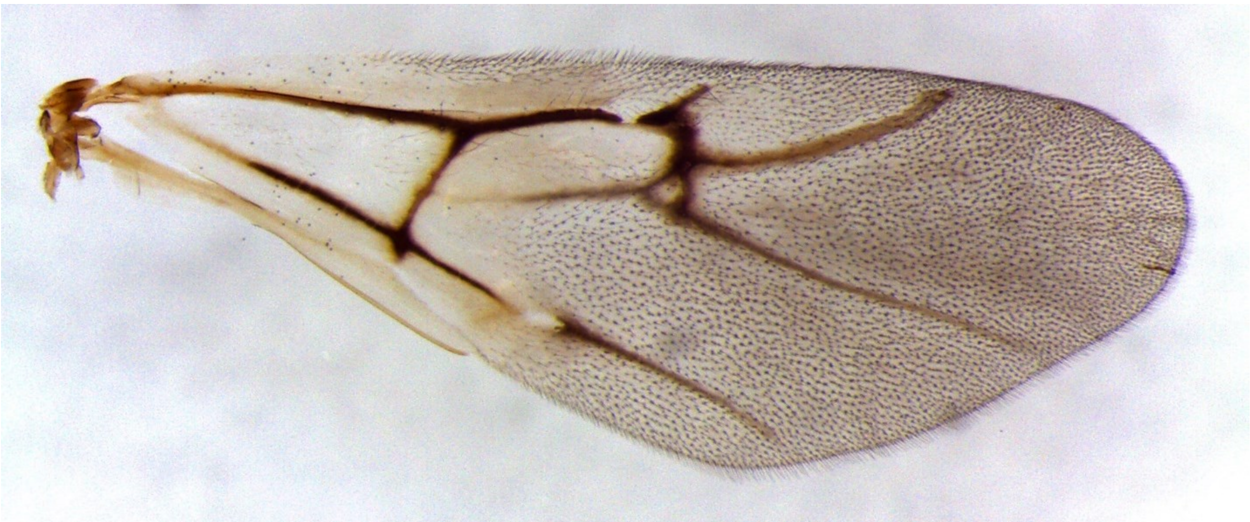

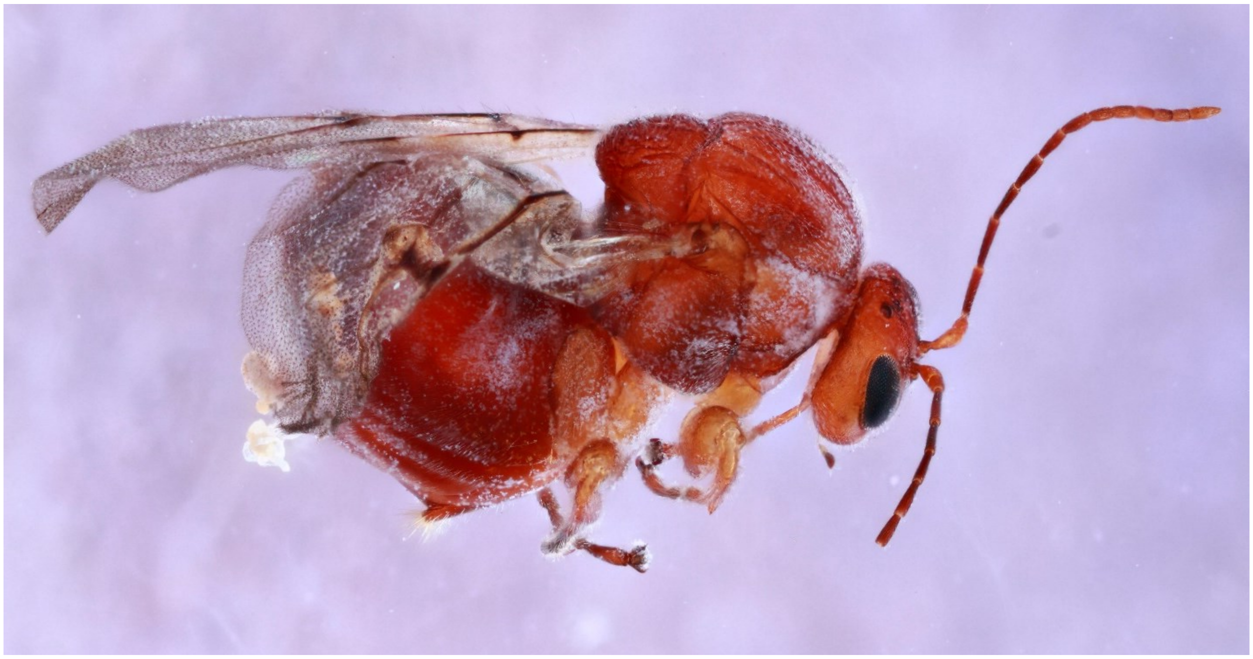

*Andricus chinquapin* - Wasp: 765-030-001

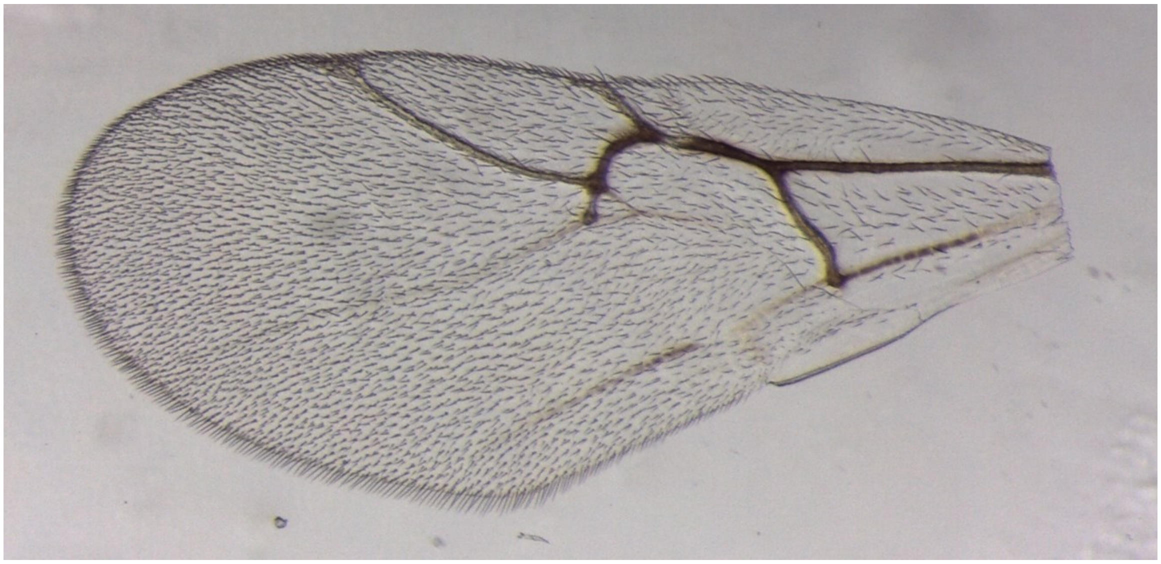

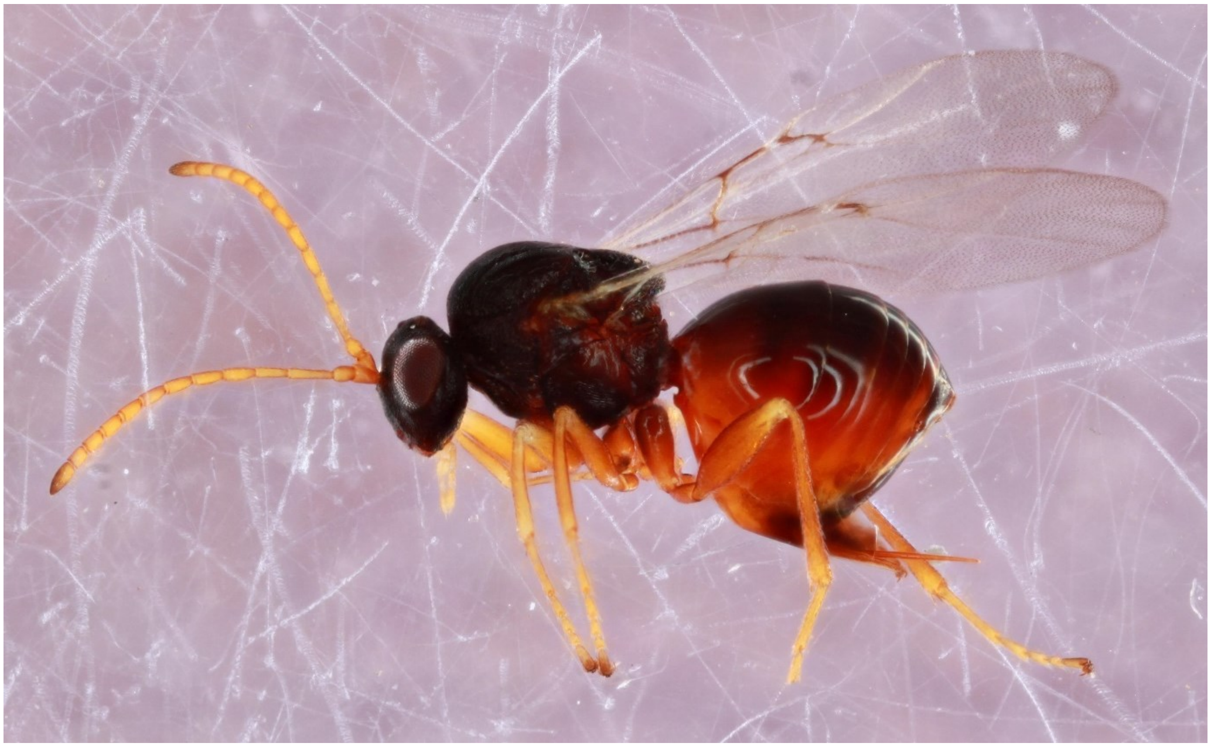

*Andricus coortus* - Wasp: Ac_1

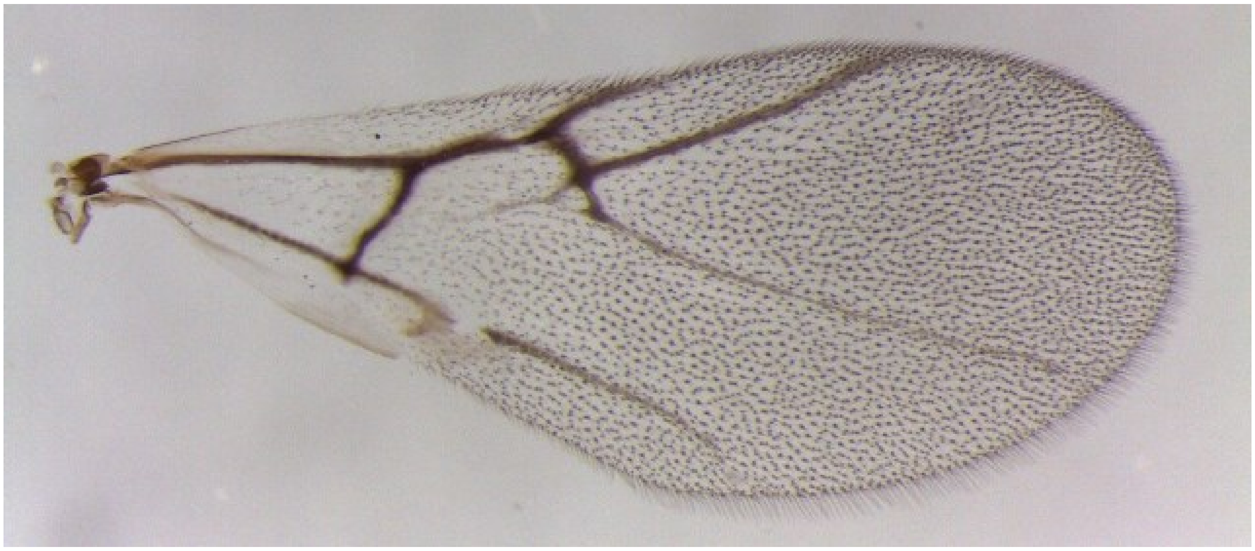

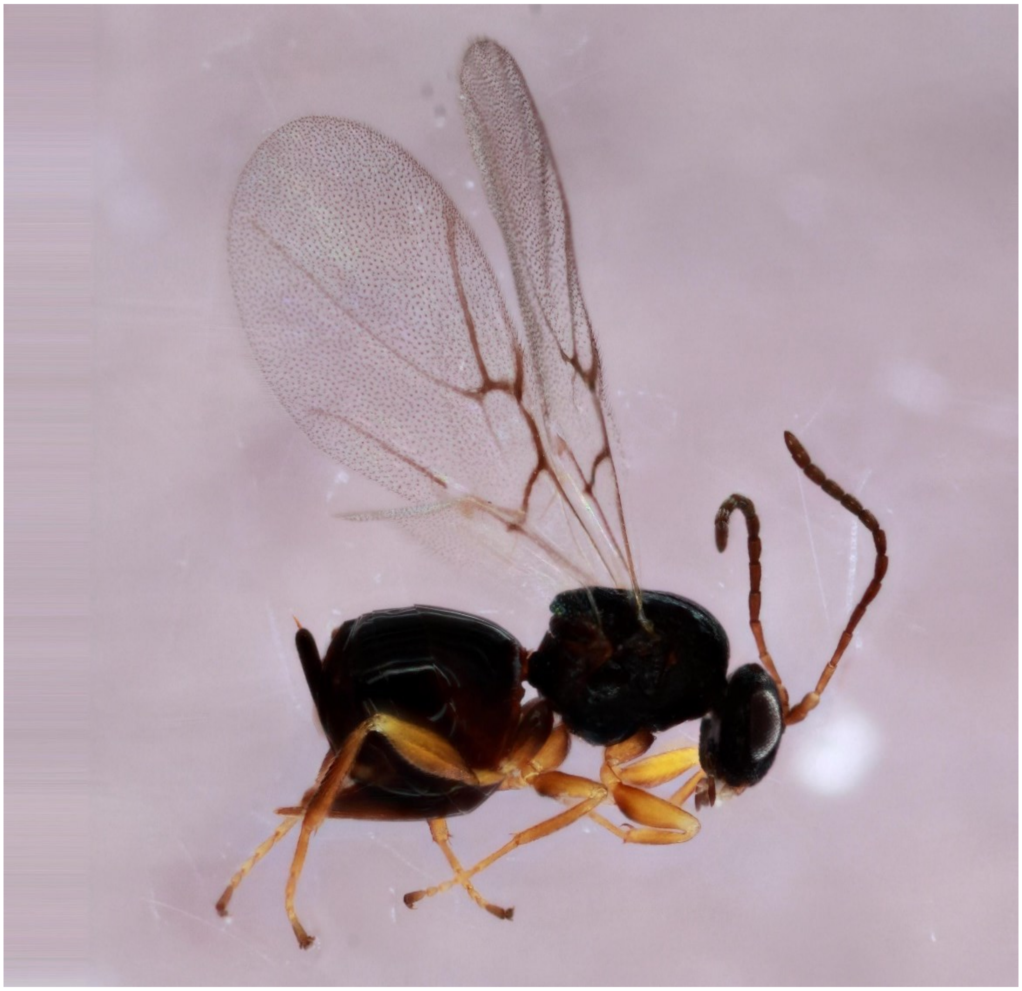

*Andricus foliaformis* - Wasp: 1324-001-1

(no wing picture)

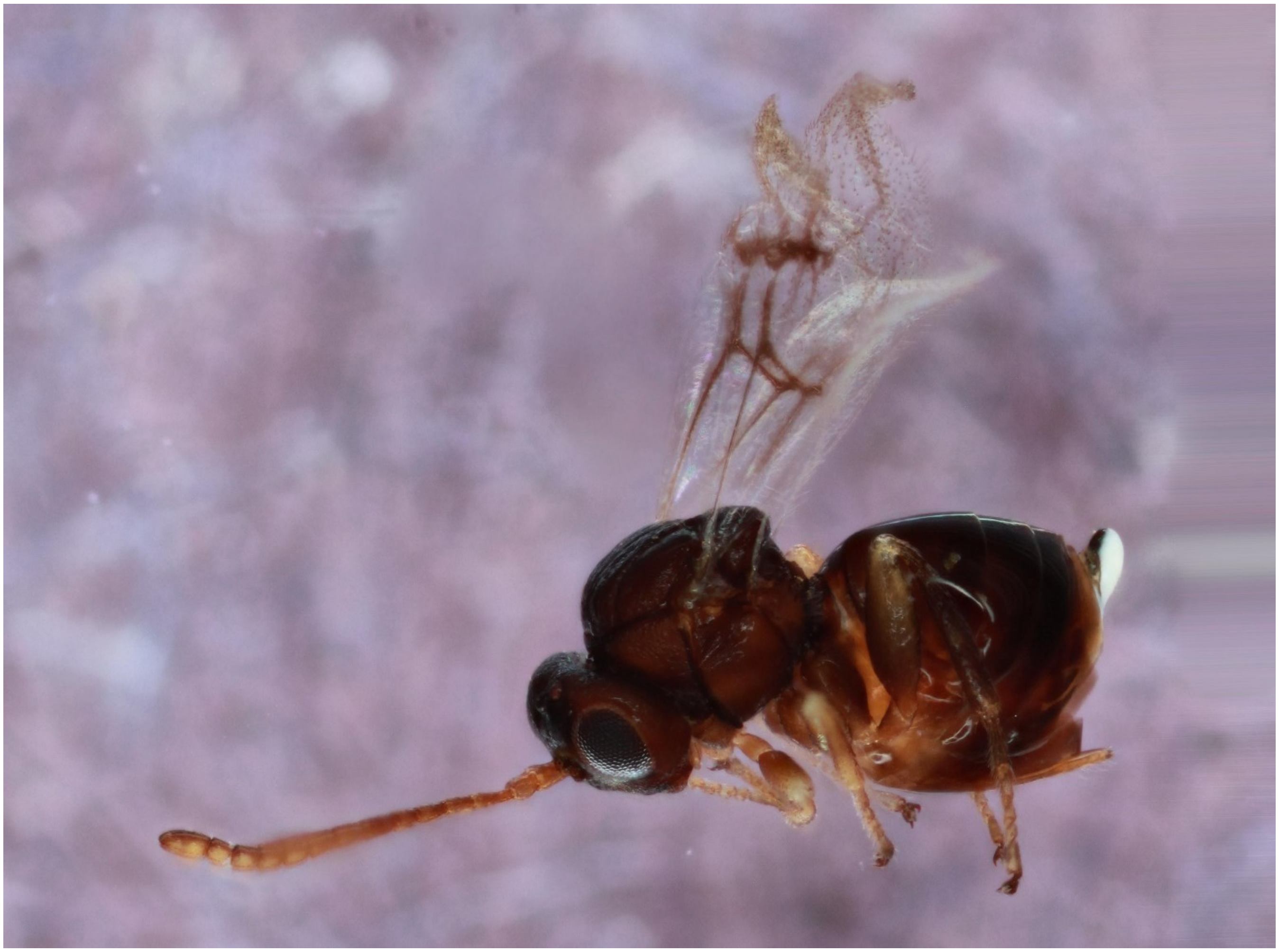

*Andricus kingi* - Wasp: Ak_1

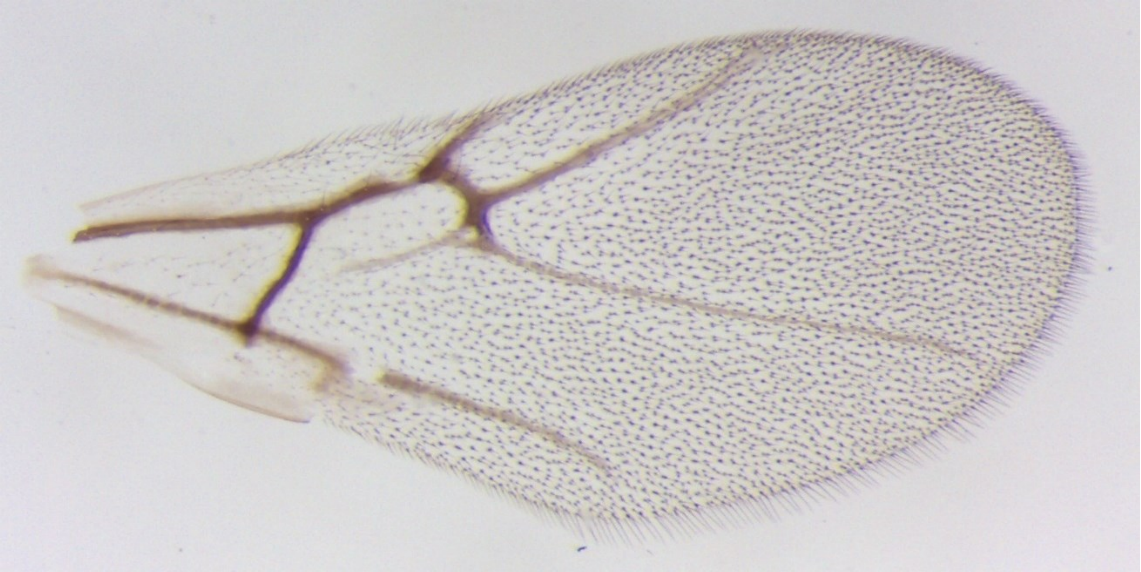

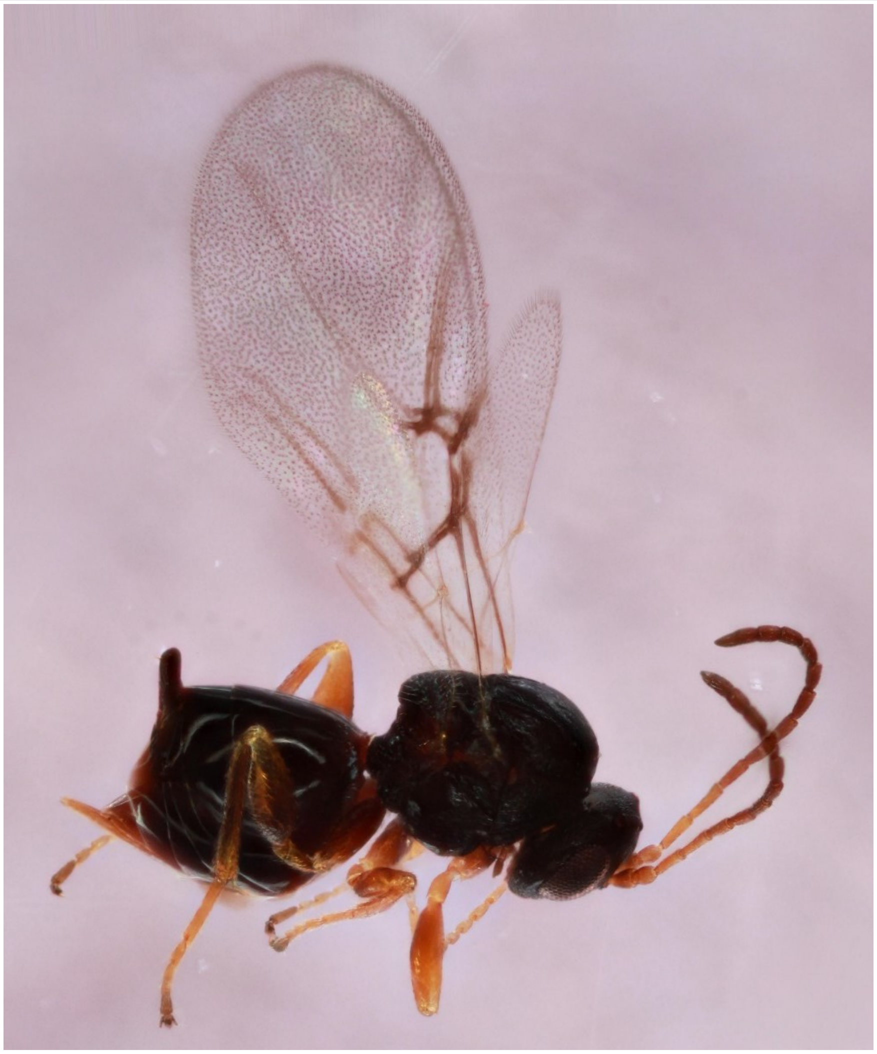

*Andricus parmula* - Wasp: 1480-136-1

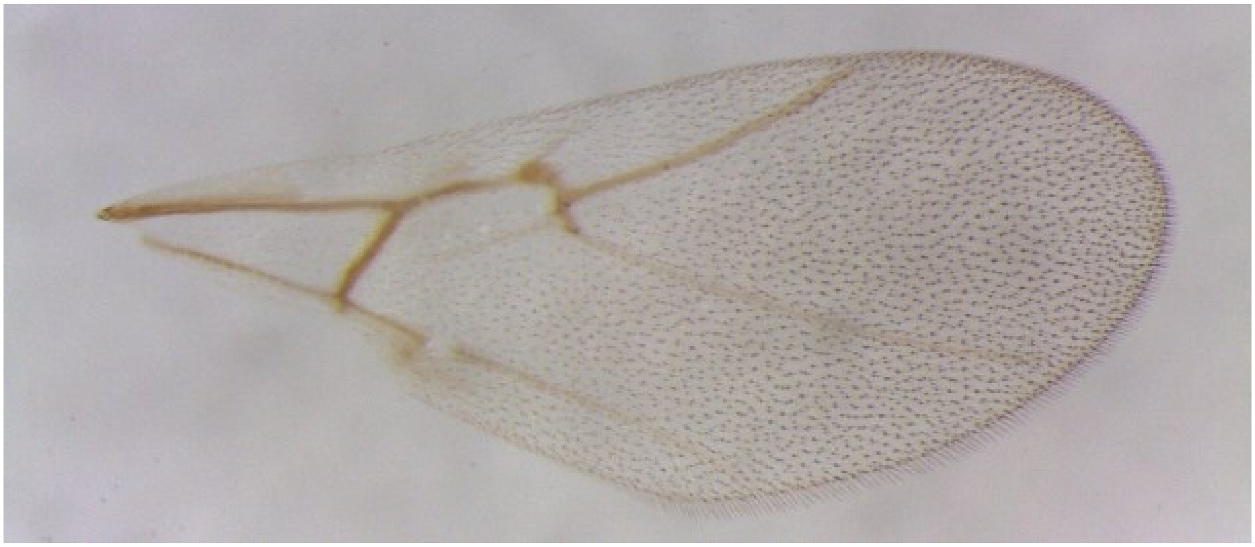

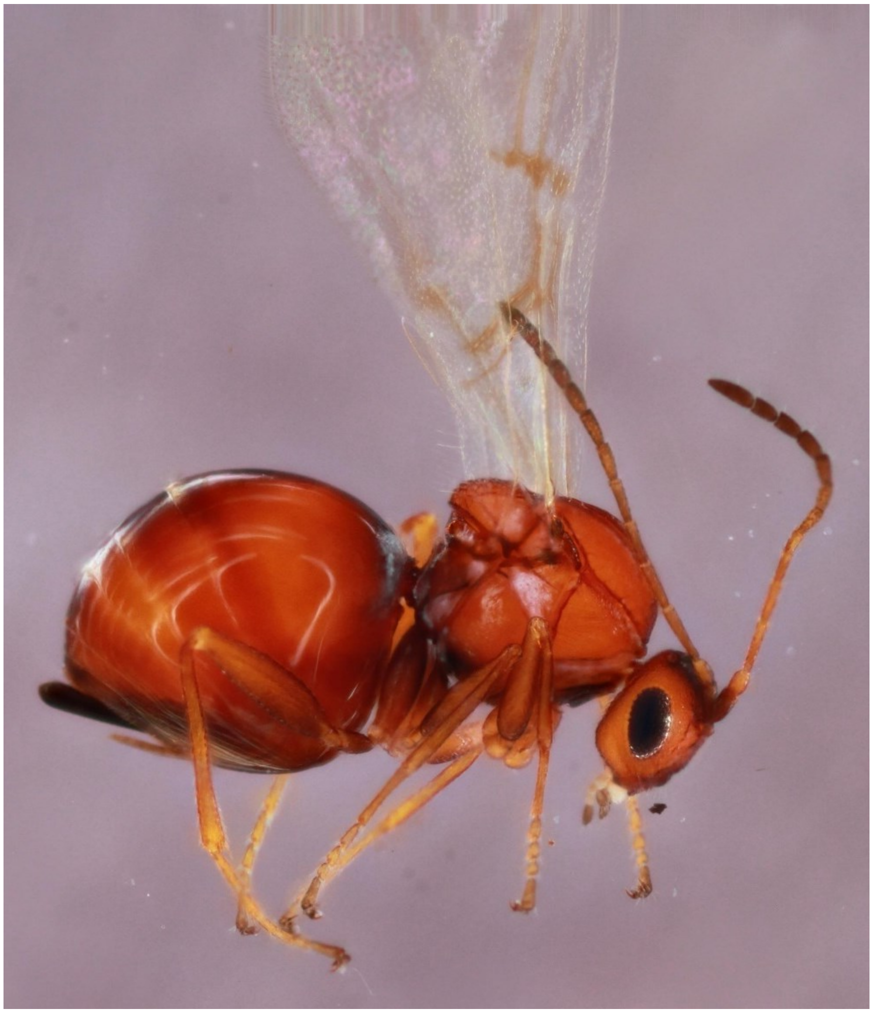

*Andricus pedicellatum* - Wasp: Dp2

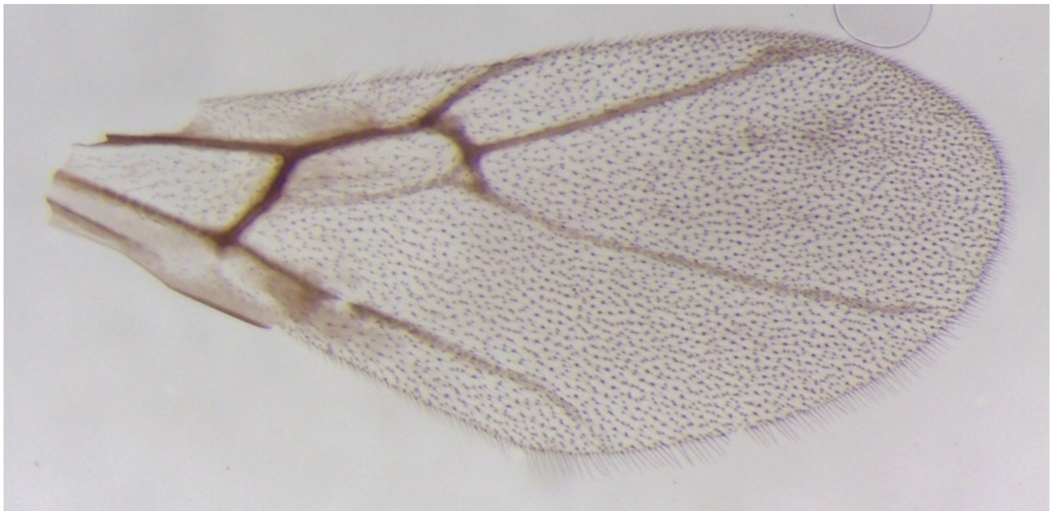

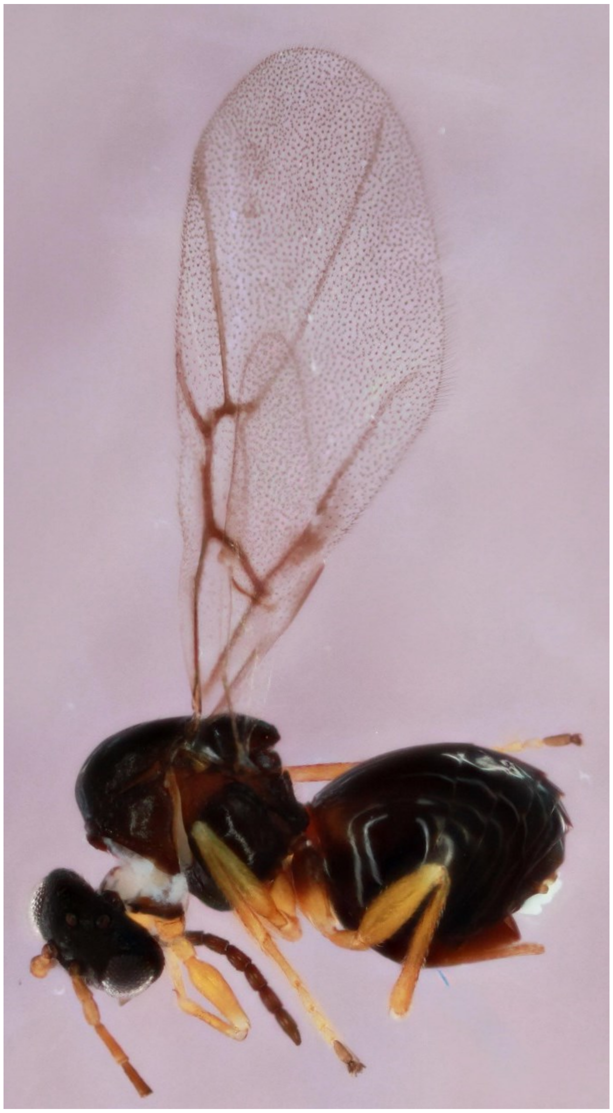

*Andricus pisiformis* - Wasp: 723-037-2

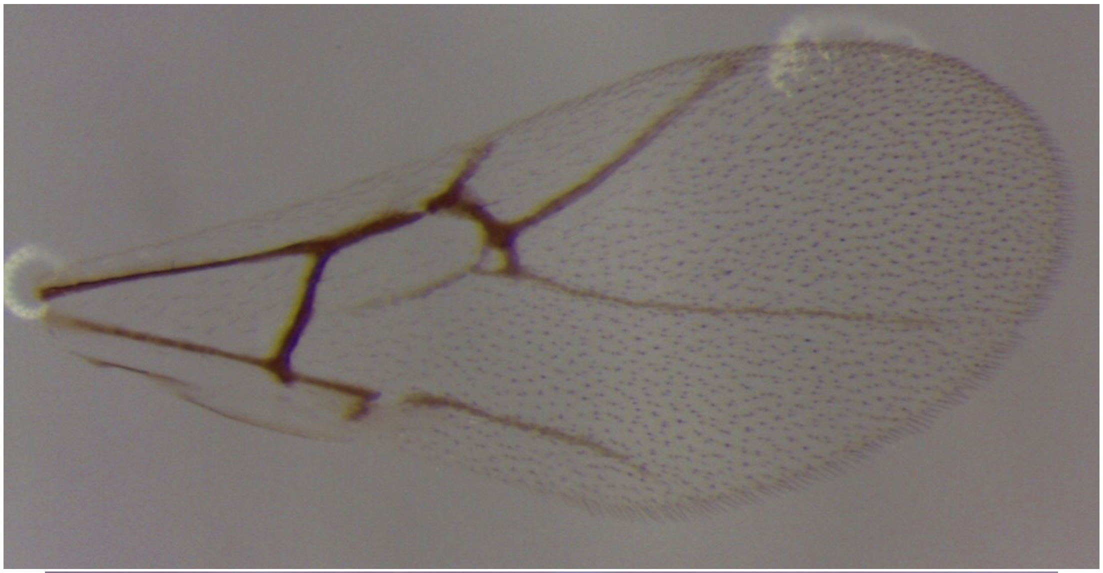

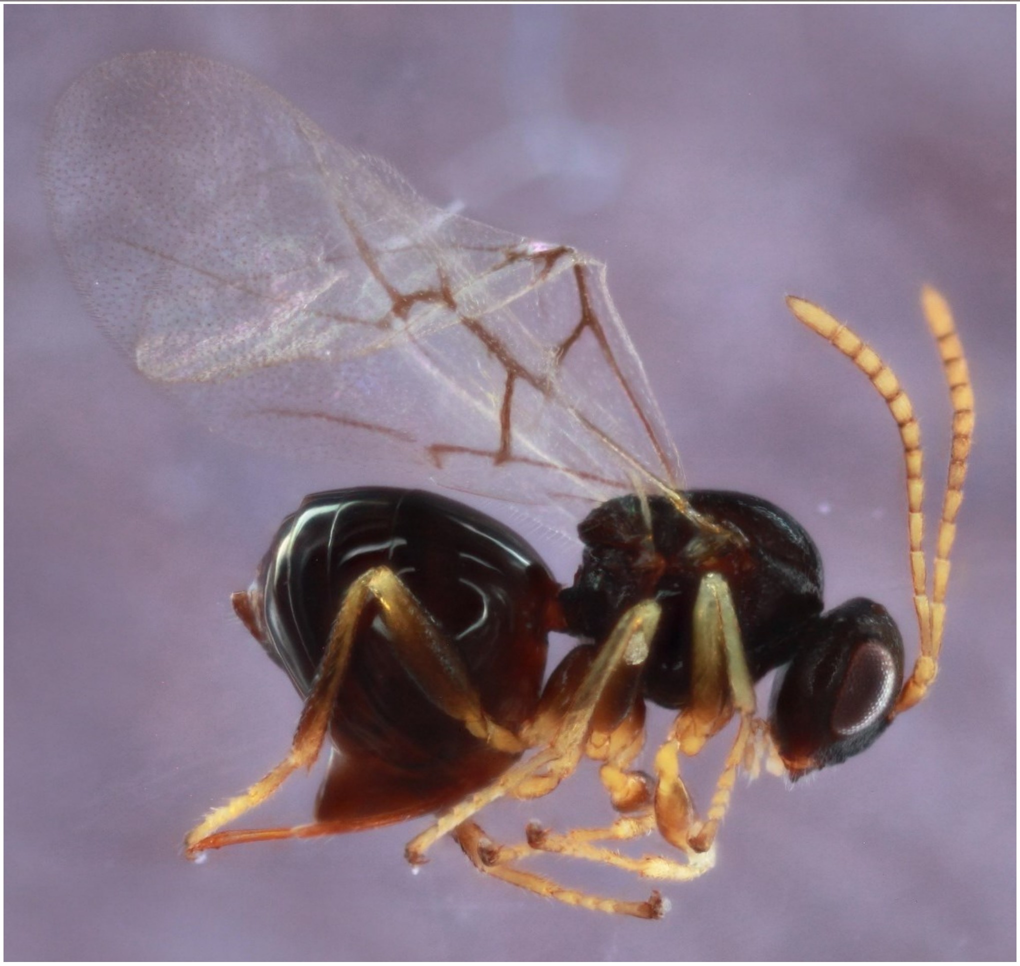

*Andricus quercuscalifornicus* - Wasp: 1470-130-5

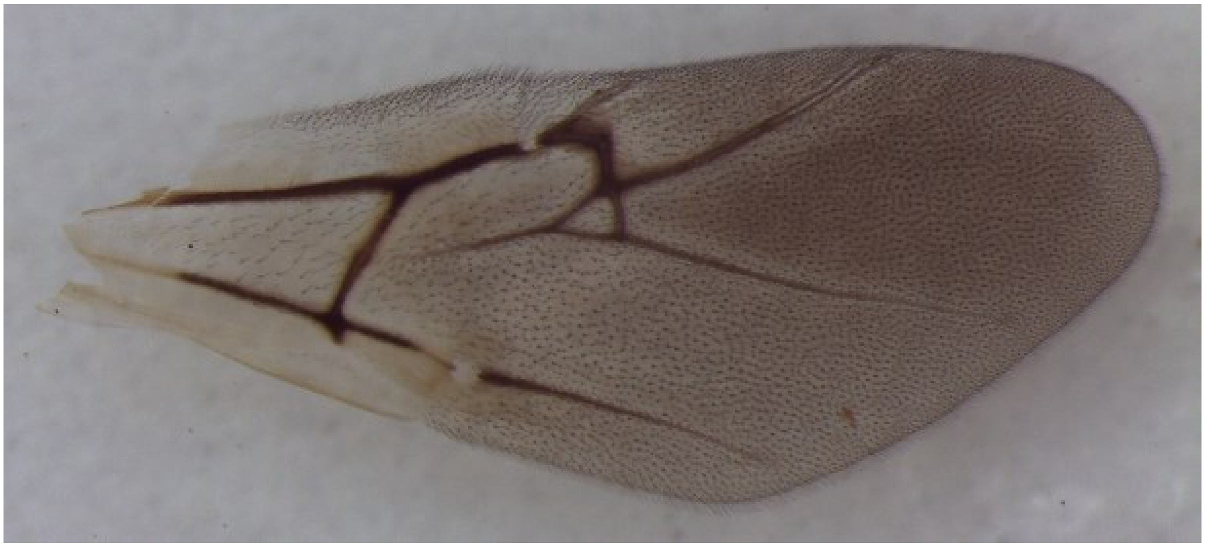

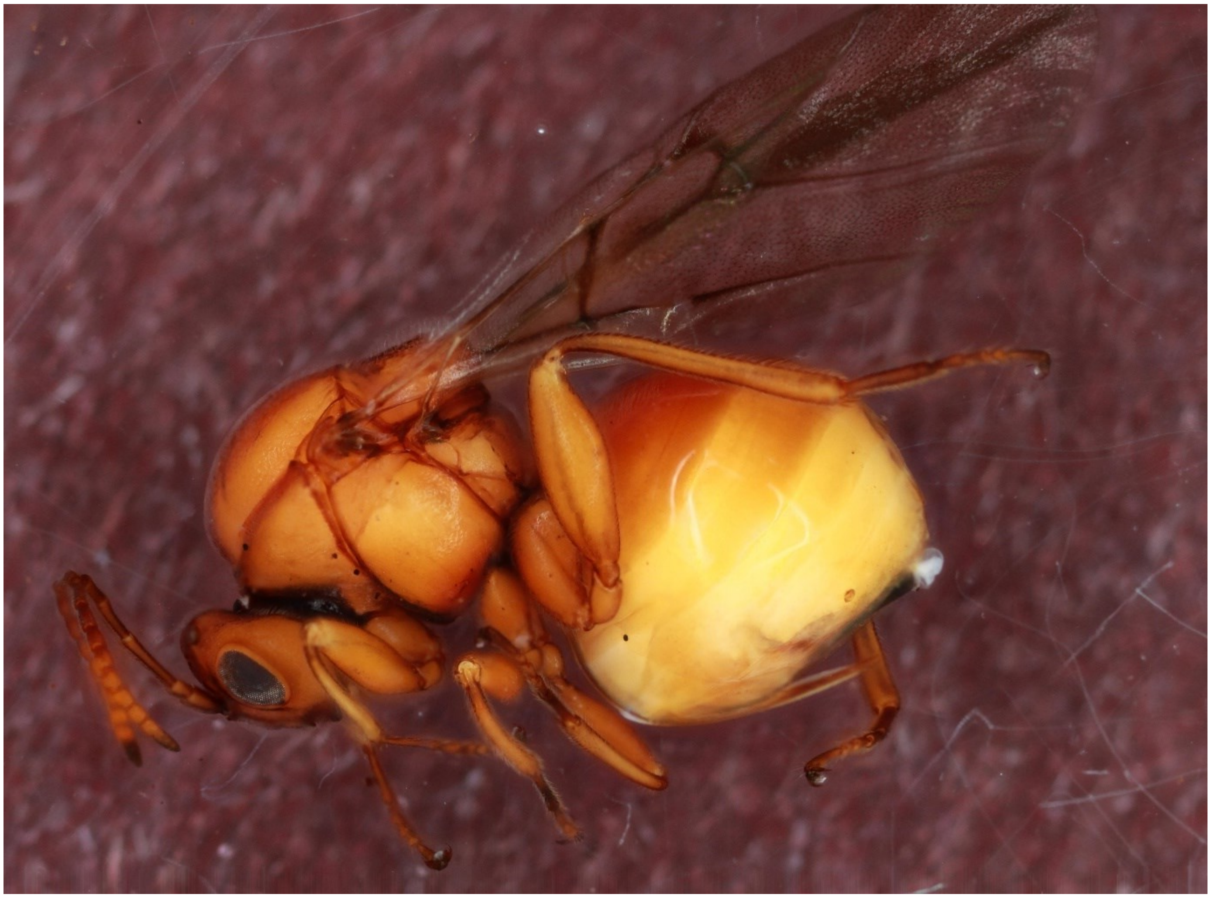

*Andricus quercusostensackenii* - Wasp: 1617-166-1

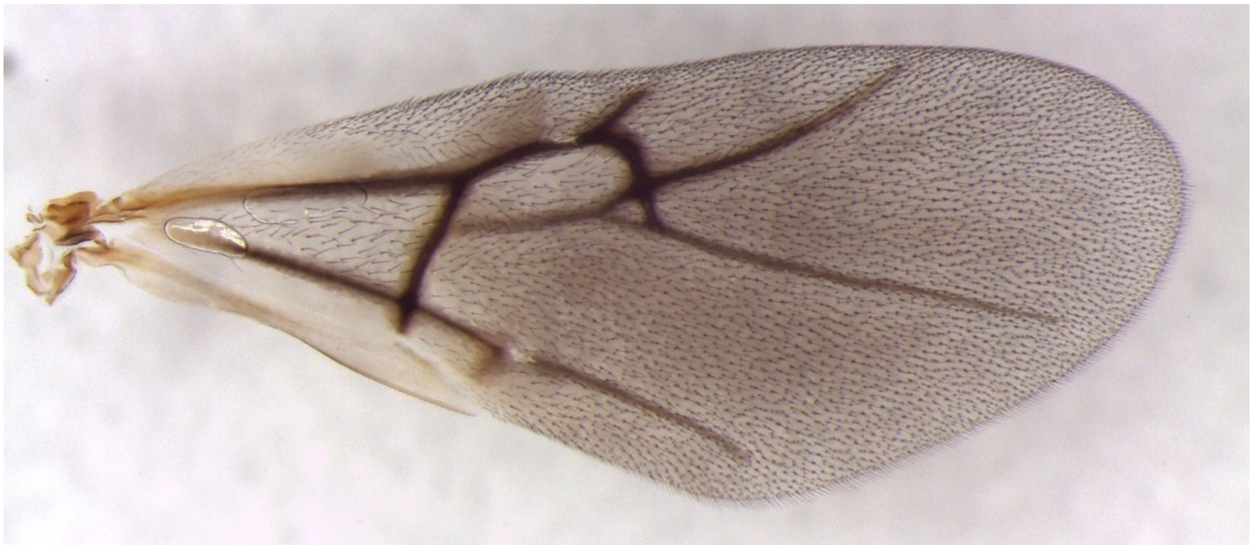

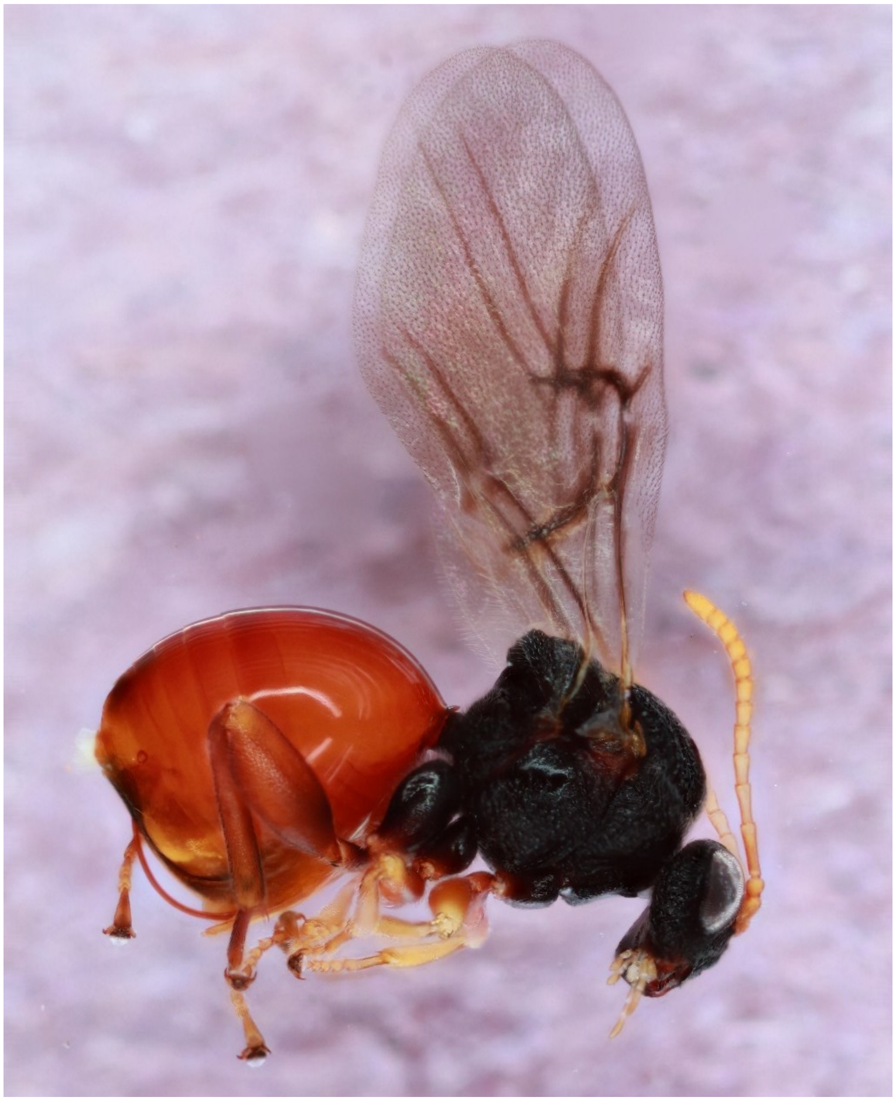

*Andricus quercuspetiolicola* - Wasp: 880-042-7A

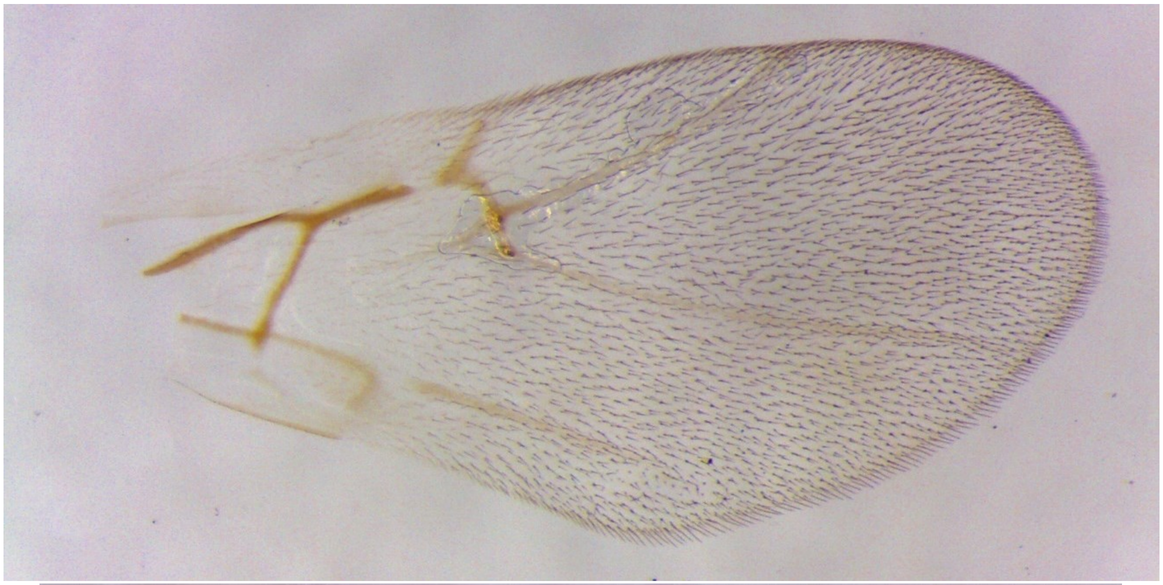

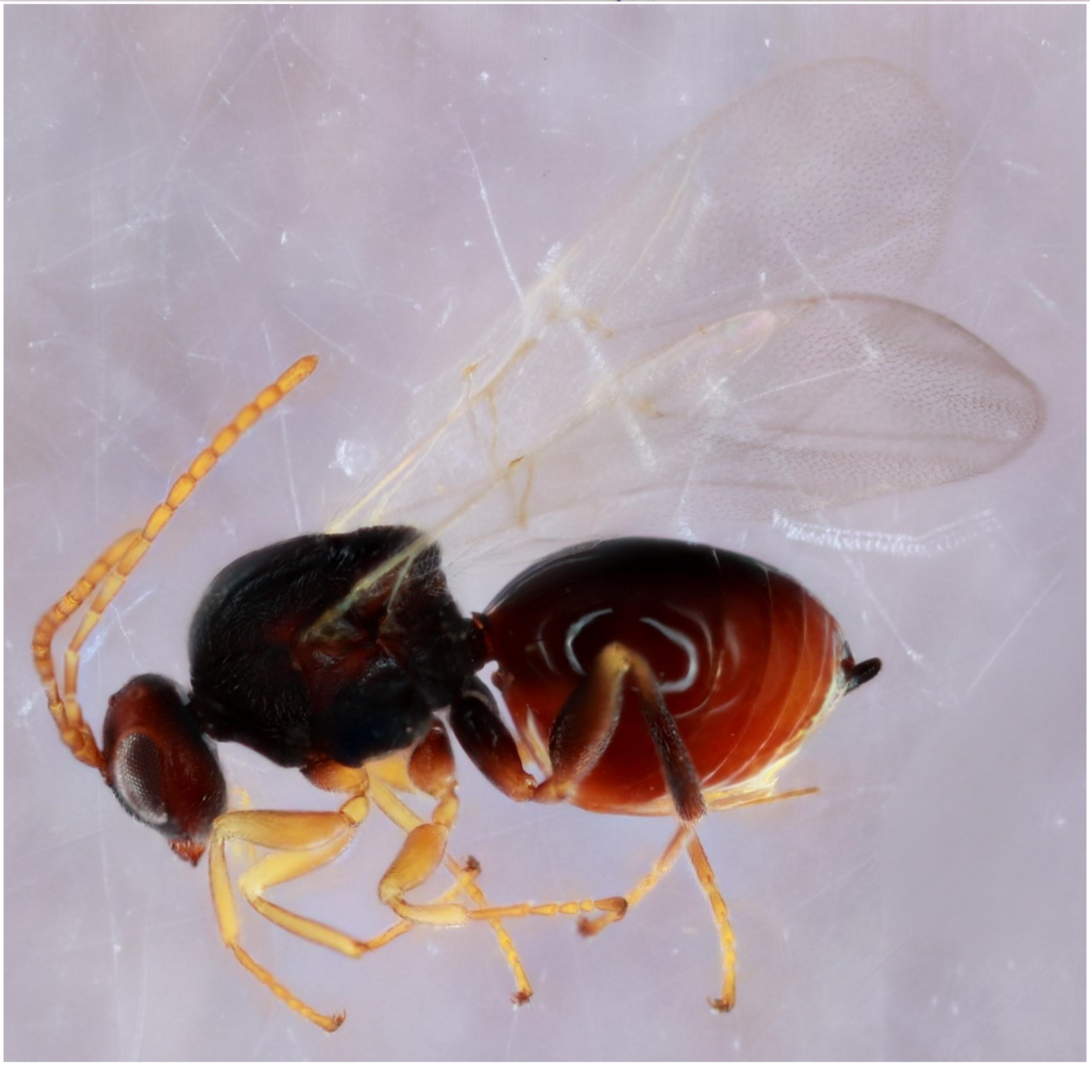

*Andricus quercusstrobilanus* - Wasp: 500-011-033

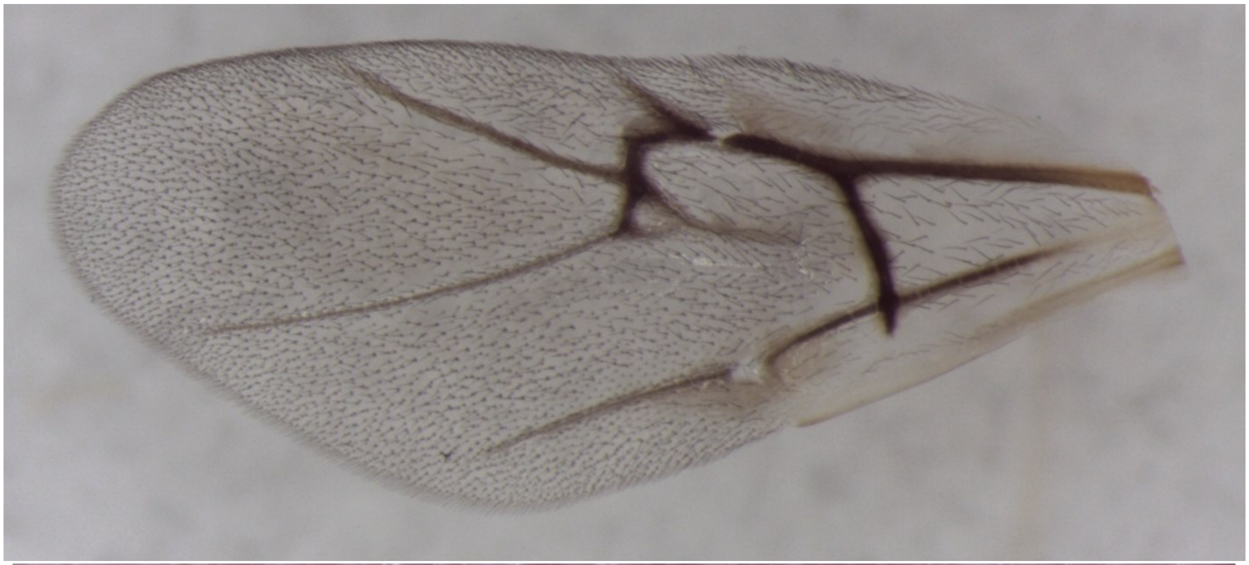

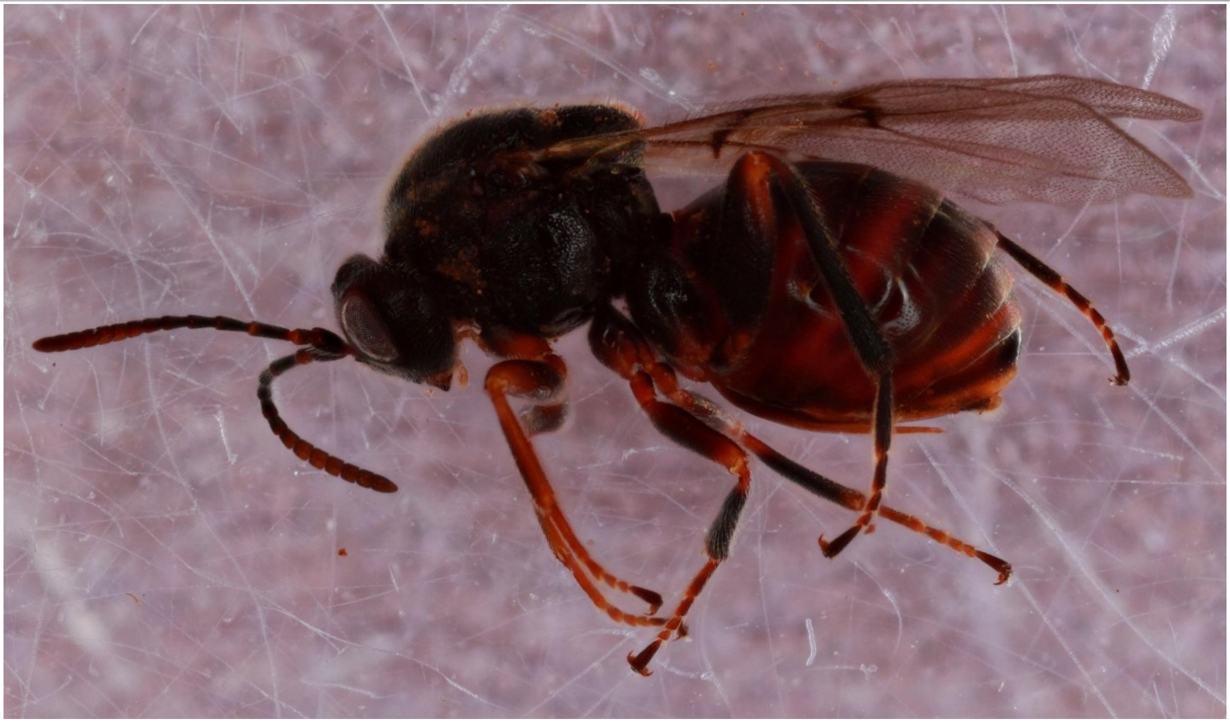

*Andricus quercusutriculus* - Wasp: 778-040-001

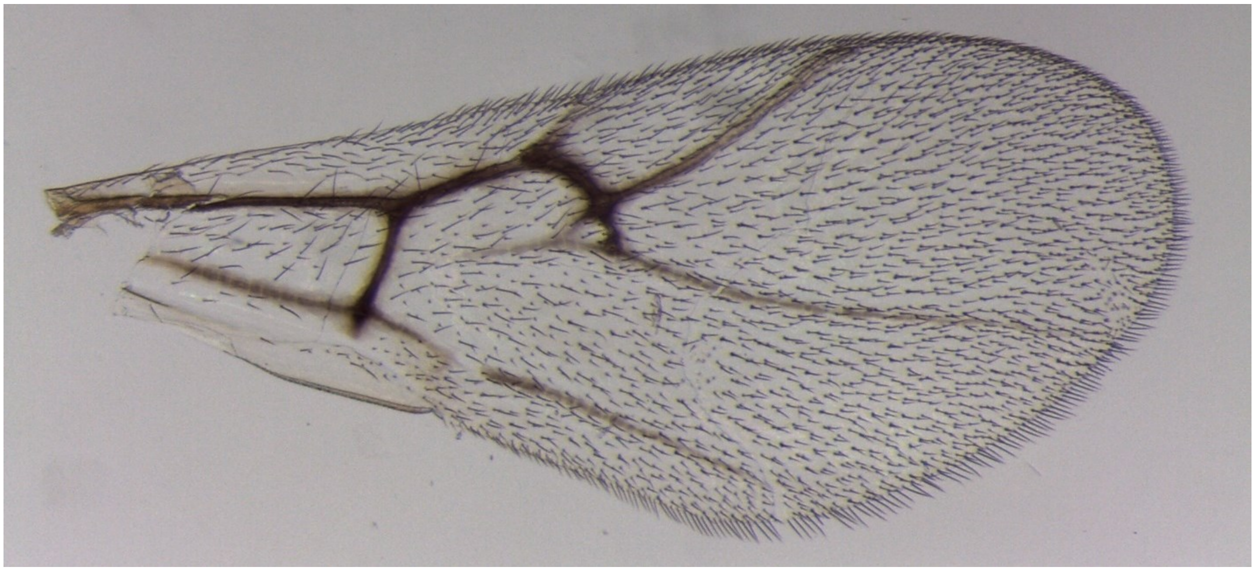

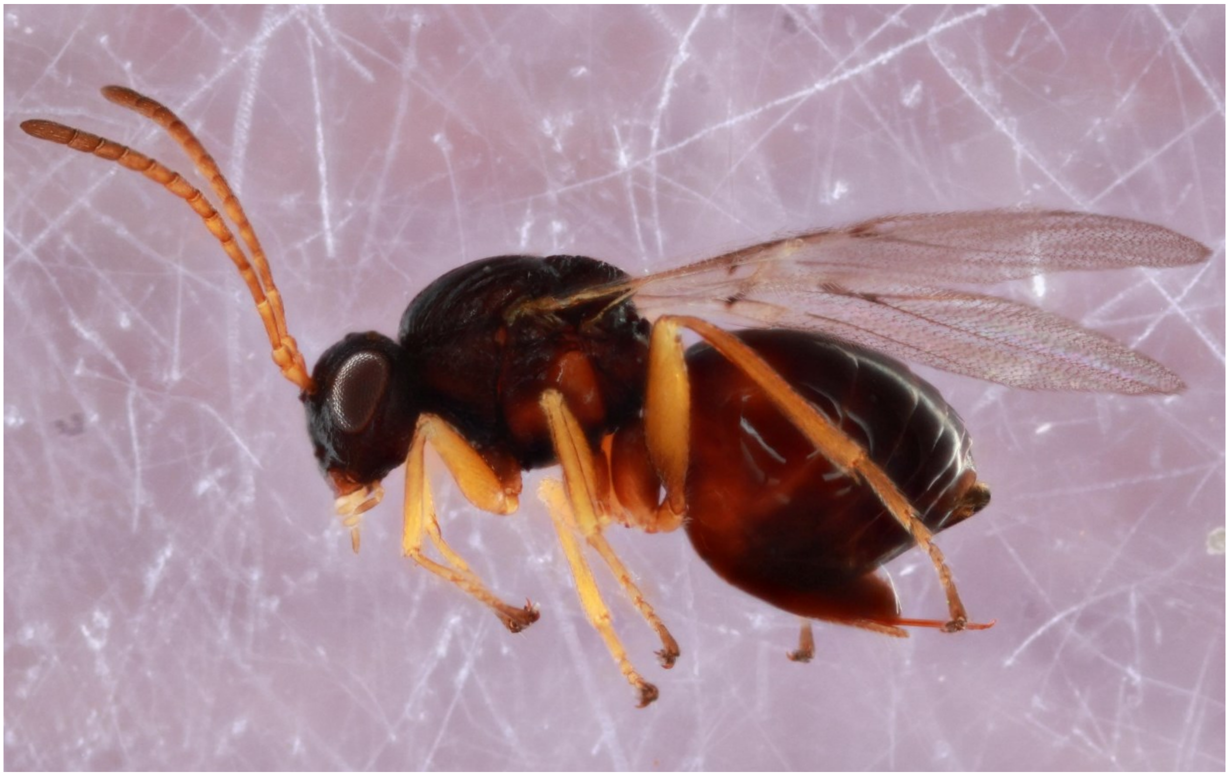

*Atrusca bella* - Wasp: 1688-212-3

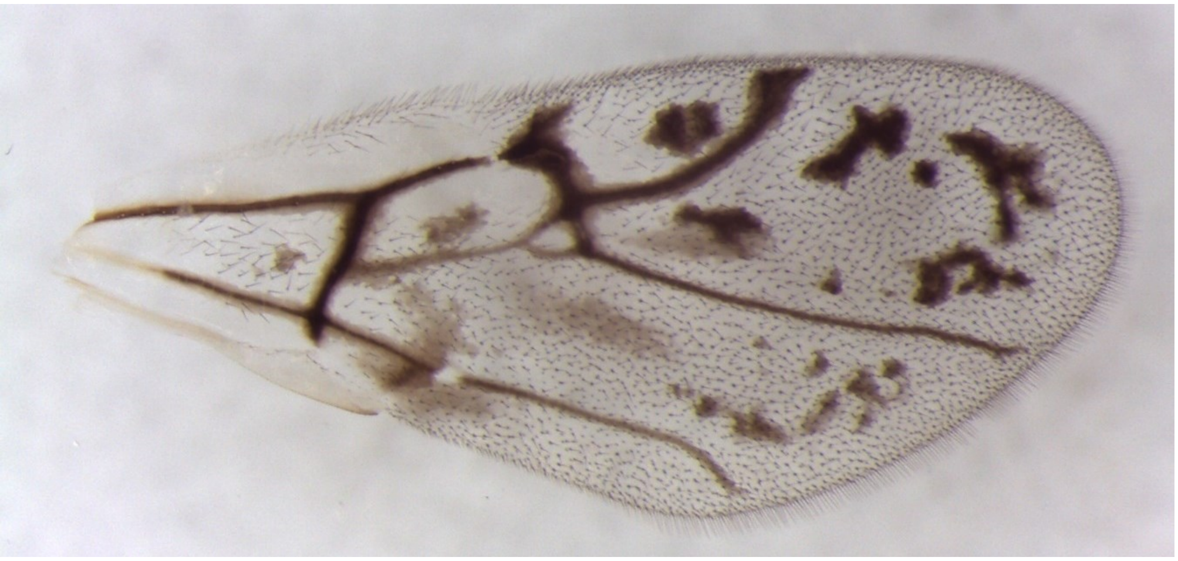

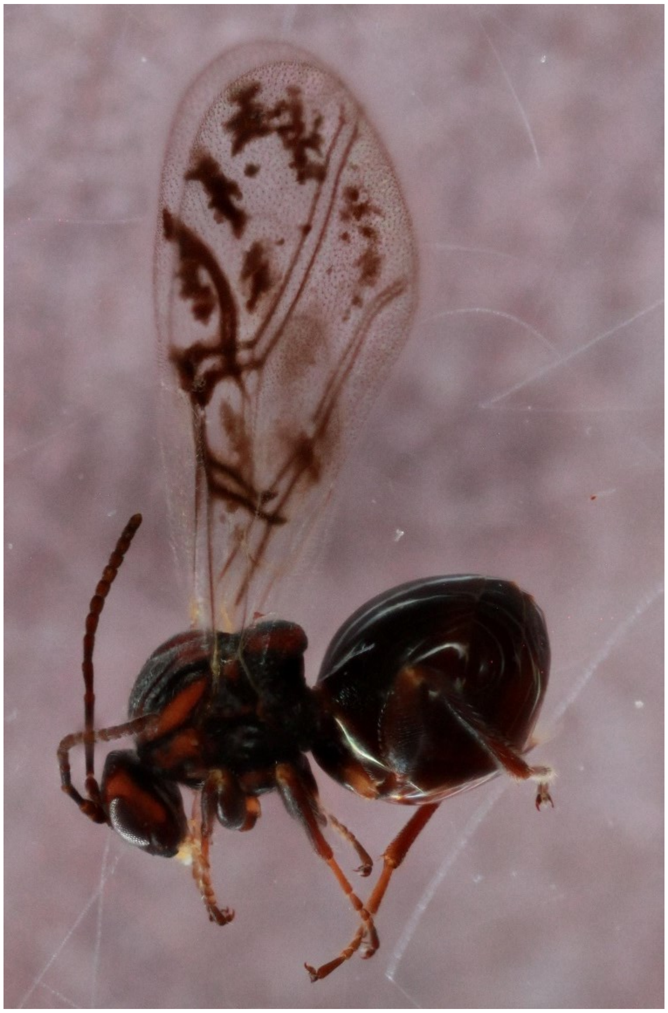

*Atrusca brevipennata* - Wasp: 1645-169-1A

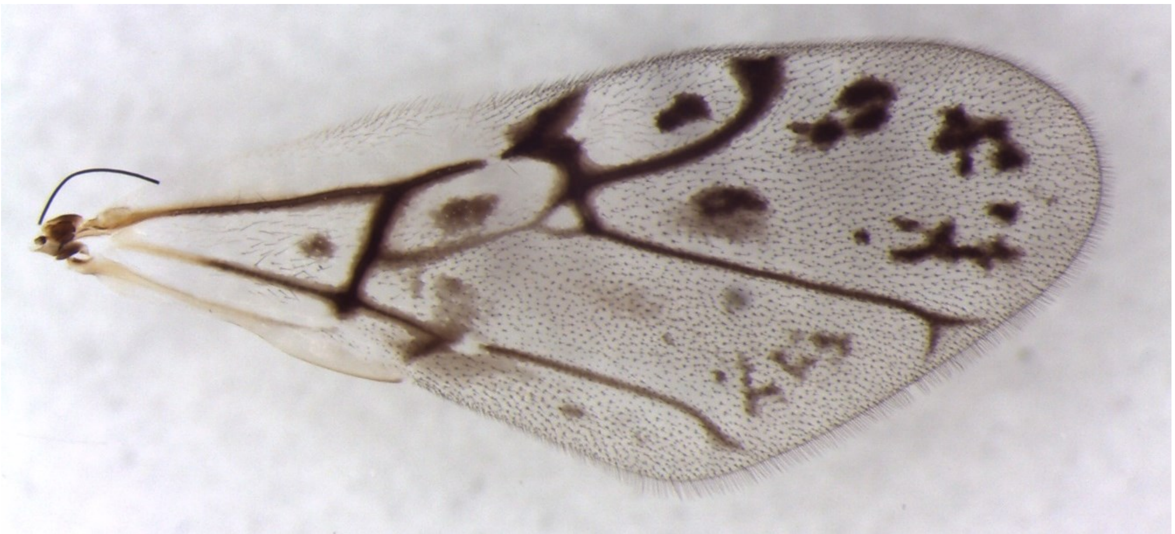

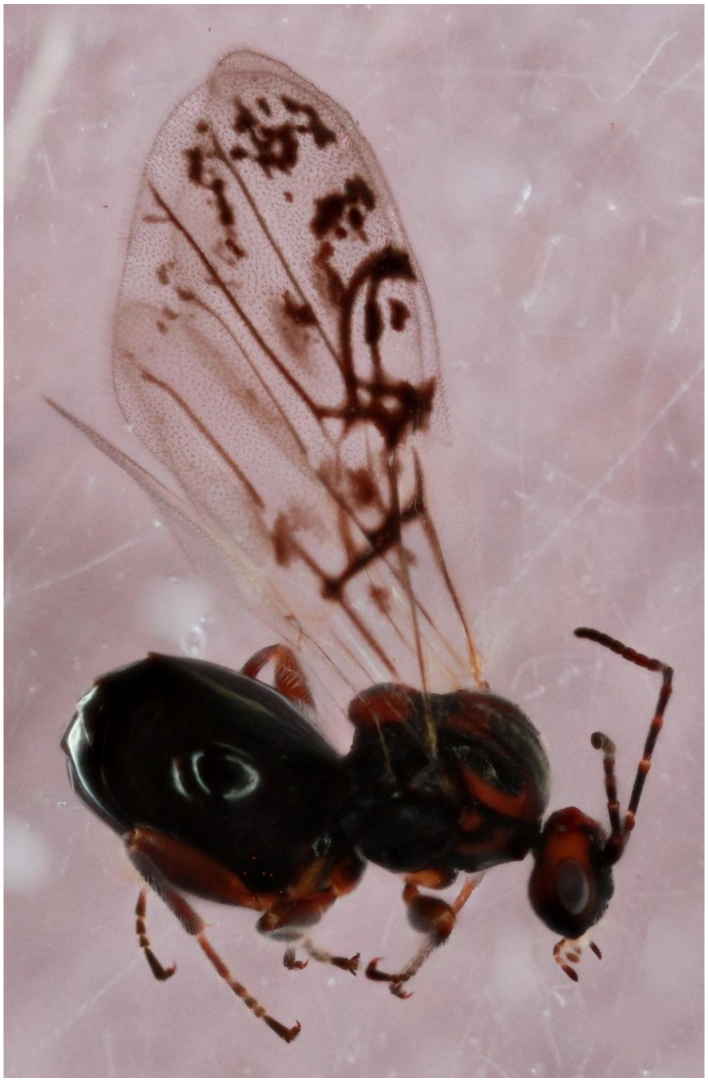

*Atrusca quercuscentricola* - Wasp: 1092-065-1

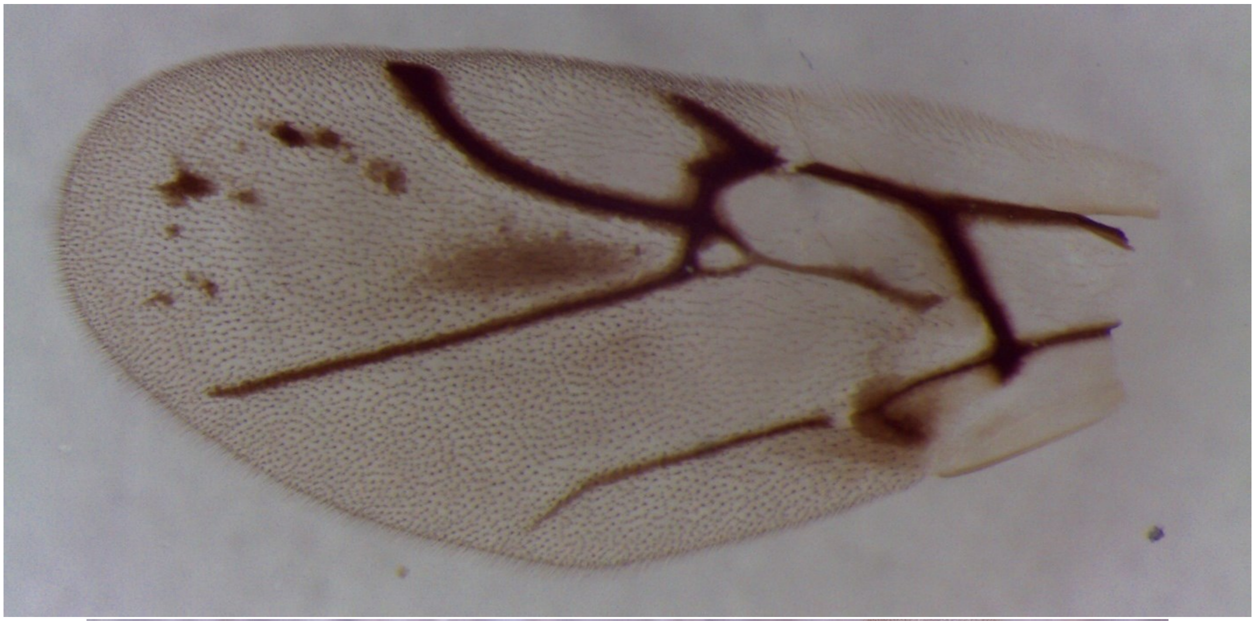

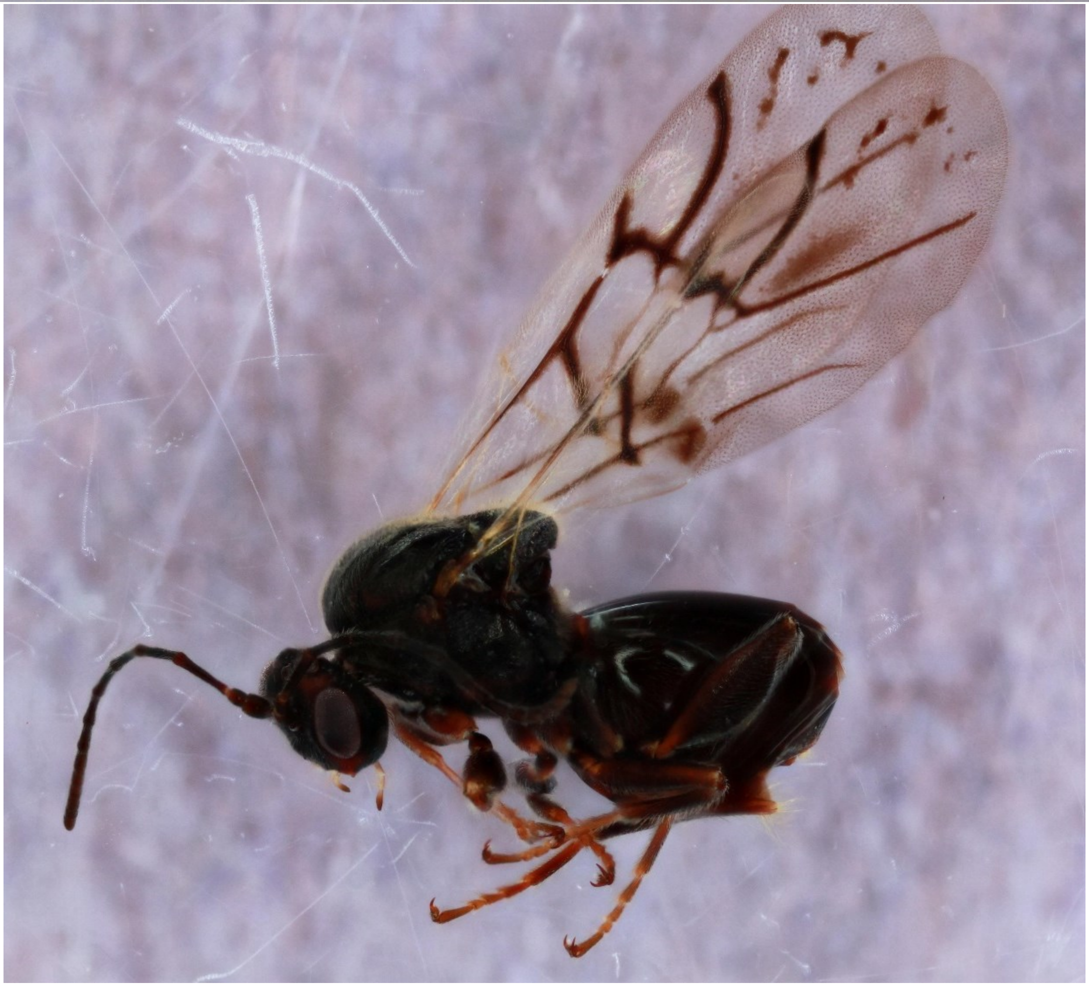

*Bassettia pallida* - Wasp: LZ6437_2

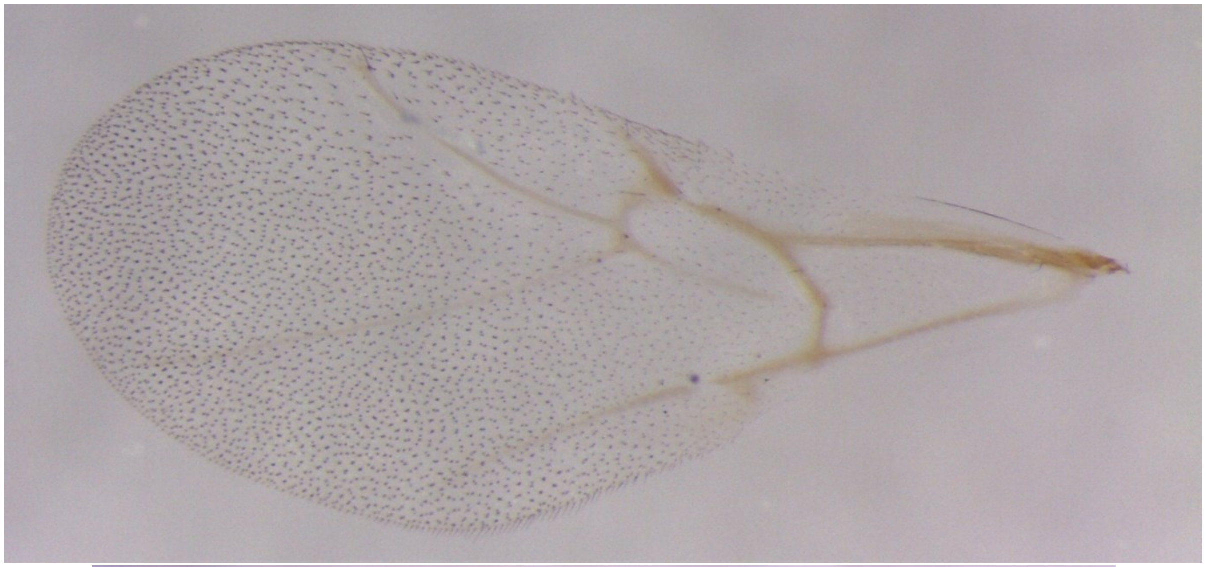

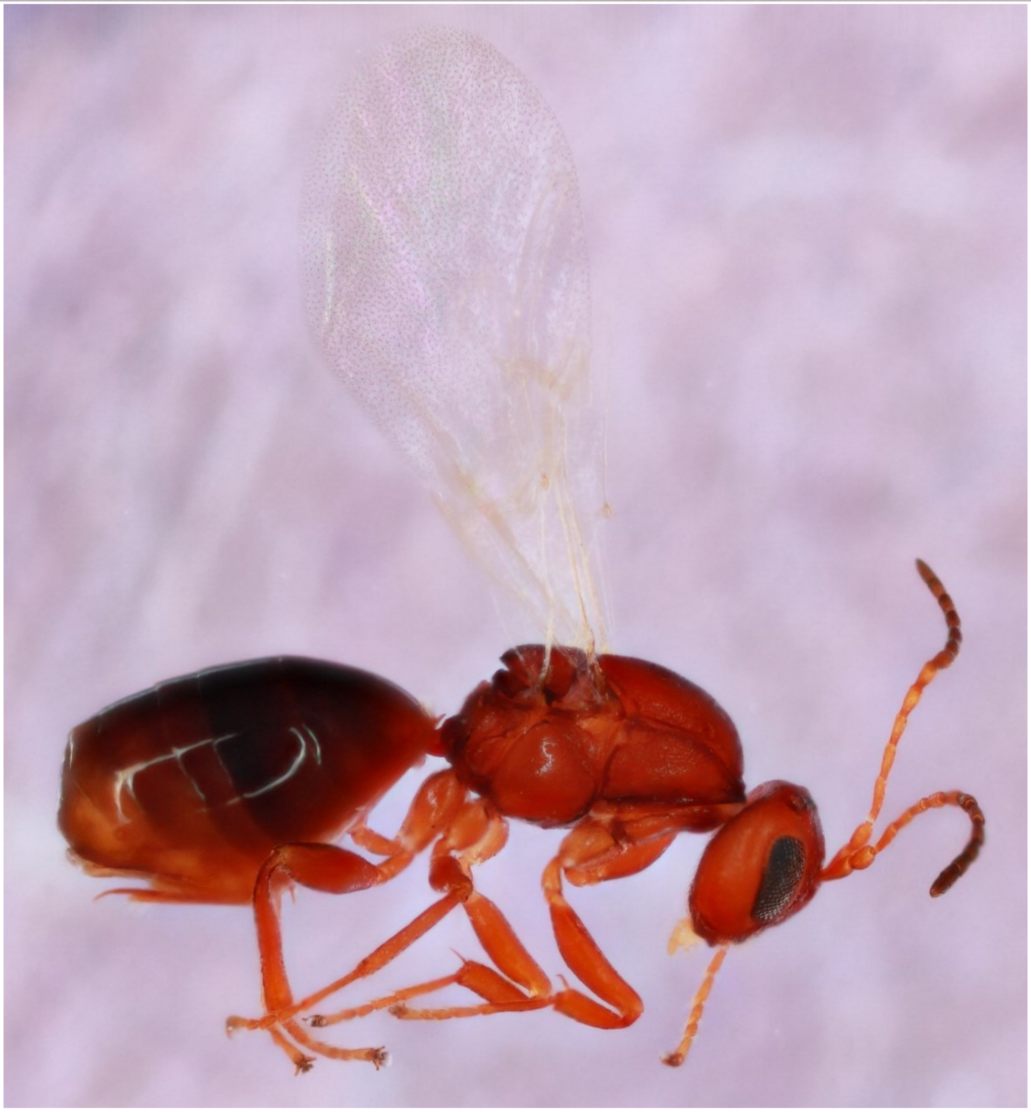

*Belonocnema fossoria* - Wasp: LZ_4370

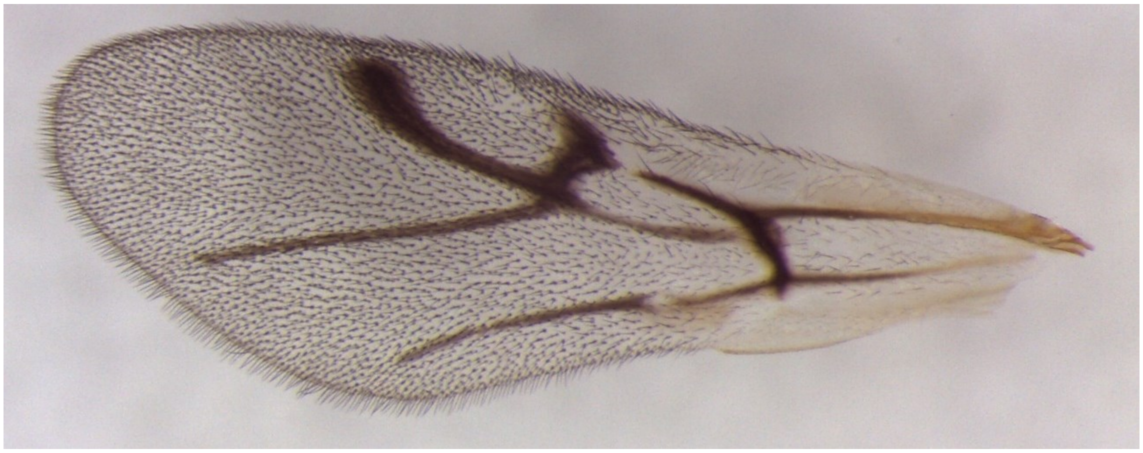

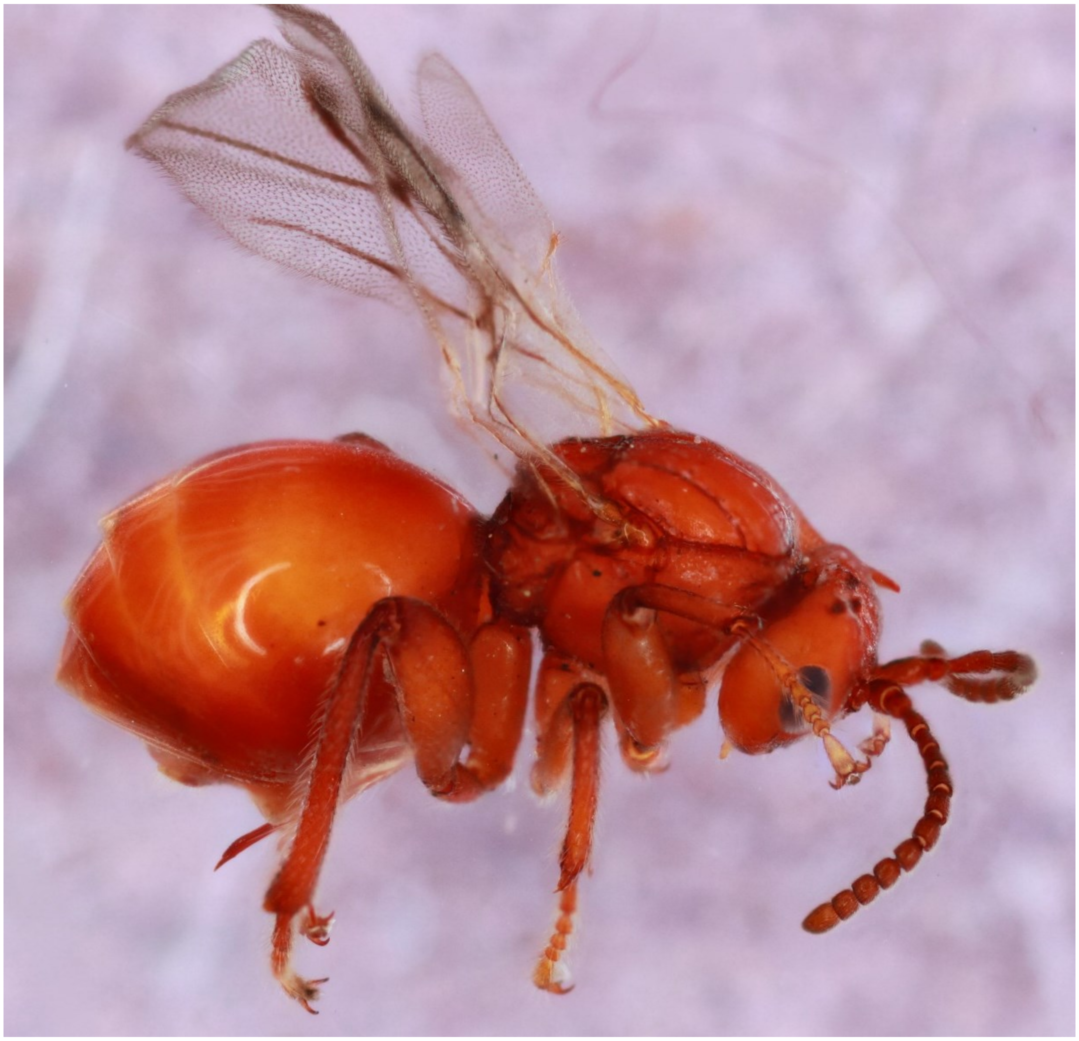

*Belonocnema treatae* - Wasp: LZ_4348

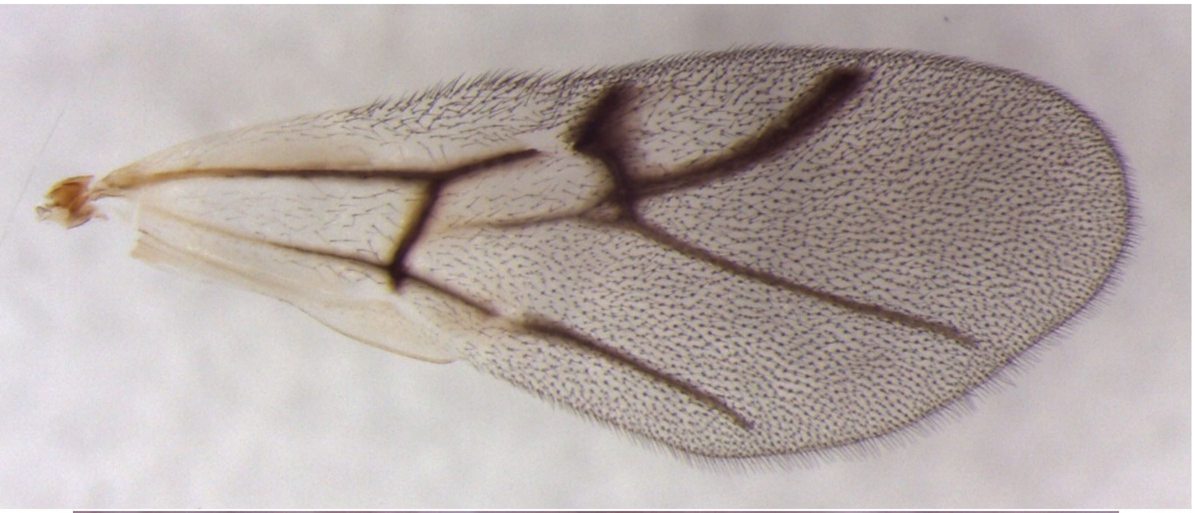

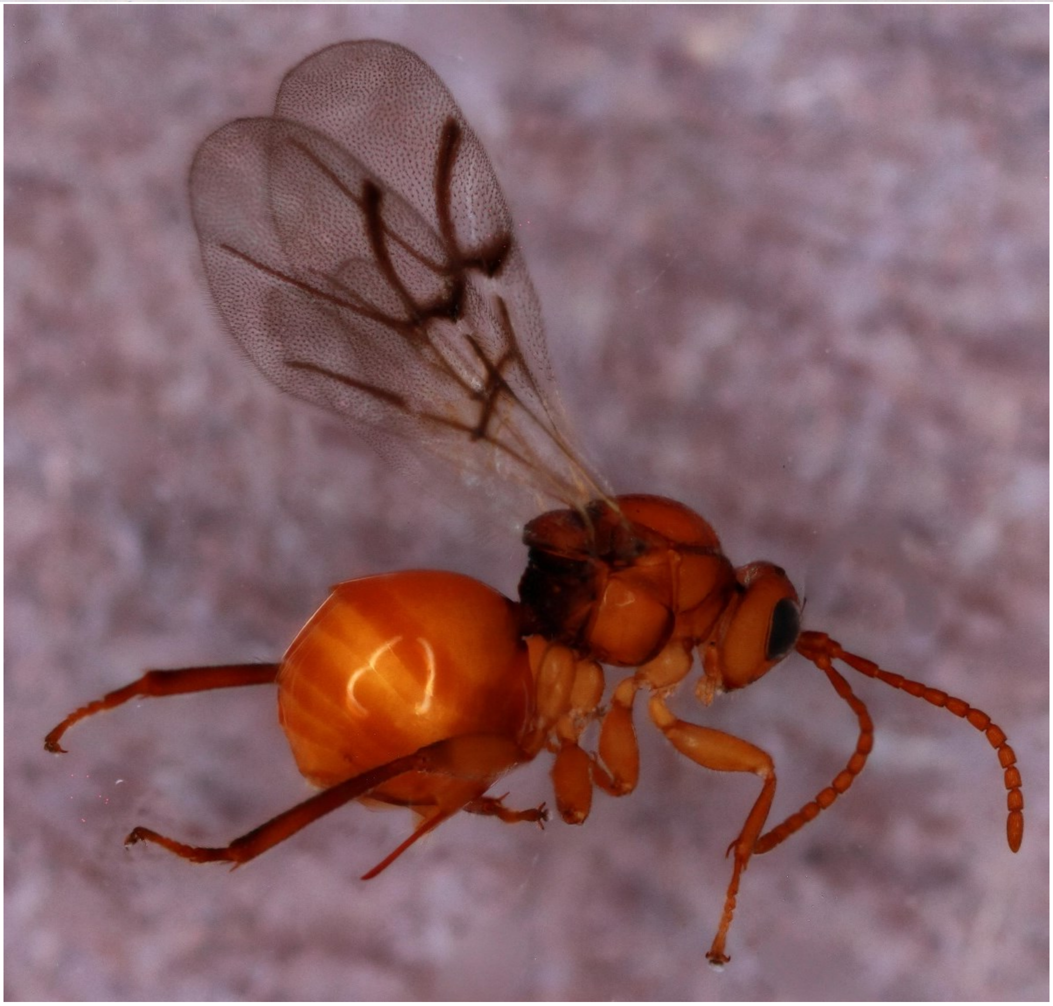

*Burnettweldia washingtonensis* - Wasp: Bc_1

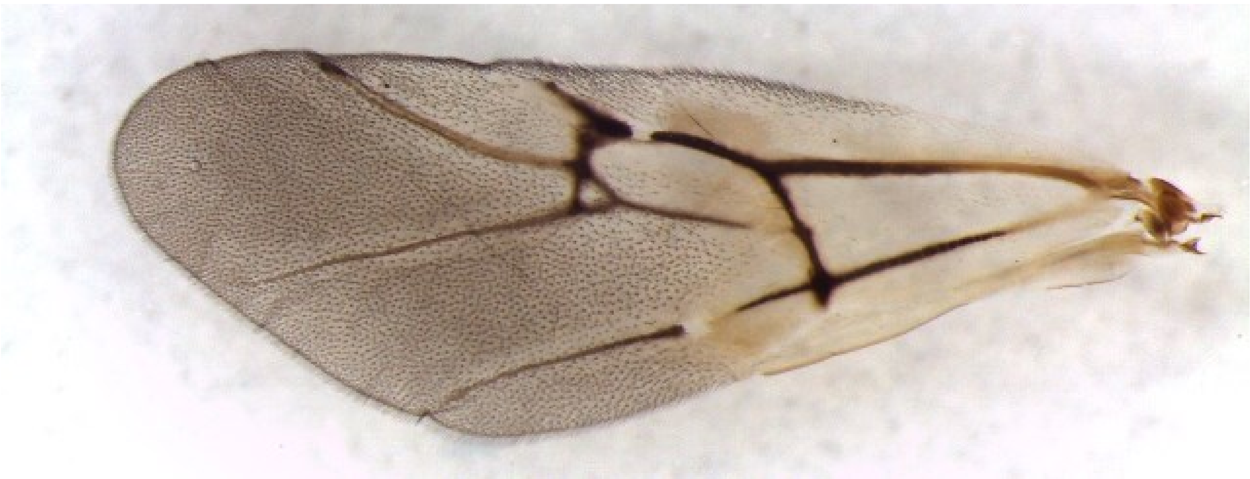

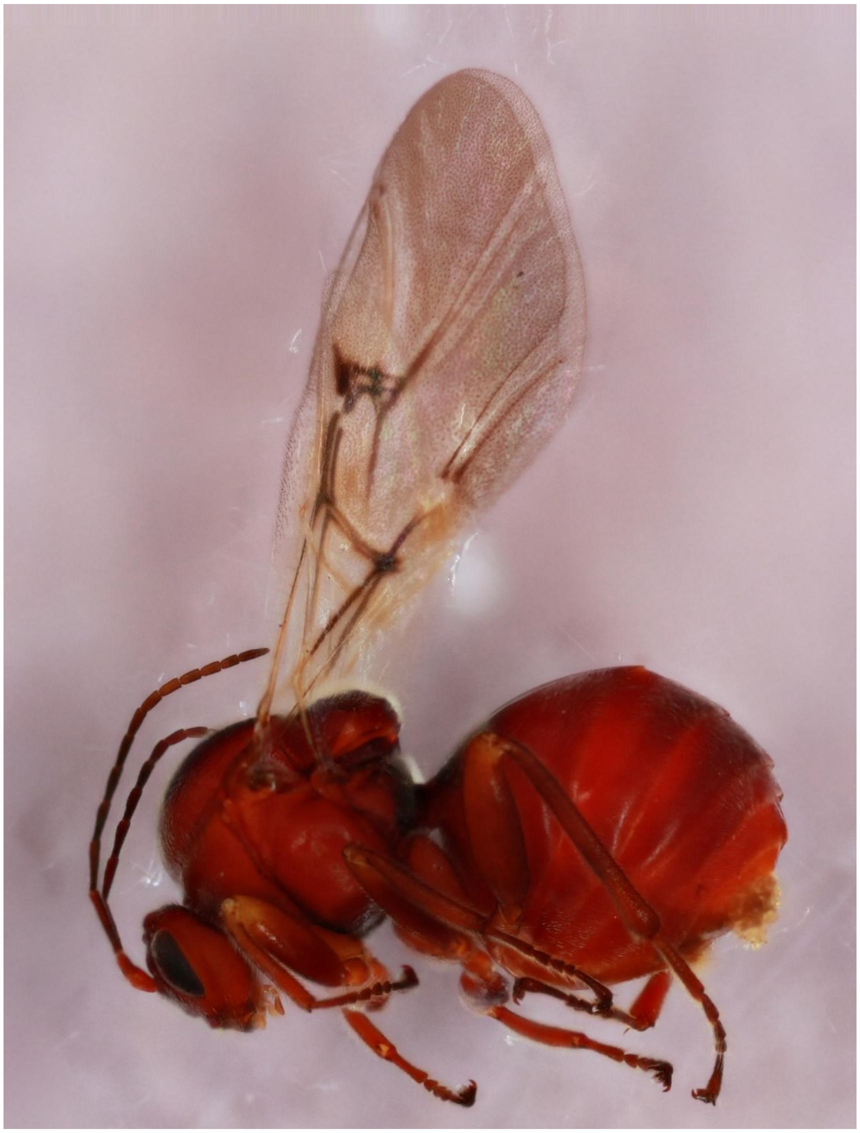

*Callirhytis attigua* - Wasp: 1579-154-10

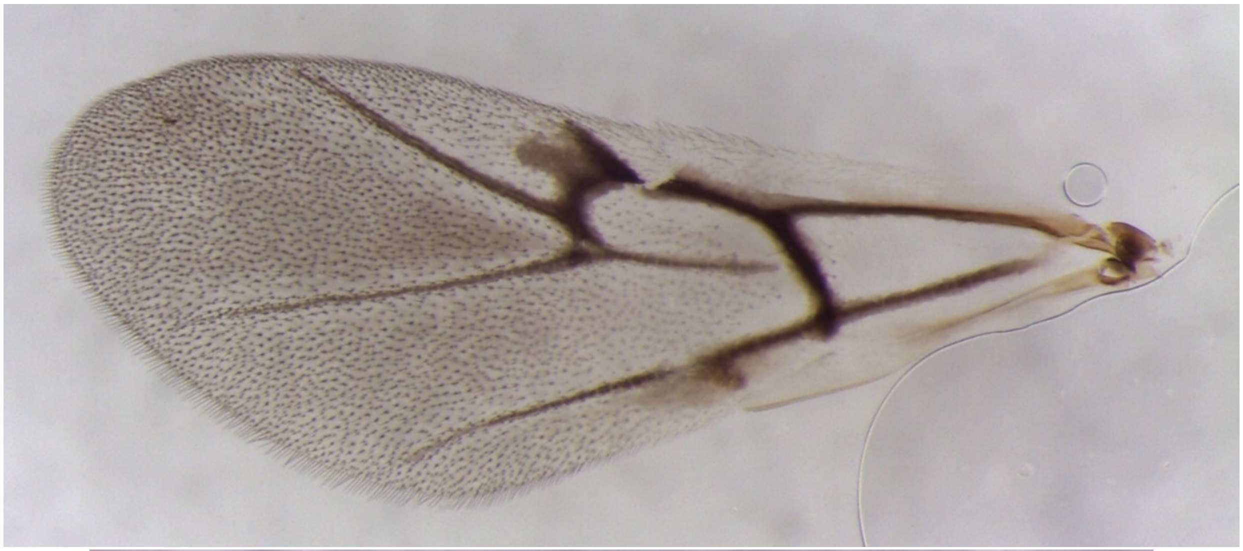

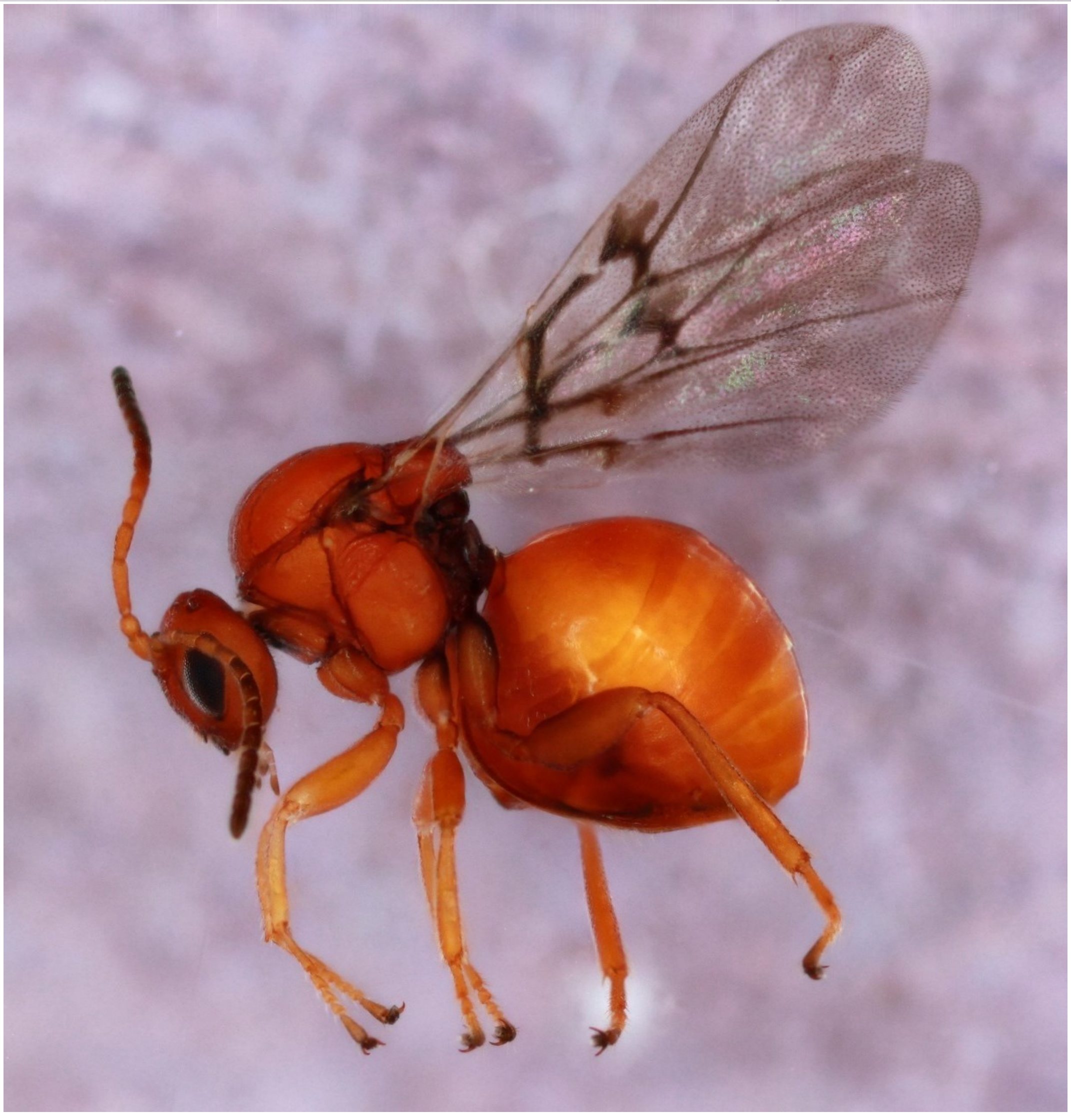

*Callirhytis exigua* - Wasp: 711-085-001A

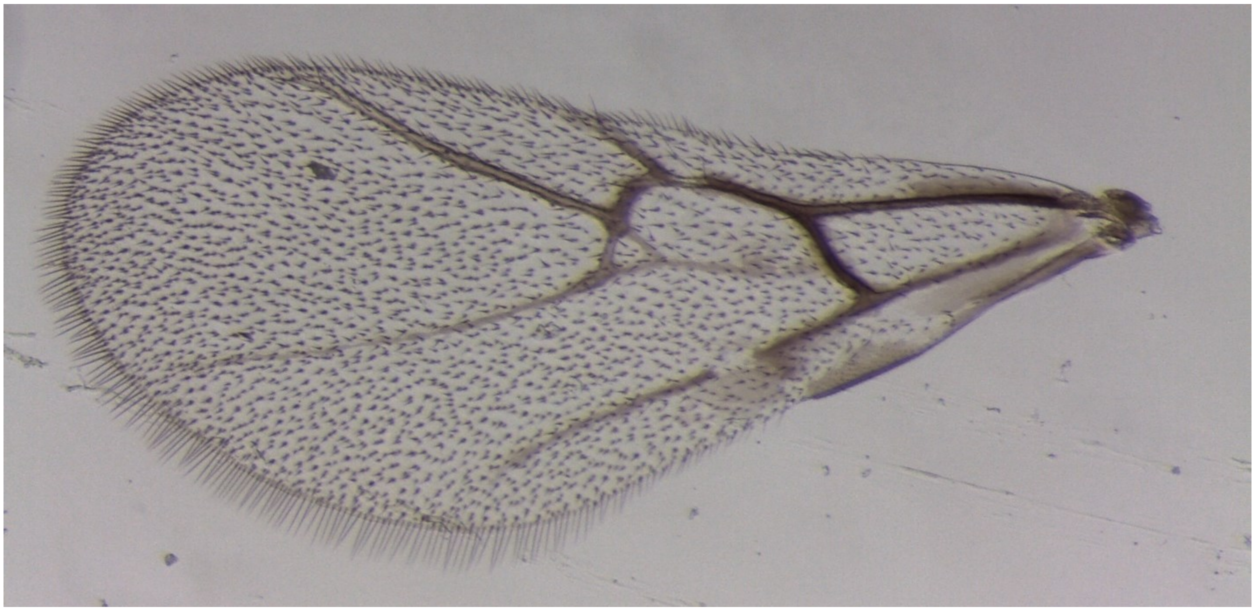

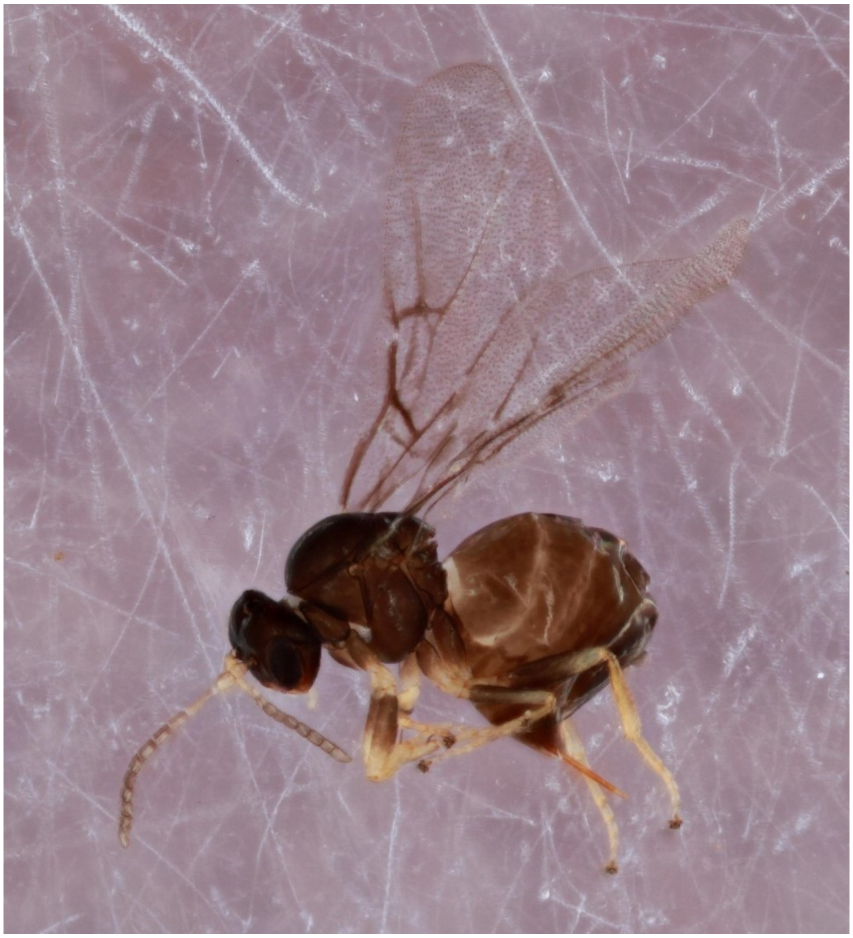

*Callirhytis flavipes* - Wasp: 881-013-1 (MALE)

(no wing picture)

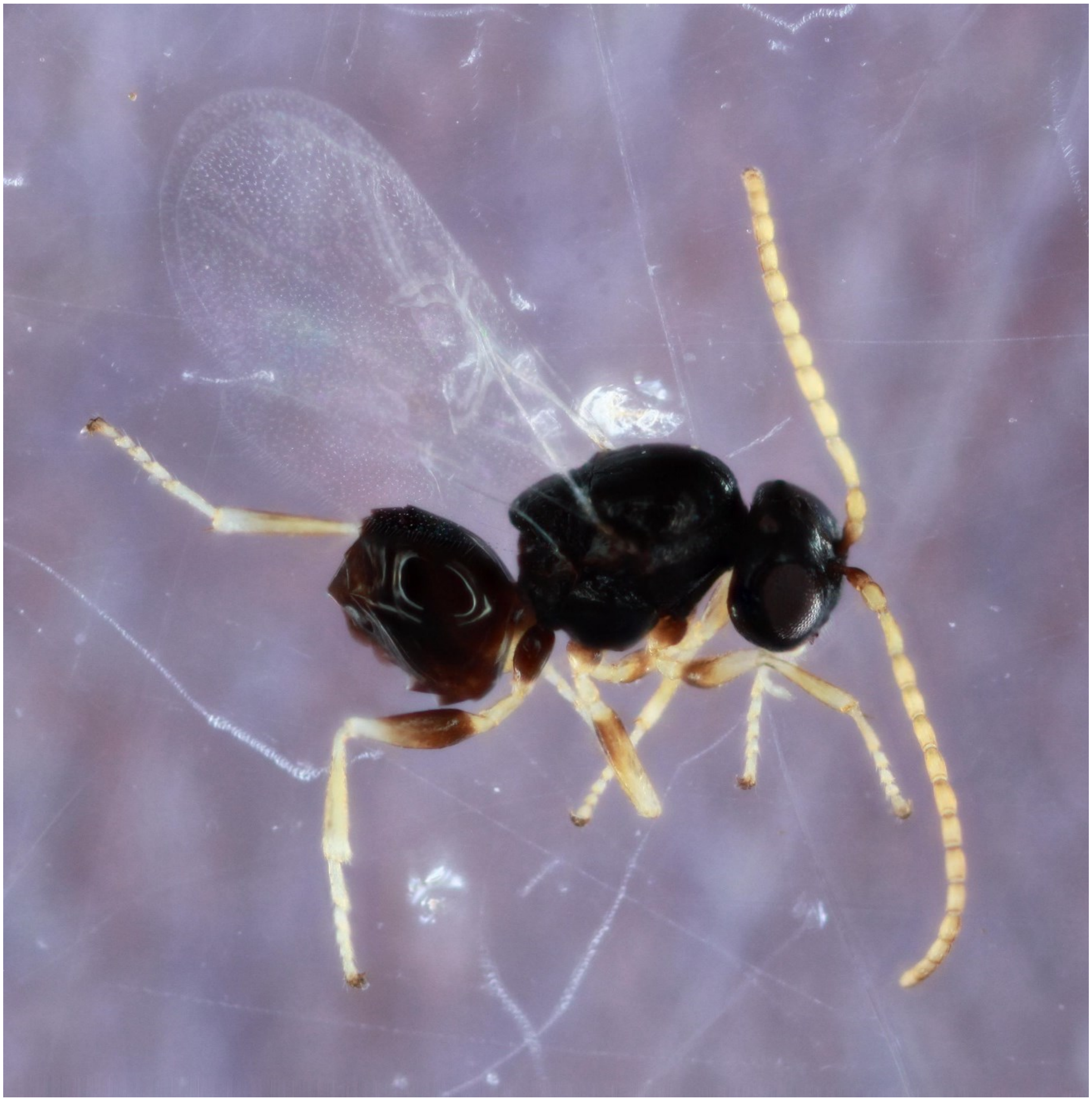

*Callirhytis glandulus* - Wasp: 1013-098-1

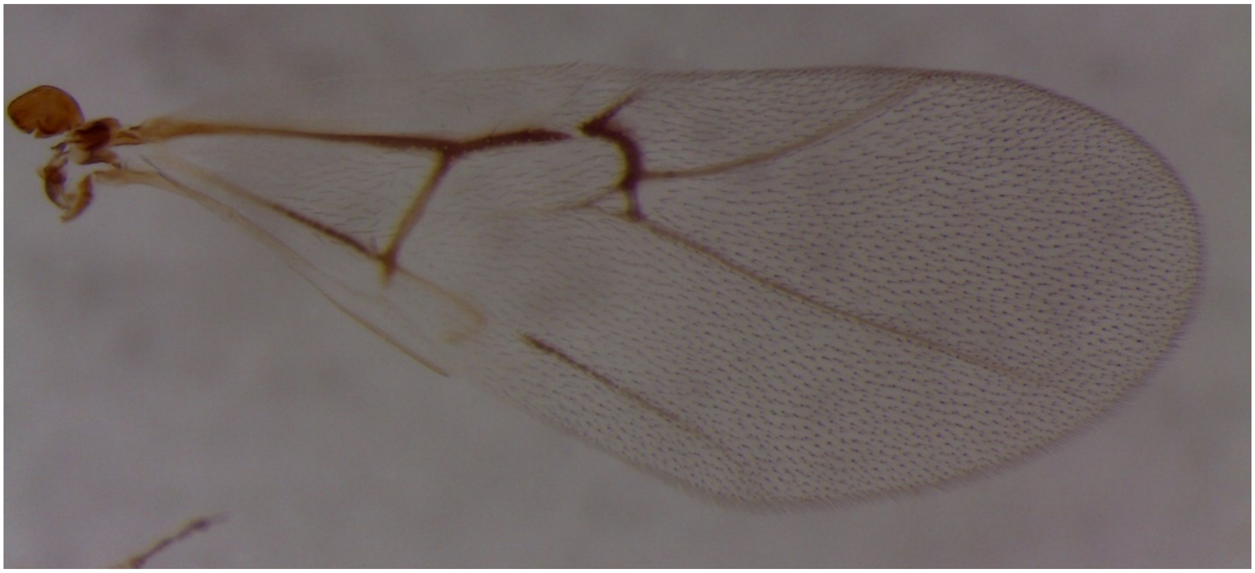

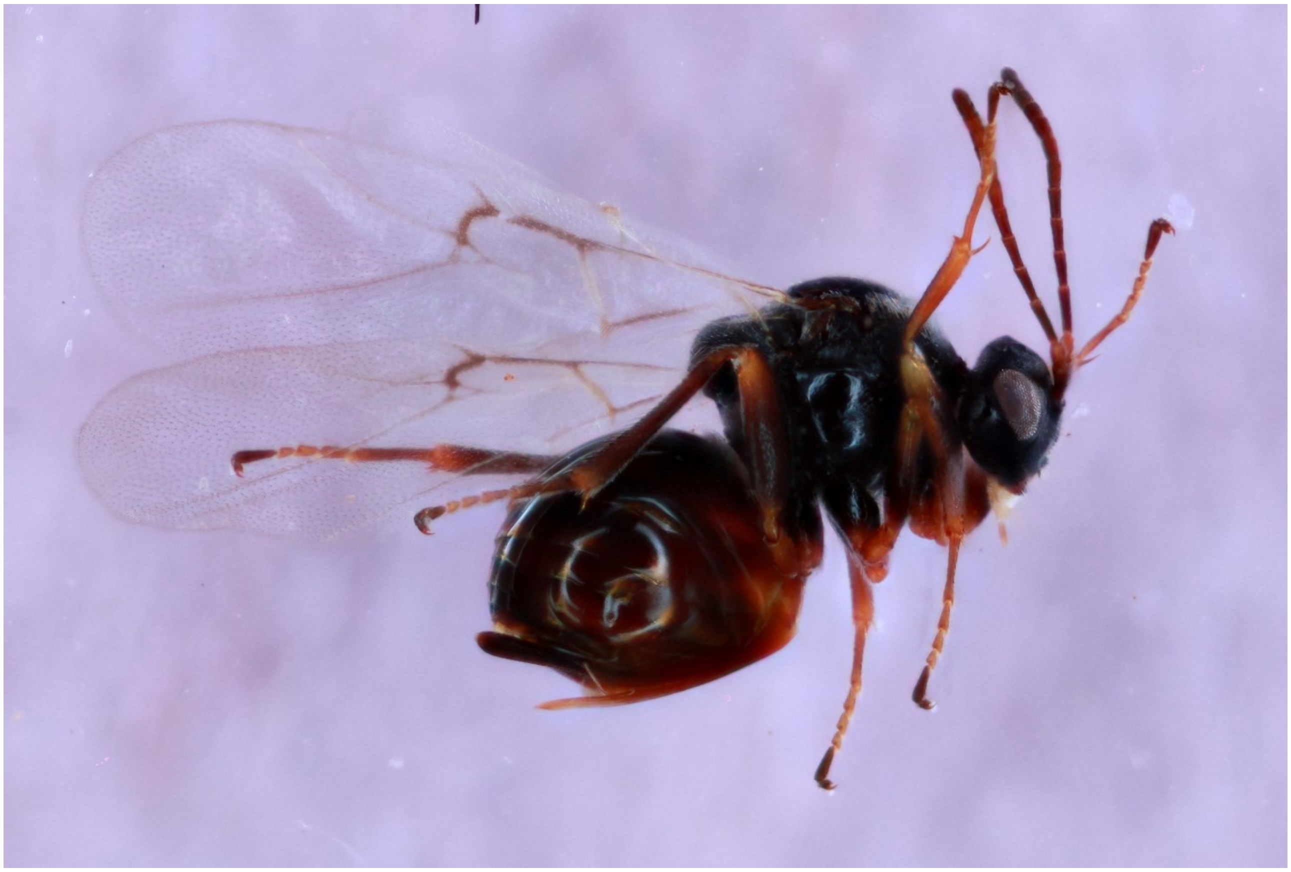

*Callirhytis pigra*- Wasp: 1574-104-3A (MALE)

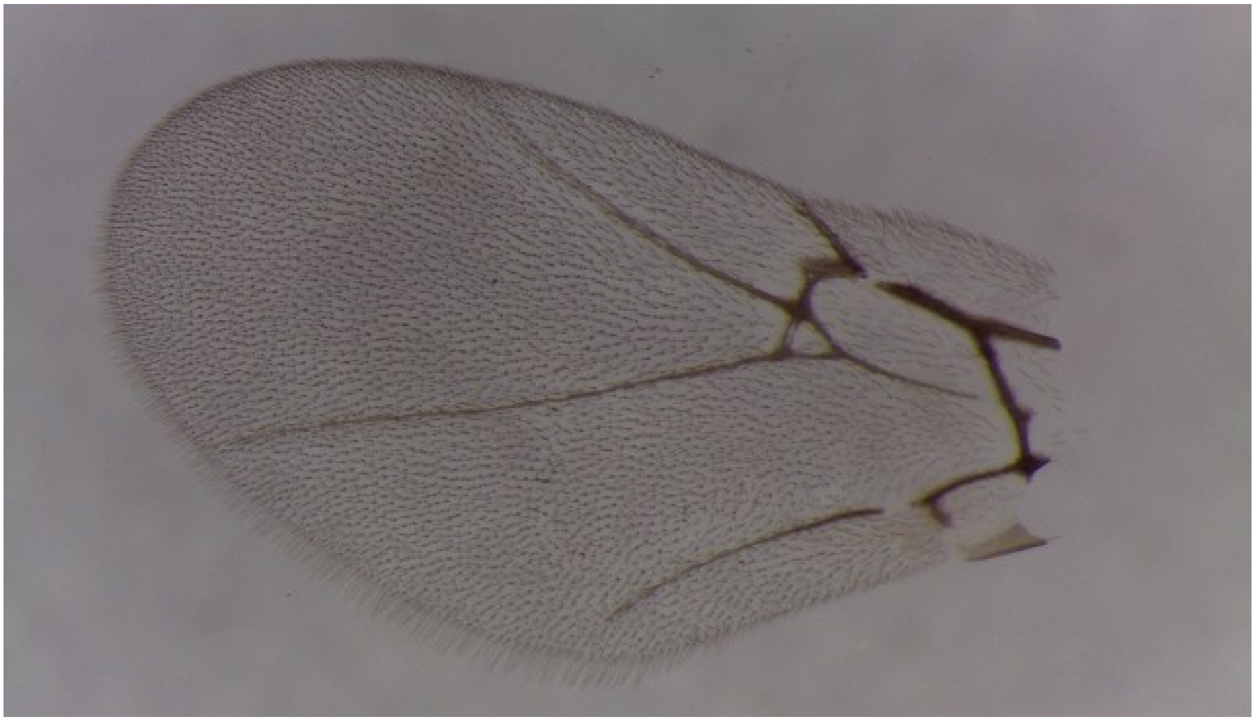

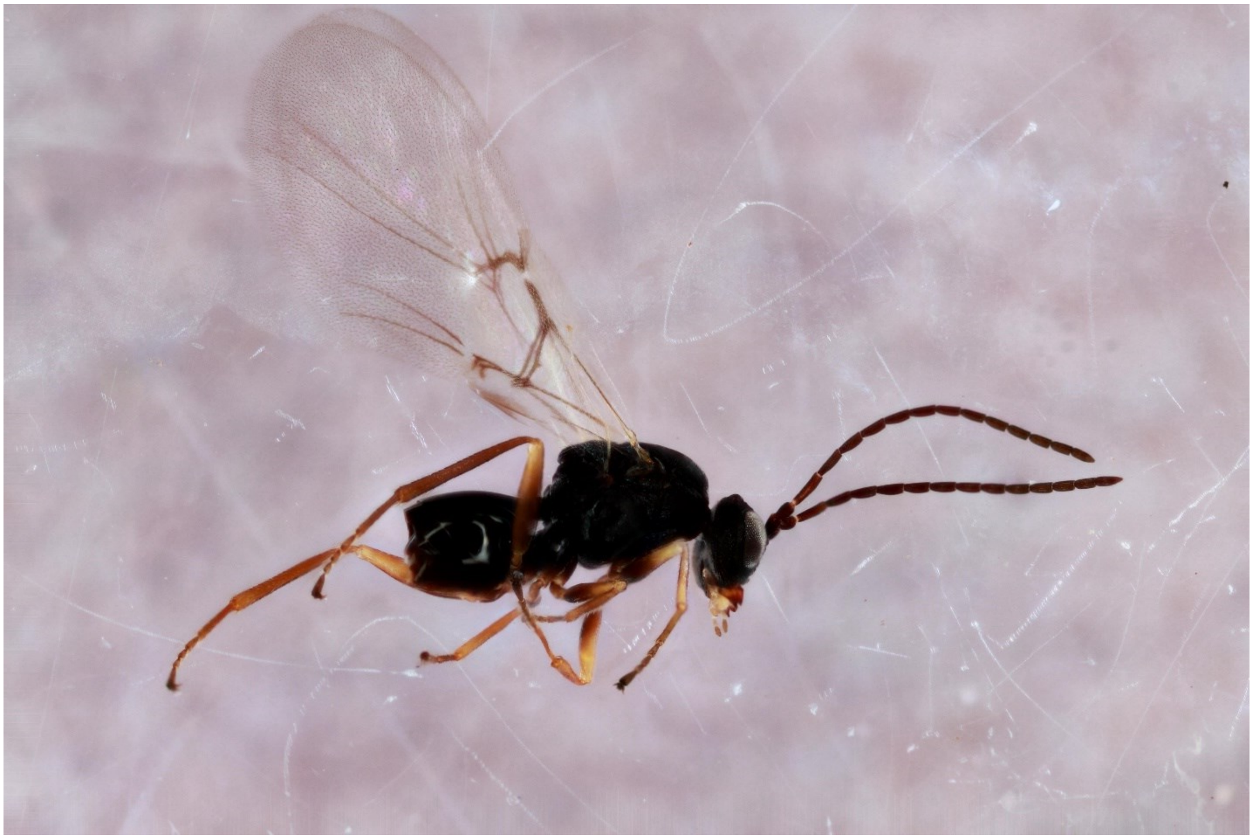

*Callirhytis pulchra* - Wasp: 1604-165-4A

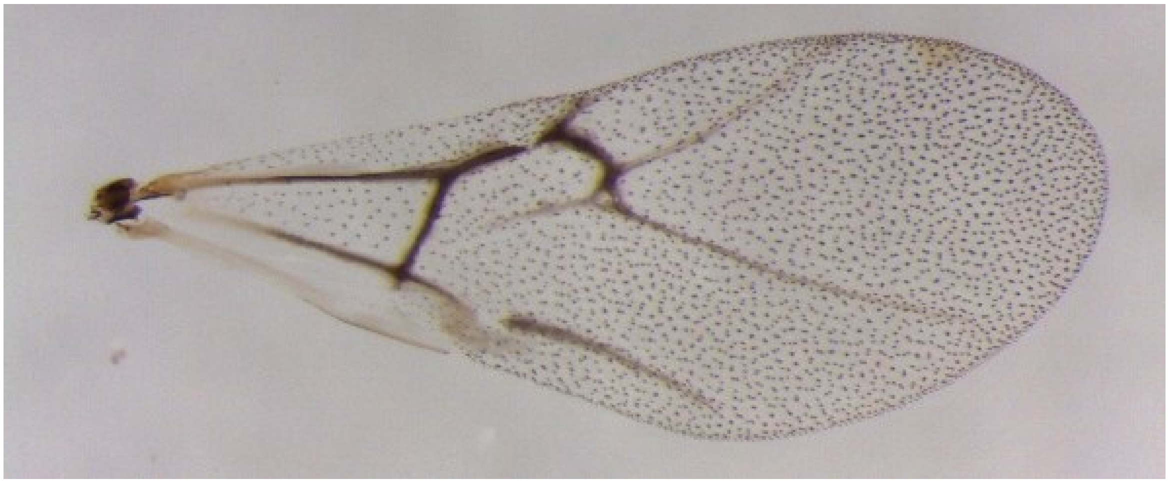

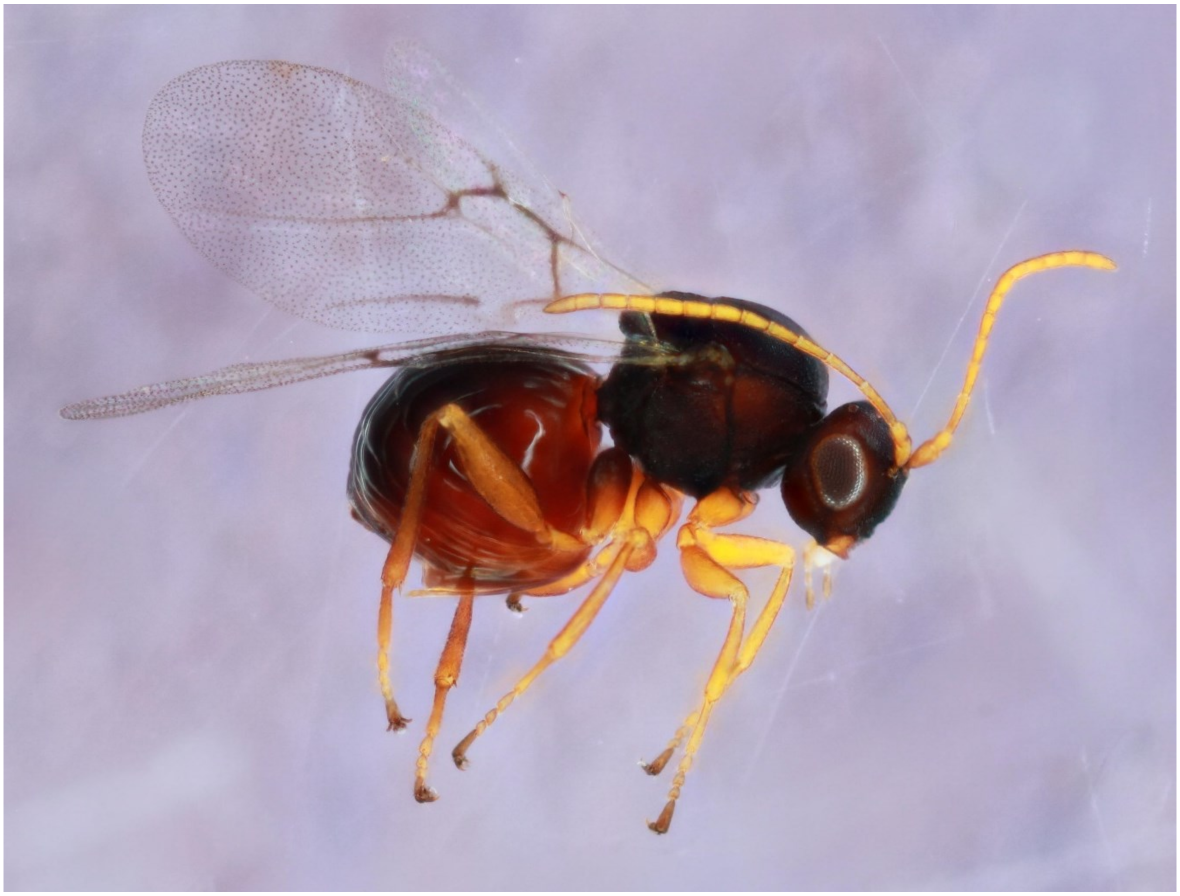

*Callirhytis quercuscornigera* - Wasp: 789-070-27A

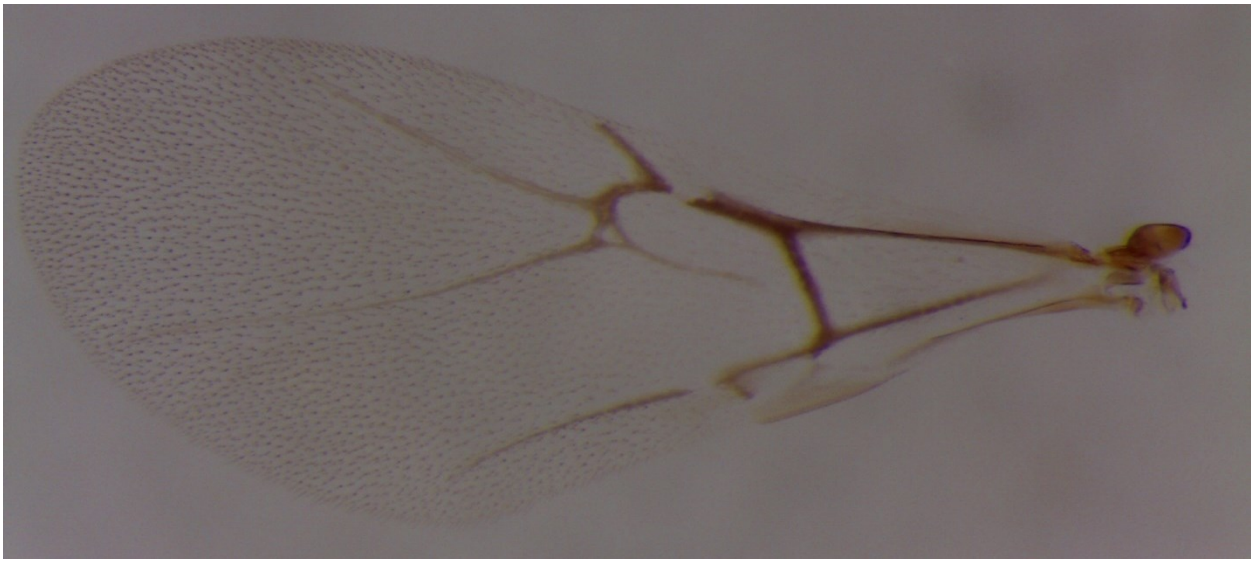

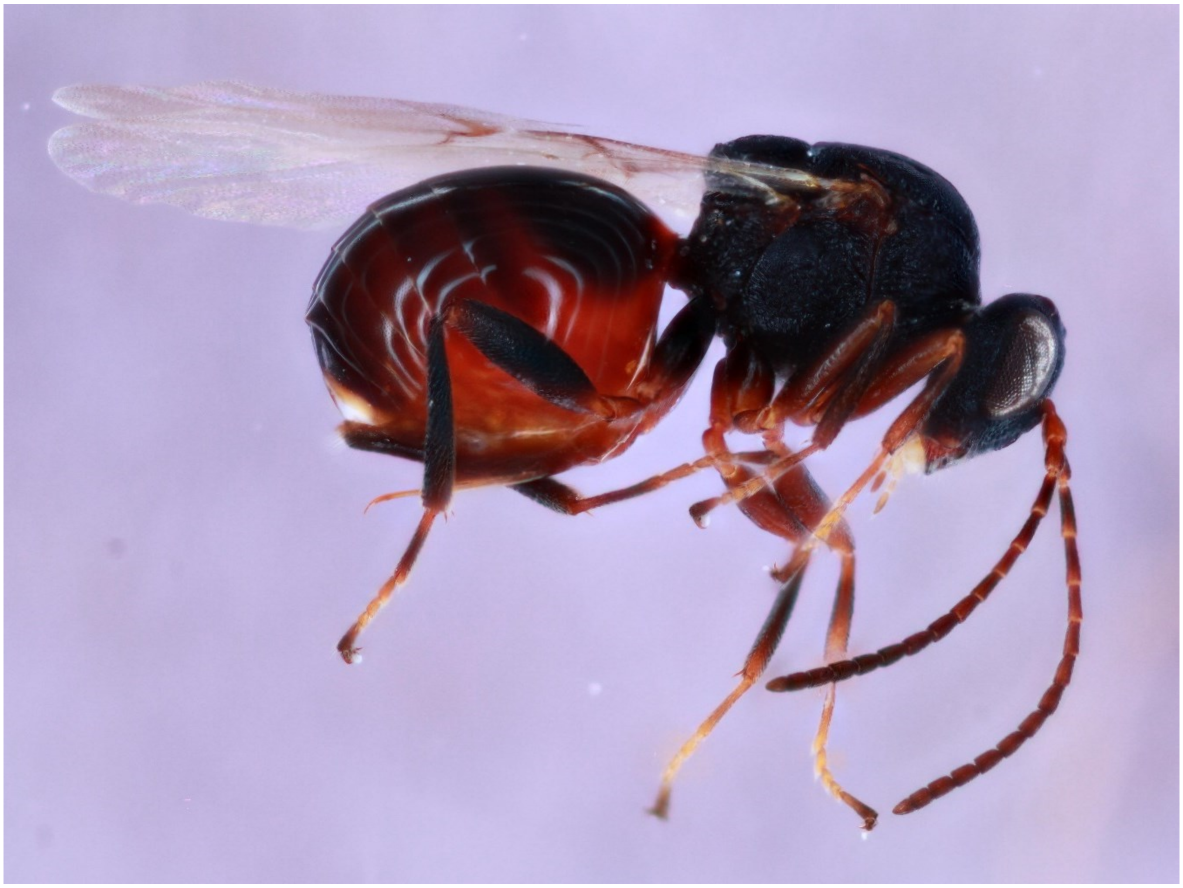

*Callirhytis quercusfutilis* - Wasp: 865-051-001A

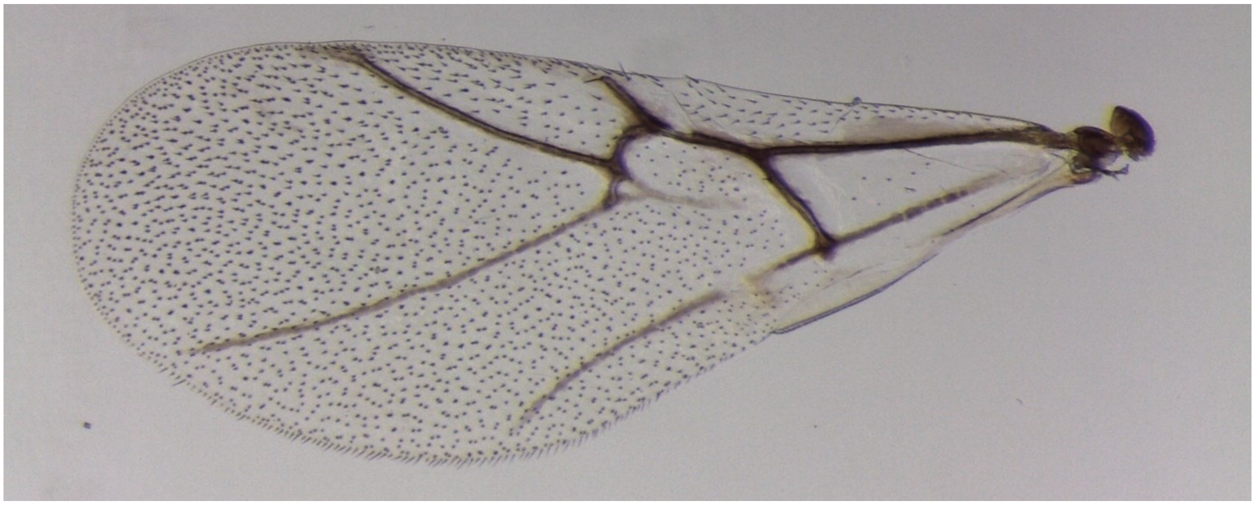

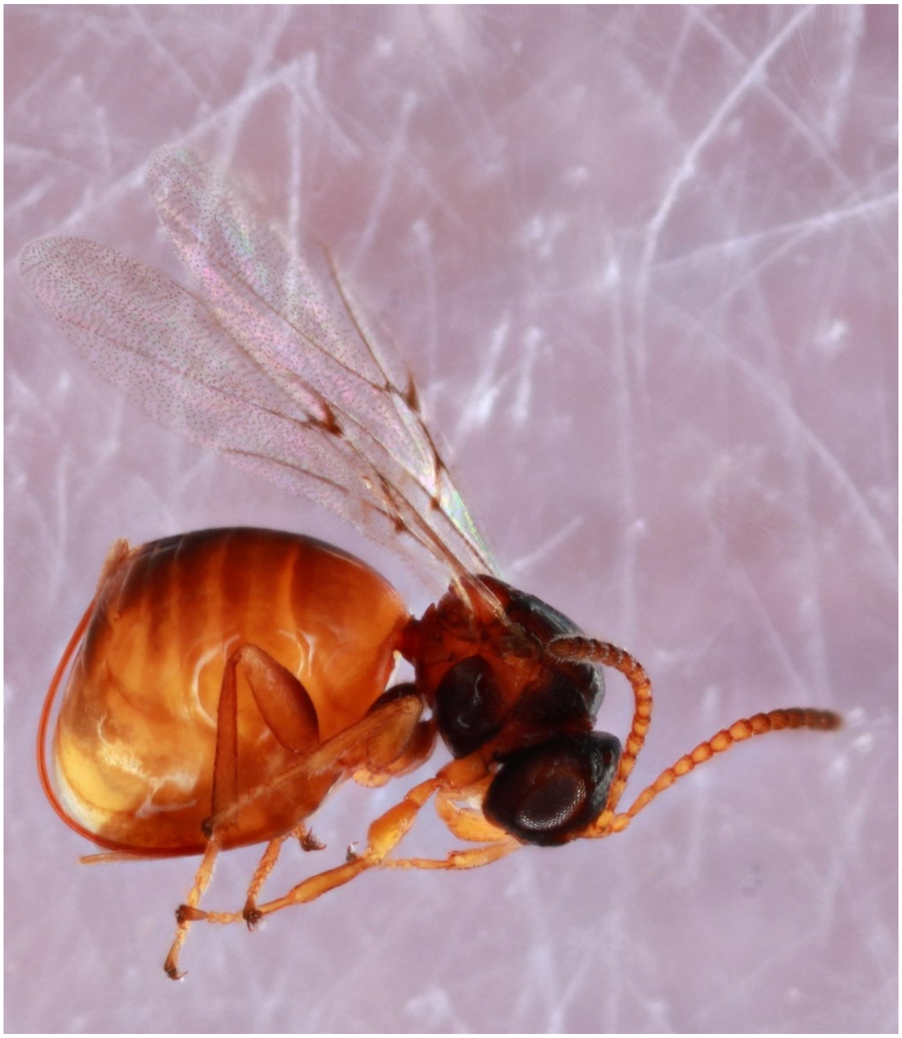

*Callirhytis quercusoperator* - Wasp: 1344-123-17A

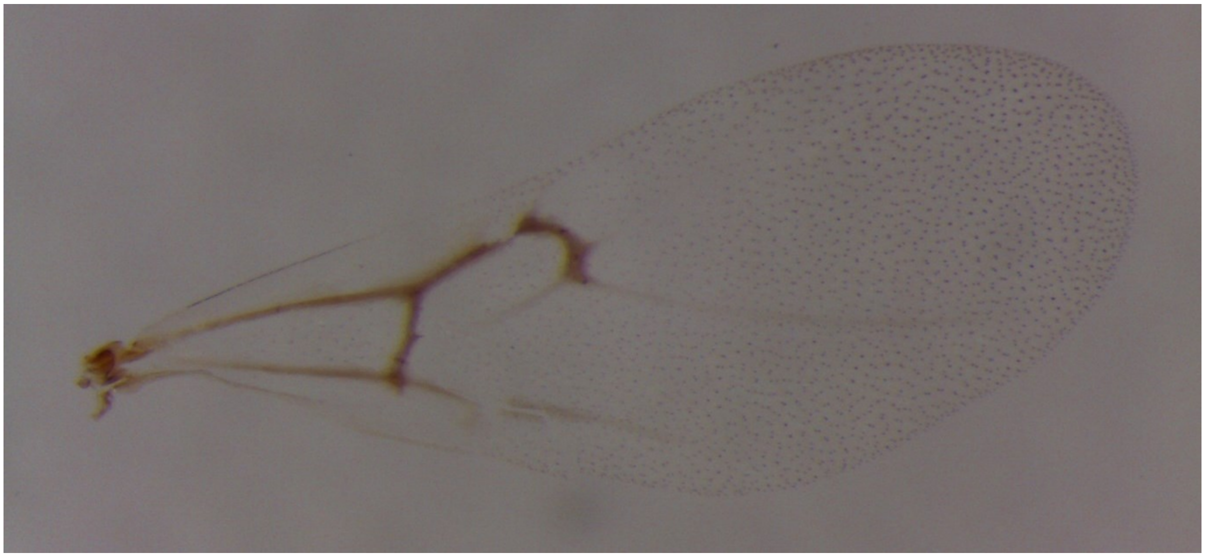

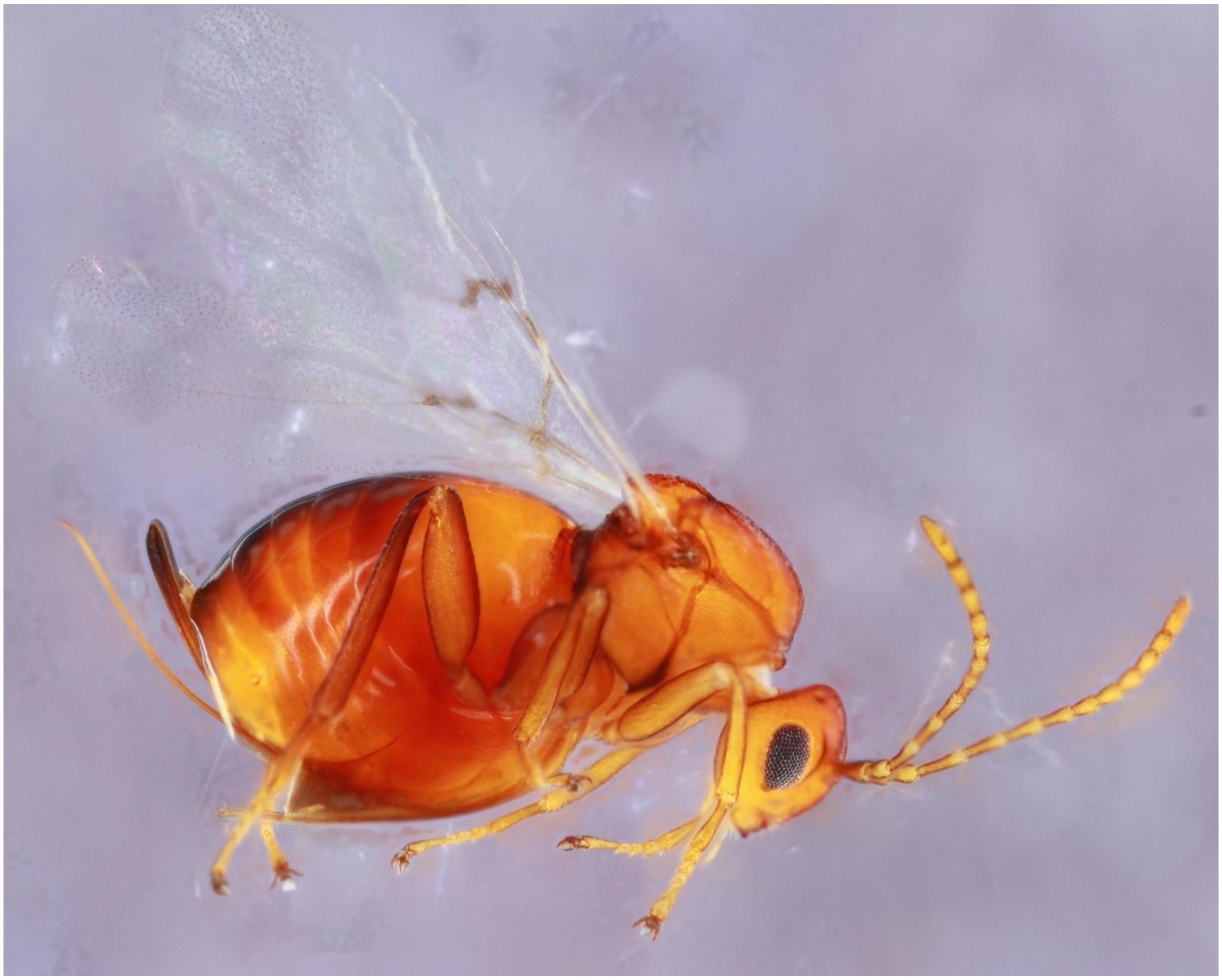

*Callirhytis quercuspunctata* - Wasp: 847-069-4A

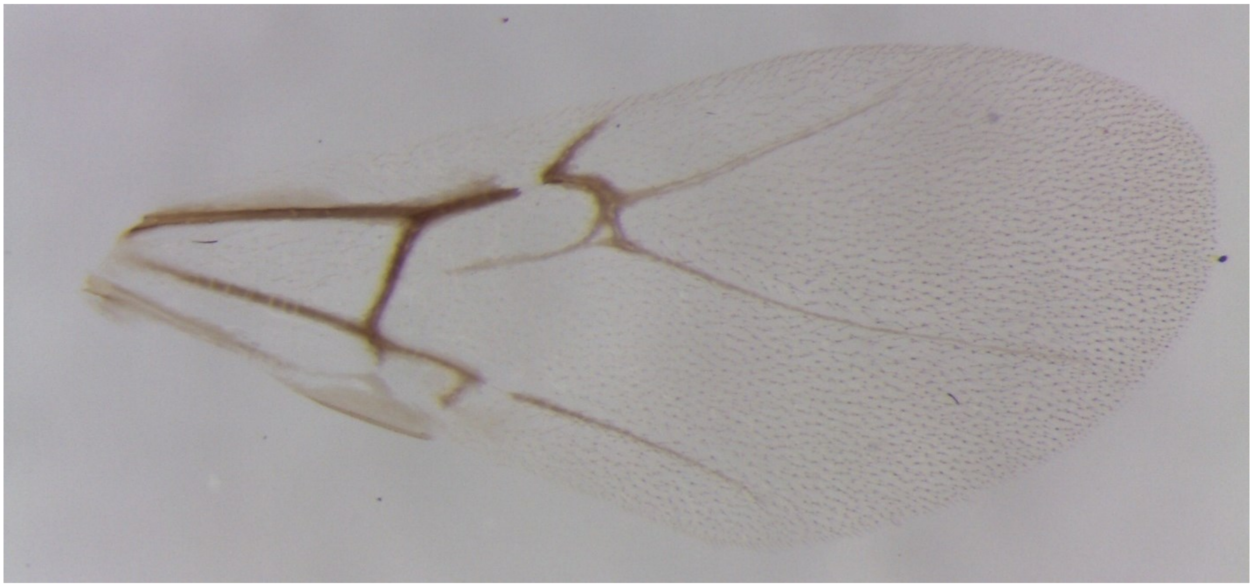

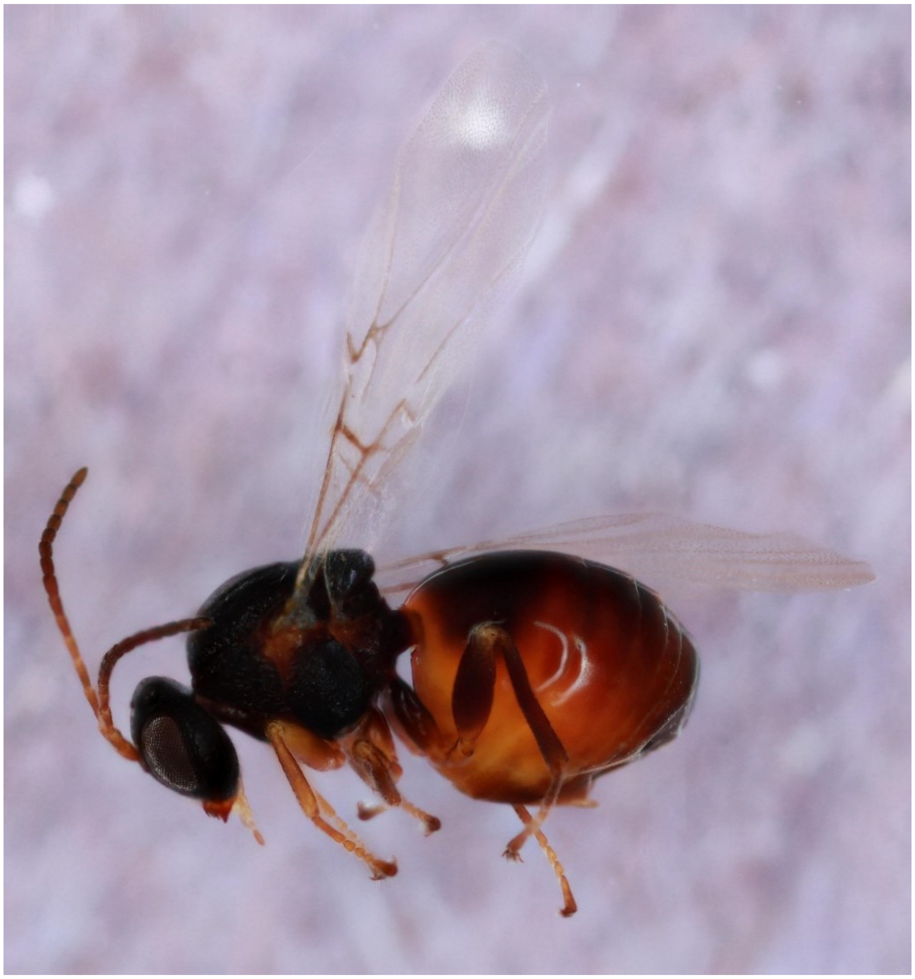

*Callirhytis quercusventricosa* - Wasp: 1404-041-4

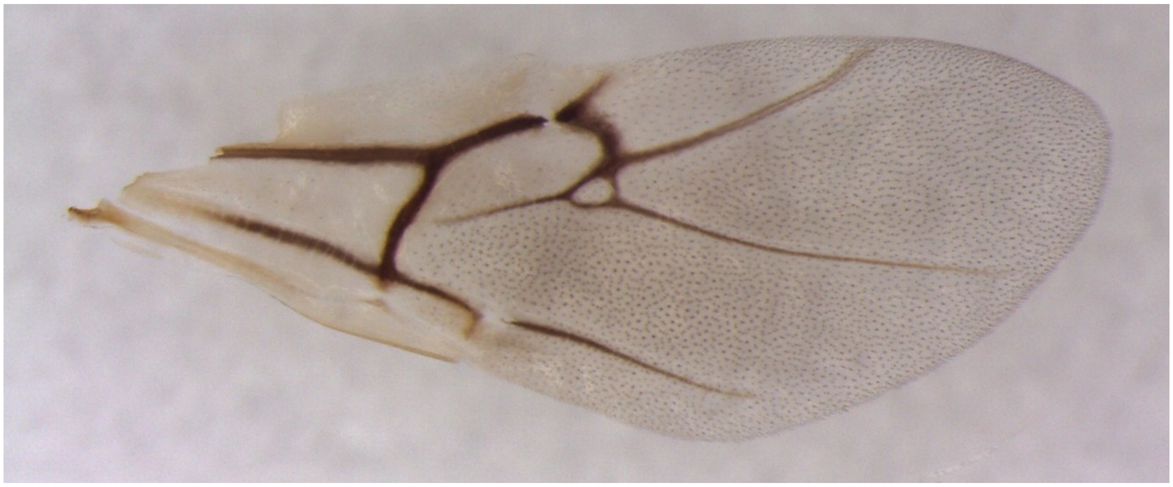

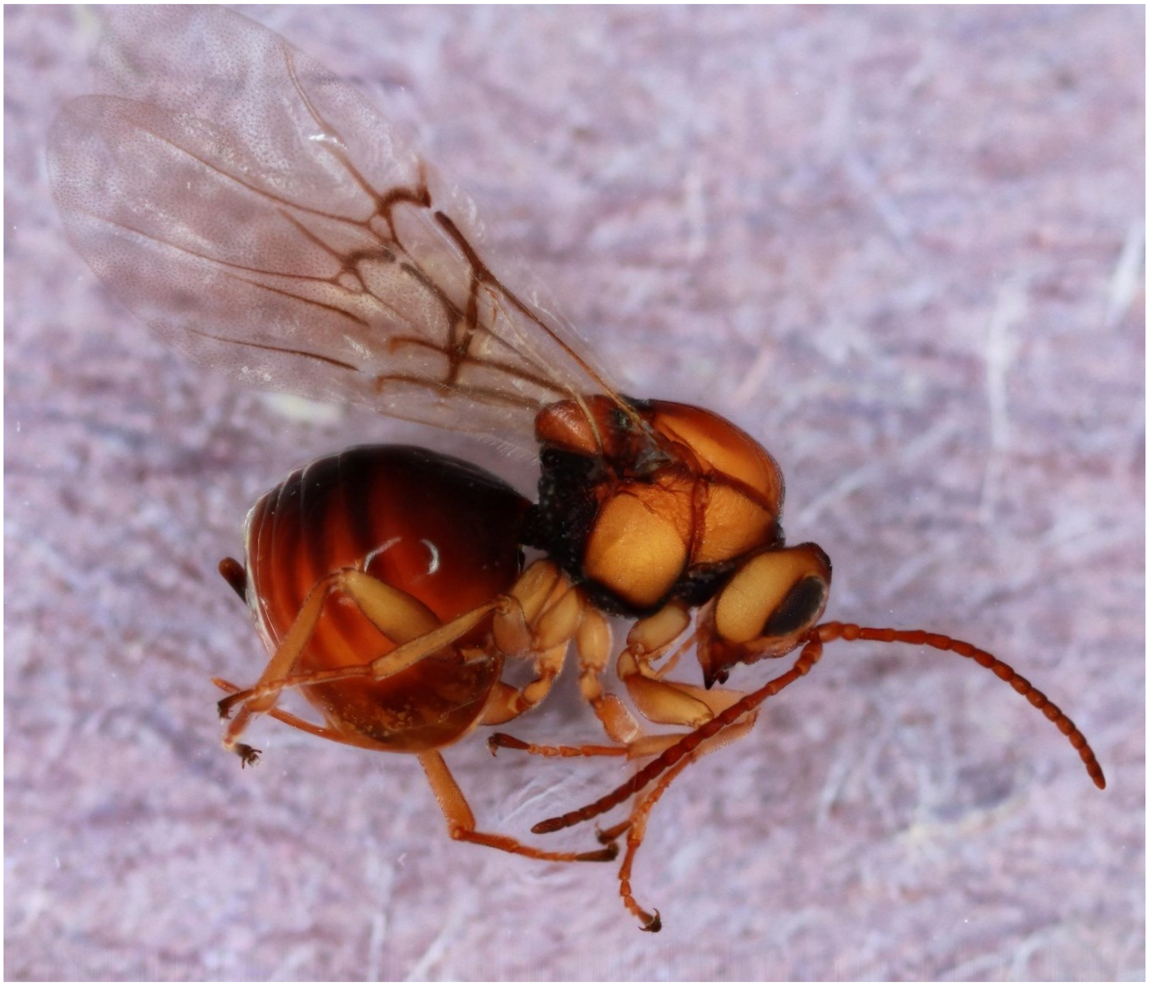

*Callirhytis scitula* - Wasp: 872-049-001A

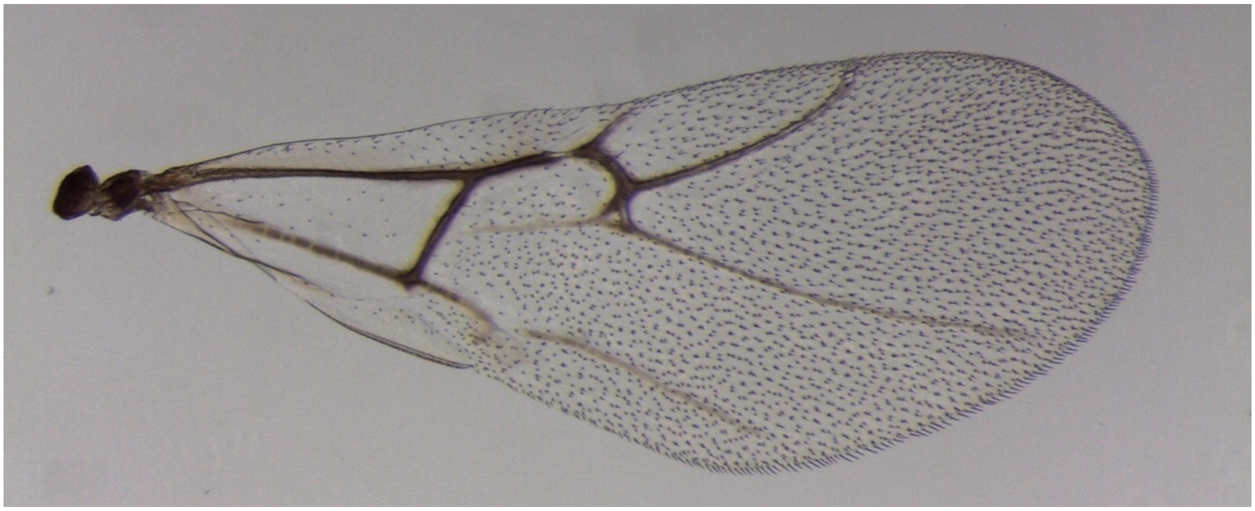

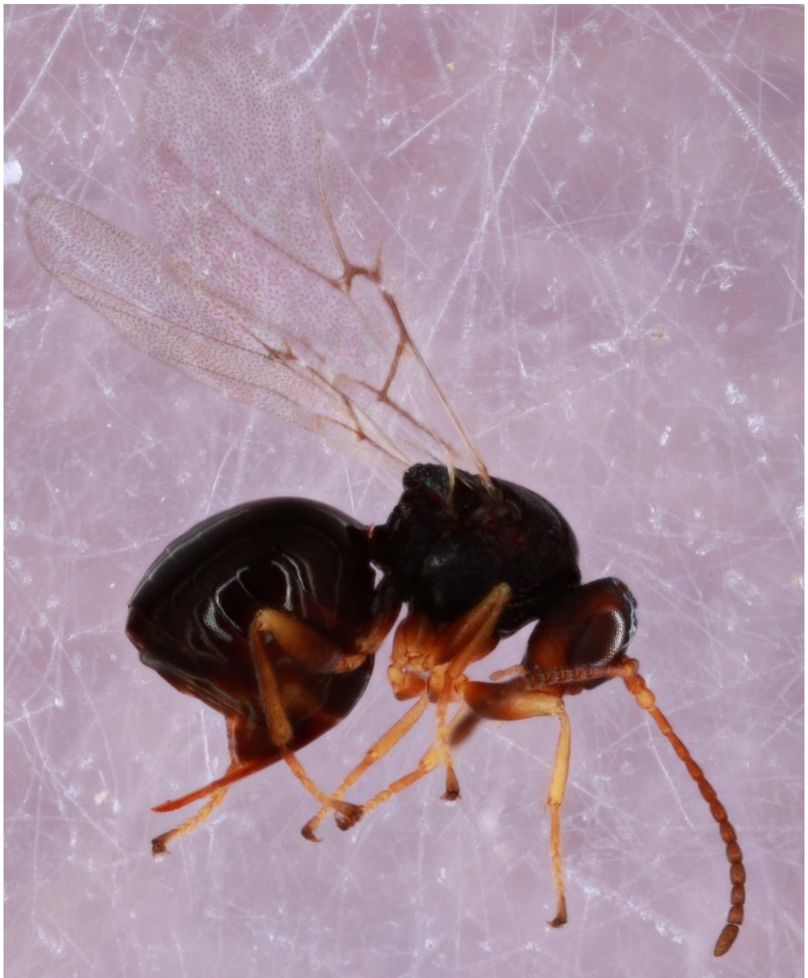

*Callirhytis seminator* - Wasp: 884-039-4

(No wing picture)

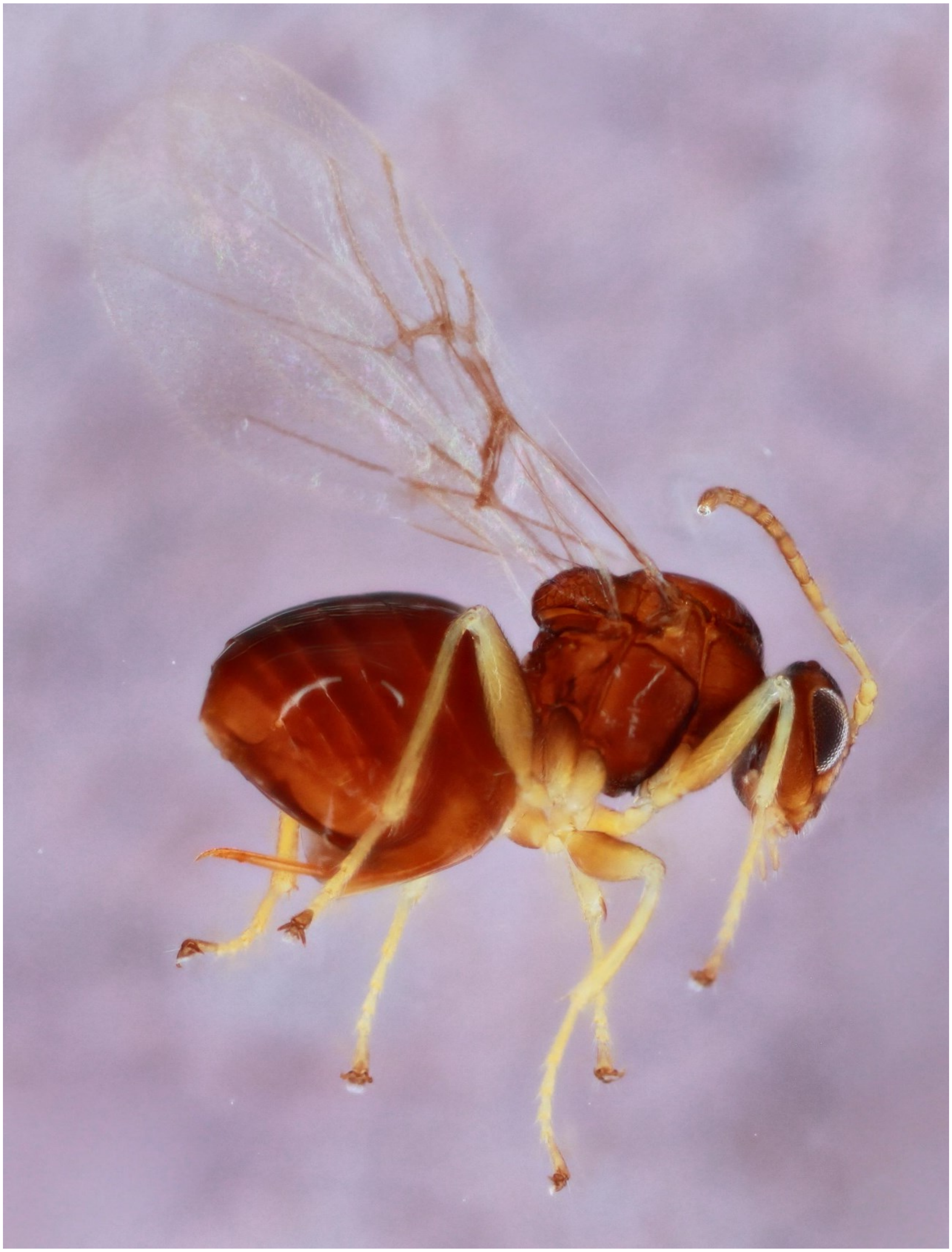

*Callirhytis* unknown leaf gall on *Quercus stellata* - Wasp: 1096-113-1

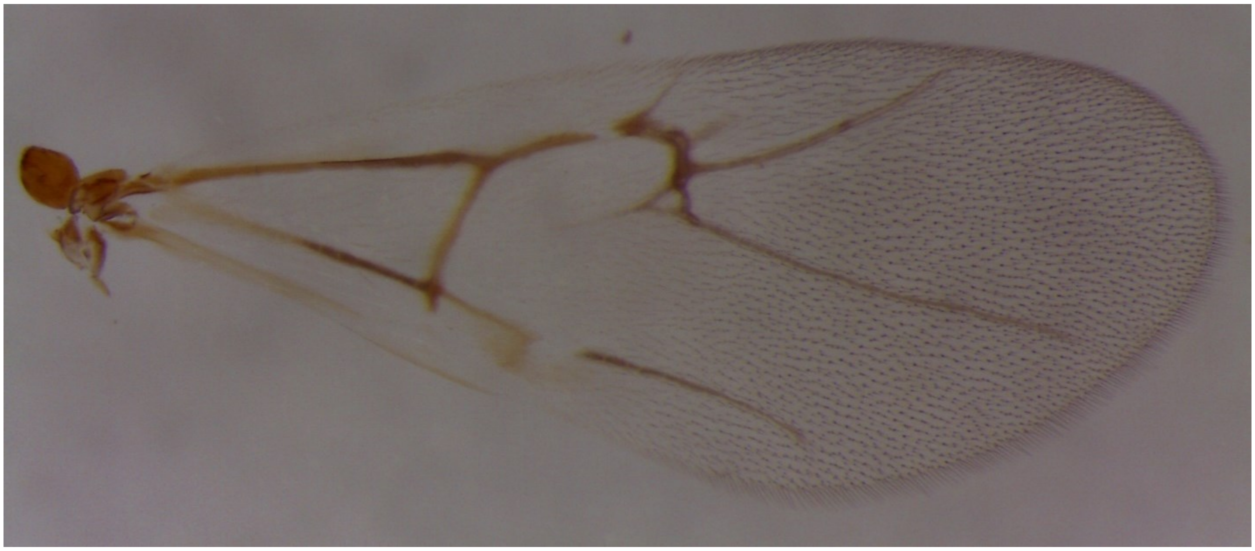

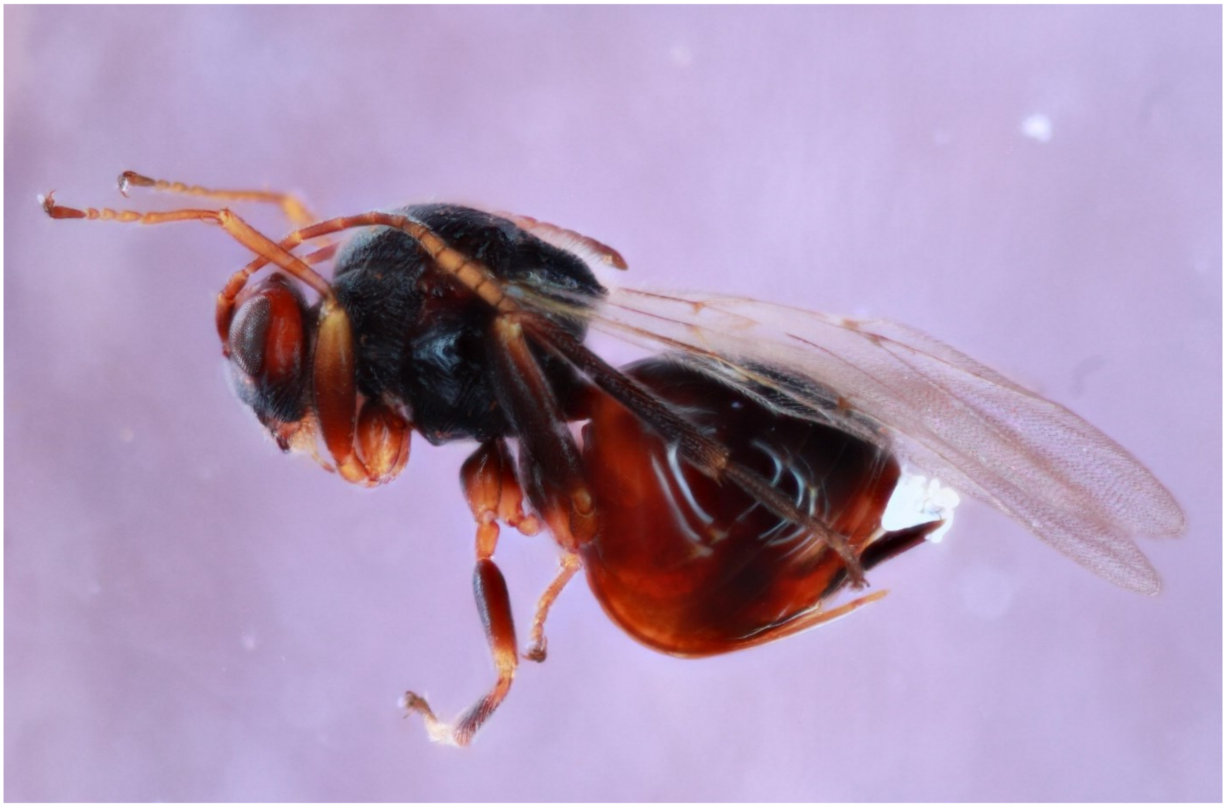

*Callirhytis vaccinii* - Wasp: 1580-155-9

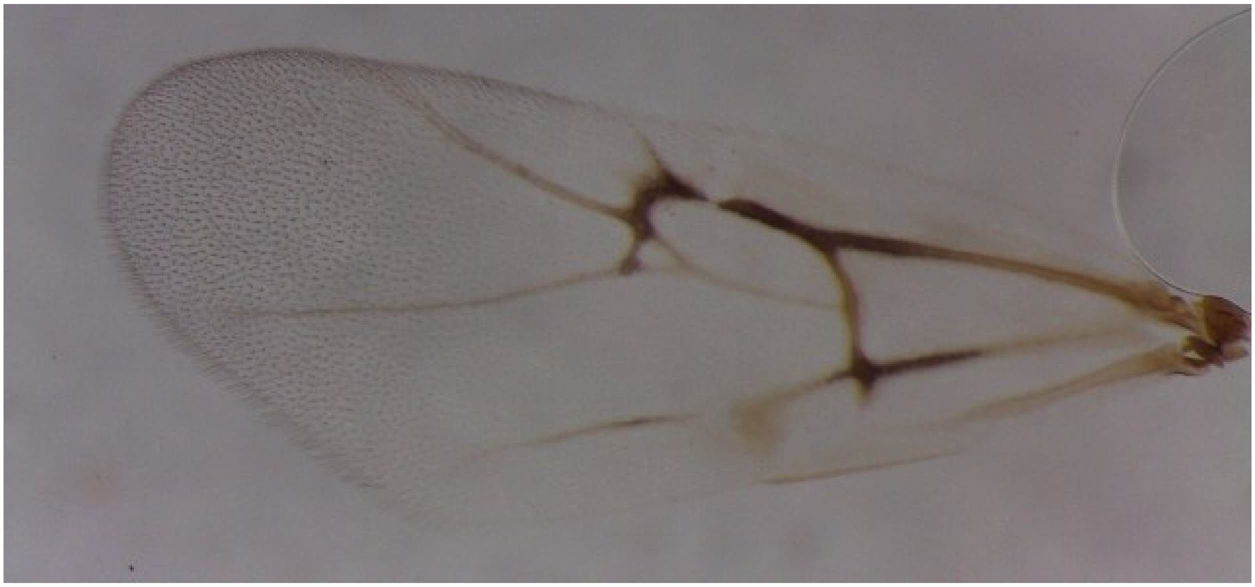

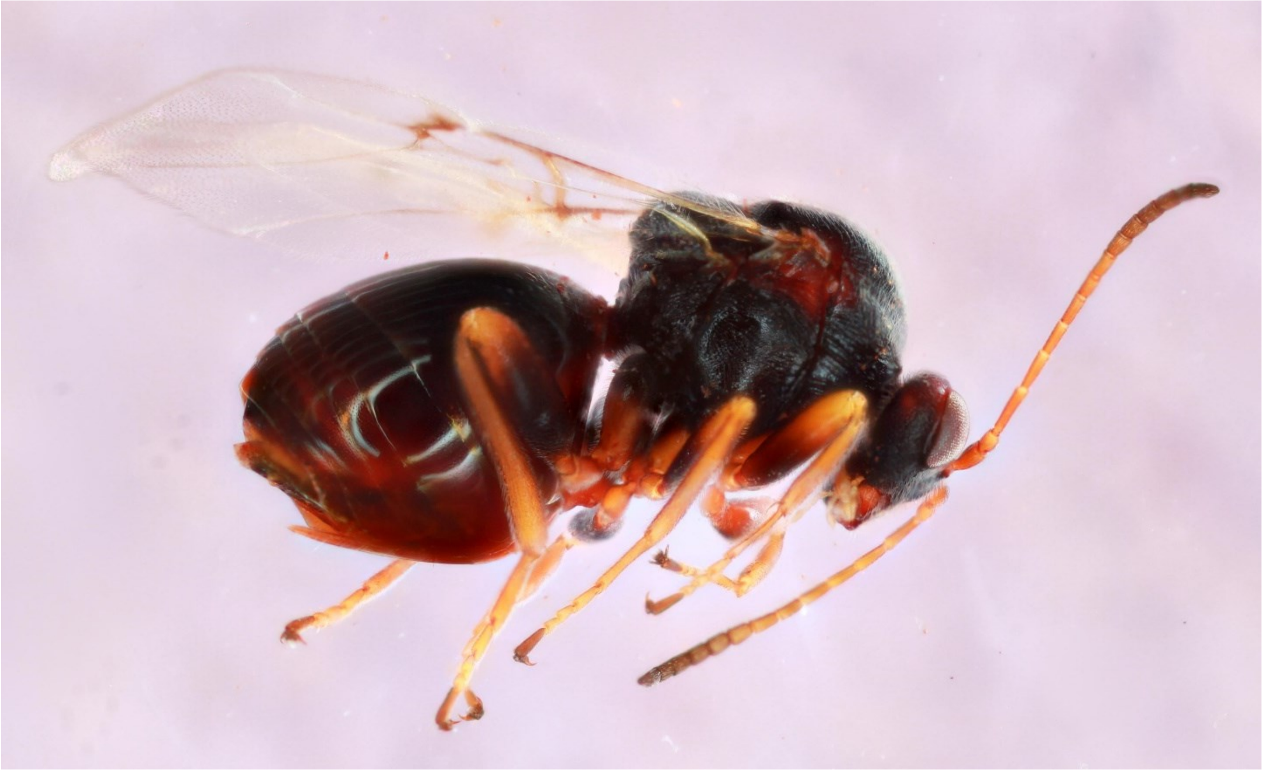

*Cynips conspicuus* - Wasp: 1489-125-2

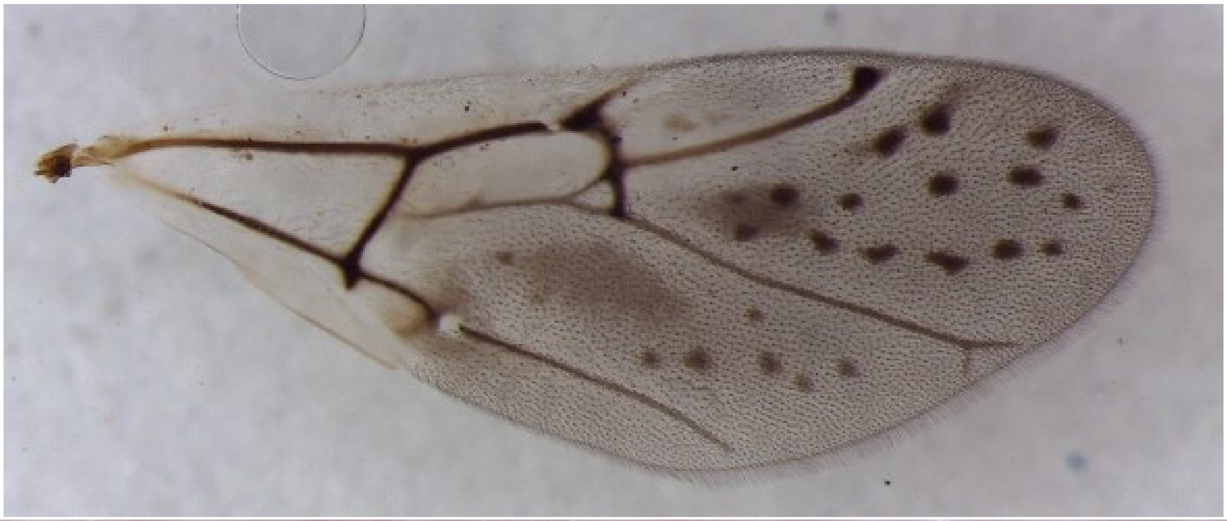

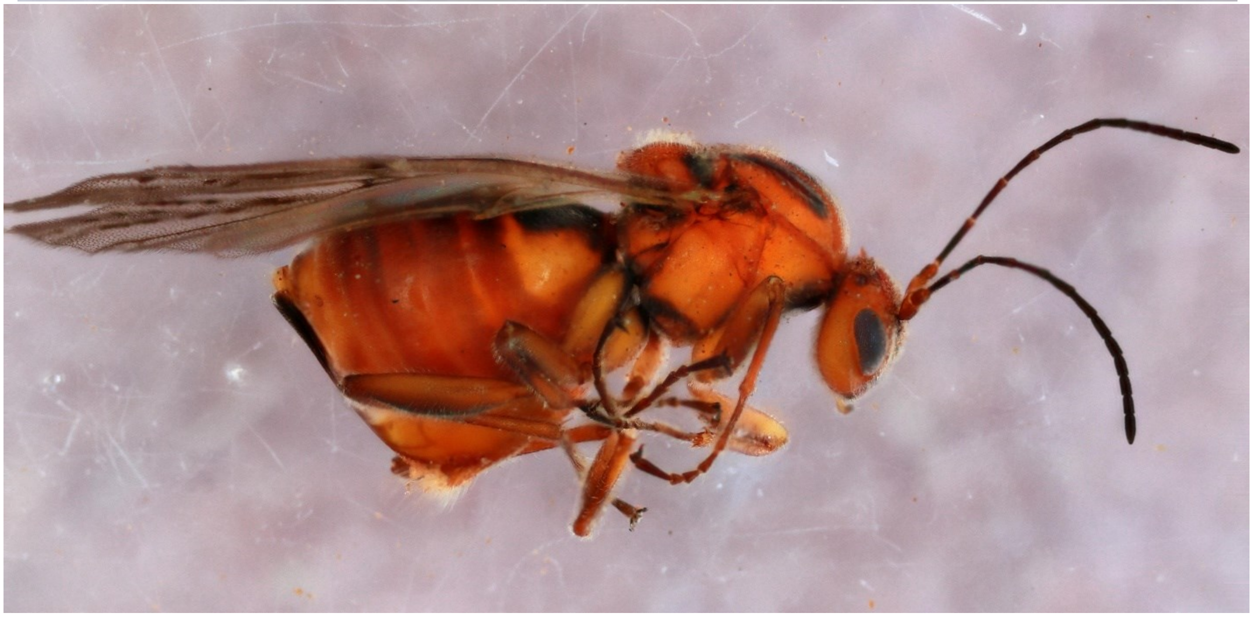

*Disholcaspis cinerosa* - Wasp: 114-1-35A

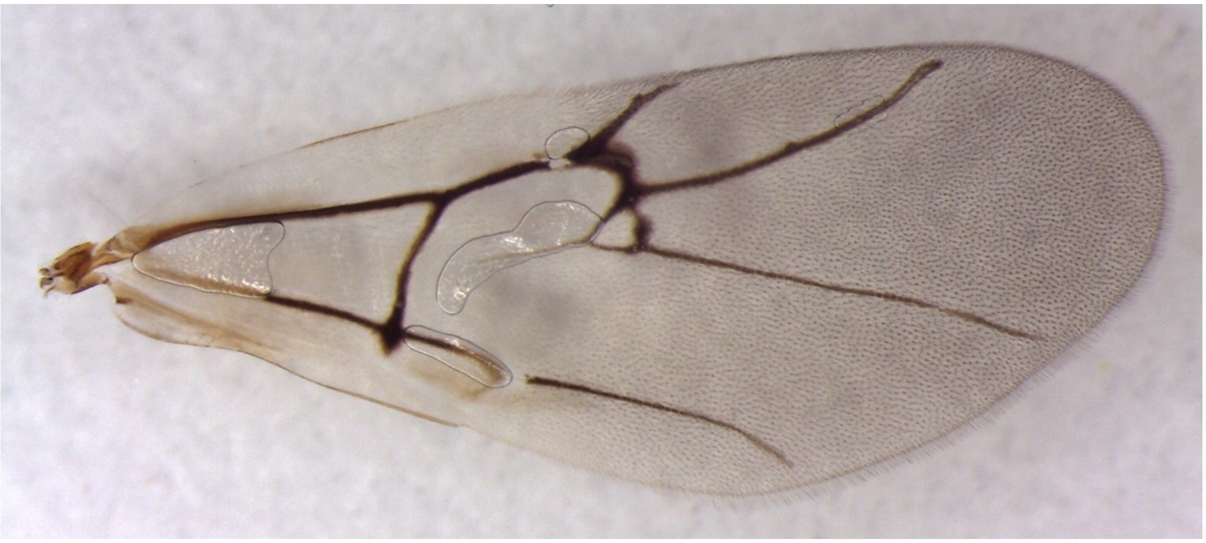

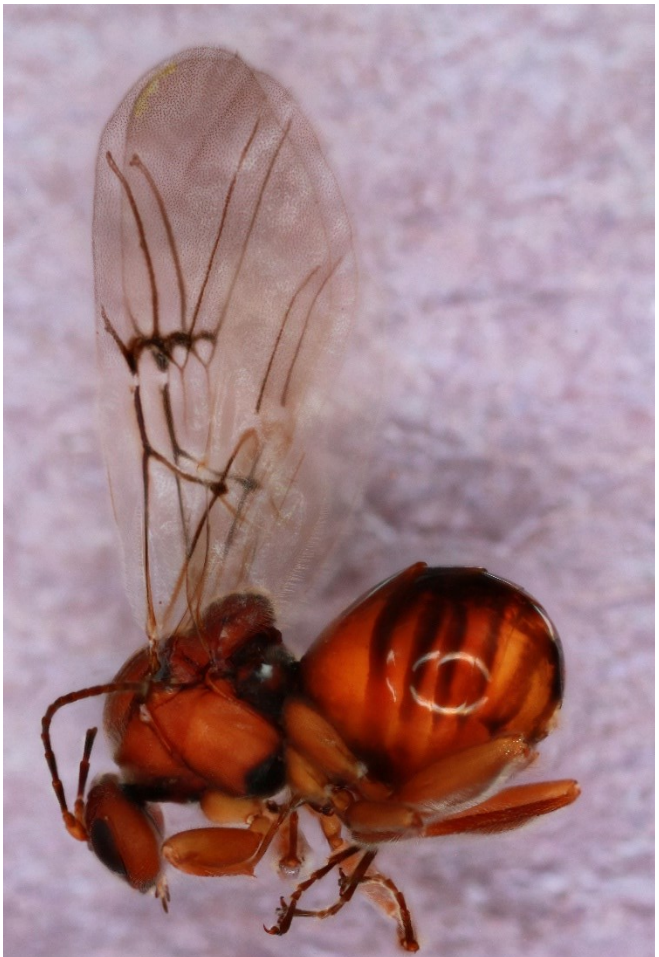

*Disholcaspis edura* - Wasp: 1685-216-1A

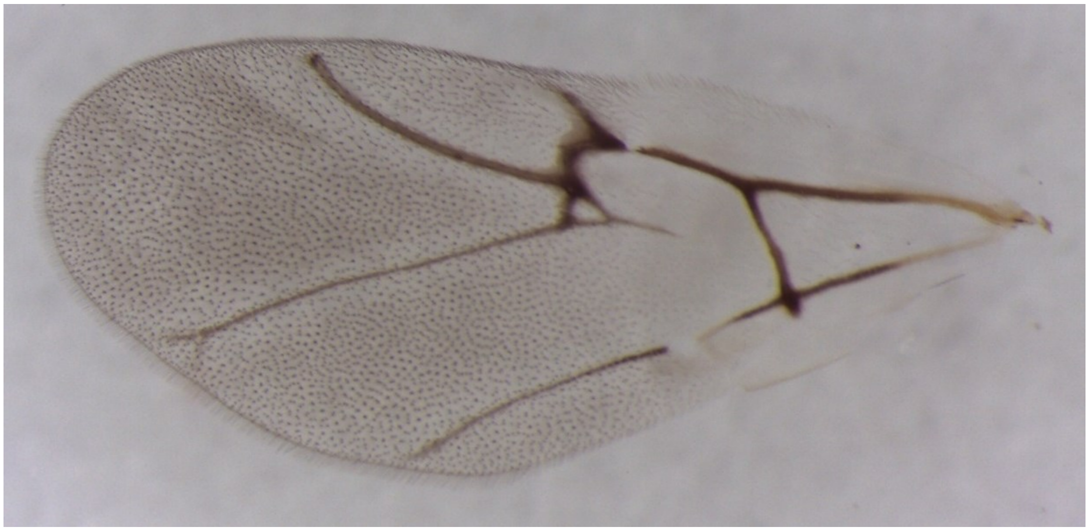

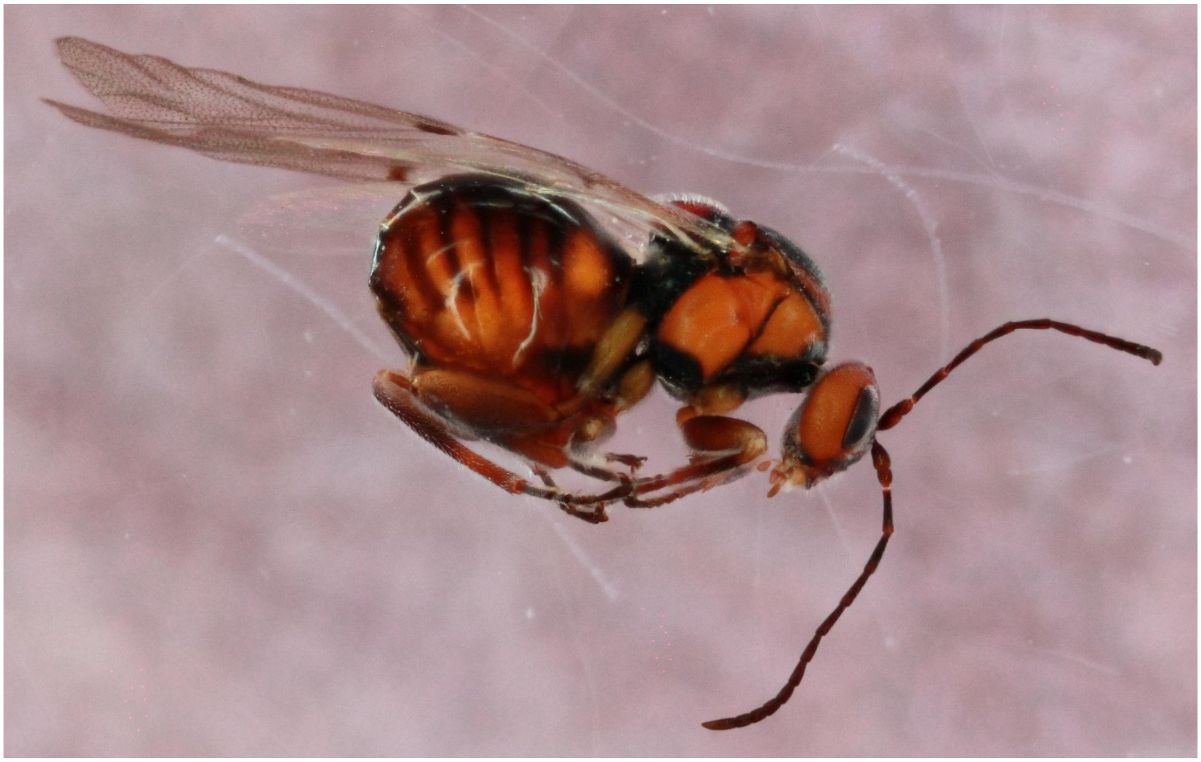

*Disholcaspis quercusglobulus* - Wasp: 1126-019-2

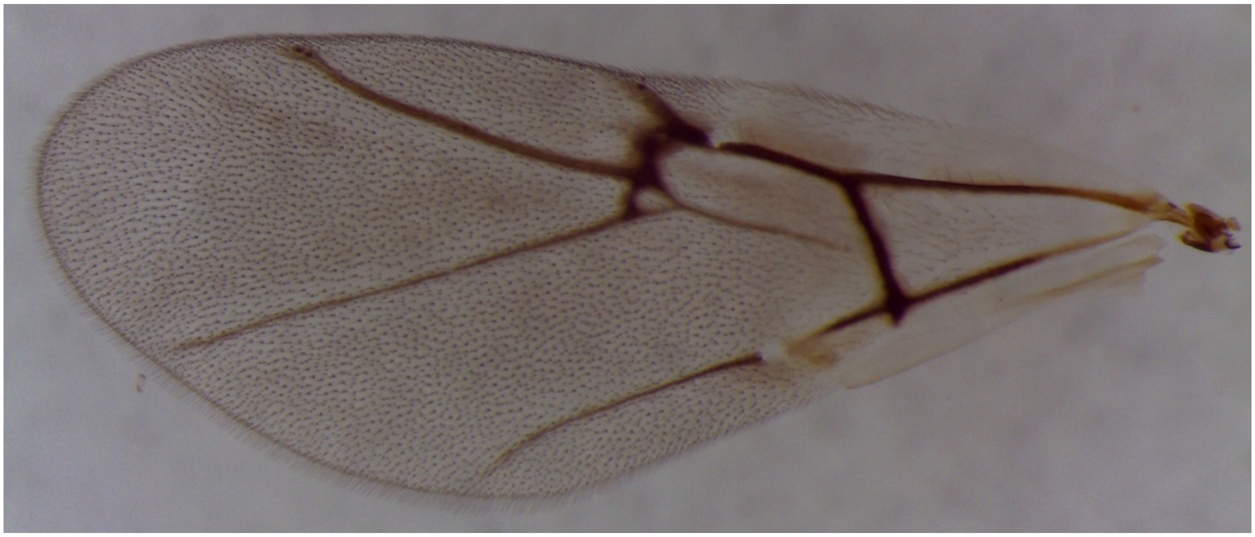

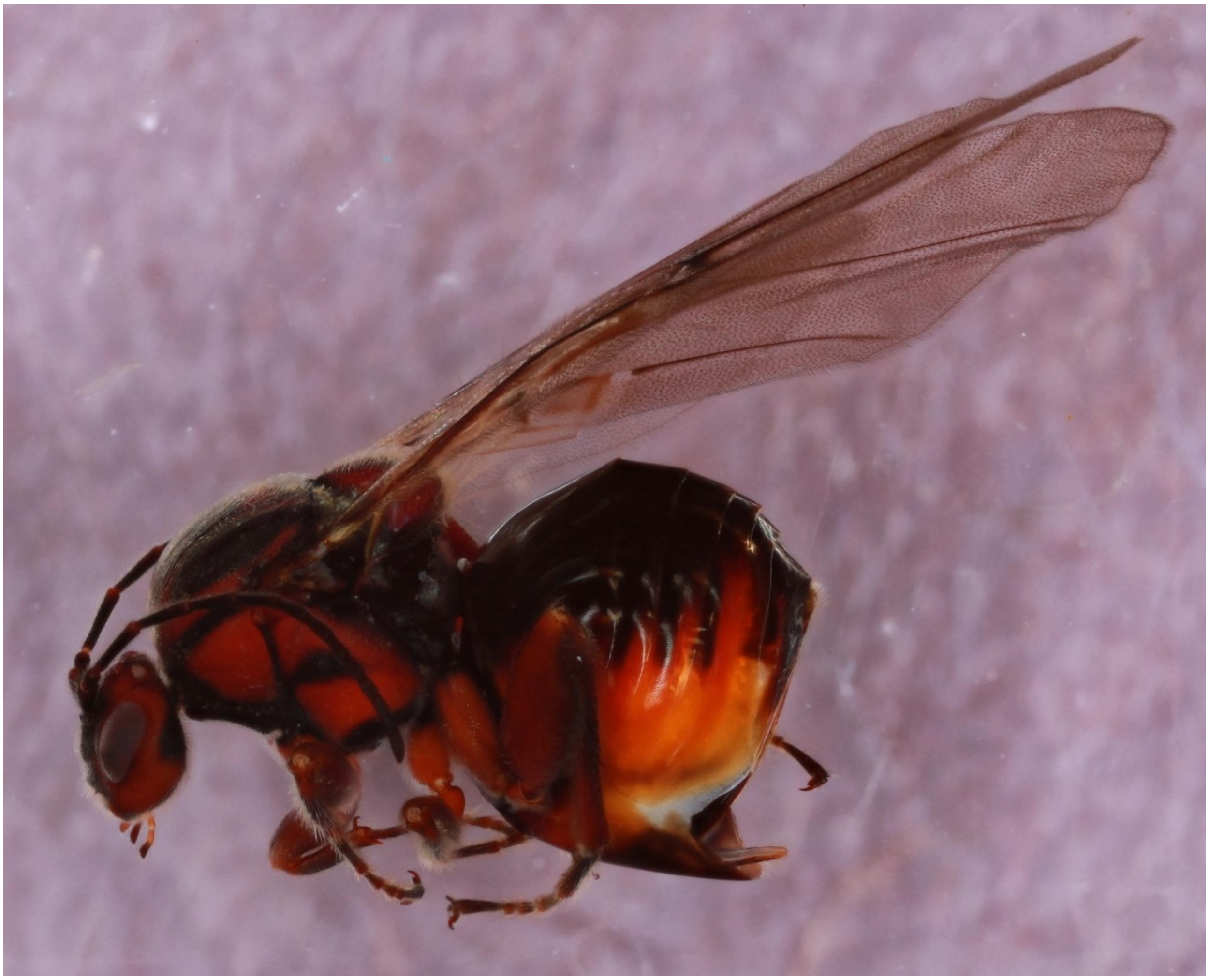

*Disholcaspis quercusmamma* (sexgen female) - Wasp: 1286-090-1A

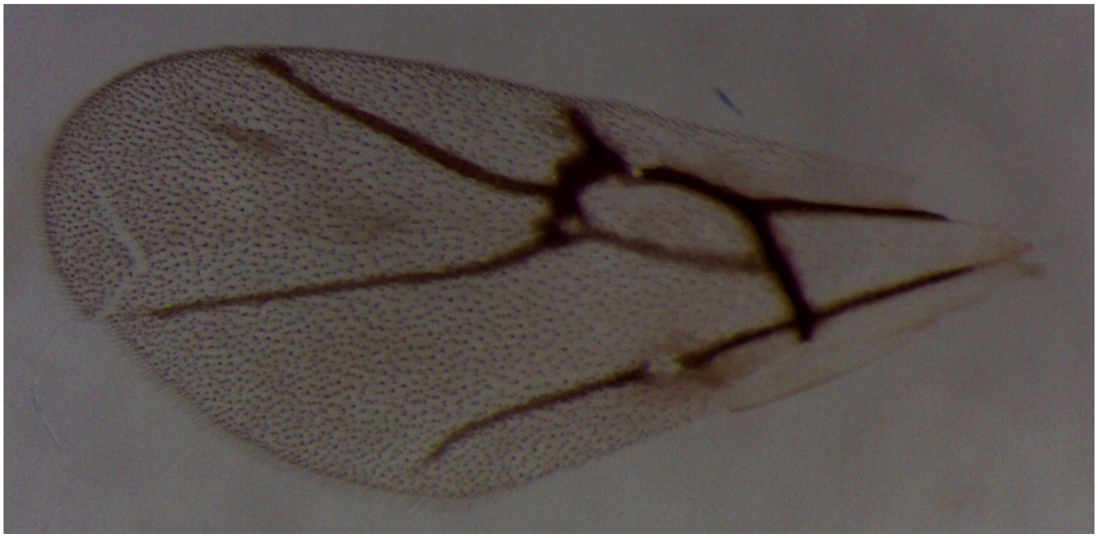

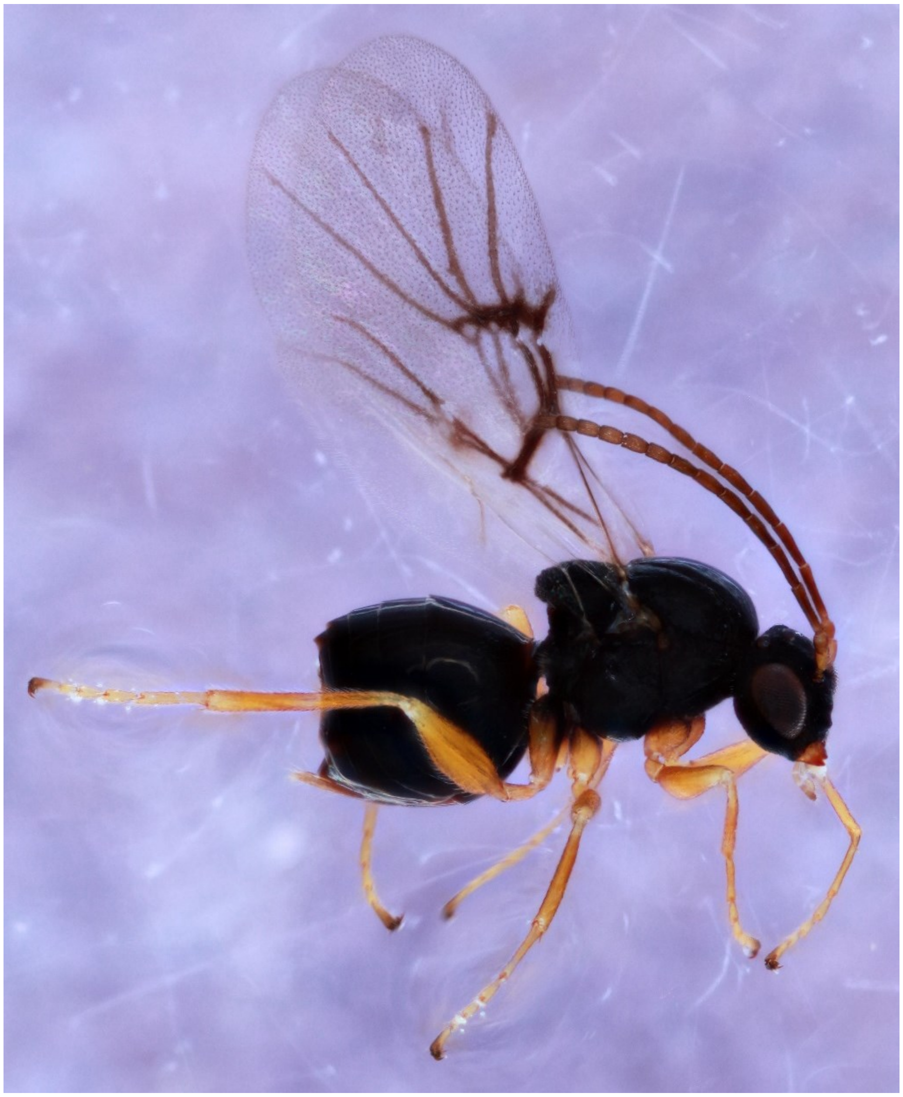

*Disholcaspis quercusmamma* (asex female) - Wasp: 644-020-001

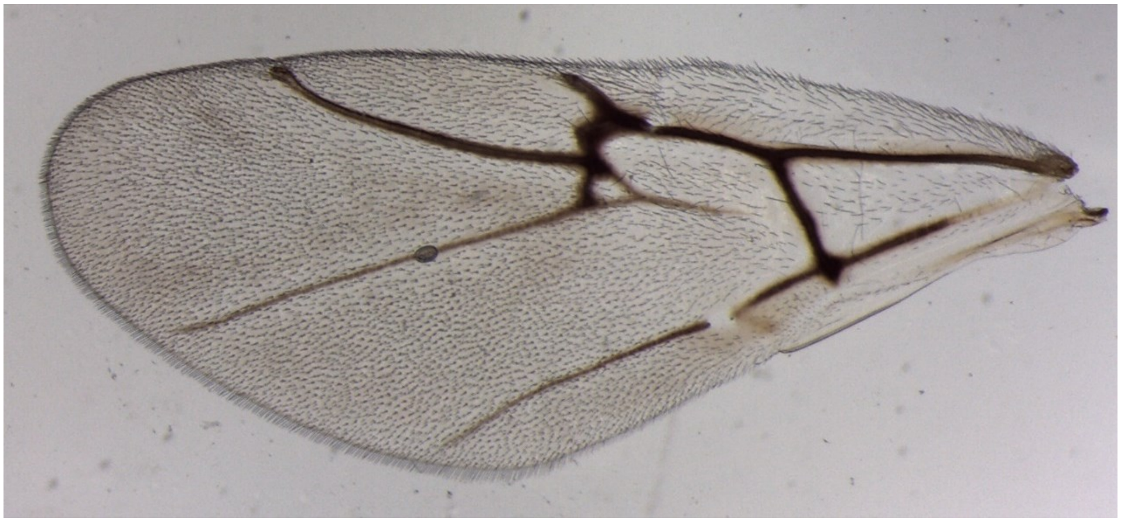

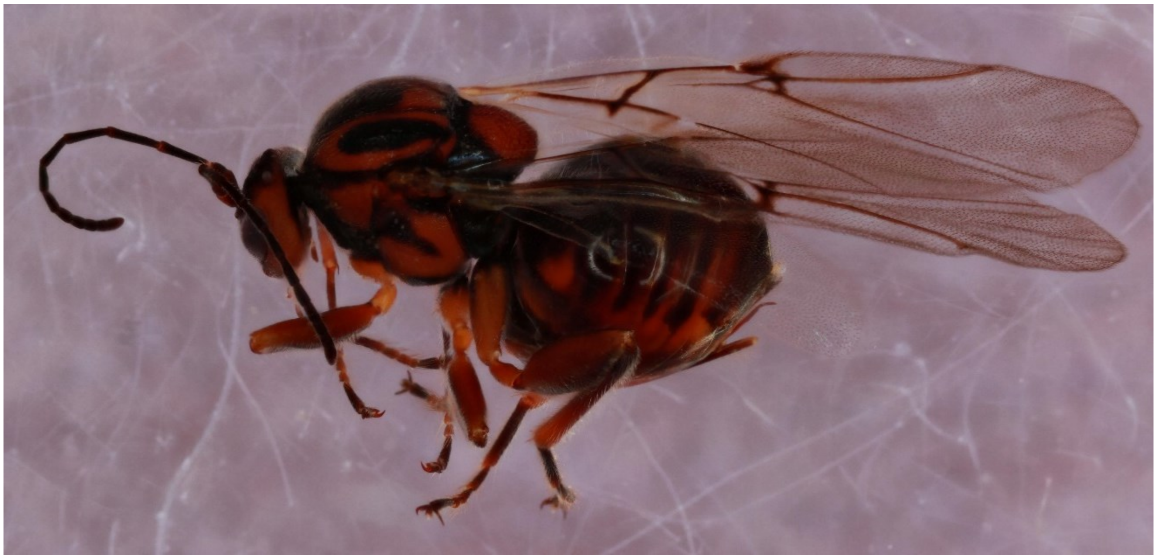

*Disholcaspis* unknown conical stem gall - Wasp: 1679-211-1

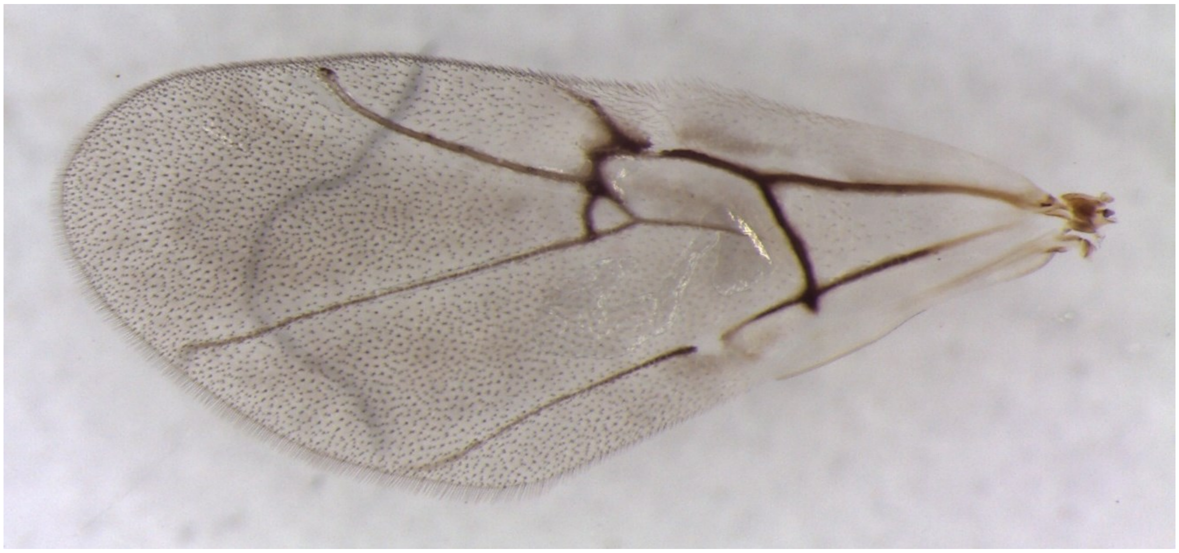

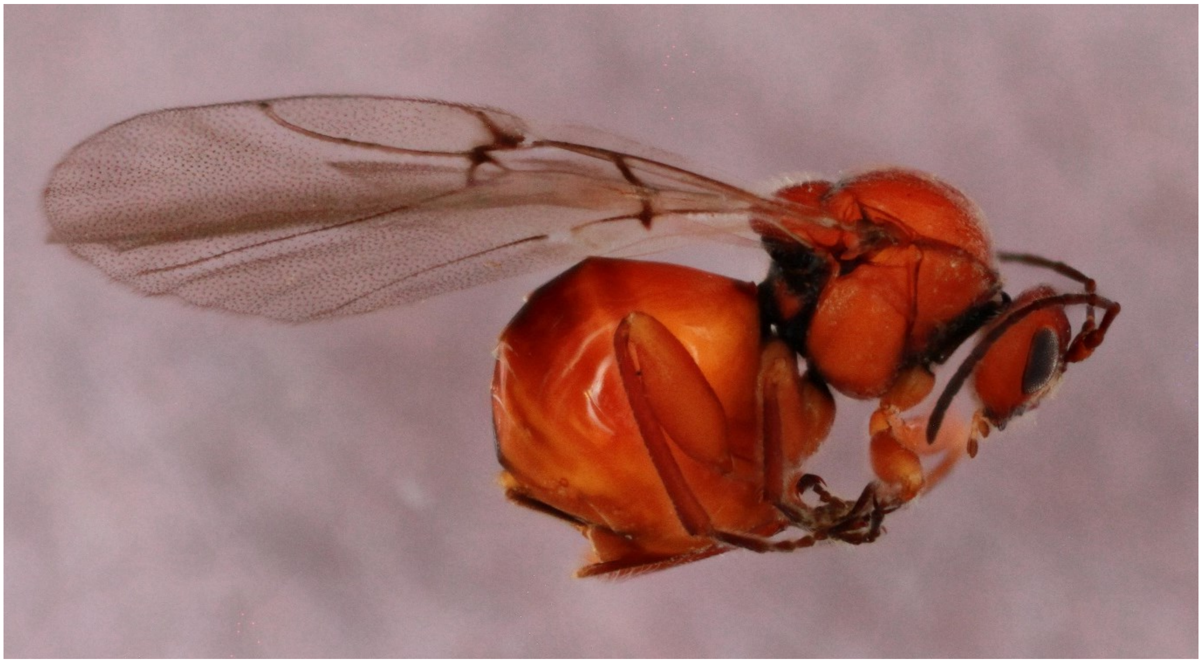

*Druon ignotum* - Wasp: 114-1-35A

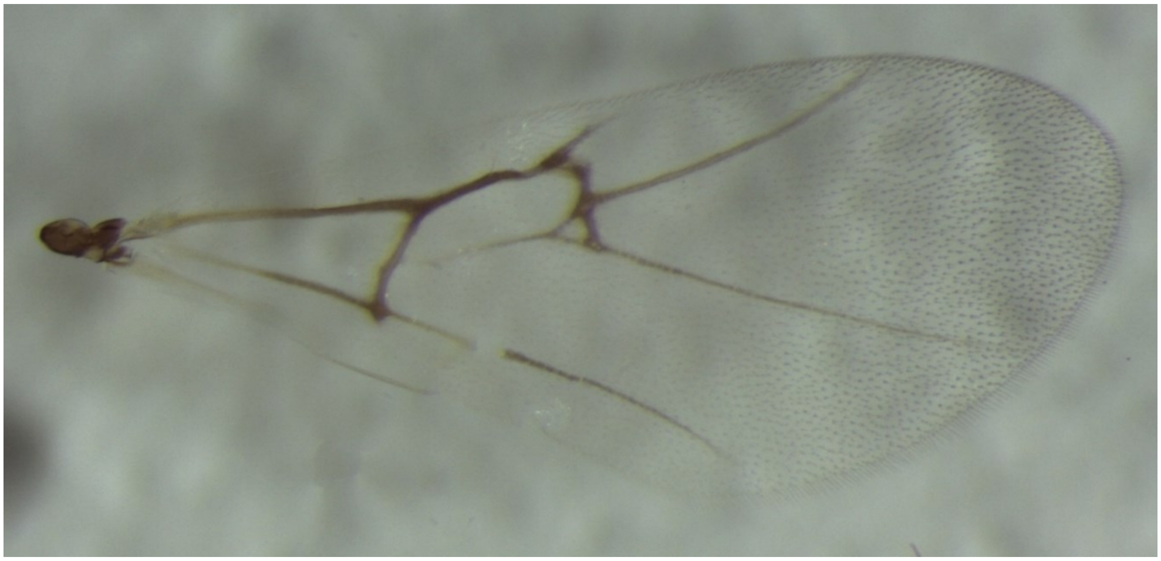

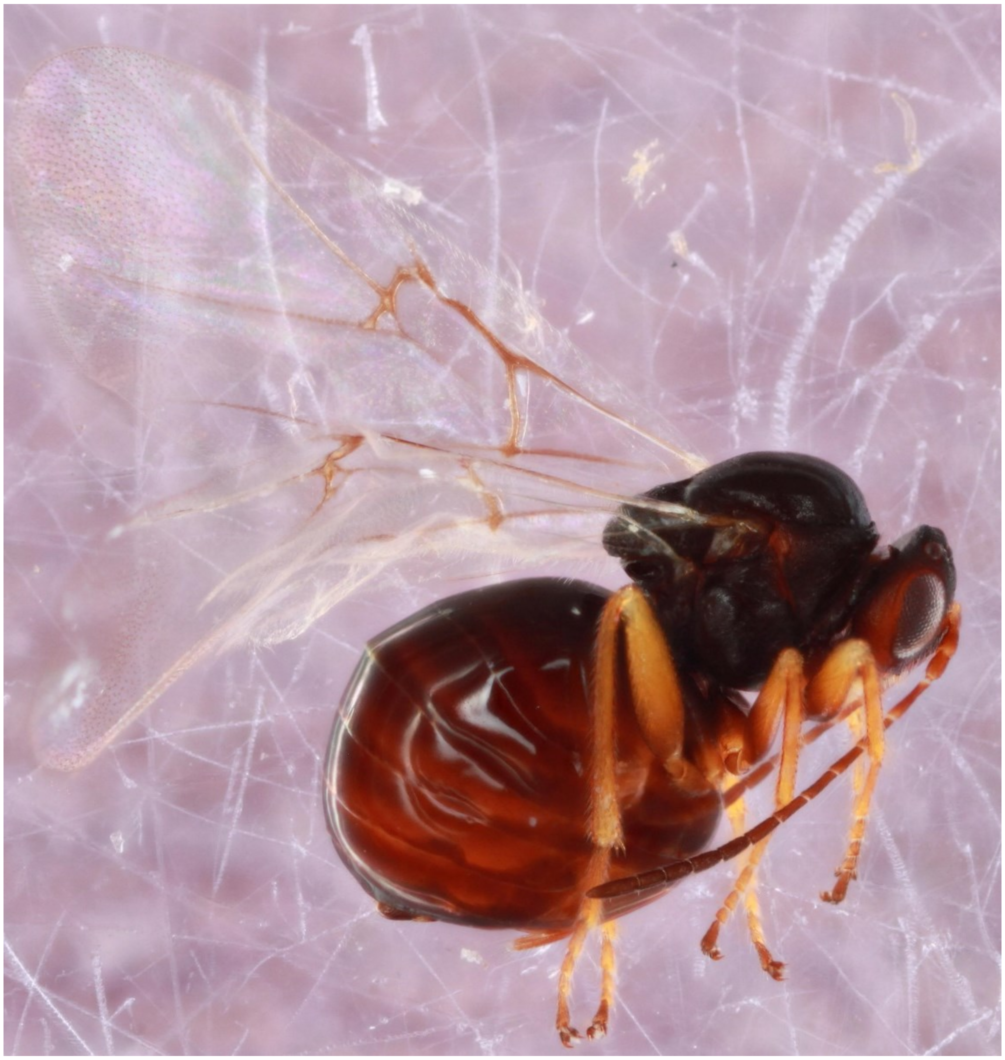

*Druon quercusflocci* - Wasp: 650-061-3

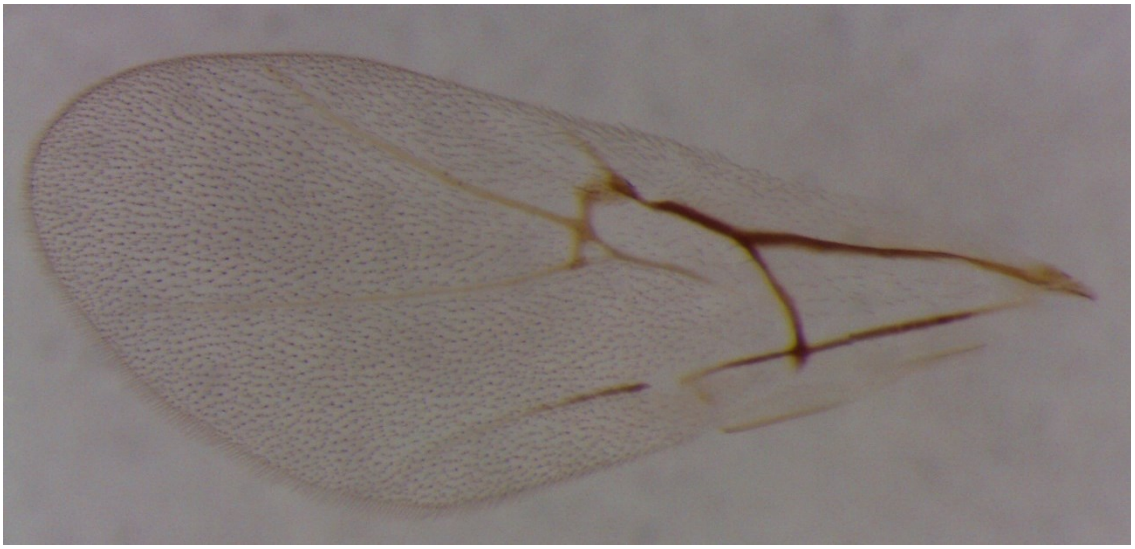

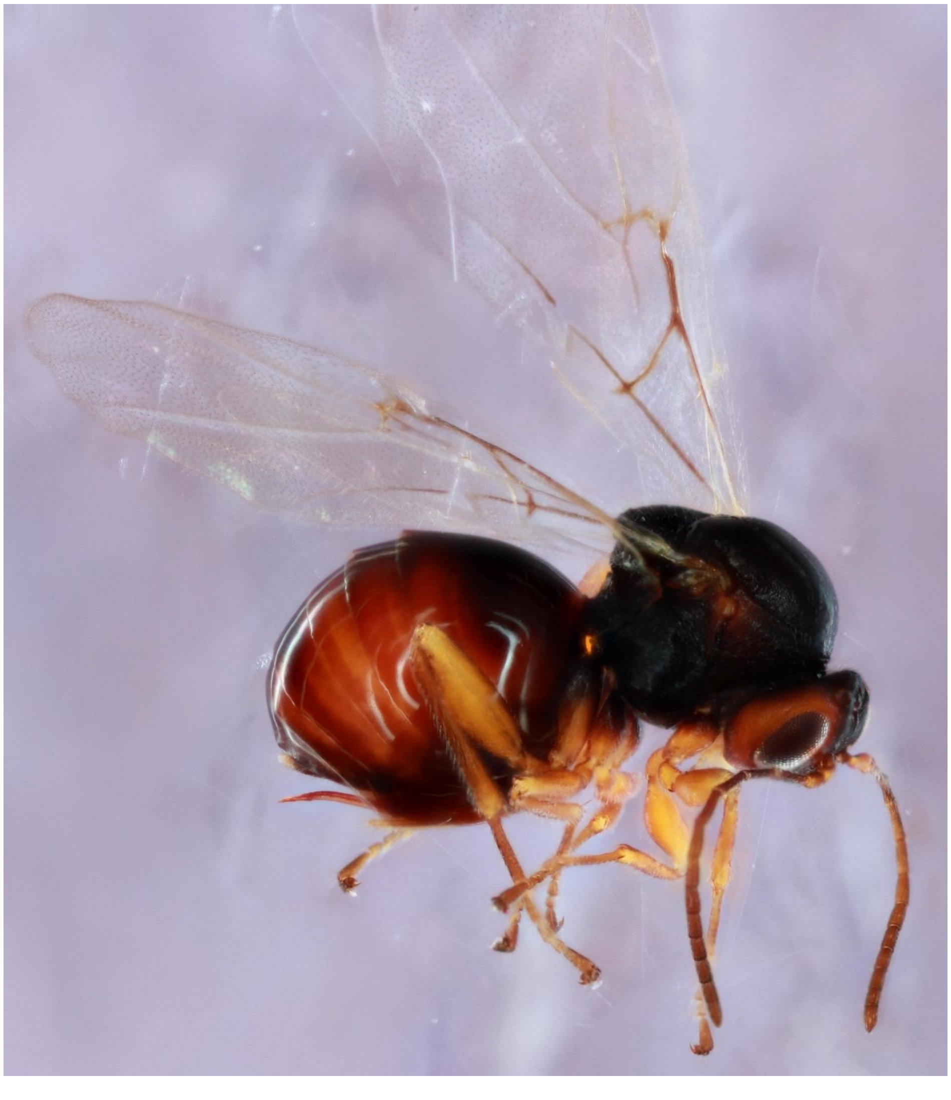

*Dryocosmus minusculus* - Wasp: 1497-141-9

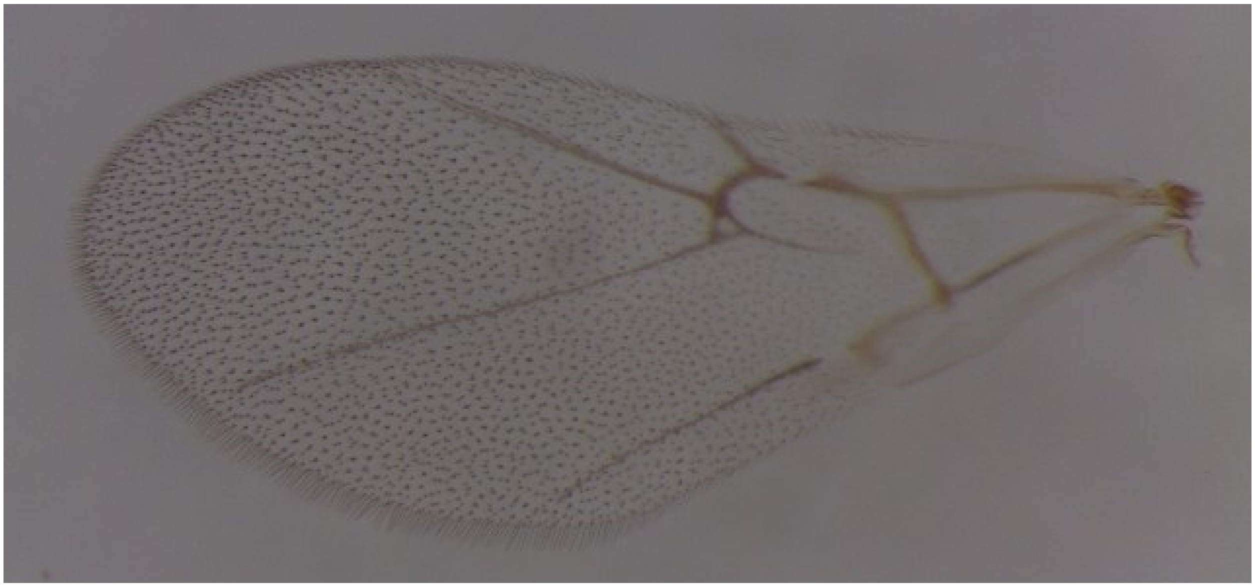

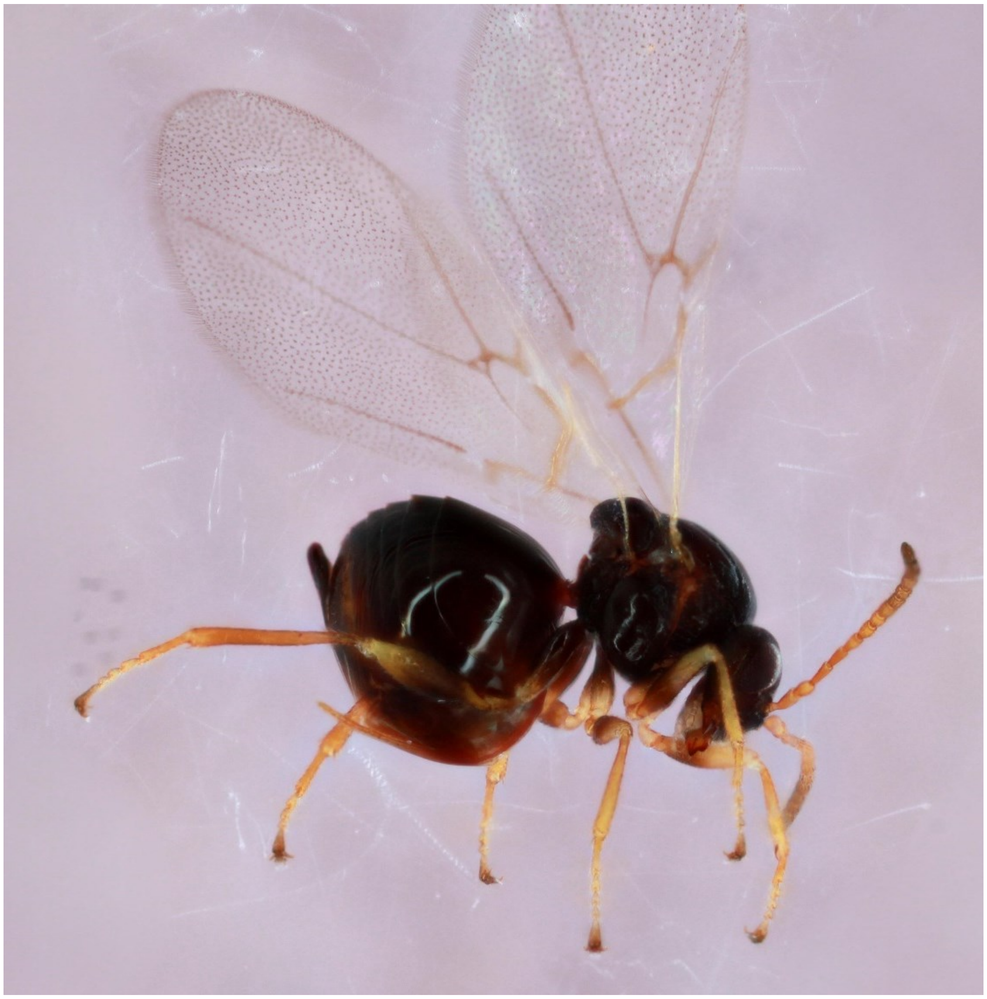

*Dryocosmus quercusnotha* - Wasp: 720-038-001A

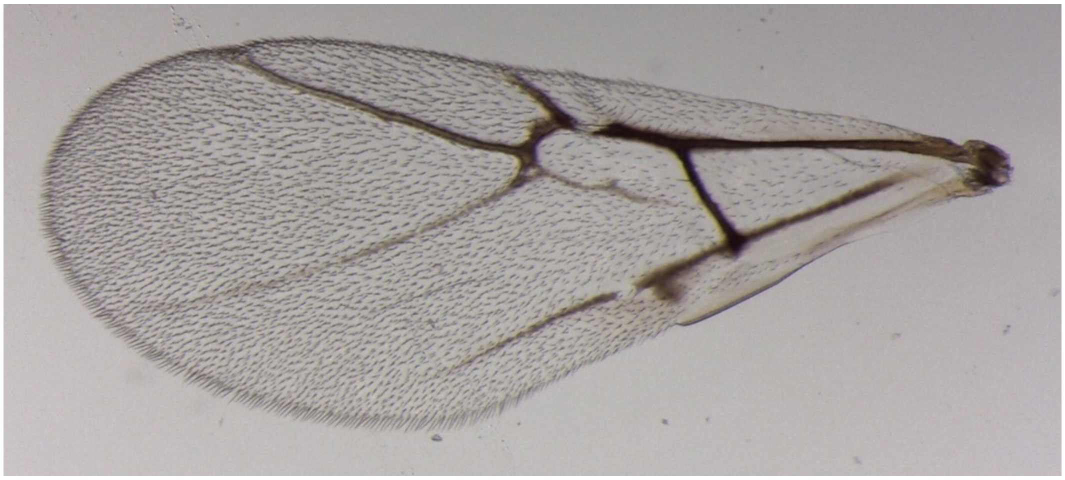

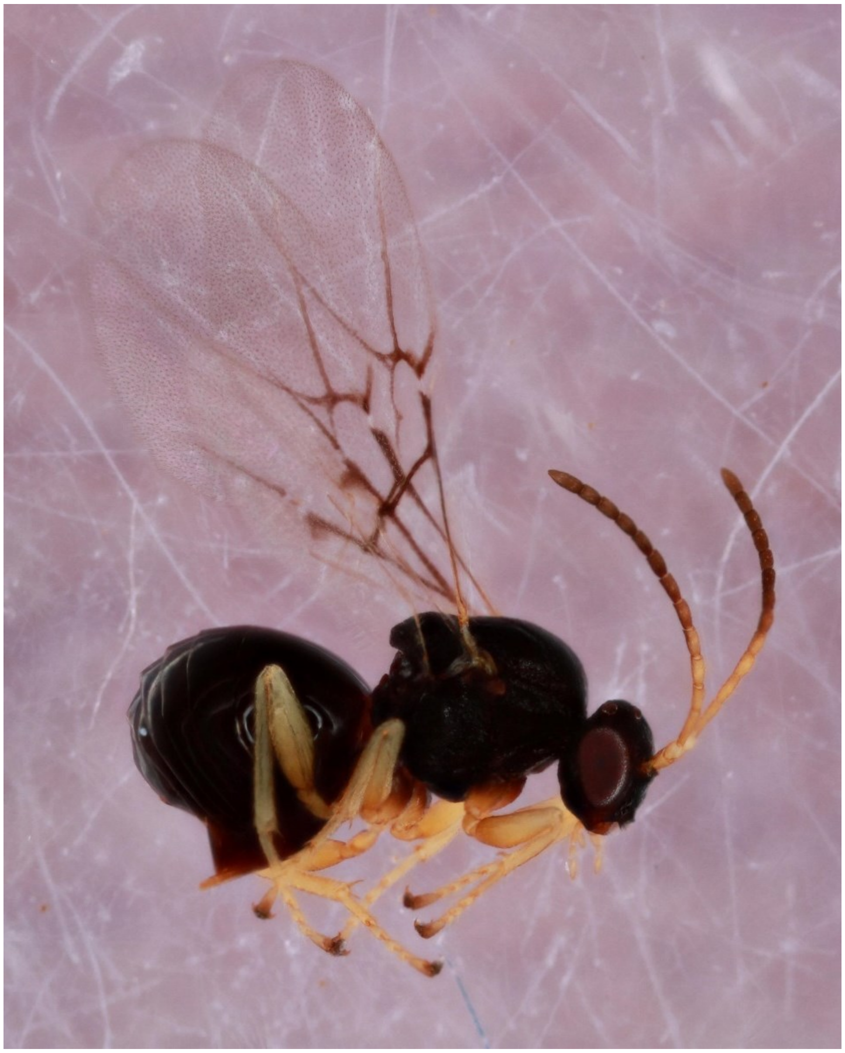

*Dryocosmus quercuspalustris* - Wasp: 720-038-001A

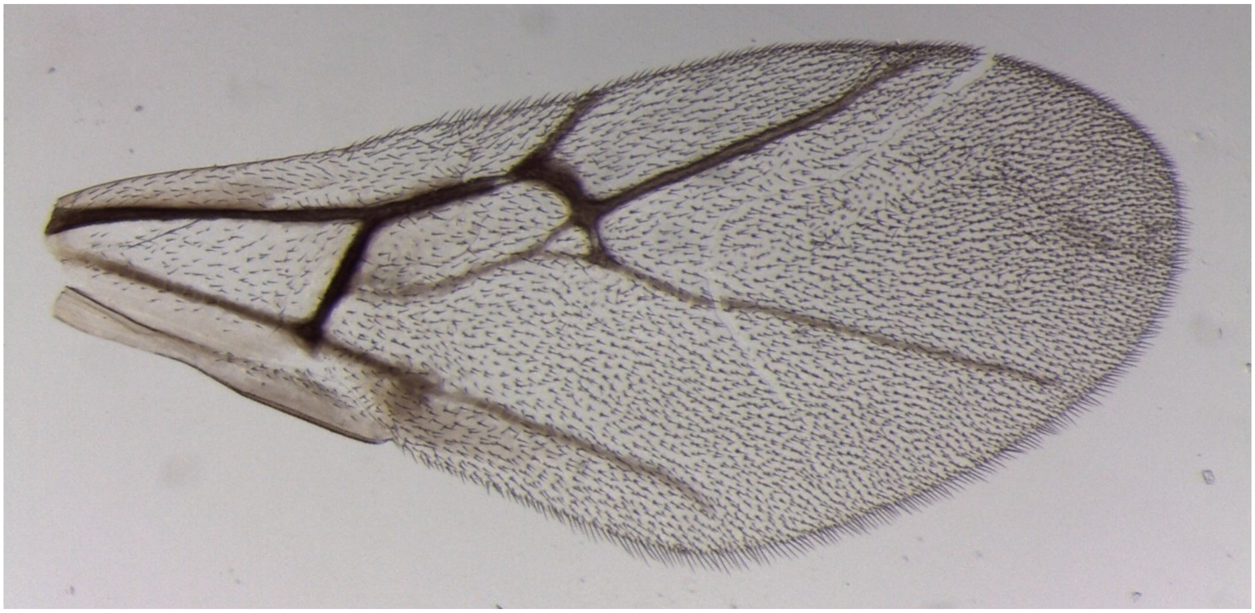

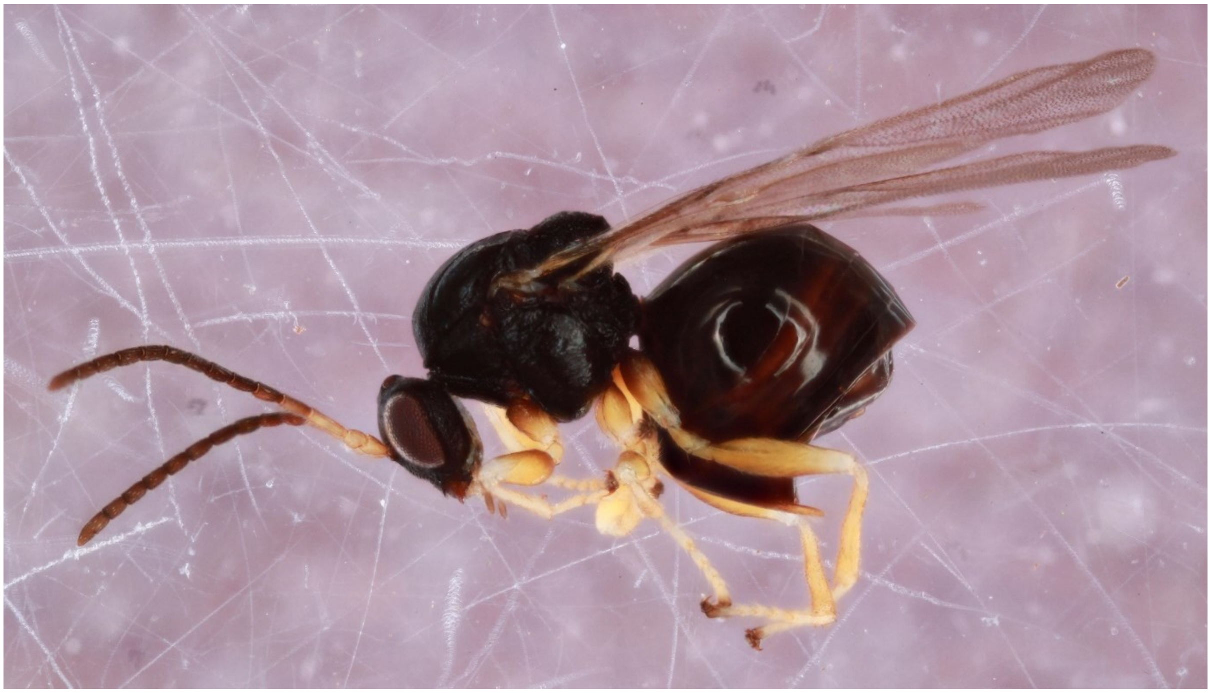

*Kokkocynips imbricariae* - Wasp: 1000-059-1

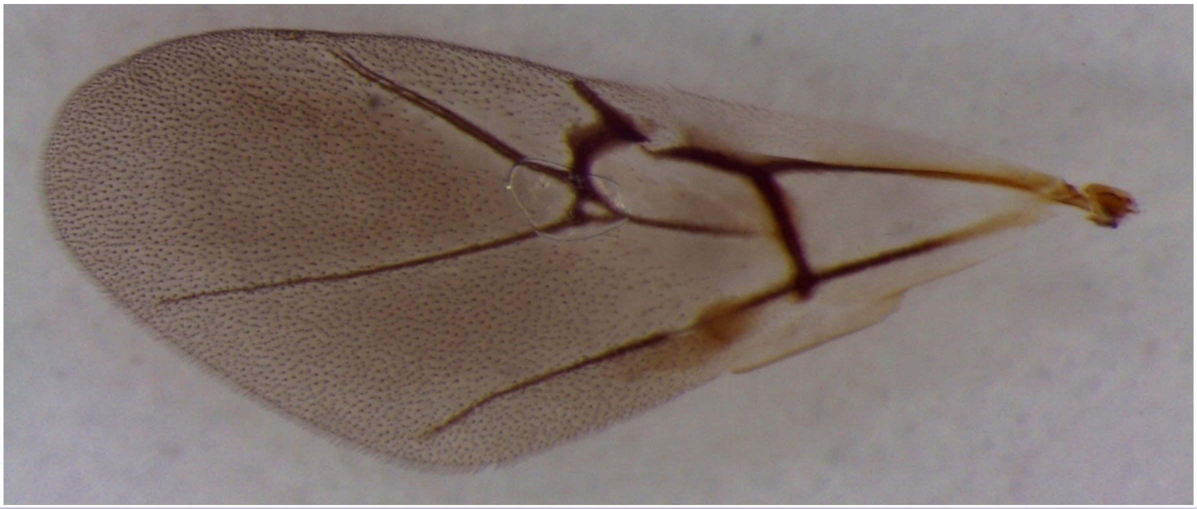

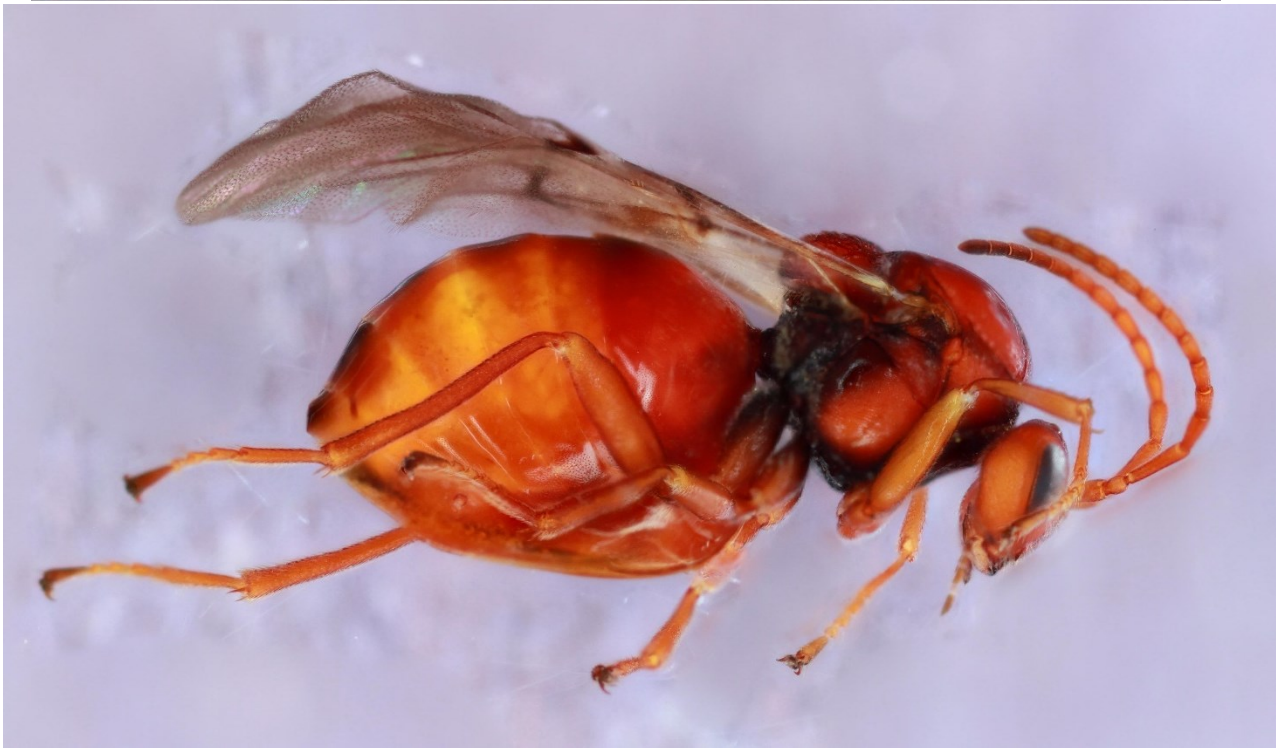

*Loxaulus quercusmammula* - Wasp: 866-035-017

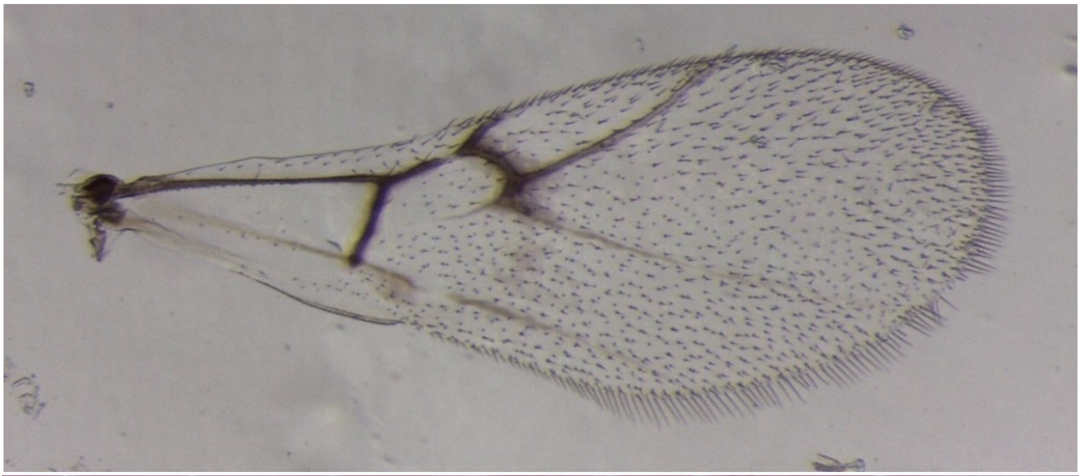

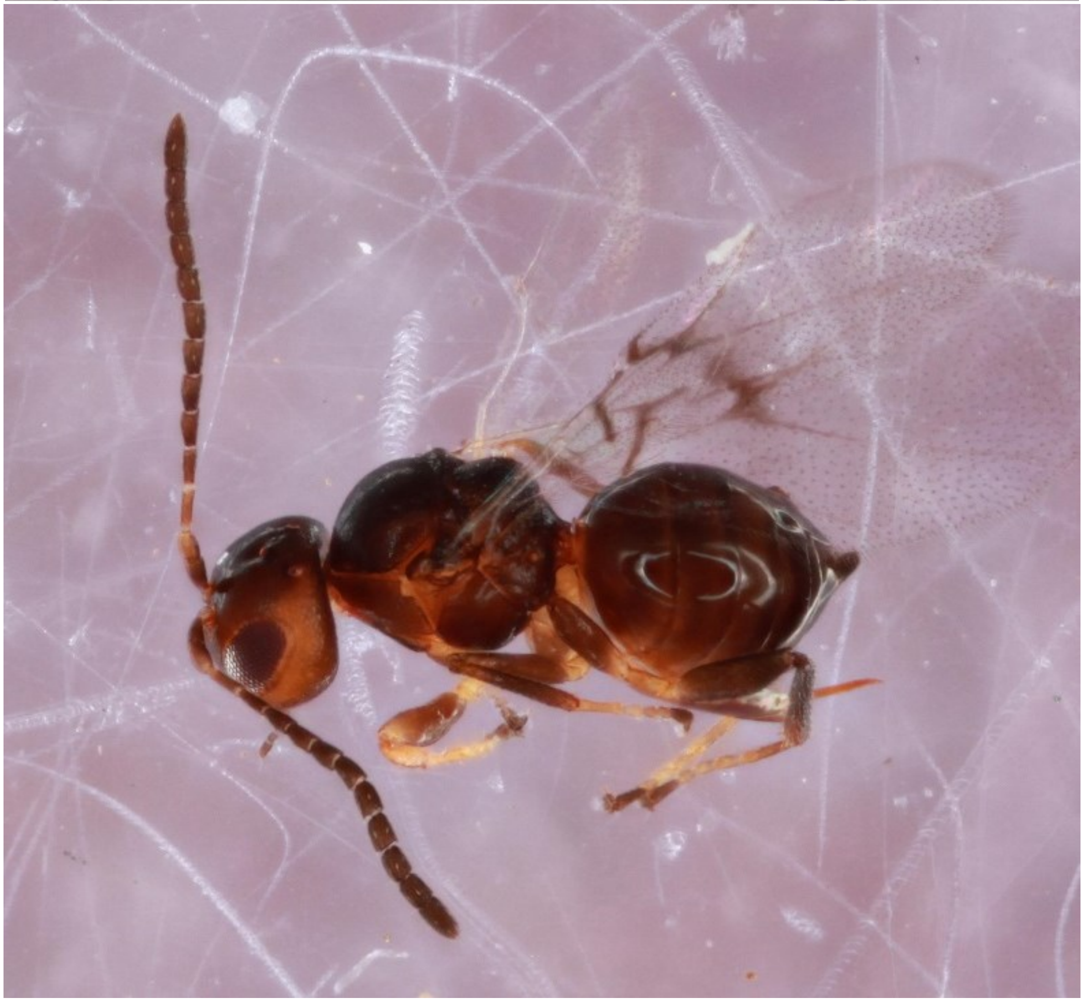

*Melikaiella ostensackeni* - Wasp: 876-016-011A

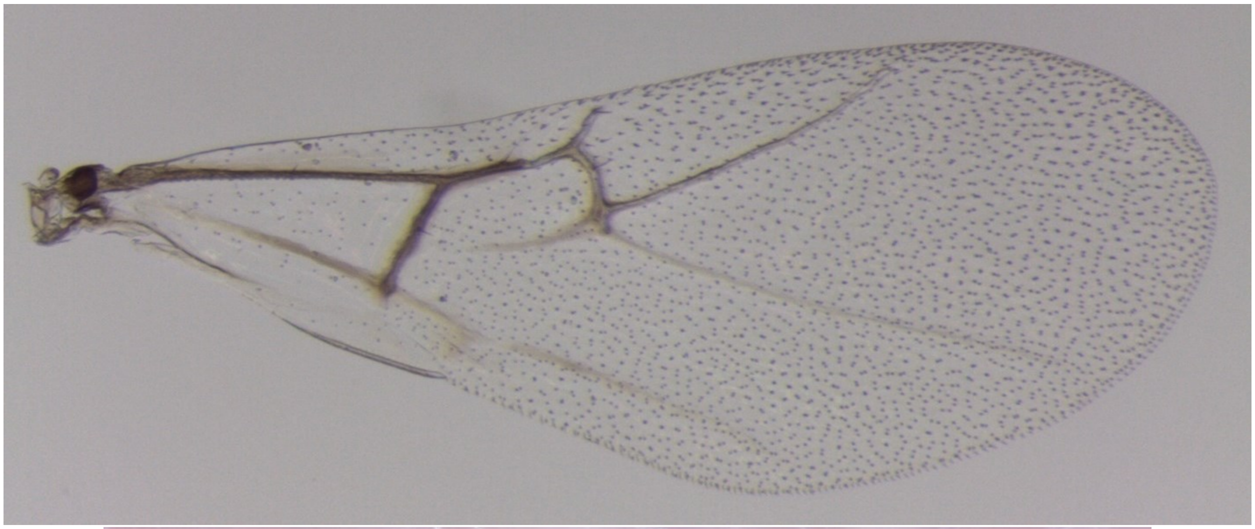

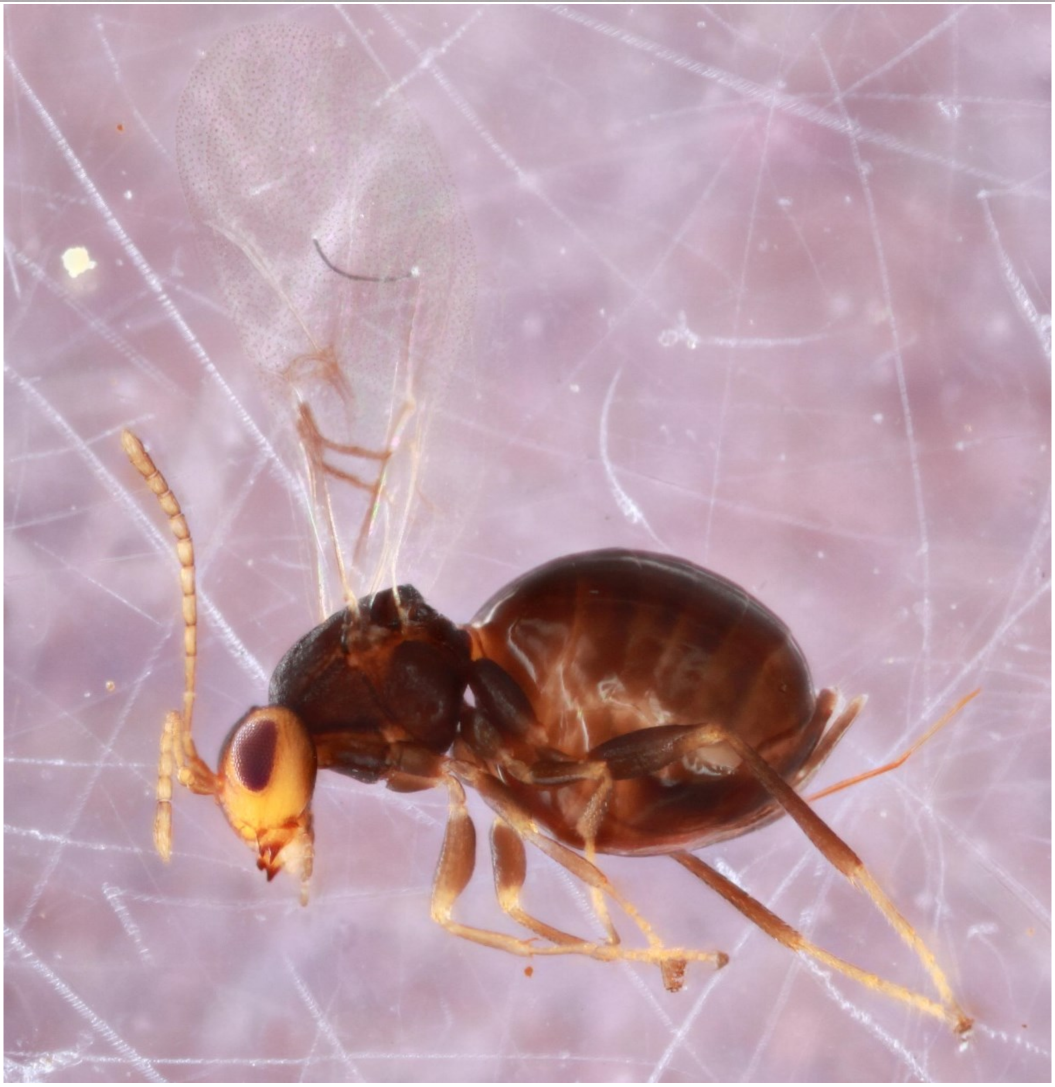

*Melikaiella tumifica* - Wasp: 810-043-002

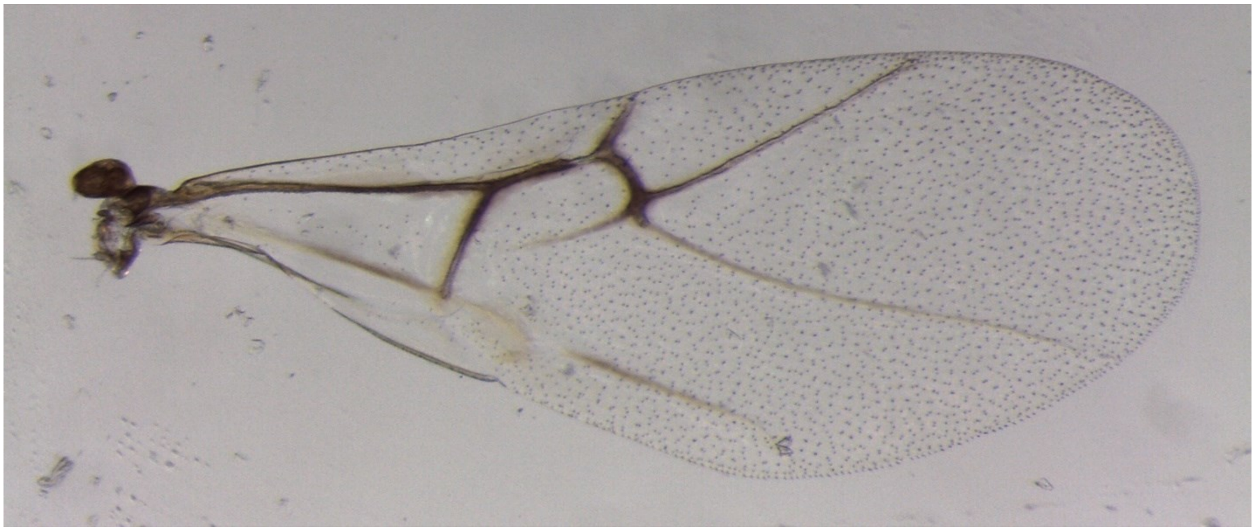

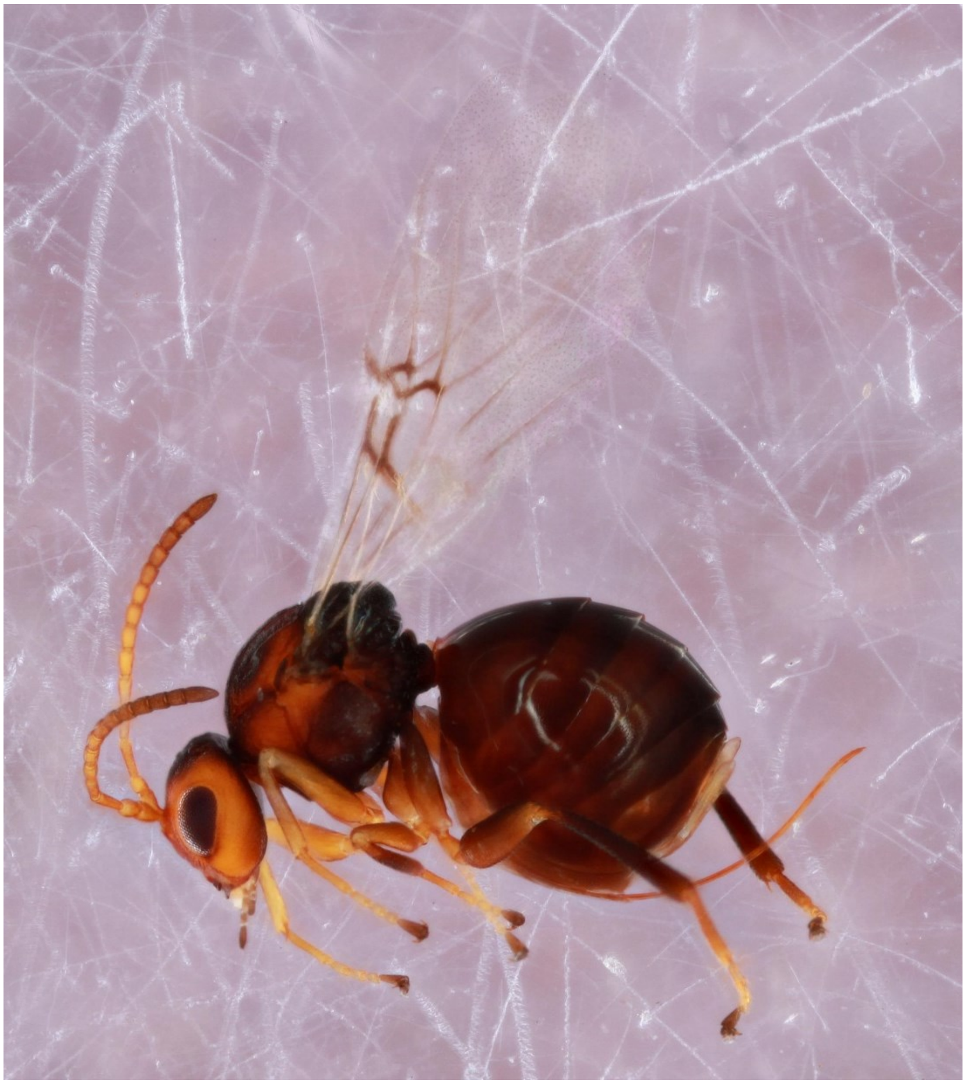

*Neuroterus fragilis* - Wasp: Nf_3

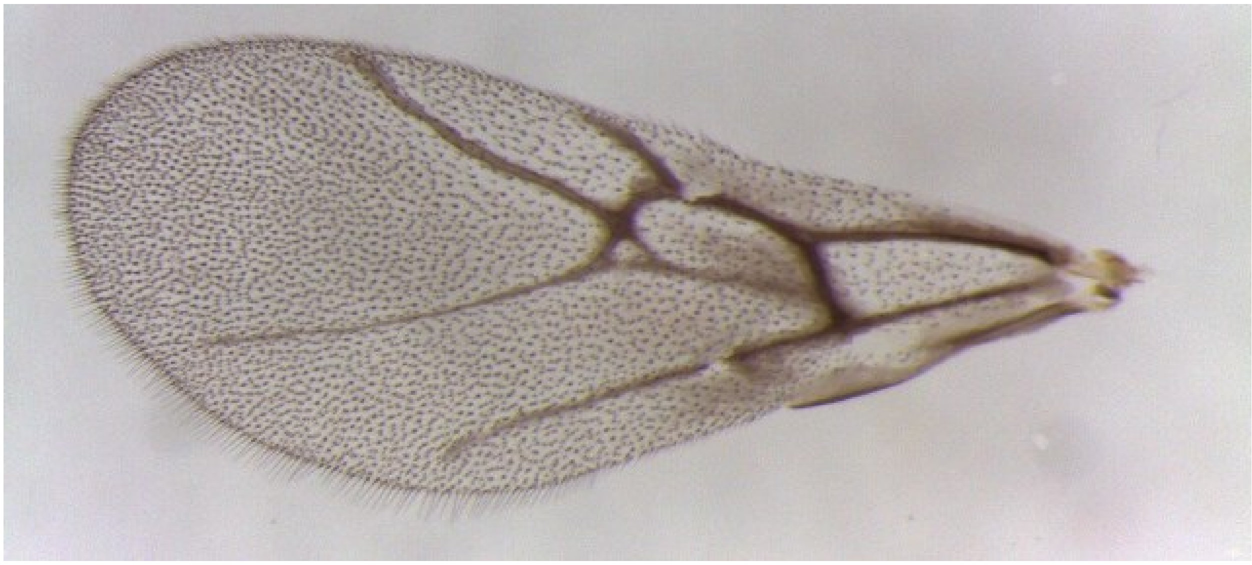

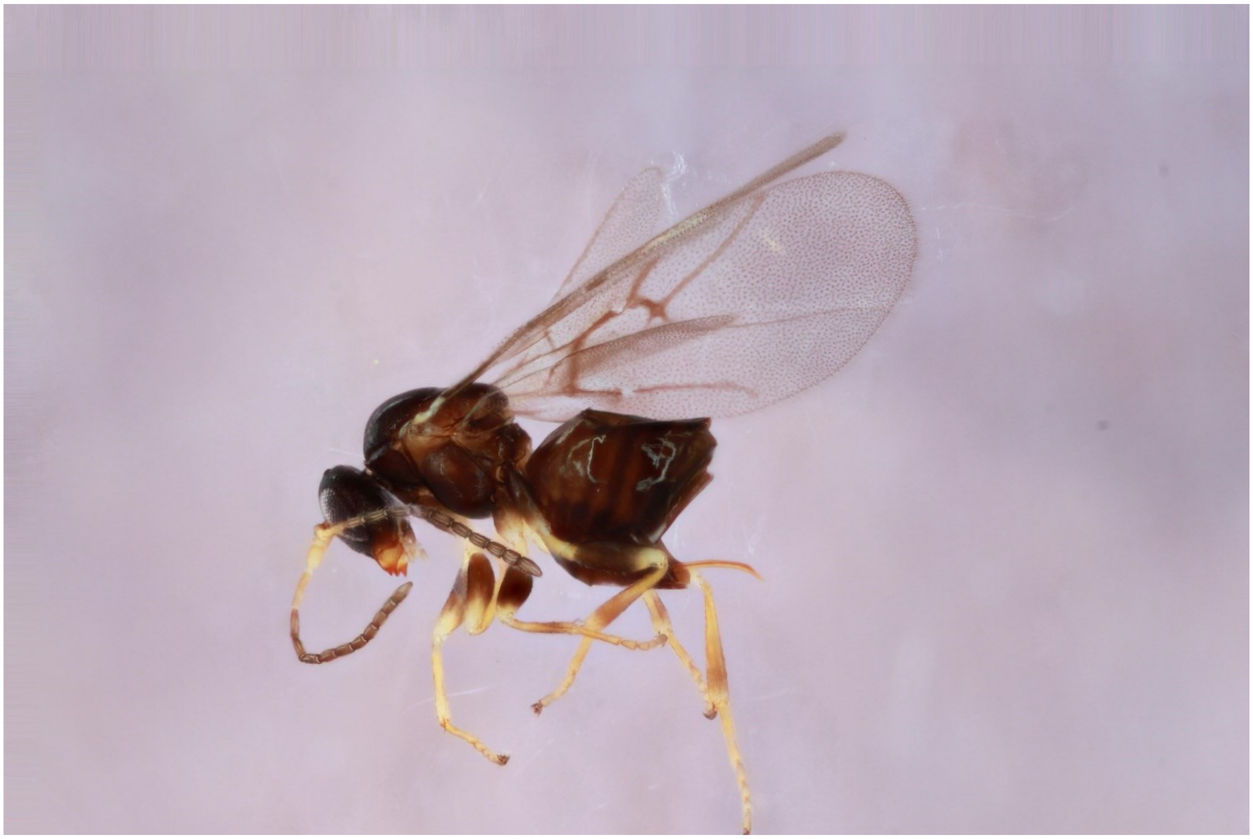

*Neuroterus pallidus* - Wasp: 1312-089-1

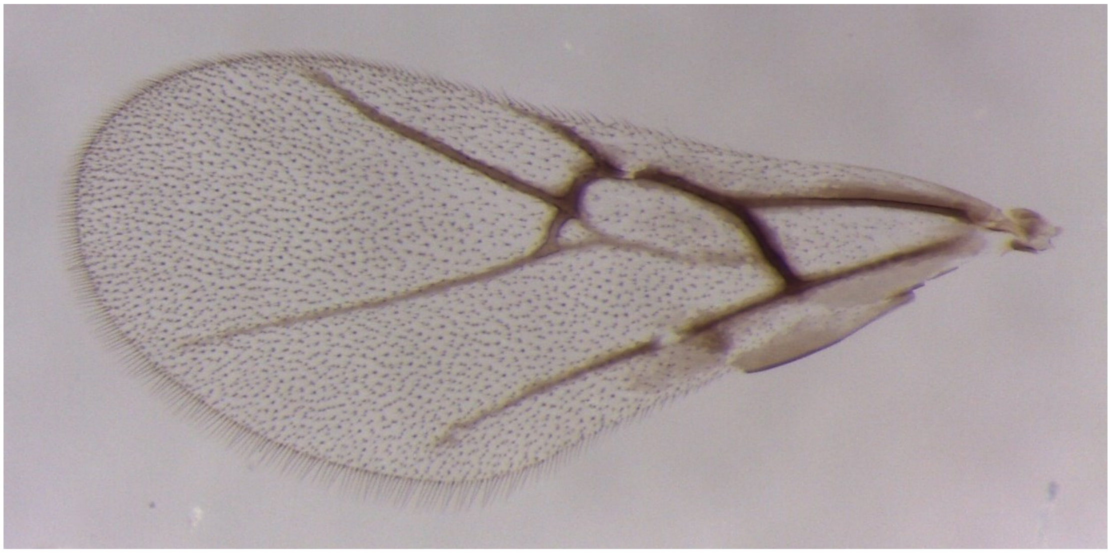

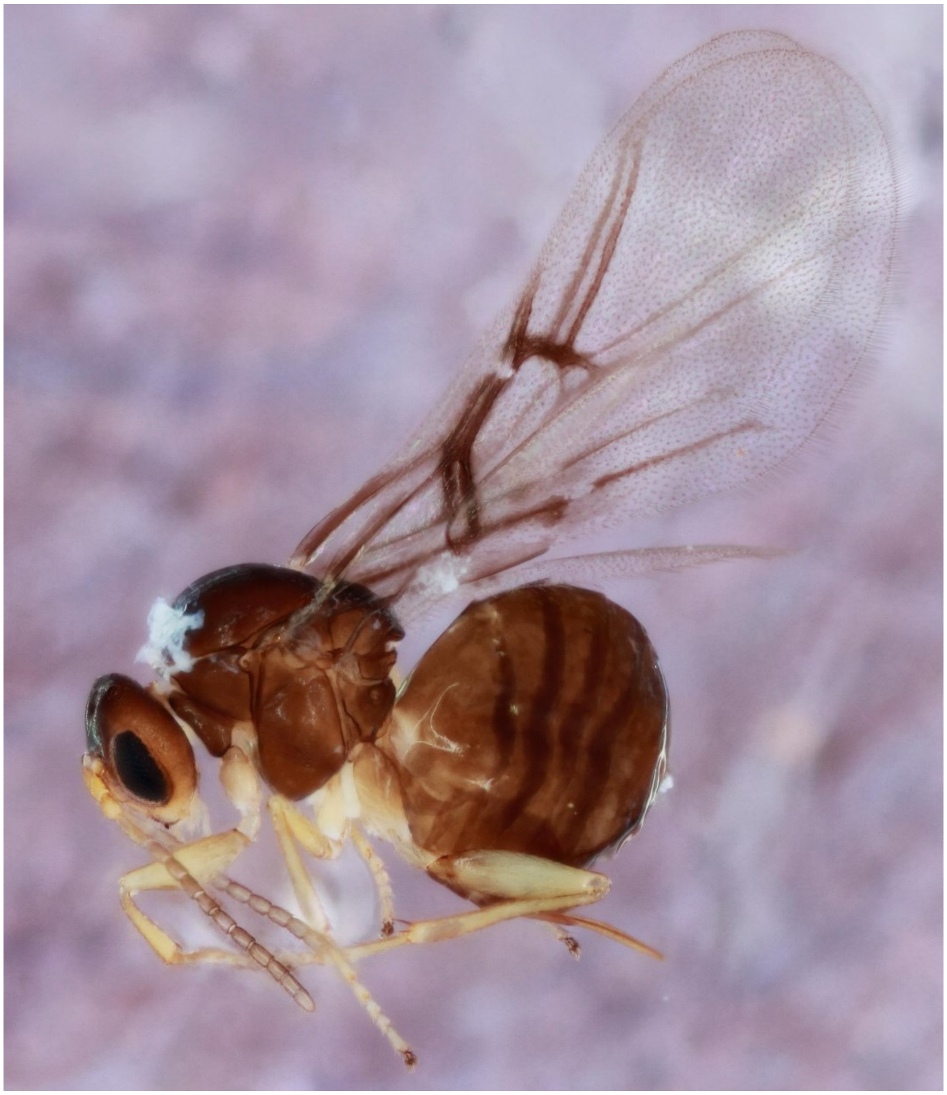

*Neuroteus* q_macrocarpa_flower_swelling_gall - Wasp: 742-088-3A

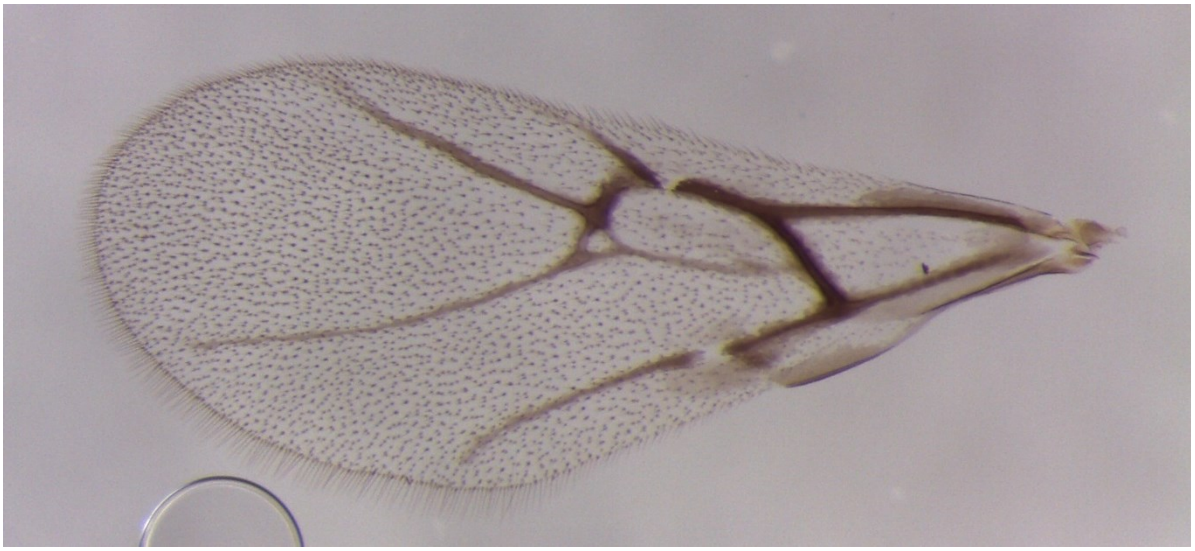

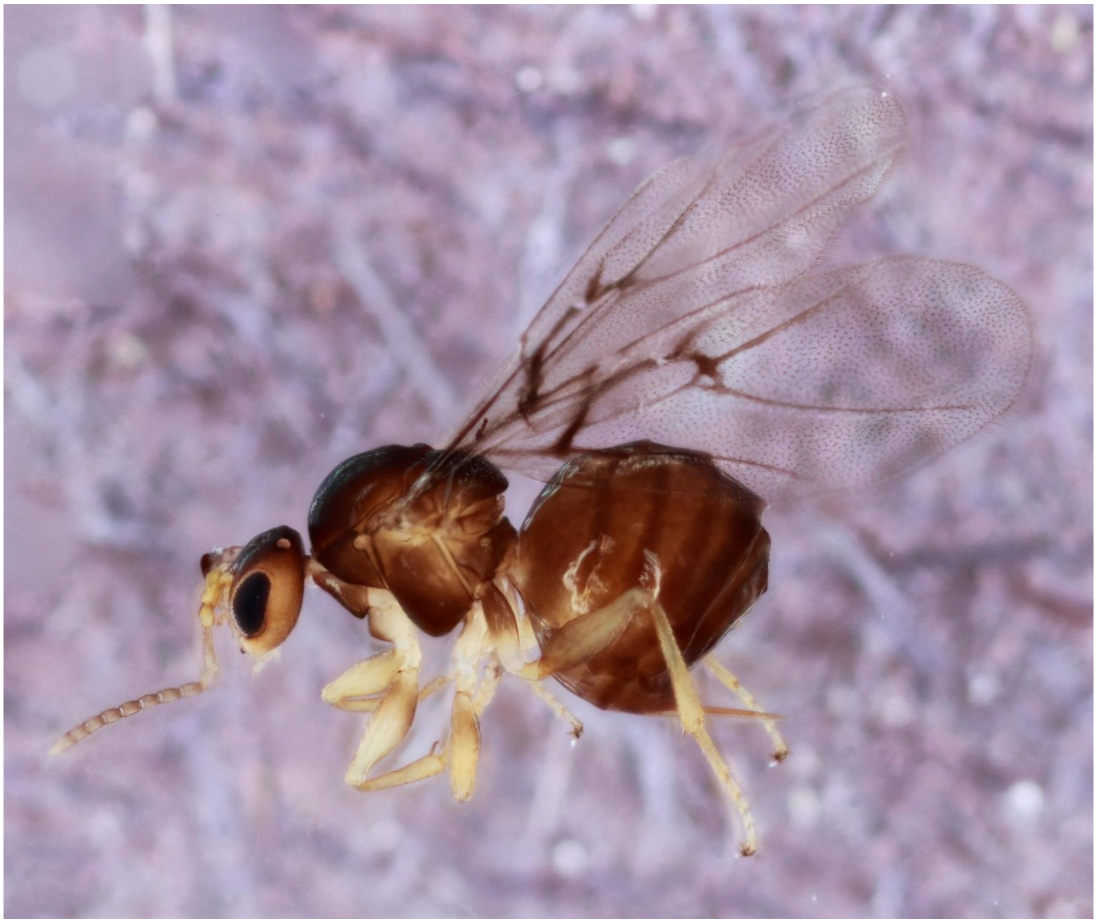

*Neuroterus quercusbatatus* - Wasp: 341-053-8

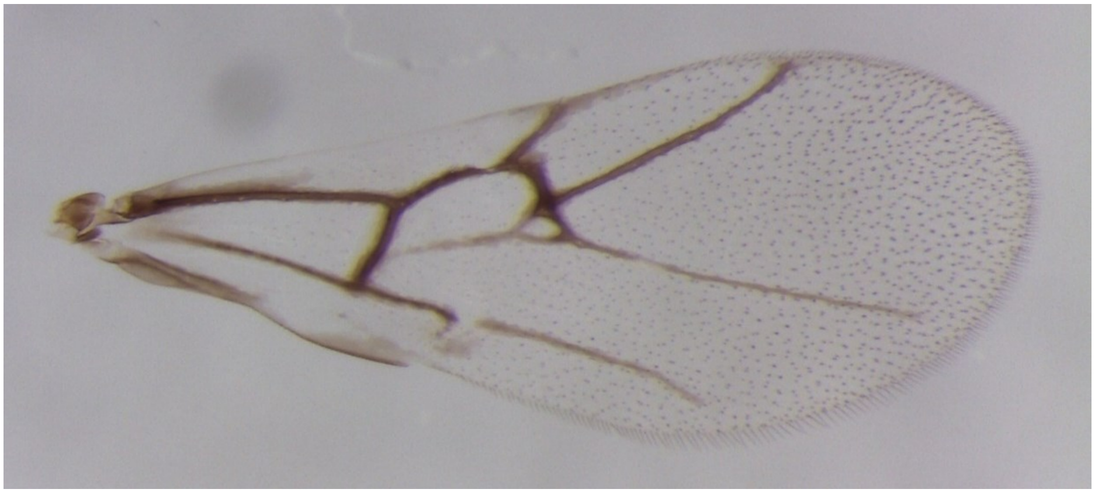

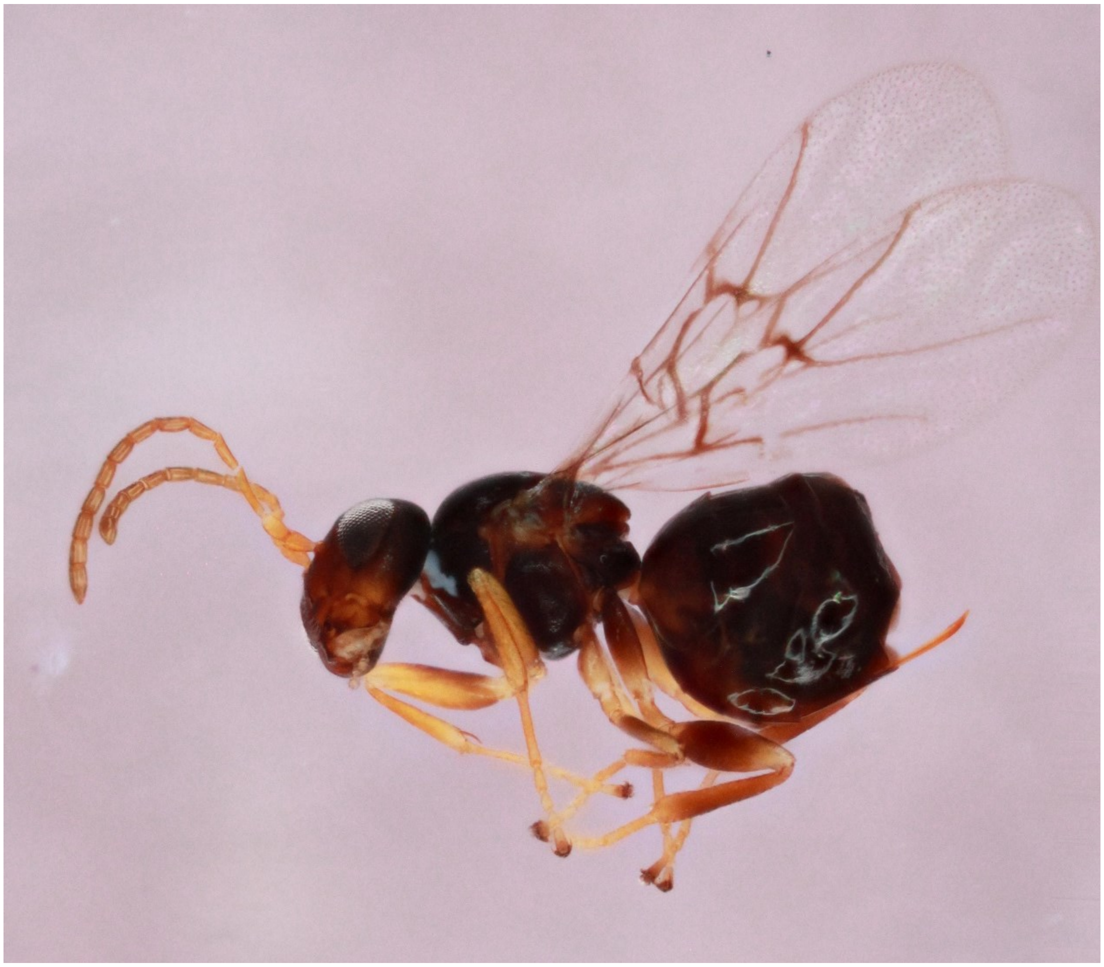

*Neuroterus quercusirregularis* - Wasp: 776-044-001A

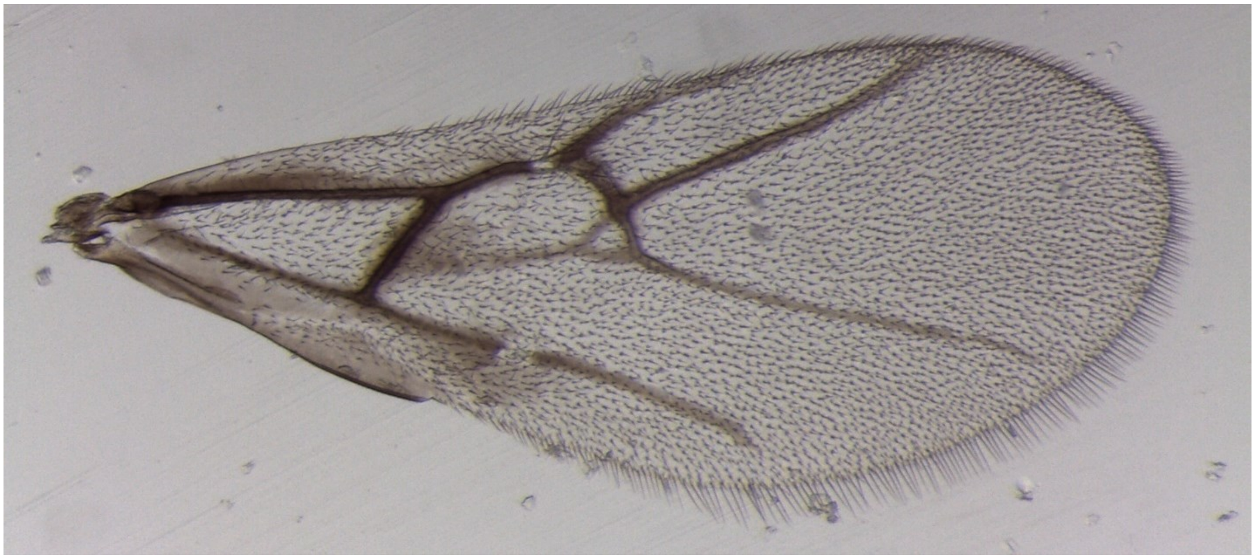

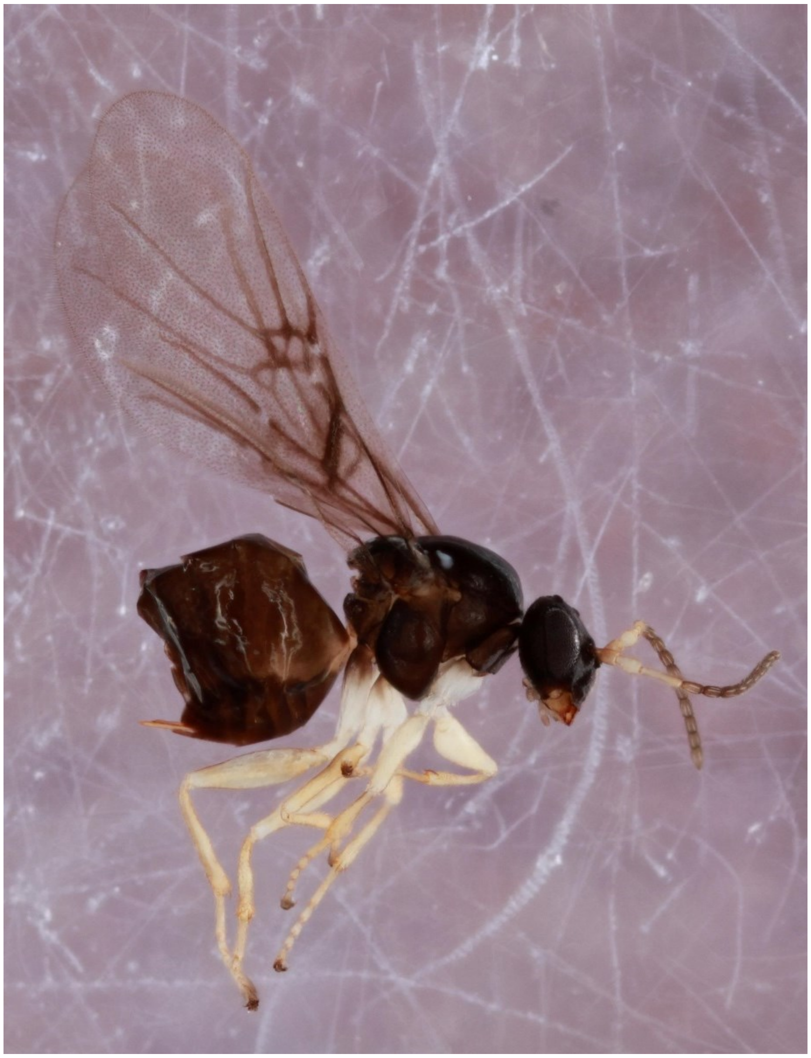

*Neuroterus quercusminutissimus* - Wasp: LZ_972_2

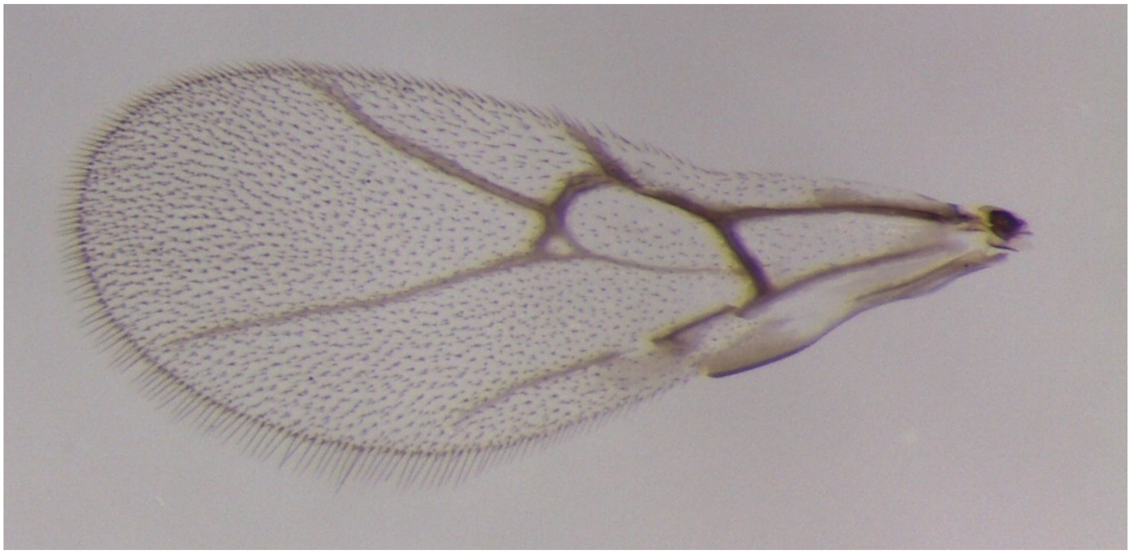

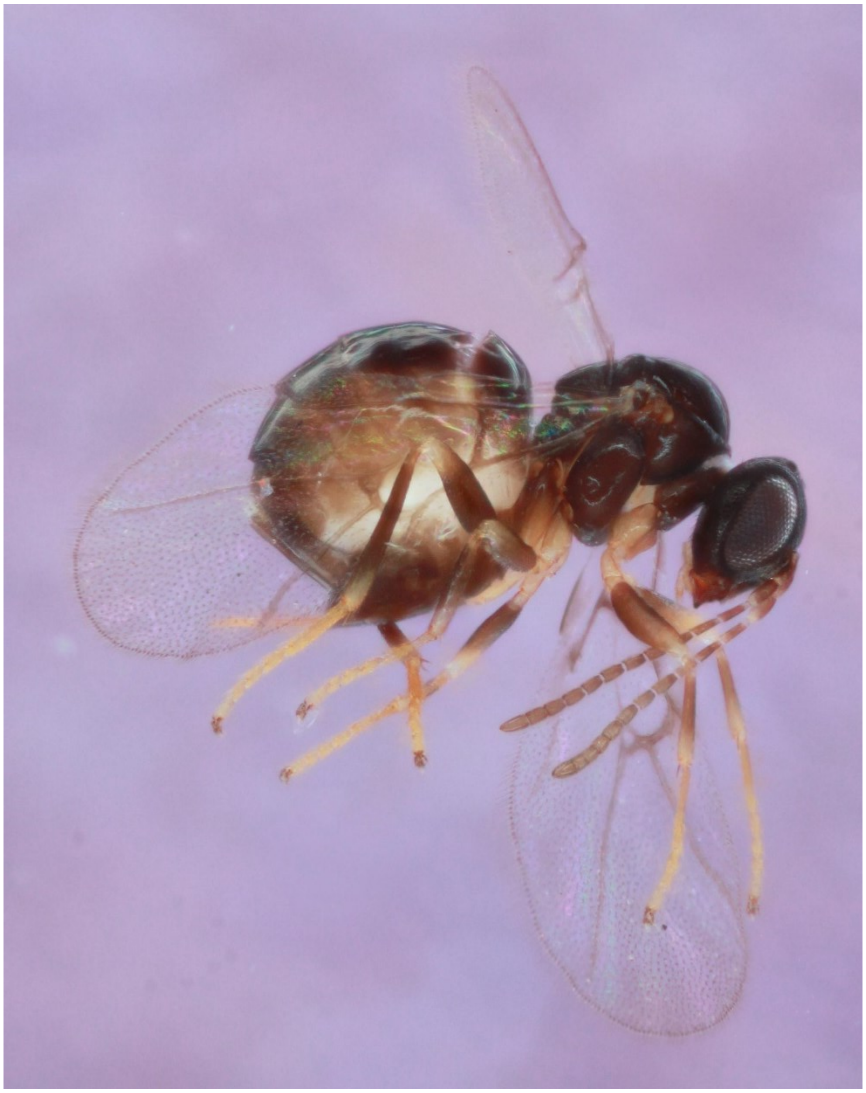

*Neuroterus quercusverrucarum* - Wasp: 1210-022-16A

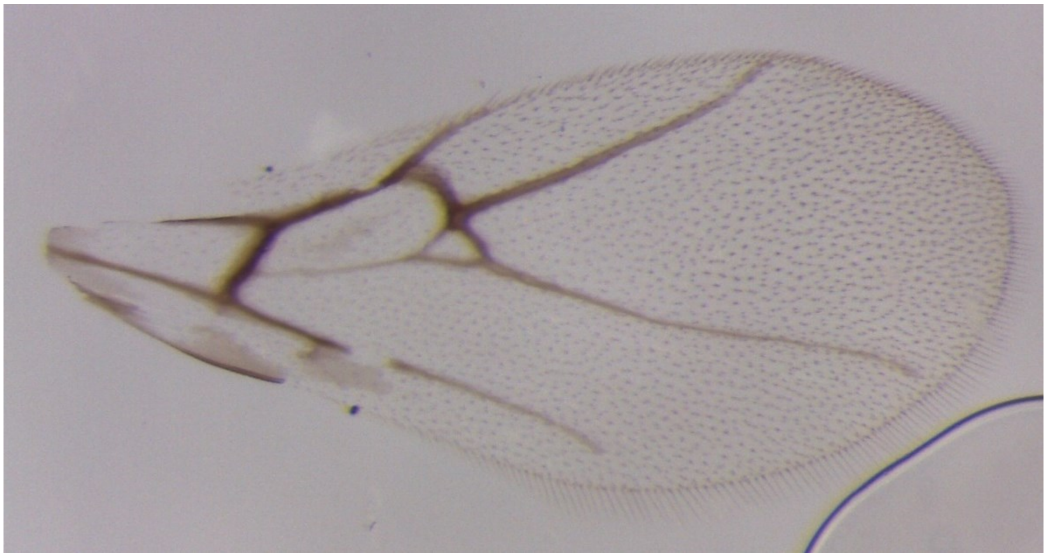

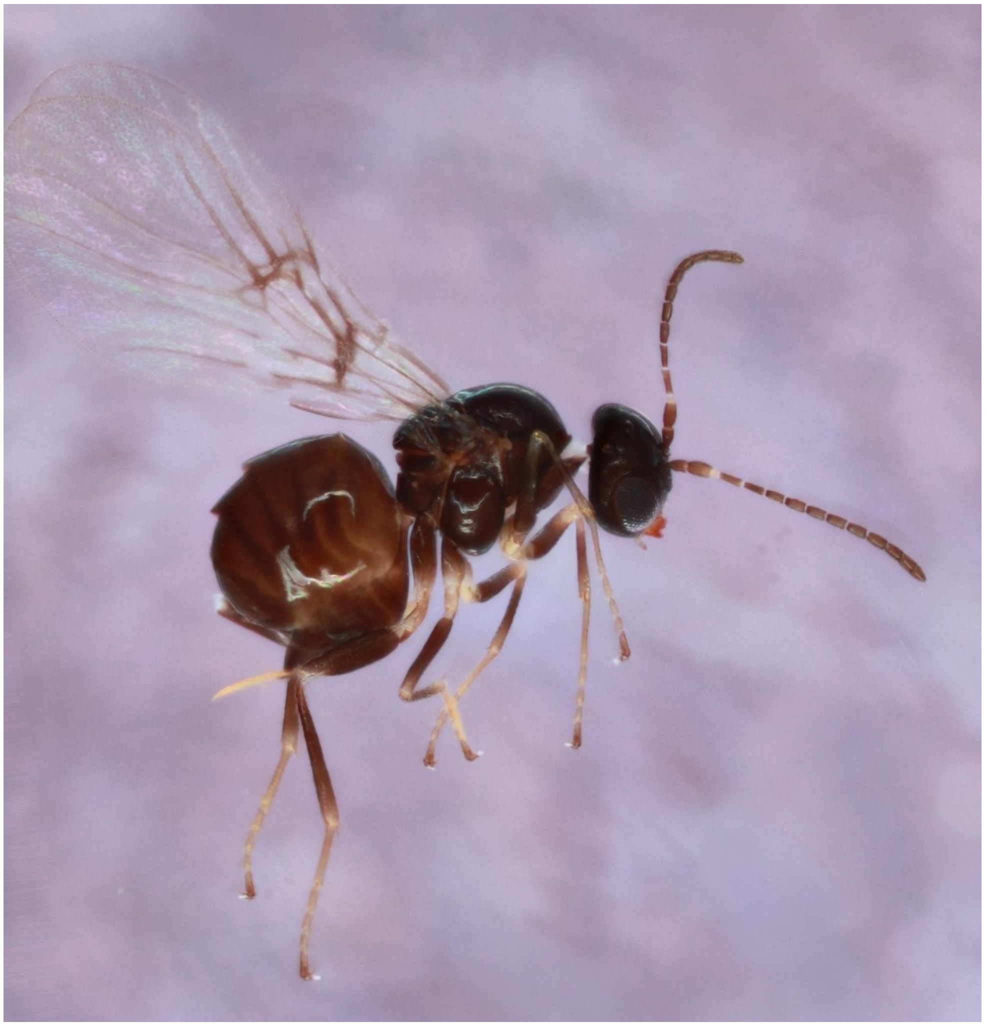

*Neuroterus saltatorius* - Wasp: Nsg1

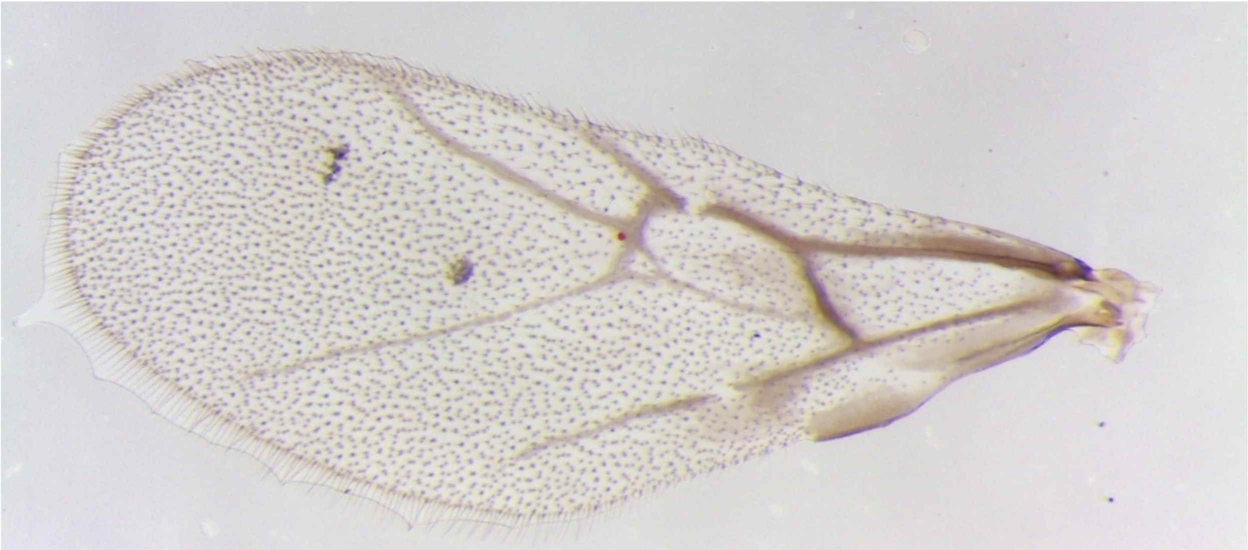

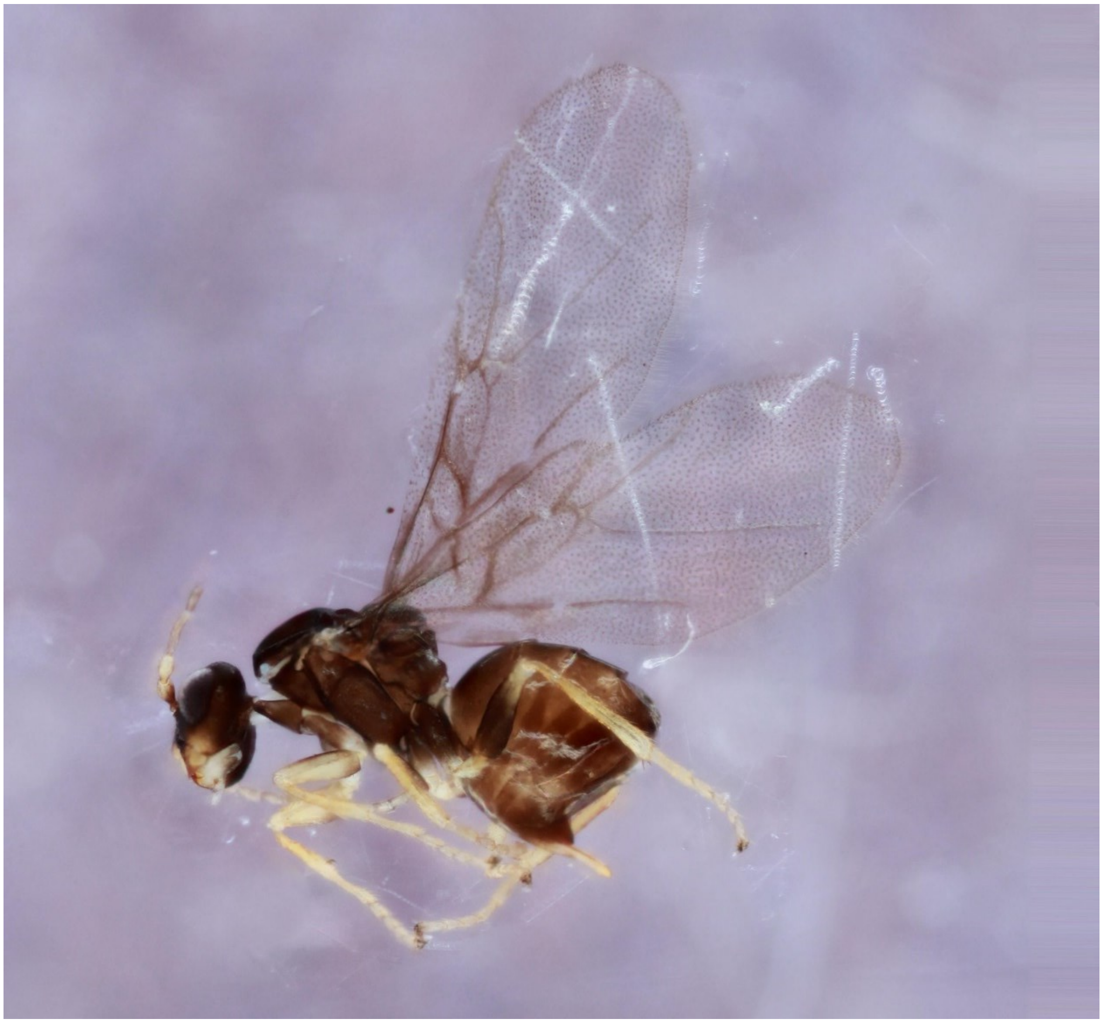

*Neuroterus saltatorius texanus* - Wasp: 1593-163-2

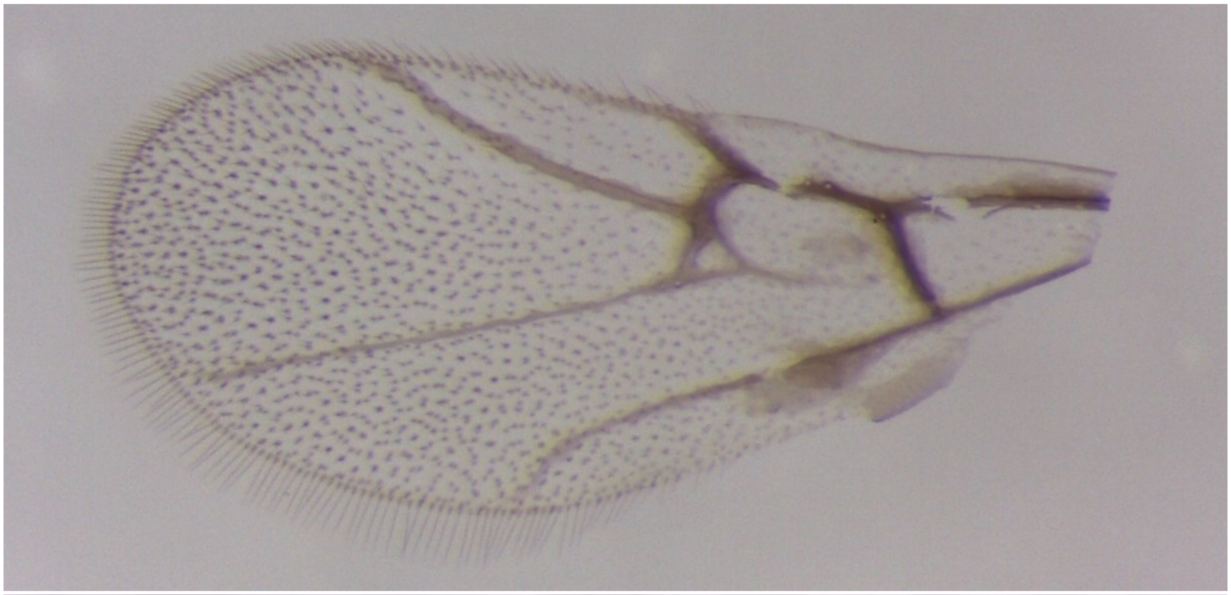

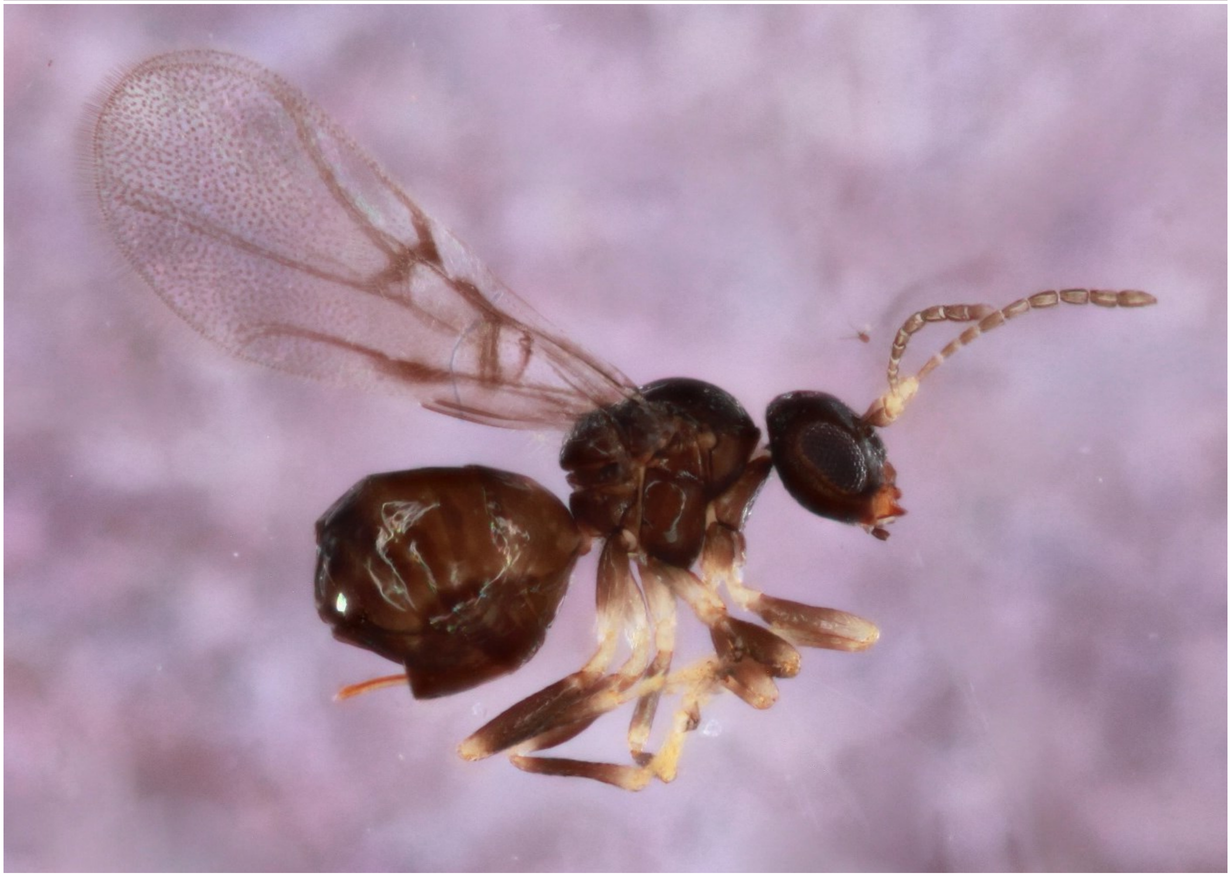

*Neuroterus vesicula* - Wasp: 694-033-001

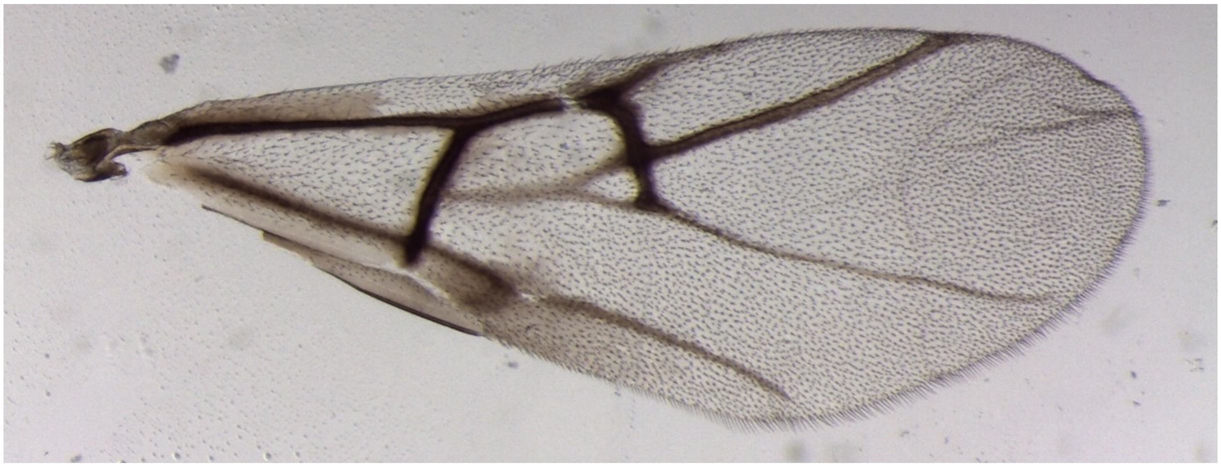

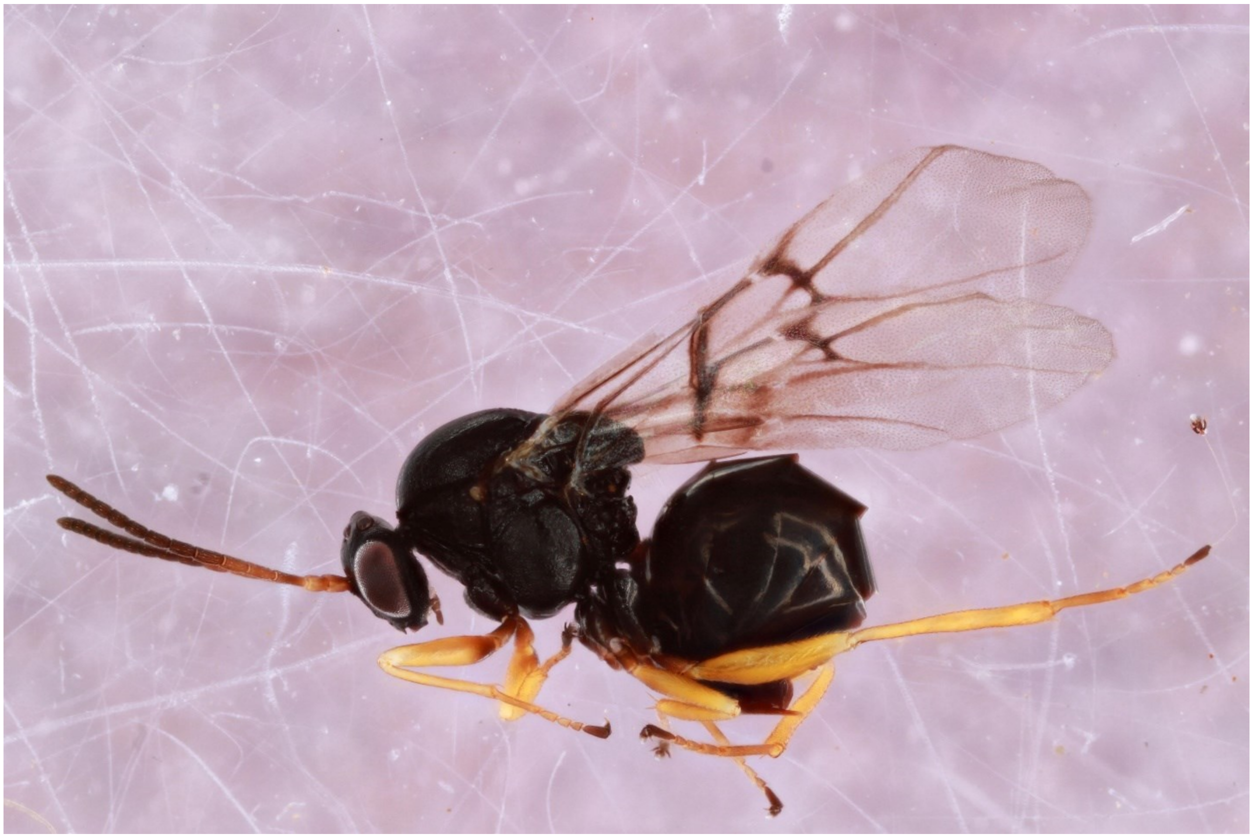

*Neuroterus washingtonensis* - Wasp: Nw_2

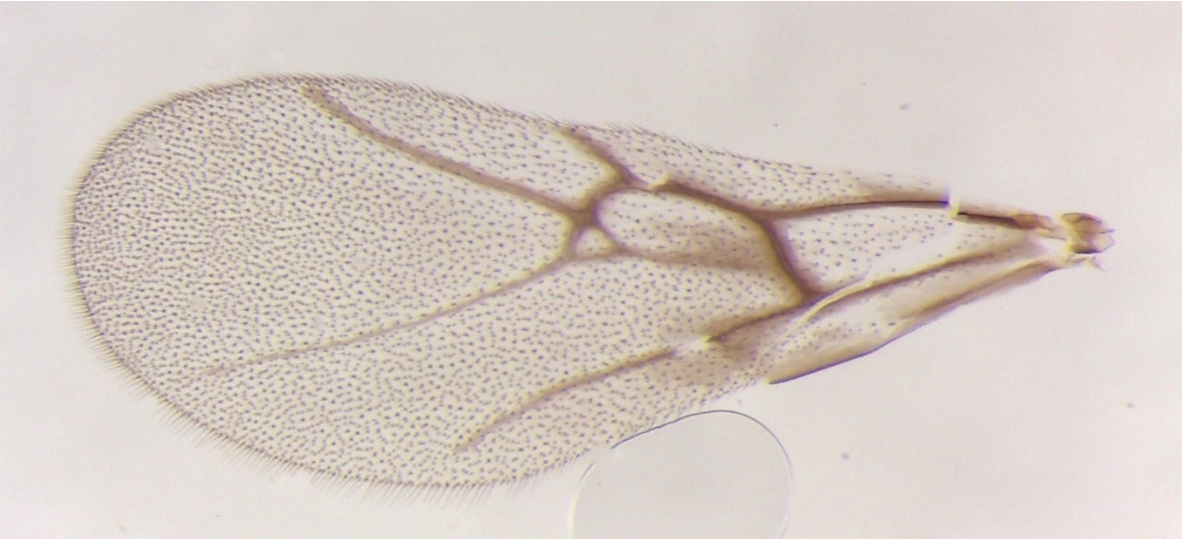

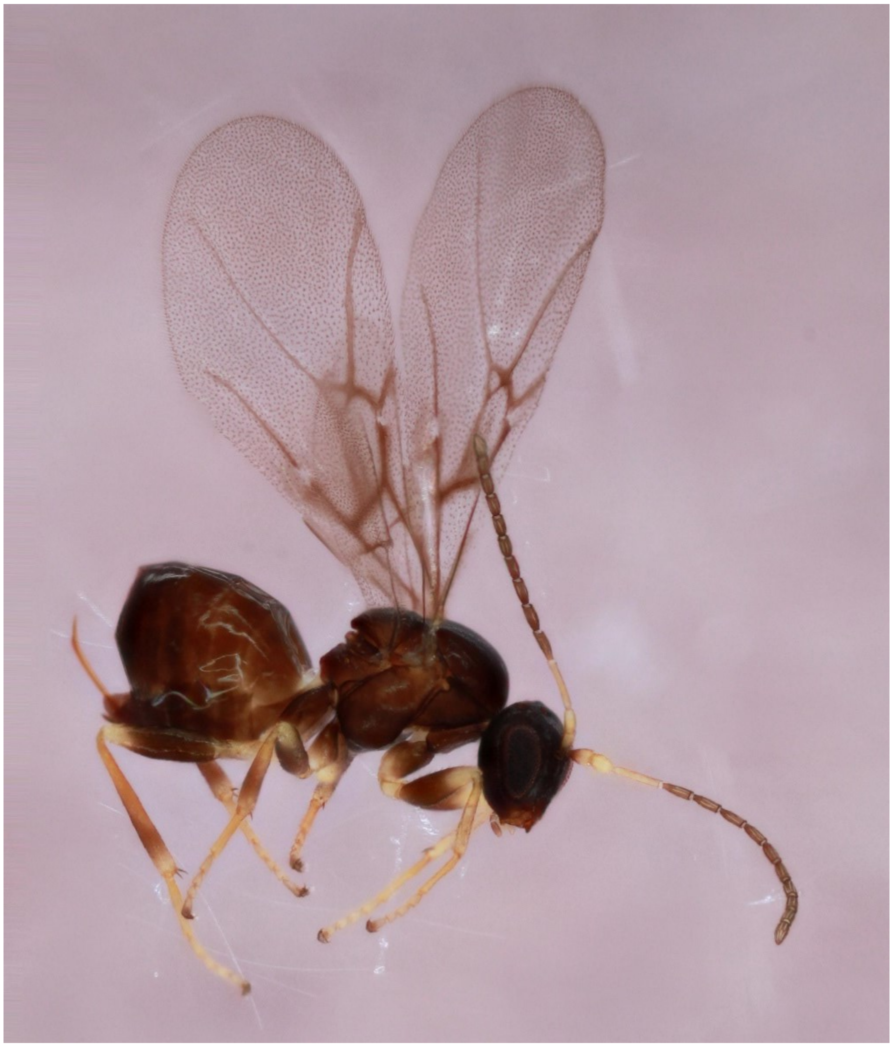

*Philonix nigra* – 630-023-7

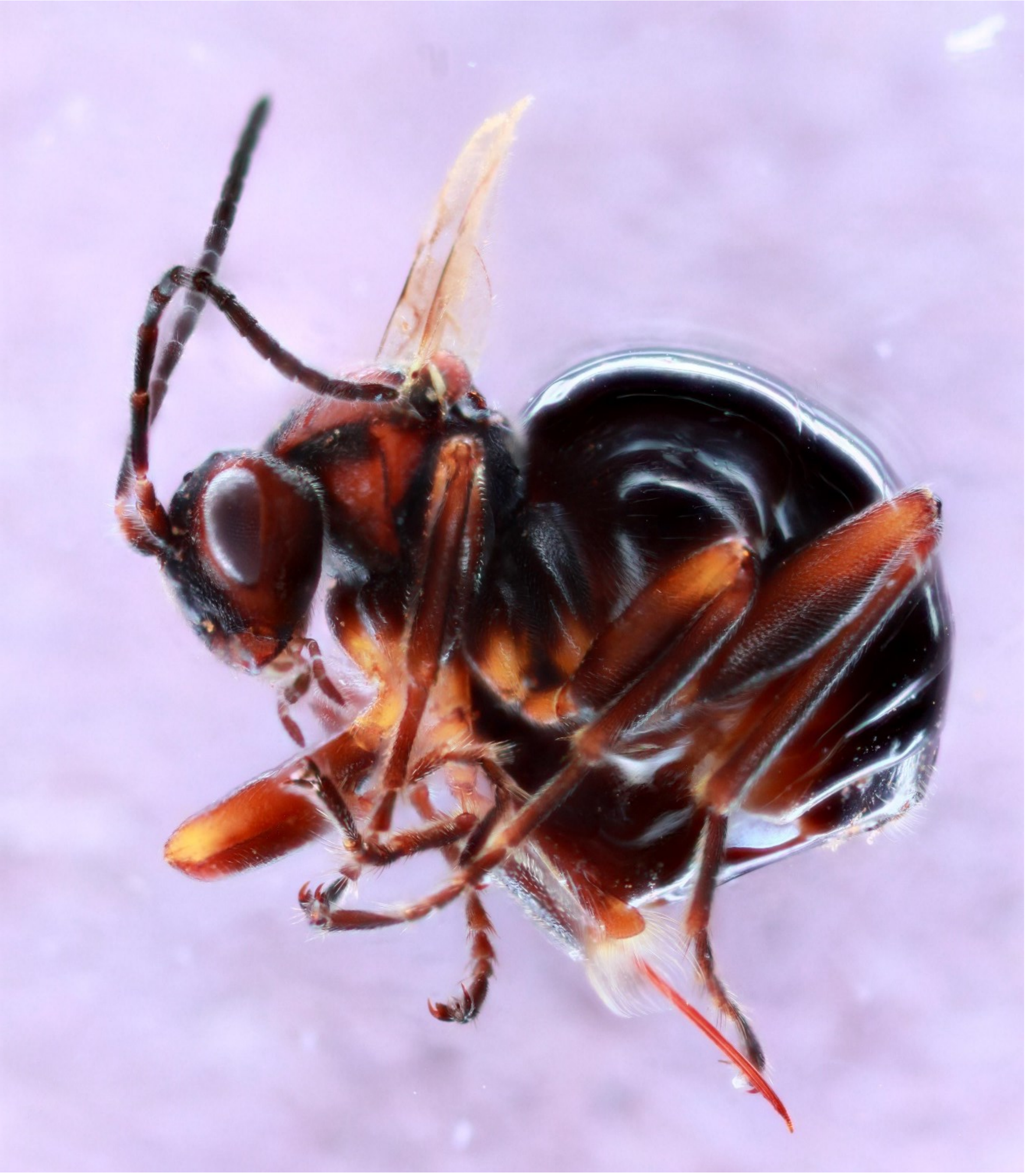

Unknown bud gall on *Quercus alba* – 1355-095-2

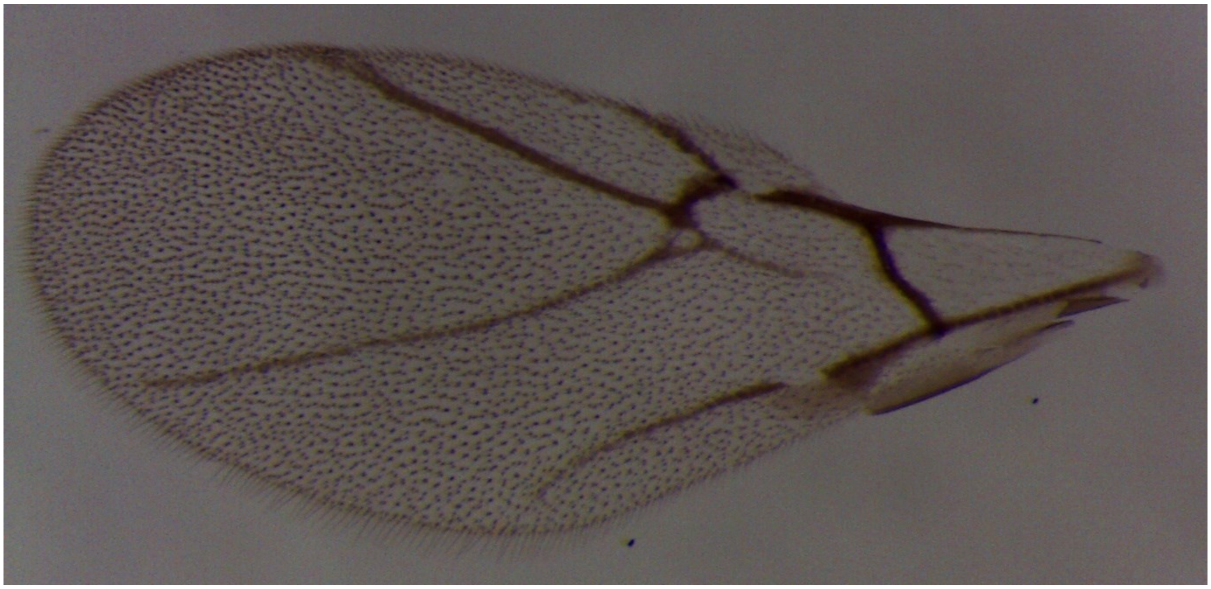

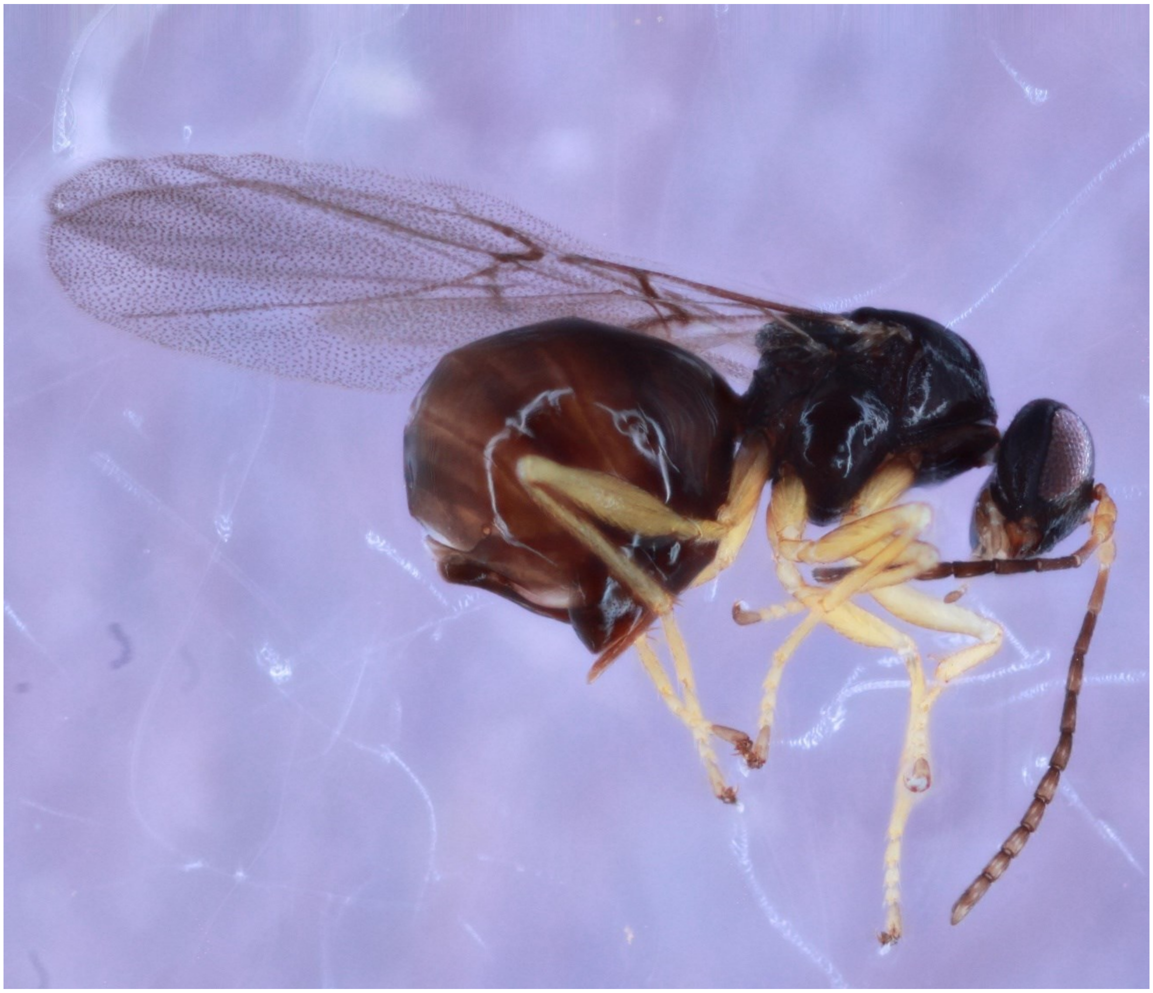

Unknown integral gall on *Quercus imbricariae* – 1291-045-1

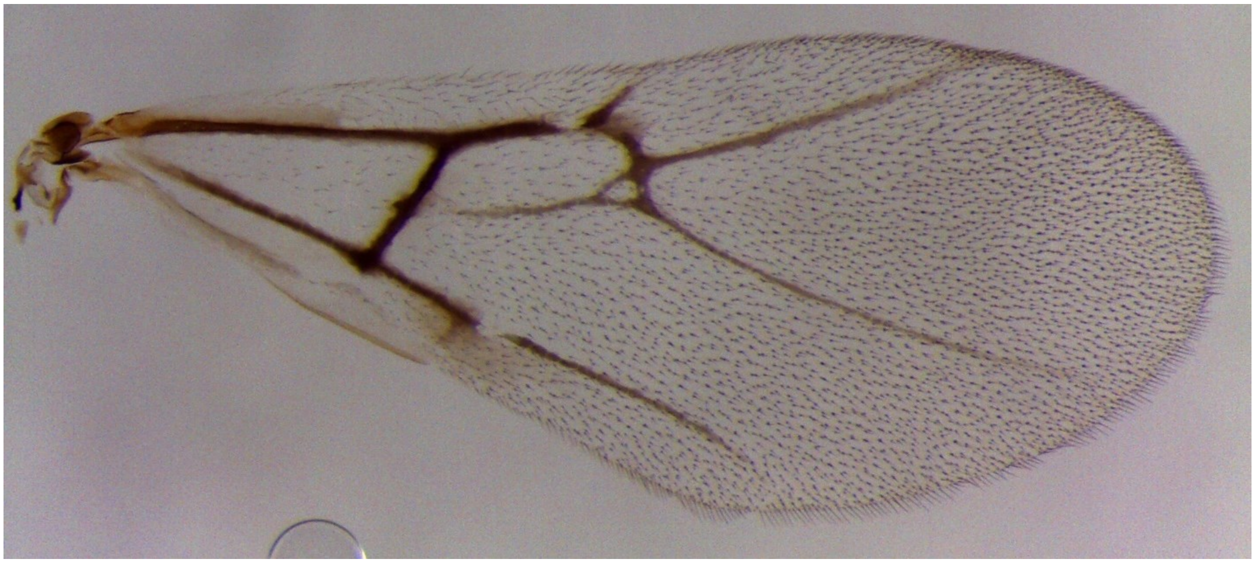

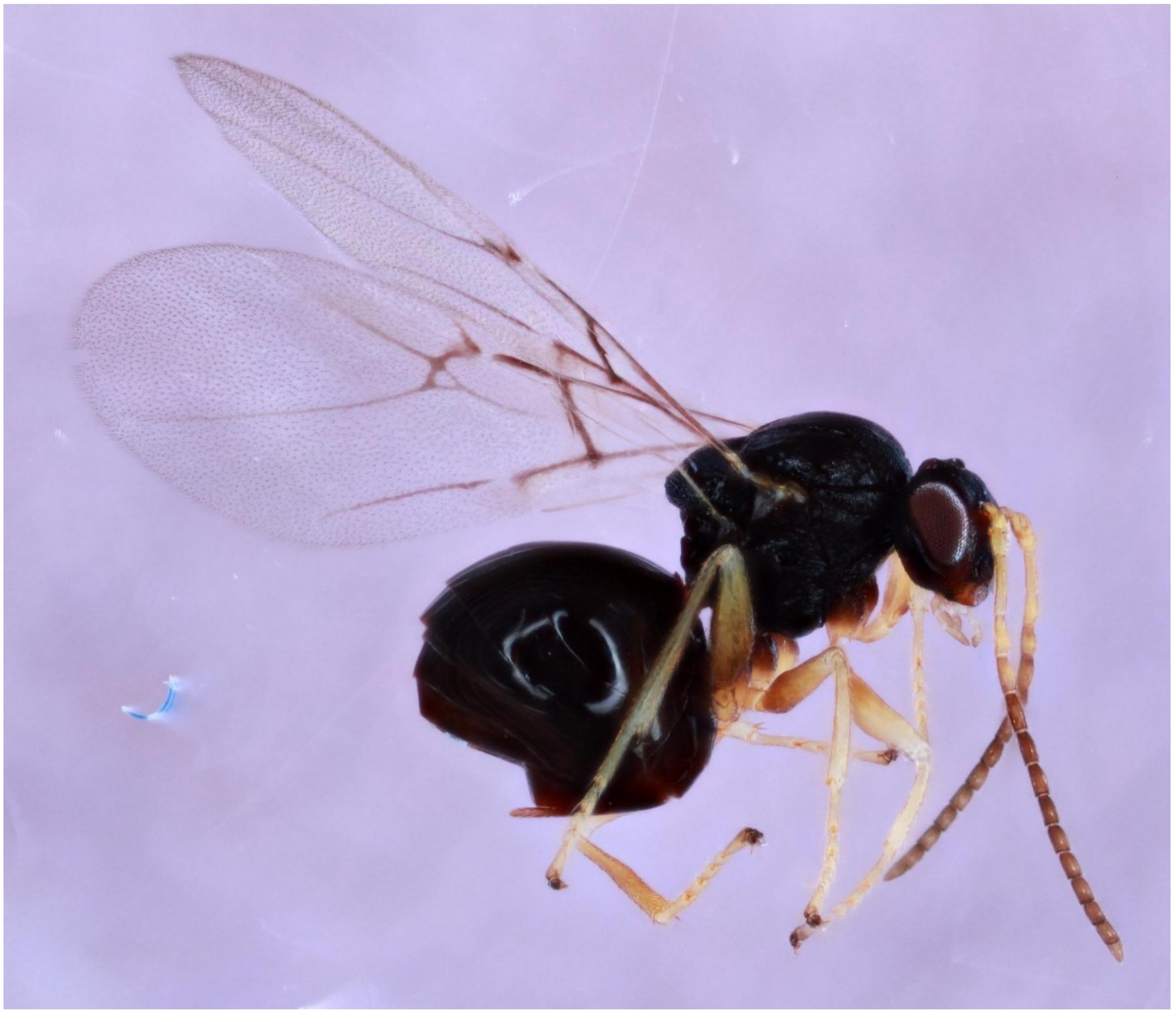

Unknown stem gall on *Quercus grisea* – 1686-217-2A

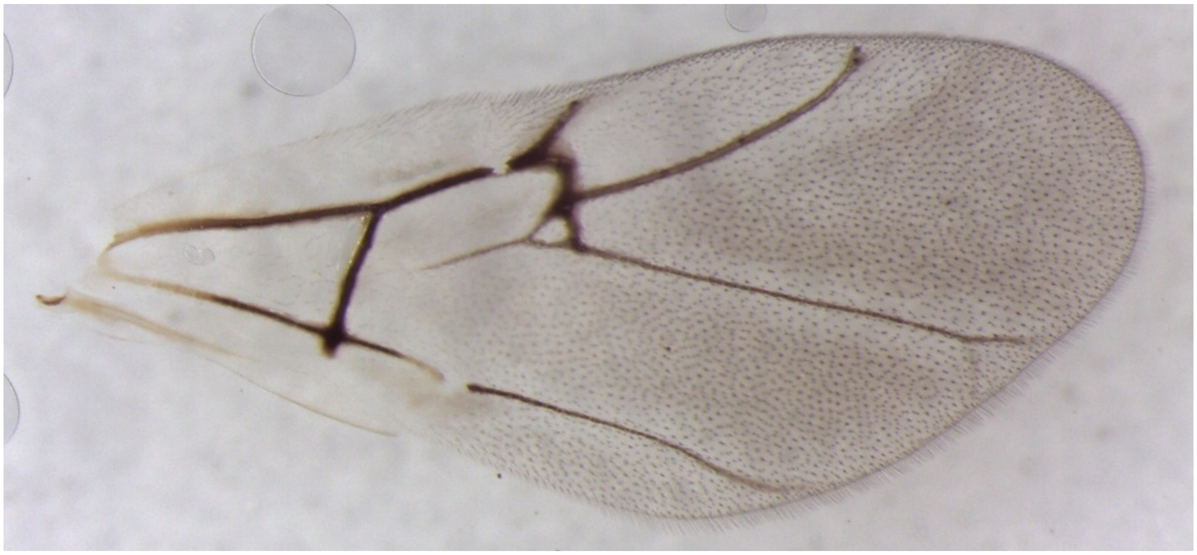

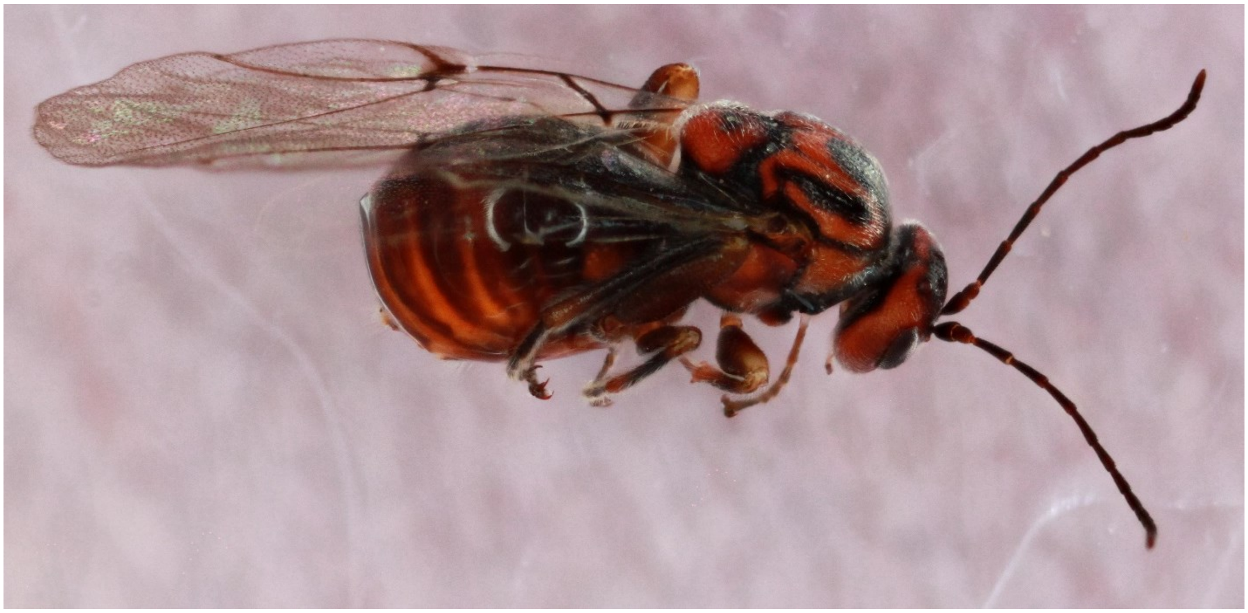

*Xanthoteras eburneum* – 1664-199-2A

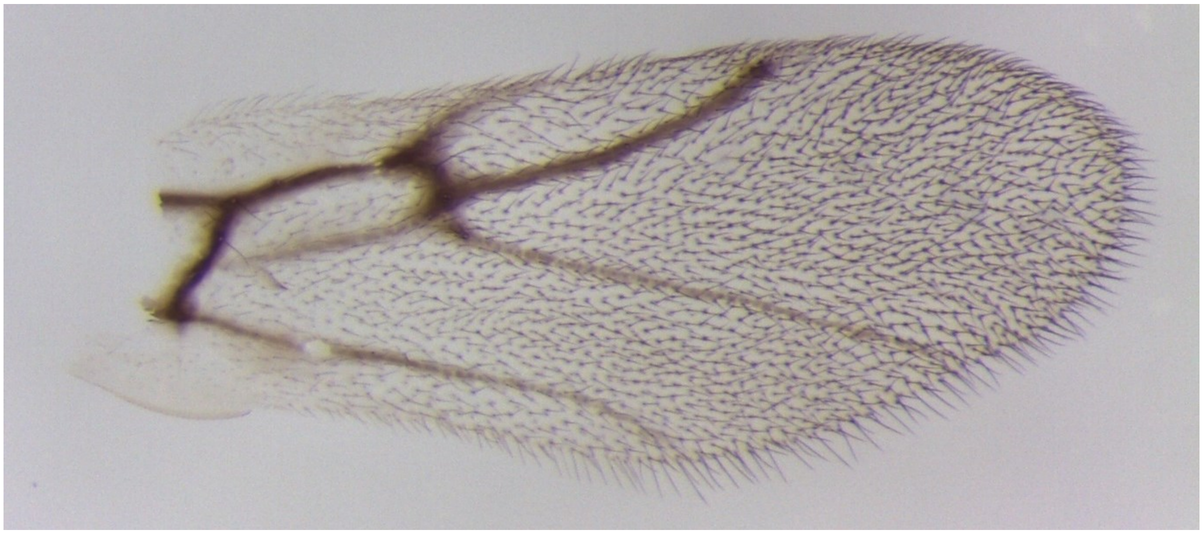

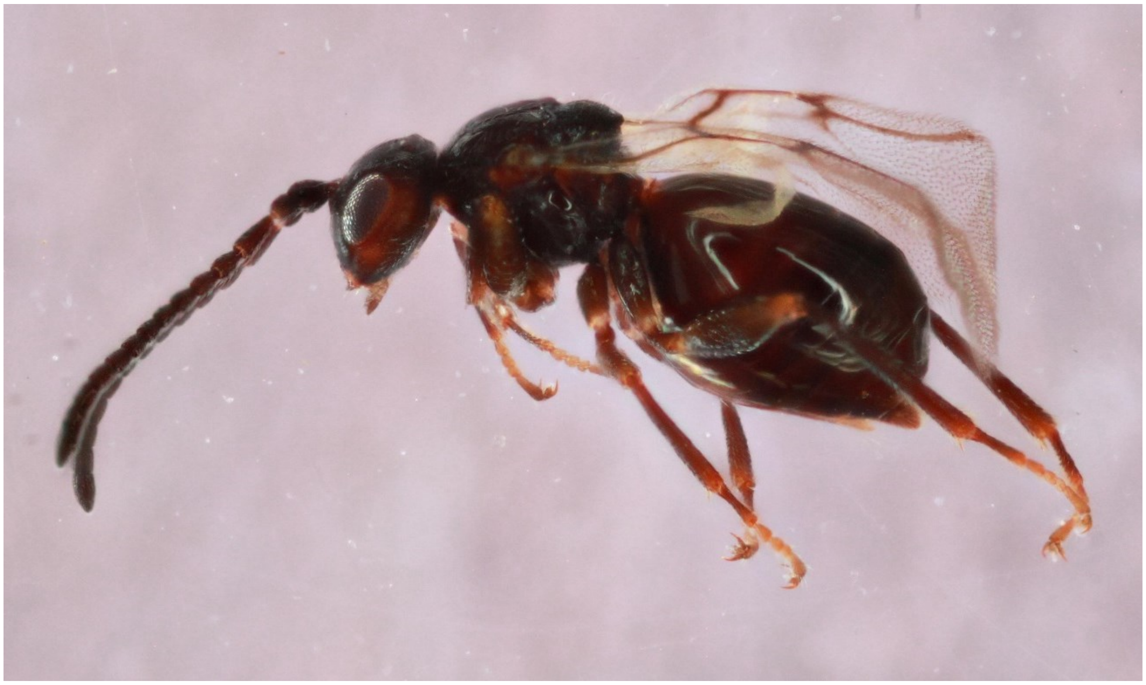

*Zopheroteras sphaerula* – 668-062-2

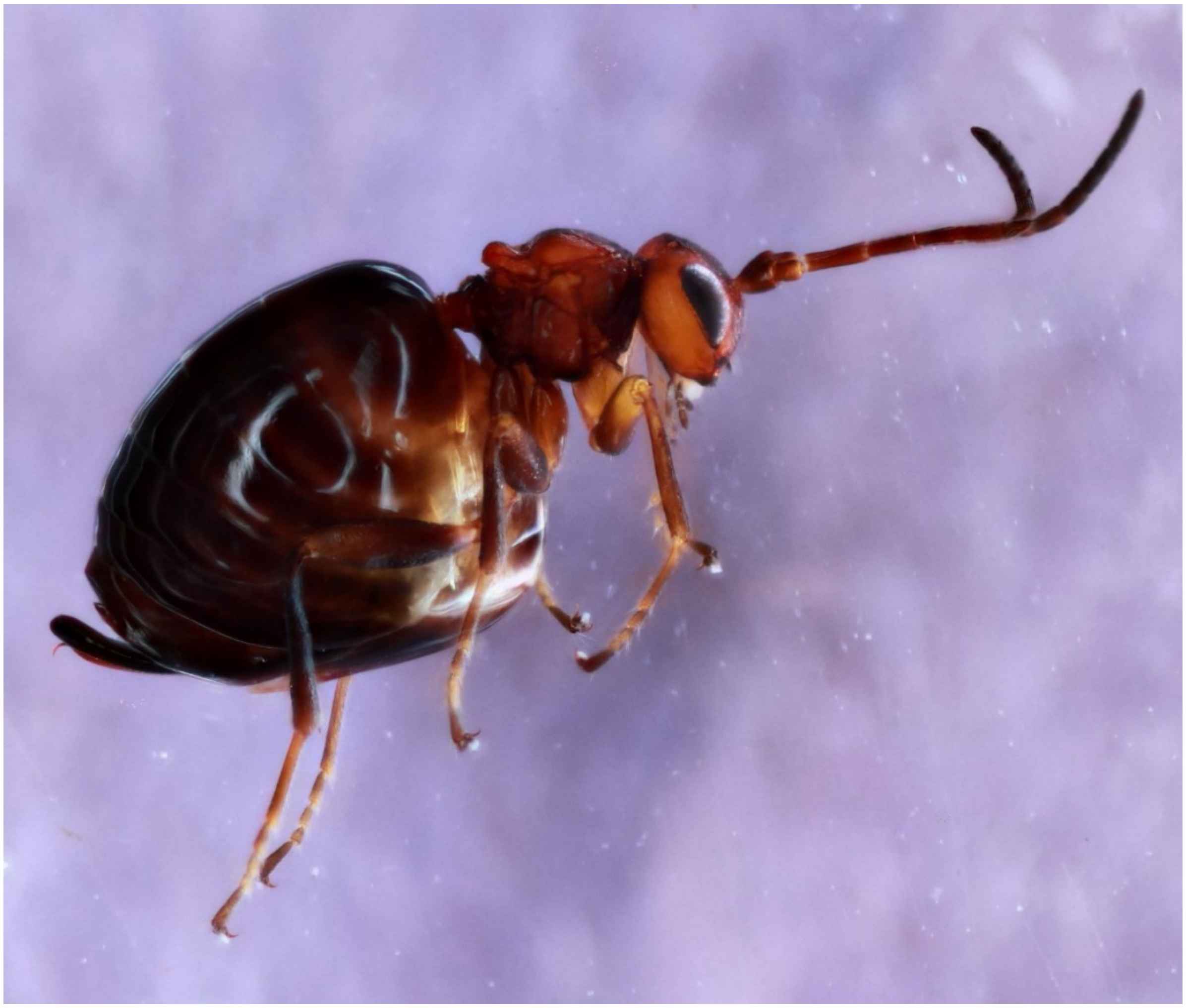
